# Cryptic genetic variation shapes the fate of gene duplicates in a protein interaction network

**DOI:** 10.1101/2024.02.23.581840

**Authors:** Soham Dibyachintan, Alexandre K Dube, David Bradley, Pascale Lemieux, Ugo Dionne, Christian R Landry

## Abstract

Paralogous genes are often redundant for long periods of time before they diverge in function. While their functions are preserved, paralogous proteins can accumulate mutations that, through epistasis, could impact their fate in the future. By quantifying the impact of all single-amino acid substitutions on the binding of two myosin proteins to their interaction partners, we find that the future evolution of these proteins is highly contingent on their regulatory divergence and the mutations that have silently accumulated in their protein binding domains. Differences in the promoter strength of the two paralogs amplify the impact of mutations on binding in the lowly expressed one. While some mutations would be sufficient to non-functionalize one paralog, they would have minimal impact on the other. Our results reveal how functionally equivalent protein domains could be destined to specific fates by regulatory and cryptic coding sequence changes that currently have little to no functional impact.

## Introduction

Proteins evolve in a vast sequence space^1^, only a tiny fraction of which is explored during evolution. This space encompasses mutations with varying effects. Even proteins under purifying selection meander in this sequence space^2^, provided their function is maintained. Although protein function is preserved during this meandering, exploration in sequence space may determine a protein’s ability to acquire novel functions in the future^3, 4^. One context in which such sequence space exploration becomes particularly important is following gene duplication.

Gene duplication generates redundancy, which relaxes the constraints imposed by purifying selection on one or both of the two copies. As a result, most duplicates accumulate loss of function mutations and revert to a single copy state, a phenomenon called pseudogenisation^5^. Among the preserved duplicates, adaptive evolution in one copy can lead to the acquisition of novel functions^6, 7^. In contrast, degenerative mutations in both duplicates^8^ can result in duplicates partitioning their ancestral functions^9^, sometimes creating dependency^10^. Although duplicates undergoing pseudogenisation and dependency have been observed in many studies, it has been estimated that approximately 30% of retained duplicates maintain a high degree of functional redundancy^11–13^. The consequences of this redundancy for the future evolution of paralogous genes are poorly understood and rarely studied.

Epistasis determines the evolutionary accessibility of a mutation by making it contingent on the genetic background in which it is introduced^14, 15^. Protein sequences can diverge over time by accumulating cryptic variation, which does not contribute to any heritable phenotypic variation but can dictate the evolvability of these proteins in the future^16–18^. Cryptic variation and epistasis, therefore, give functionally equivalent proteins access to different neutral and adaptive mutations^19–21^, which together determine the extent of evolvability of a protein. One demonstration of the impact of epistasis is that many substitutions are incompatible between orthologous proteins performing similar functions^22, 23^, even when their coding genes functionally complement between species^24^. Studies that have primarily examined orthologous proteins that evolved to perform the same function in different cellular contexts^25, 26^ suggest that proteins with similar functions could, early after speciation, be imprisoned into different areas of the sequence space compatible with their molecular functions. Similarly, functionally redundant paralogs^13^ might have accumulated such cryptic divergence. Thus, mutational paths that are evolutionarily accessible to one paralog may not be accessible to the other. The impact of such cryptic divergence between paralogs on their evolvability is largely unknown.

To assess the potential impact of cryptic divergence on paralogous protein evolution, we examined eukaryotic proteins which contain peptide recognition modules (PRMs). Specifically, our model gene pair contains Src-Homology 3 (SH3) domains. These domains mediate transient protein-protein interactions that localize cellular components and regulate pathways^27–29^. SH3 domains are numerous in eukaryotic proteomes due to multiple gene duplication events^30^. Their differences in sequence largely determine the binding of these domains to specific motifs on other proteins as they bind to distinct peptides when studied in isolation *in vitro*^31, 32^.

Our experimental system comprises a pair of duplicated motor proteins involved in endocytosis (Myo3 and Myo5), called type-1 myosins, in the model organism *Saccharomyces cerevisiae*. These paralogous genes originated from an ancient whole genome duplication event in ascomycetes^33^. Type-I myosins are crucial for cellular viability, with their C-terminal SH3 domains playing critical roles in protein localization and actin polymerization^34–38^. In fungi that diverged before the whole genome duplication, the Myo3/Myo5 single ortholog is either essential or leads to severe growth defects upon deletion^39–41^. A single deletion has no measurable fitness defects in *S. cerevisiae.* In contrast, a double deletion of the two duplicates leads to synthetic lethality^29, 42, 43^, which makes Myo3/Myo5 an excellent model system for understanding sequence space exploration for redundant paralogs and their evolutionary trajectories. Recent research has demonstrated that the SH3 domains in these duplicates are structurally similar (RMSD = 0.386) and functionally equivalent as they can be interchanged between the two paralogs without detectable effects on the binding of Myo3 or Myo5 to cognate interaction partners^44^. The resurrection of the ancestral domain before the gene duplication event showed that the domain has remained functionally unchanged in either paralog since duplication despite a divergence of 10% of sites (6/59 residues) from the ancestral sequence on either paralog^44^. These diverged sites represent potential cryptic divergence that could shape the future evolution of these proteins in the protein network.

Leveraging this model gene pair, we aimed to determine whether cryptic divergence between the duplicates has accumulated, thereby constraining or facilitating their evolution in the future (Fig 1A). The impact of cryptic divergence on future evolution would manifest as differences in the impact of the same mutations in the two paralogous domains on binding to their partners. Although the two domains have preserved redundant roles so far, the few amino acid substitutions they have accumulated may determine how they will functionally diverge in the future. We assess this by introducing all possible single amino acid substitutions in the two paralogous SH3 domains using saturation mutagenesis and CRISPR-Cas9-based homology repair (Fig 1B). We inserted complete single amino acid mutant libraries directly into the yeast genome^29, 45, 46^ and quantified the functional impact of these mutations on their ability to interact with eight representative cognate partners in the protein interaction network regulating yeast endocytosis (Fig 1C). To assess the impact of regulatory divergence between the two genes^47^ and changes in sequence context in the rest of the protein^29, 48, 49^, we measured the same effects of all mutations by swapping the domains between the paralogous myosins (Fig 1B), that is, expressing a Myo5 protein carrying the Myo3 SH3 domain and vice versa. Our assays reveal that the two domains have accumulated cryptic divergence that shapes how mutations affect their binding to partners in a way that is specific to the identity of the binding partners, allowing the duplicates to partition their function. Our results also show that the divergence in other parts of Myo3 and Myo5 and their regulation influence the impact of mutations on binding. The future evolution of gene duplicates, even when redundant, is therefore highly contingent on many factors shaping the architecture of cellular networks.

**Figure 1:**
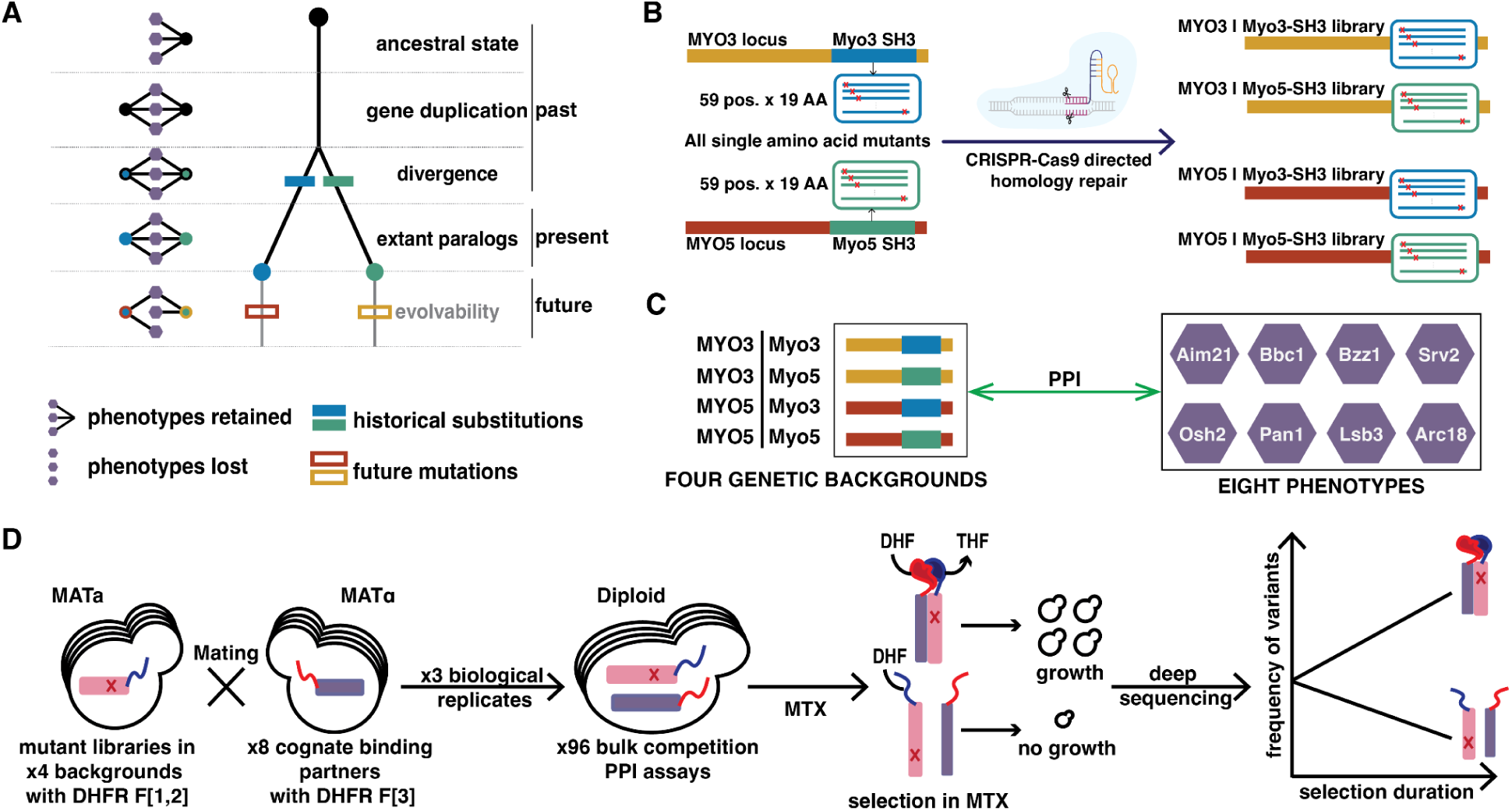
Experimental design to study evolution following gene duplication. **(A)** Schematic of long-term evolutionary trajectories involving multiple binding phenotypes following gene duplication. A gene duplicates and accumulates mutations in both copies while they maintain their ancestral phenotypes up to the present day. In the future, how evolvable each of the duplicates is contingent on the mutations historically fixed, with the same mutation having varying effects on the ability of a duplicate to maintain its ancestral phenotypes. **(B)** Saturation mutagenesis strategy of paralogous SH3 domains followed by genomic integration by CRISPR-Cas9 directed homology repair at their native loci (Myo3 SH3 in *MYO3* and Myo5 SH3 in *MYO5*) and swapped paralogous loci (Myo3 SH3 in *MYO5* and Myo5 SH3 in *MYO3*). **(C)** The experimental model system contains two mutated SH3 domains integrated into their native genomic loci and swapped paralogous loci (for four backgrounds) and eight protein-protein interaction phenotypes. The impact of each mutation is assayed for the binding phenotype to 8 cognate partners in three replicates. **(D)** The dihydrofolate reductase protein-fragment complementation assay (DHFR PCA) is a growth-based selection assay for measuring the effect of mutations on the protein-protein interaction phenotypes. Mutated genes are tagged with the DHFR F[1, 2] fragment in the MAT**a** mating type, and cognate interaction partners are tagged with the DHFR F[3] fragment in the MAT**α** mating type. These two strains are mated in three biological replicates to generate libraries for performing the growth-based phenotypic selection in the presence of methotrexate (MTX), an inhibitor of yeast DHFR. The tagged DHFR fragments reconstitute a mutated murine DHFR resistant to MTX. Hence, the reconstituted complex is essential for growth in MTX. Samples from before and after selection are sequenced on Illumina Novaseq to quantify the frequency of mutants and calculate a functional score.

## Results and Discussion

### Quantifying the functional effects of mutations

Myo3 and Myo5 share the same arrangement of protein domains: an N-terminal myosin motor domain, a neck containing two IQ motifs, and a C-terminal tail consisting of a TH1 domain, an SH3-domain and a CA domain. The SH3 domains are short (59 amino acids long) and dictate binding to multiple partners^29, 44^, making them excellent candidates for genetic manipulation.

We created complete variant libraries for the two SH3 domains (Fig 1B). First, we inserted the variant libraries at their native loci (Myo3 SH3 library at the *MYO3* locus and the Myo5 SH3 library at the *MYO5* locus) using CRISPR-Cas9 mediated homology-directed repair^29, 45, 50^. We also introduced the Myo3 SH3 library at the *MYO5* locus and the Myo5 SH3 library at the *MYO3* locus. This way, we could also directly compare the impact of mutations on the two paralogous domains when expressed from the same locus.

After successfully integrating the variants into the yeast genome, we performed bulk competition assays to estimate the effect of mutations on protein-protein interactions with known functional binding partners involved in endocytosis, vesicle trafficking and actin cytoskeleton organization. Each interaction partner reports on a specific binding phenotype (eight) in this network (Fig 1C). The assay we used is a dihydrofolate reductase protein-fragment complementation assay (DHFR-PCA)^51^ in bulk competition^29, 45^, which quantitatively measures the amount of protein complex formed between two interaction partners by allowing strains bearing variants which bind strongly to grow faster in the selection media (Fig 1D). We measured the frequency of a variant in the competition assay before and after selection by deep sequencing. We defined the effect of a mutation (F) as the natural logarithm of the ratio between the variant frequency after and before selection with reference to the wild-type protein sequence (Supplementary File 2).

We observed that five phenotypes have the same dynamic range of measurements across all four combinations of SH3 library and locus, and the remaining phenotypes had the same dynamic range for either the *MYO5* or *MYO3* locus (Fig Suppl 1). We scaled the scores between 0 and 1 using the median of synonymous wild-type variants and nonsense variants for every experiment, and we refer to this as the functional score of a mutation (ΔF) (Supplementary File 3,5).

Validation experiments and computational analyses strongly support the reliability of the functional scores. The three biological replicates show high reproducibility (median Pearson’s r = 0.95 [0.91-0.97]) (Fig Suppl 2). In addition, we independently reconstructed 60 mutants (15 in each genetic background) with a range of effects. We assayed the effect of these mutations in single cultures on three phenotypes in three biological replicates. We obtained a high correlation with the results from our bulk competition experiments (median Pearson’s r = 0.93 [0.85-0.93]) (Fig Suppl 3A). For further validation, we took datasets from the published literature on functional mutagenesis studies of homologous SH3 domains^29, 52^, and our data correlates well with these datasets (median Pearson’s r = 0.61 [0.5-0.65] (Fig Suppl 4A, B). Moreover, our experiments recapitulated the loss of function mutations at key binding interface residues (positions 7, 9, 36, 49 and 54 on the SH3 domain) from these studies. Our experiments correlate strongly with folding stability measurements of mutants (ΔG)^53^ in both the Myo3 (Pearson’s r = −0.71) and Myo5 (Pearson’s r = −0.65) SH3 domains (Fig Suppl 5A, D), with destabilizing mutants being recapitulated very well. We also used evolutionary models, which infer the effect of mutations from coevolution between sites across natural protein sequences, and our data correlate very well with both Myo3 and Myo5 GEMME models^54^ (Pearson’s r = 0.73 [0.69-0.74]) (Fig Suppl 4C). Our assays, therefore, capture the functional impact of mutations in a highly reproducible manner.

### Cryptic variation preserves function but alters evolvability

Next, we looked into how cryptic divergence affects the response to mutation of different binding phenotypes. We focused on the five phenotypes (binding to Aim21, Arc18, Bbc1, Osh2 and Pan1) with the same dynamic range of measurements across genetic backgrounds. We hierarchically clustered correlation coefficients of functional scores of mutations for all four locus-domain combinations for the five phenotypes. Five major clusters emerged, grouping based on the locus and SH3 domain instead of the phenotypes for four of these clusters (Fig 2A, 2B, Fig Suppl 6). However, the Osh2 phenotypes were the exception as they formed a separate fifth cluster with the SH3 domains forming sub-groups (Fig Suppl 6). The structure of the four major groups where four phenotypes (binding to Bbc1, Aim21, Arc18, Pan1) clustered together was the same for each domain-locus combination (Fig 2A, 2B, Fig Suppl 6). These results suggest that cryptic divergence can alter the effect of mutations for all phenotypes we assessed but to different extents.

**Figure 2:**
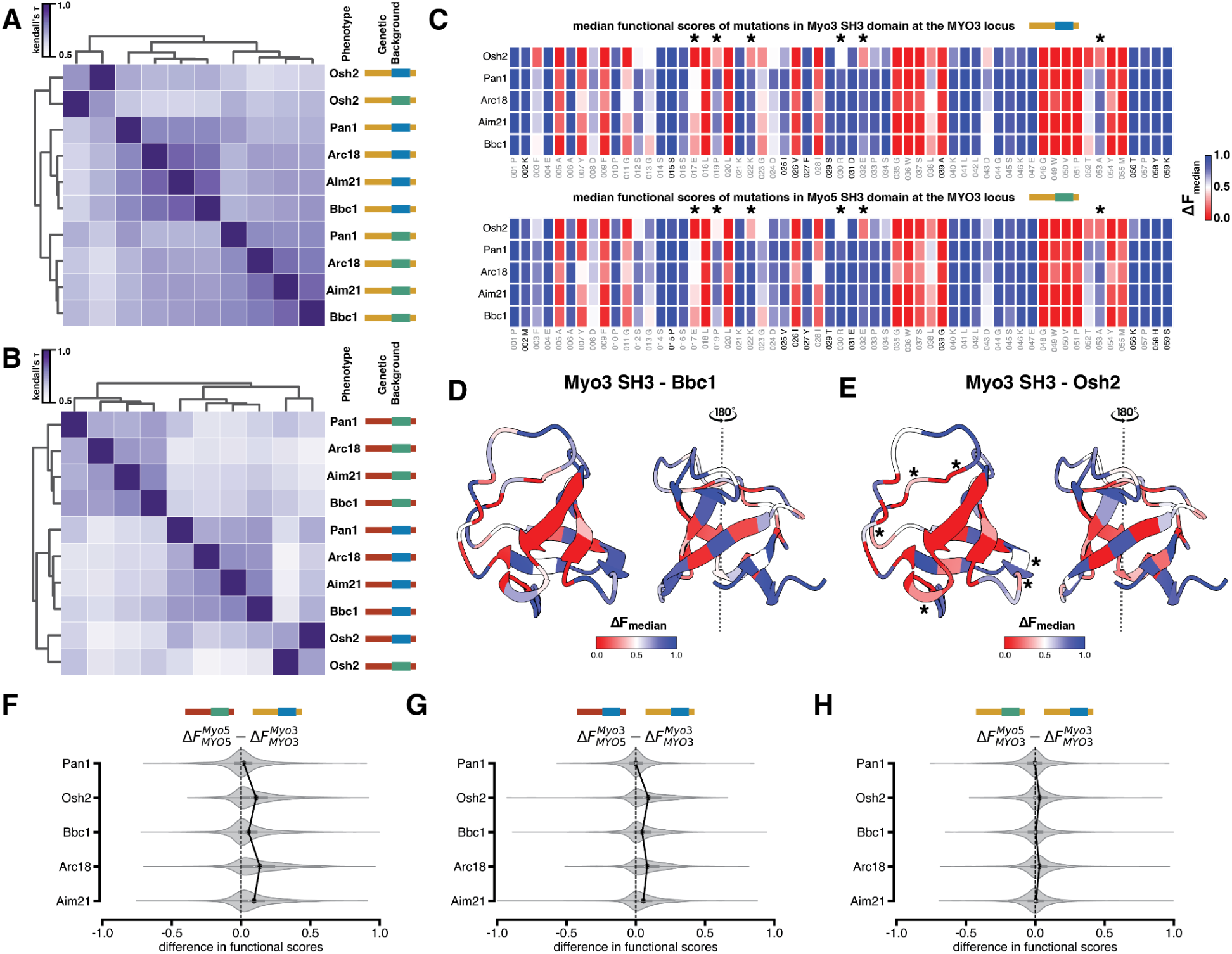
Cryptic variation and regulatory divergence shape the binding evolvability of gene duplicates. **(A)** Clustering of correlation of functional scores for five phenotypes at the MYO3 locus for mutations introduced in either SH3 domain. **(B)** Clustering of correlation of functional scores for five phenotypes at the MYO5 locus for mutations introduced in either SH3 domain. **(C)** The median functional score of mutations at a given position where 1 represents the function of the wild-type protein and 0 represents a total loss of function for five different phenotypes at the *MYO3* locus for the Myo3 SH3 domain (top) and Myo5 SH3 domain (bottom). Divergent amino acid positions are marked in bold. **(D)** Median functional scores at each site of the Myo3 SH3 domain expressed from the *MYO3* locus for the Bbc1 phenotype mapped onto the Myo3 SH3 domain structure (PDB: 1RUW). **(E)** Median functional scores at each site of the Myo3 SH3 domain expressed from the *MYO3* locus for the Osh2 phenotype mapped onto the Myo3 SH3 domain structure (PDB: 1RUW). * represent the sites where median functional scores significantly differ between Bbc1 and Osh2 phenotype (two-sided Mann-Whitney U test, Benjamini-Hochberg adj. p value < 0.01). **(F)** The difference in scores between the same mutations introduced in the two paralogous domains at their native loci. The solid black line joins the mean difference in functional scores for different binding phenotypes. Cohen’s d: Pan1=0.14; Osh2=0.64; Bbc1=0.29; Arc18=0.75; Aim21=0.57. **(G)** The difference in functional scores between the same mutations introduced in the Myo3 SH3 domain introduced at the higher expressed locus (*MYO5*) and the lower expressed locus (*MYO3*). The solid black line joins the mean difference in functional scores for different phenotypes. Cohen’s d: Pan1=0.01; Osh2=0.62; Bbc1=0.37; Arc18=0.62; Aim21=0.41. **(H)** Difference in functional scores between the same mutations introduced in the Myo5 SH3 domain and the Myo3 SH3 domain when expressed from the same genetic locus. The solid black line joins the mean difference in functional scores for different phenotypes. Cohen’s d: Pan1=0.02; Osh2=0.24; Bbc1=0.03; Arc18=0.25; Aim21=0.05.

Previous studies have experimentally identified the intrinsically disordered regions (IDRs) through which Bbc1, Pan1 and Osh2 bind to type-1 myosin SH3 domains^35, 36, 38^, while other binding partners have been less studied. Although there is variation in the exact binding motif sequence, the sequences in these IDRs satisfy the consensus polyproline binding motifs (see Methods for details) identified for both domains^31^. Hence, it was surprising that a single phenotype, i.e. Osh2 (APKHAPPPVP), clustered separately when the binding motif was highly similar to those of Bbc1 (MPNTAPPLPR) and Pan1 (PPAGIPPPPP). To explain this, we decided to look at how the average effect of a mutation varied between phenotypes for any domain-locus combination. We observed 18 positions highly intolerant to most mutations for all phenotypes in both paralogous domains (Fig 2C). However, specifically for Osh2, positions 17, 19, 22, 32, 52 and 54 were also intolerant to most mutations when introduced in both SH3 domains (Fig 2C) at both loci (Fig Suppl 7). The interaction partners, therefore, modulate the impact of mutations in the domains. When we mapped the median functional scores for Bbc1 (Fig 2D) and Osh2 (Fig 2E) to the Myo3 SH3 domain structure (PDB: 1RUW), all these additional constrained sites mapped to the surface of the domain. Moreover, mutations at these positions had a weaker effect on folding stability than those that caused loss of function for all phenotypes (Fig Suppl 8). These additional constraints imposed on the evolvability of both domains explain the separate clustering of the functional scores for the Osh2 phenotype.

These analyses reveal that although the binding motifs satisfy the consensus SH3 binding peptide profile^31^, the binding partners in the protein-protein interaction network are largely independent phenotypes that can be affected disparately by mutations in the paralogs. This suggests that variation in the motifs among the binding partners may not impact the binding specificity but still influence evolvability. Amino acid substitutions that impair binding to all phenotypes would eventually lead to the pseudogenisation of one duplicate, with the organism returning to a single copy state (Fig 2C, Fig Suppl 17 B). Meanwhile, the substitutions that specifically impair binding to Osh2 or Bbc1 can set up conditions for the duplicates to partition their functions and ultimately subfunctionalize (Fig 2C, Suppl Fig 17 C).

### Regulatory divergence biases binding evolvability

Following gene duplication, many pairs of duplicates undergo regulatory divergence over time, resulting in the two copies being expressed at different levels *in vivo*^55–57^. By altering protein abundance, regulatory changes can alter the distribution of effects of protein-coding mutations on fitness, thereby affecting protein evolvability^47, 58, 59^. Similarly, as protein-protein interaction depends on protein abundance and binding affinity, regulatory divergence could also strongly impact how mutations in SH3 domains translate into changes in protein complex formation. Although type-1 myosins are largely functionally redundant^60^, *MYO5* has a slightly higher expression level than *MYO3 in vivo*^44^, which we find is at least partially caused by *MYO5* having a stronger promoter than *MYO3* (Fig Suppl 9B). Hence, we hypothesized that mutations introduced at the *MYO5* locus would cause less functional impairment than those introduced at the *MYO3* locus.

We analyzed five phenotypes with the same dynamic range across genetic backgrounds. We found, as hypothesized, that mutations caused systematically less functional impairment when introduced at the *MYO5* locus for all phenotypes but one. The sole exception is the Pan1 interaction phenotype, for which we observed a minimal bias in functional scores between loci (p<0.01, two-sided paired t-test for all phenotypes, Cohen’s d = 0.14 for Pan1, Cohen’d = [0.36−0.75] for remaining phenotypes) (Fig 2F). The same pattern was observed when functional scores were compared between the same SH3 domain integrated at the higher and lower expressed loci (p<0.01, two-sided paired t-test for all except Pan1) (Fig 2G), with the bias in favour of *MYO5* locus being medium to large for all phenotypes (Cohen’s d=0.37−0.62), except Pan1 (Cohen’s d=0.01). This bias favouring the higher expressed gene disappeared when the two SH3 domains were expressed from the same locus, with the bias in functional scores being very small to small (Cohen’s d=0.02−0.25, p>0.01, two-sided paired t-test for all except Osh2 and Arc18) (Fig 2H).

Our results show that a mutation in the same protein sequence impairs protein function to different extents based on *in vivo* expression levels. We also observed that this difference in effect is not uniform across phenotypes, which is likely a function of the binding affinity and local protein abundances underlying specific interactions. These results show that evolutionary contingency can arise in paralogous proteins due to regulatory divergence, independently of sequence divergence (Fig 2G). Although the same mutations are accessible to both paralogs following duplication, regulatory divergence makes the same mutations have different impacts on the paralogous genes, thereby altering their probability of fixation. Interestingly, divergence in expression could be neutral at first^57^ but could later dictate which mutations will be neutral or not in the coding sequences.

### Binding evolvability is contingent on cryptic divergence

We next asked whether the evolvability of these two SH3 domains is impacted by the amino acid changes they have accumulated since duplication. We compared the functional scores of the mutations in the SH3 domains expressed from the same genetic locus to control for regulatory divergence. We focused on the three phenotypes well characterized experimentally involving direct physical interactions between the type-1 myosin SH3 domains and Bbc1, Pan1 and Osh2^35, 36, 38^. First, we classified mutations as functional and non-functional, with non-functional mutations behaving similarly to proteins with the SH3 domains removed (see Methods for details). Next, we defined contingent mutations as mutations whose functional scores significantly differ between domains (at the same locus) and non-contingent mutations as two types: (1) functional mutations which do not differ significantly between domains and (2) mutations that are not functional in both domains for a given binding phenotype (Fig 3A).

**Figure 3:**
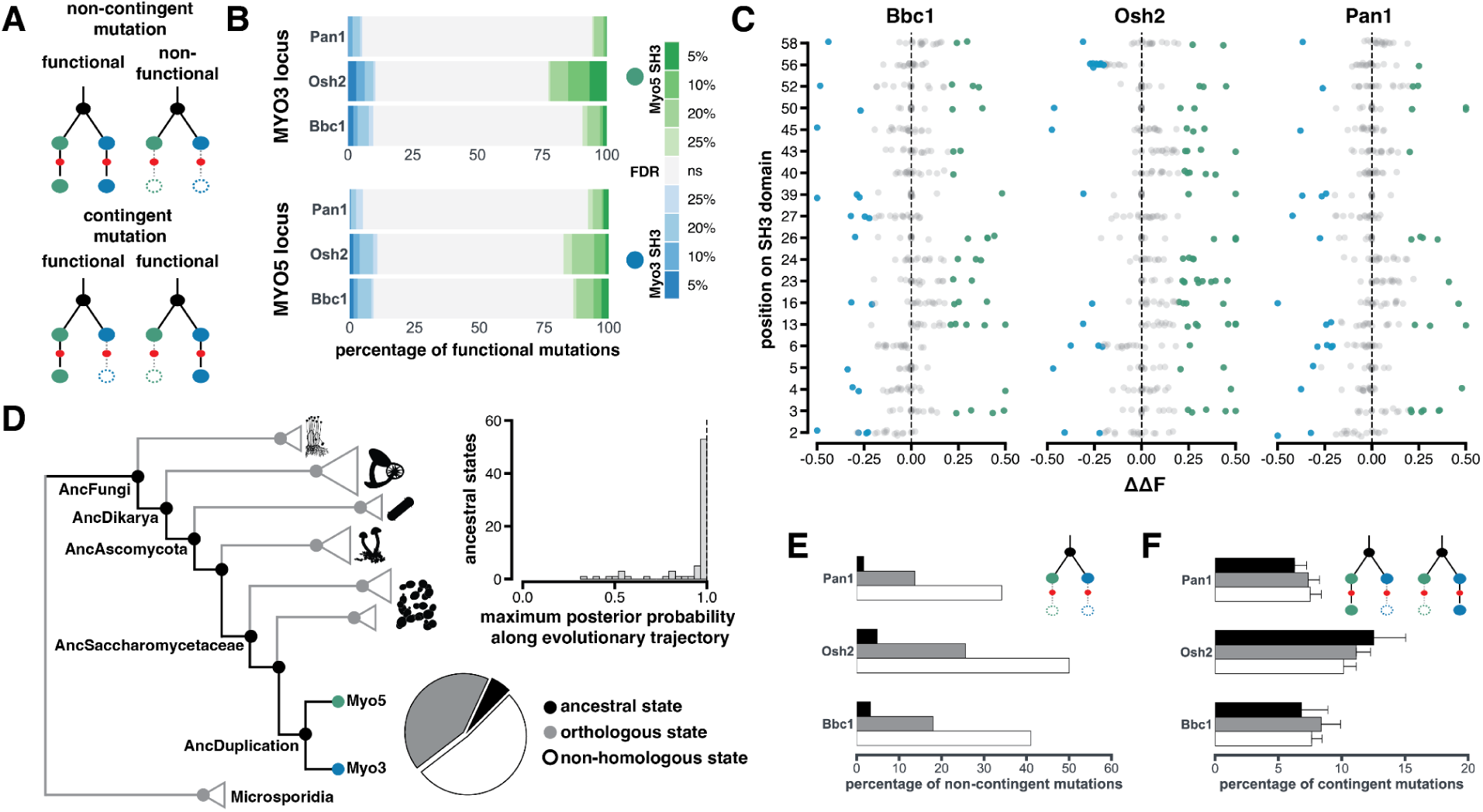
Binding evolvability is contingent on cryptic divergence. **(A)** Contingent mutations have effects on protein function that depend on the genetic background. Non-contingent mutations: mutations whose effect on protein functions is independent of the genetic background. **(B)** Percentage of significant contingent mutations among all functional mutations for all phenotypes at different False Discovery Rate thresholds (Benjamini-Hochberg correction) using two-sided Welch’s t-test. Significant mutations with a higher functional score in the Myo5 SH3 domain are shaded in green, and mutations with a higher functional score in the Myo3 SH3 domain are shaded in blue. **(C)** Contingent mutations (|ΔΔF|>0.2) for all phenotypes at sites enriched for such mutations in at least one phenotype for the Bbc1, Osh2 and Pan1 interaction. Blue dots represent contingent mutations with a higher functional score in the Myo3 SH3 domain, green dots represent contingent mutations with a higher functional score in the Myo5 SH3 domain, and gray dots represent non-contingent mutations. To simplify the illustration, the outlier ΔΔF measurements are rounded at −0.5 and 0.5. **(D)** Inferred maximum likelihood phylogenetic tree of type-1 fungal myosins. The histogram represents the maximal posterior probabilities of every ancestral state reconstructed along the evolutionary trajectory pre-duplication (represented by the path in black in the phylogenetic tree). The pie chart represents the proportions of mutations classified as ancestral states (black), non-ancestral orthologous states (gray), and non-homologous states (white) **(E)** Percentage of mutations that are non-functional in both domains in each evolutionary category at the *MYO3* locus. All pairwise comparisons are statistically significant (p < 0.05, Fisher’s exact test) **(F)** Percentage of mutations that are contingent (|ΔΔF|>0.2) in each evolutionary category at the *MYO3* locus. Standard error bars were calculated using the number of contingent mutants (|ΔΔF|>0.2) from the difference in functional scores of biological replicates between domains. All pairwise comparisons are statistically not significant (p > 0.1, Fisher’s exact test).

At the *MYO3* locus, approximately 24% of mutations for Pan1, 29% for Bbc1 and 37% for Osh2 are non-contingent as they are non-functional in both domains (Fig Suppl 10A). Similarly, at the *MYO5* locus, approximately 21% of mutations for Pan1, 25% for Bbc1 and 31% for Osh2 are non-functional (Fig Suppl 10B). We observe that the fraction of non-functional mutations in both domains is lower when the mutations are introduced into the higher expressed locus, which can likely be attributed to the higher amount of protein complex formed by certain loss of function mutants at higher abundance^47^. Among functional mutations across phenotypes in the paralogous domains, we performed a two-sided Welch t-test to identify mutations with statistically significant differences in functional scores. This allowed us to estimate the fraction of contingent mutations at different false discovery rate thresholds (Fig 3B). Almost equal percentages of contingent mutations are found in both domains, with a significantly higher functional score in one domain over the other for the Bbc1 and Pan1 phenotypes. We observed a larger percentage of mutations with a higher functional score when introduced in the Myo5 SH3 domain as opposed to the Myo3 SH3 domain for the Osh2 phenotype.

Having observed multiple statistically significant contingent mutations, we next tested whether contingent mutations were enriched at specific sites in the domain. We defined ΔΔF as the difference between the functional score of a mutation introduced in the Myo5 SH3 domain and the Myo3 SH3 domain at each locus separately to control for the locus effect. We calculated ΔΔF of mutations for all three phenotypes at both loci. However, we focused on functional scores measured at the *MYO3* locus for further analysis because measurements at this locus were more reproducible (Pearson’s r = 0.97 for all experiments) than at the *MYO5* locus (Pearson’s r = 0.95 for all experiments). We identified 19 sites on the SH3 domain that had multiple contingent mutations (|ΔΔF| > 0.2) (see Method for details) for at least one phenotype (Fig 3C). The residues at positions 2 and 58 were enriched for contingent mutations, which could arise from the proximity of amino acid changes at positions flanking the domain. However, contingent mutations at the remaining sites likely occur due to direct or distant interactions with one or more divergent residues within a domain. We see several contingent mutations at sites 6, 13, 16, 23, 26, 39 and 52 that are common across all phenotypes, most likely due to specific epistatic interactions between the introduced mutation and one or more of the 11 divergent SH3 residues. Since these mutations are present at both the core and surface of the domain and impact all binding phenotypes, the contingency in their functional scores likely arises from their effect on protein folding stability and the shared binding interface Fig 3C, Fig Suppl 17D, E). There were also several contingent mutations specifically for the Osh2 phenotype, which were enriched at positions 23, 40, 45 and 56, all of which mapped to the surface of the SH3 domain, suggesting contingency at that specific binding interface on the SH3 domain (Fig 3C, Fig Suppl Fig 17F, G).

Our analysis above considered all mutations that were introduced artificially in the domains. However, mutations compatible with functional SH3 domains, such as substitutions observed at homologous sites, are most relevant to evolution. We hypothesized that mutations that were never fixed during the evolutionary history of the protein are more likely to be non-functional for both paralogs and, hence, non-contingent. We also hypothesized that mutations sampled along the evolutionary trajectory before duplication would be less likely to be contingent than mutations sampled at random. We tested these hypotheses with a recently inferred maximum likelihood phylogeny^44^ of over 300 type-1 fungal myosins. We reconstructed the ancestral sequences for the SH3 domain at each ancestral node along the evolutionary trajectory. Along this trajectory, substitutions occurred at 47 of the 59 sites in the SH3 domain. Owing to multiple substitutions, 74 unique ancestral amino acid states existed at these sites at some point in the past, which have now been replaced by the extant amino acids of the two duplicates. Most of these 74 ancestral states were reconstructed with very high confidence (posterior probability > 0.95) in one or more ancestors along the evolutionary trajectory (Fig 3D). We further defined substitutions that occurred in the phylogenetic tree but not on the evolutionary trajectory as non-ancestral orthologous states and substitutions that did not occur in the phylogeny as non-homologous states.

As expected, we found that non-functional, non-contingent mutations were enriched in non-homologous states (∼20x) and orthologous states (∼8x) when compared to ancestral states (p < 0.05 for all pairwise comparisons, Fisher’s exact test) (Fig 3E). However, among contingent mutations (|ΔΔF| > 0.2), we did not find any statistically significant difference between the three different groups of mutations (p > 0.1 for all pairwise comparisons, Fisher’s exact test) (Fig 3F). This suggests that contingency in the effect of mutations can arise regardless of shared evolutionary history. Our findings from the phylogenetic analyses were robust to multiple |ΔΔF| thresholds and the two different loci we tested (Fig Suppl 11, 12). Overall, these results demonstrate that historical substitutions along an evolutionary trajectory are a good indicator for predicting the functionality of a mutation following duplication. In contrast, they cannot determine which mutations will become contingent following a duplication event, suggesting that idiosyncratic epistatic interactions between residues play a significant role in determining which mutations will be differentially accessible in the future.

Broadly, these results illustrate that the functional effect of certain mutations is contingent on substitutions contributing to the cryptic divergence between duplicates. Contingent mutations which impair binding to all phenotypes when they arise in one duplicate but not the other will eventually lead to pseudogenisation of that duplicate, assuming organismal fitness is not adversely affected (Fig 3C, Fig Suppl 17D, E). However, mutations that specifically impair binding to only one of the phenotypes will allow the duplicates to partition their ancestral functions and eventually subfunctionalize (Fig 3C, Fig Suppl 17 F, G). Therefore, although the variation in binding motifs allows for the partitioning of functions, cryptic divergence determines what mutations are accessible for the said partition to occur.

### Pairwise epistasis is sufficient for evolutionary contingency

Our previous results showed how cryptic divergence between duplicates can alter accessibility to mutations following duplication, which led us to test whether the substitutions that drove cryptic divergence (Fig 4A) between paralogs were contingent on each other. To test this, we synthesized chimeric variants of both SH3 domains by introducing divergent residues (single and double substitutions) from one duplicate into the other. These substitutions in chimeric variants can represent either a reversion to an ancestral state pre-duplication or a derived state post-duplication (Fig 4A). We integrated these chimeric SH3 domains at their native loci using CRISPR-Cas9 mediated homology-directed repair (Fig 4B). We tested the functional effects of these variants for the three representative binding phenotypes analyzed previously: Bbc1, Osh2 and Pan1 (Fig 4B) using the DHFR-PCA bulk competition assay. Since we observed an unimodal distribution as opposed to a bimodal distribution in our previous experiments for the distribution of log fold changes, we defined *F_C_* (the functional score of a chimeric variant) as the difference between the log fold change of a variant and the median of the log fold changes of all chimeric variants (see methods for details). The functional scores strongly correlated among three biological replicates (r = [0.95-1.0] for Myo3 phenotypes and r = [0.9-0.97]) (Fig Suppl 13).

**Figure 4:**
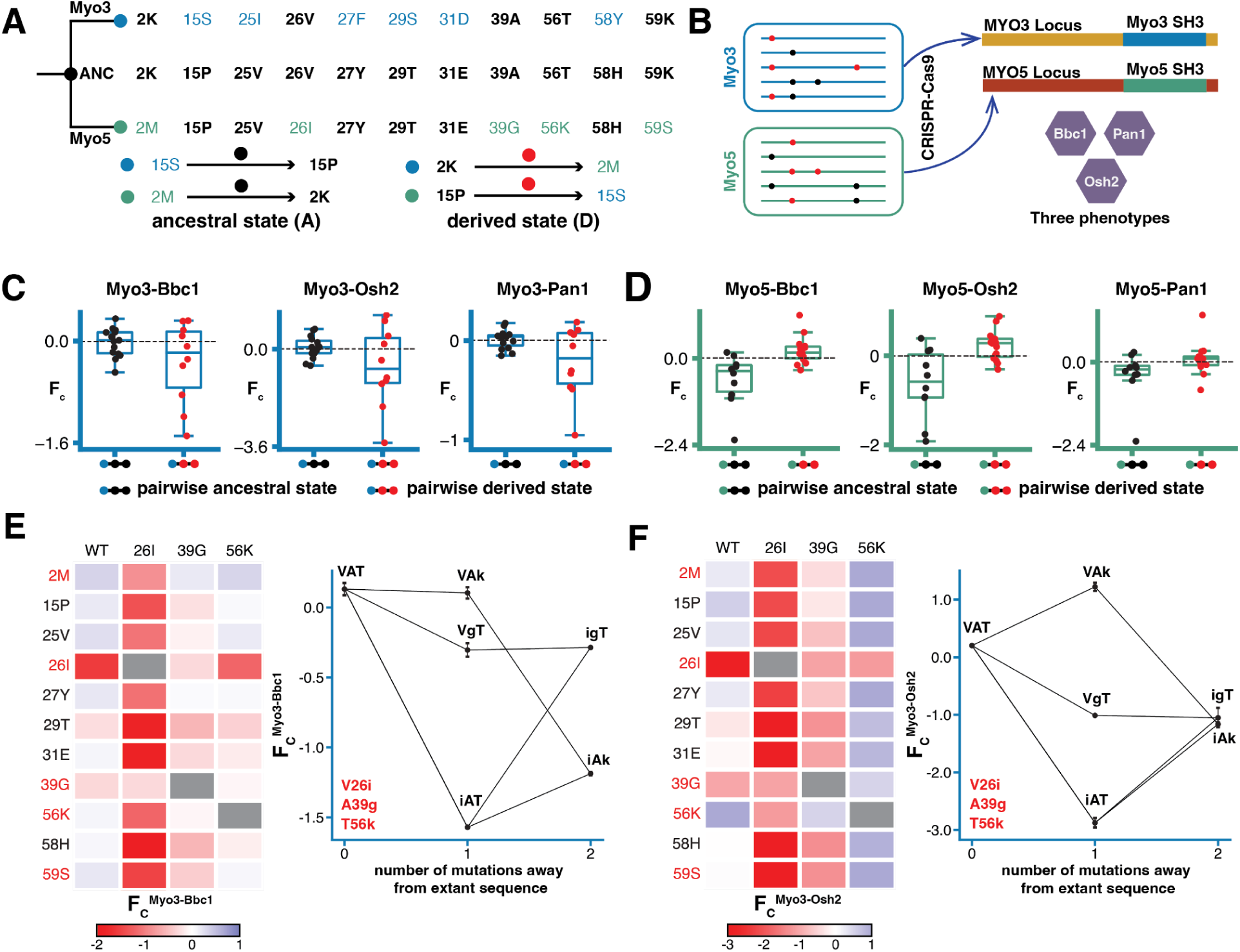
Pairwise epistasis is sufficient for evolutionary contingency. **(A)** Ancestral and divergent sequence states for the two paralogous SH3 domains with reference to their ancestor prior to duplication. **(B)** Variant synthesis strategy for the bulk competition assay CRISPR-Cas9 mediated integration of variants at native genomic loci and the phenotypes tested using DHFR PCA. **(C)** Boxplots representing functional scores for double substitutions involving pairwise ancestral or pairwise derived states for all three phenotypes in the Myo3 SH3 domain (blue axes). There is no significant difference in the median of functional scores between the two categories (p>0.05, two-sided Mann-Whitney U test) for all phenotypes. **(D)** Boxplots representing functional scores for double substitutions involving pairwise ancestral or pairwise derived states for all three phenotypes in the Myo5 SH3 domain (green axes). There is a significant difference in the median of functional scores between the two categories (p<0.05, two-sided Mann-Whitney U test) for all phenotypes. **(E)** The heatmap represents the functional score of chimeric substitutions when introduced in the wild type Myo3 SH3 sequence background, V26I background, A39G background and the T56K background for the Myo3-Bbc1 interaction. The plot on the right displays how the loss of function due to V26I substitution is rescued by the A39G substitution specifically but not by the T56K substitution. **(F)** The heatmap represents the functional score of chimeric substitutions when introduced in the wild type Myo3 SH3 sequence background, V26I background, A39G background and the T56K background for the Myo3-Osh2 interaction. The plot on the right displays how the loss of function due to V26I substitution is rescued to the same extent by both the A39G and T56K substitution, where A39G specifically restores the functionality of V26I, T56K showcases a global increase in functional score across all substitutions including V26I.

We compared the functional scores for pairwise ancestral and derived states to assess whether diverging substitutions were contingent on each other. We observed that for the Myo3 SH3 domain, most pairwise ancestral states did not cause loss of function, while a few led to minor losses of function (Fig 4C) due to interactions among three ancestral states (Fig Suppl 14A, B, C). In contrast, some pairwise derived states caused severe losses of function (Fig 4C), which is primarily driven by two derived substitutions (V26I and A39G) (Fig Suppl 15A, B, C). These observations were held across all three phenotypes (Fig 4C), suggesting structural destabilization since the residues are at the core of the SH3 domain. However, we found no significant differences in the median functional score between pairwise ancestral and derived states across phenotypes for the Myo3 SH3 domain (p-value>0.05 across phenotypes, two-sided Mann-Whitney U test). For the Myo5 SH3 domain, we observed the reverse trend, where pairwise ancestral states were linked to substantial loss of function, whereas pairwise derived states were not linked to severe losses of function for all three phenotypes (Fig 4D). However, the extent of this loss of function for the pairwise ancestral states was lower for the Pan1 phenotype, likely due to it being a stronger interaction than the other two^44^. In contrast to the Myo3 SH3 domain, we found the median functional score of pairwise derived states was significantly higher than that of pairwise ancestral states across phenotypes for the Myo5 SH3 domain (p-value<0.05 across phenotypes, two-sided Mann-Whitney U test). These results suggest that reverting to ancestral states from the Myo5 SH3 domain sequence is less probable than from the Myo3 SH3 domain sequence without causing loss of function.

To elucidate how pairwise epistasis introduces evolutionary contingency, we focused on the Myo3-Bbc1 and Myo3-Osh2 phenotypes since they had two strong loss of function derived states: V26I and A39G. We observe that for the Bbc1 phenotype, although multiple ancestral and derived states can rescue the loss of function due to A39G (Fig 4E), only A39G rescues the loss of function due to V26I (Fig 4E). On the other hand, for the Osh2 phenotype, two derived states (A39G and T56K) rescue the loss of function due to V26I to the same extent (Fig 4F). The key difference between how these two states rescue loss of function is that T56K, which is a surface substitution, increases the functional score globally in combination with all other states but is limited to the Osh2 phenotype, which points to its role in stabilizing the Osh2 specific binding interface. In contrast, A39G, which is a core substitution, only has this effect for the V26I state, but this effect is present across phenotypes (Fig 4E, 4F, Fig Suppl 15C), suggesting it stabilizes the protein’s fold. Thus, two permissive substitutions are needed for the V26I substitution to fix post-duplication without perturbing the protein interaction network. These results illustrate that seemingly neutral substitutions can not be interchanged even between redundant duplicates, as they are contingent on specific substitutions fixed before them.

## Conclusion

Complex cellular systems have multiple configurations and gene sequences that can give rise to the same phenotypes^61^. This causes such systems to accumulate cryptic genetic variation^62^ that, although currently silent, can, in the future, influence how such systems will respond to mutations and thus evolve. Here, we show that the apparent conservation of protein-binding preference between two duplicated protein-binding domains implicated in endocytosis has accumulated such cryptic variation.

Our results reveal that regulatory and cryptic divergence can contribute to which mutations become accessible for a paralogous protein following gene duplication. Regulatory divergence can lead to divergence in protein abundance levels, which alters the functional effects of mutations at a global level for a given phenotype. Cryptic protein divergence between paralogs alters functional effects more locally through specific epistatic interactions. Moreover, some of the historical substitutions that led to this divergence between paralogs were contingent on each other due to epistatic interactions between them.

If the effect of a mutation arising in either duplicate is uniform across binding partners, specific protein interactions cannot evolve without changing all interactions. The fate of duplicates would be full redundancy or return to a single copy state. The variation in the peptide motifs recognized by a given SH3 domain makes the genetic landscape of each interaction slightly different, which allows for mutations to have partner-specific effects. In combination with the cryptic divergence of the domains, this determines which mutations will allow redundant paralogs to subfunctionalize.

Our results show that in a protein interaction network, the mutations prone to fixation at any given time depend on the permissive substitutions fixed during the long redundancy phase following a gene duplication event. Predicting the fate of gene duplicates and how the network will reorganize as paralogs subfunctionalize will become more difficult as this redundancy period lasts in time and cryptic variation accumulates.

## Materials and Methods

### Strains, plasmids and culture media

All oligonucleotides used in this study are listed in Supplementary Data 1. All strains used in this study are listed in Supplementary Data 10. All plasmids used in this study are listed in Supplementary Data 11. Recipes for all culture media used in this recipe are listed in Supplementary Data 12. All yeast growth experiments were performed at 30 °C unless stated otherwise, and all bacteria were grown at 37 °C. Liquid cultures were incubated with agitation at 250 rpm. All Sanger sequencing was performed at the Centre Hospitalier de l’Université Laval sequencing platform (Université Laval, Québec). All high-throughput sequencing was performed using the Novaseq Reagent Kit v1.5 on an Illumina Novaseq for 500 cycles for paired-end 250 sequencing at the Centre Hospitalier de l’Université Laval sequencing platform (Université Laval, Québec) and the MiSeq Reagent Kit v3 on an Illumina MiSeq for 600 cycles for paired-end 300 sequencing (IBIS sequencing platform, Université Laval).

### Strain construction

All strains were constructed in the BY4741 and BY4742 genetic backgrounds. For the bulk competition DHFR-PCA experiments, we constructed BY4741 (MATa, *his3Δ leu2Δ met15Δ ura3Δ*) strains with *MYO3* or *MYO5* locus fused at its 3’ to the DHFR F[1,2] with a nourseothricin-resistance marker (NAT 100 μg/ml, Jena Bioscience GmbH, Germany). To insert the variant libraries instead of the wild-type SH3 domains, we first replaced the SH3 domain with a flexible linker (GGSSGGGG) using yeast competent cells that were co-transformed with a pCAS plasmid (Addgene plasmid 60847) expressing both the gRNA of interest and *Streptococcus pyogenes* Cas9 and a donor DNA sequence (stuffer) with 60 bp homology arms surrounding the SH3 DNA sequence. The stuffer DNA was PCR amplified^29, 45^. This allowed us to ensure that all the wild-type sequences we observed at the end of the experiments were reconstructed in the same way as the variants.

The prey strains were either constructed or retrieved from the yeast protein interactome collection^51^ (Horizon Discovery, UK). The BY4742 (MATα, *his3Δ leu2Δ lys2Δ ura3Δ*) prey strains are each expressing a gene of interest fused at its 3′ end to the DHFR F[3] with a hygromycin B-resistance marker (HPH 250 μg/ml, Bioshop Canada). We deleted the *MYO3* or *MYO5* loci in these prey strains to sequence only the loci from the BY4741 bait strain. We used a standard lithium acetate transformation protocol^63^ to construct gene deletion strains. The *MYO3* and *MYO5* deletion strains were constructed by amplifying the *URA3* cassettes from pUG72^64^, using the oligonucleotides with respective homology arms for the two loci (primers 327, 328, 329 and 330) and transforming the product in competent cells of the parental strain. Gene deletions were verified by polymerase chain reaction (PCR) using oligonucleotides 323 and 332 for *MYO3* and 325 and 332 for *MYO5,* respectively.

### Creation and validation of SH3 domain variants libraries

As described below, a PCR-based saturation mutagenesis method^45^ was used to generate the mutant libraries. Mutagenesis was carried out on the pUC19 plasmid containing the SH3 domain sequence of *MYO3* and *MYO5* flanked by their native homology arms (Myo3 SH3 in *MYO3* and Myo5 SH3 in *MYO5*) and swapped homology arms (Myo3 SH3 in *MYO5* and Myo5 SH3 in *MYO3*) subsequently used in the CRISPR-mediated genomic re-integration. We used oligos containing degenerate nucleotides (NNN) to perform the mutagenesis at each codon. In the first step of the two-step PCR procedure (short PCR step), an amplicon was generated with an oligo containing the degenerate codon positioned within the domain sequence and another oligo lying outside the SH3 domain nucleotide sequence in the plasmid. The short PCR step was performed with the following settings: 5 min at 95 °C, 35 cycles: 20 s at 98°C, 15s at 60°C, 30s at 72°C, and a final extension of 1min at 72°C. The amplicon generated in this step was used as a mega primer in the second step (long PCR step) to amplify the whole plasmid. A long PCR step was performed with the following settings: 5 min at 95°C, 22 cycles: 20s at 98°C, 15s at 68°C, 3min30s at 72 °C, and a final extension of 5 min at 72 °C. The long PCR product was digested with DpnI for 2 h at 37 °C. The digestion product was then transformed into *E. coli* MC1061 competent cells. In order to obtain most of the 64 possible codons, we aimed to recover ∼1000 colonies from each plate of single mutant libraries. Saturation mutagenesis was performed in a codon-position-wise manner. Mutants at each position were stored separately. Following transformation, the plasmid DNA was extracted from bacteria using a plasmid extraction kit (FroggaBio) to validate the plasmid single mutant libraries. Library quality control was assessed by amplifying the purified plasmid DNA preparations by PCR, followed by validation using MiSeq Reagent Kit v3 on an Illumina MiSeq for 600 cycles for paired-end 300 sequencing (IBIS sequencing platform, Université Laval).

### CRISPR-Cas9 knock-in of the SH3 domain mutants at the native and swapped loci

The genomic insertion of the mutant libraries (one library per amino acid position) was performed by targeting the stuffer sequence in the *MYO3* and *MYO5* SH3-deleted strains, as described in the strain construction section. *MYO3* and *MYO5* SH3-deleted yeast competent cells were co-transformed with pCAS (Addgene plasmid 60847) containing the stuffer-specific gRNA and *Streptococcus pyogenes* Cas9 with the PCR amplified *MYO3* and *MYO5* SH3 single position mutated libraries from the pUC19 preparations acting as the donor DNA^29, 45, 50^. We aimed to obtain ∼1000 colonies from each plate, which were retrieved by washing with sterile YPD and scraping. Glycerol stocks of yeast cells were kept for each position (4 x 59 different stocks corresponding to the 59 amino acid positions of *MYO3* SH3 in *MYO3*, *MYO5* SH3 in *MYO3*, *MYO3* SH3 in *MYO5*, and *MYO5* SH3 in *MYO5*). A master pool for each library type was generated by pooling 5U OD of each codon. For validating the libraries after CRISPR transformation in yeast, yeast genomic DNA extraction of the libraries of the two *MYO3* (native and swapped SH3) and the two *MYO5* (native and swapped SH3) mutants for the deep mutational scanning experiments were performed using a standard phenol/chloroform protocol. Library quality control was assessed by amplifying the genomic DNA preparations by PCR followed by validation using MiSeq Reagent Kit v3 on an Illumina MiSeq for 600 cycles for paired-end 300 sequencing (IBIS sequencing platform, Université Laval, Québec).

### CRISPR-Cas9 knock-in of the SH3 domain mutants validation set

The SH3 variant strains tested as part of the validation studies were constructed using the same strategy as that used to generate the DMS libraries. We initially selected 60 variants for validation studies (15 variants in each of the four genetic backgrounds), encompassing a wide range of phenotypic effects across three binding partners. To generate mutated donor DNA, we used a fusion PCR strategy where oligonucleotides containing the mutations of interest are used to generate two overlapping amplicons. We started from genomic DNA extracted from strains of BY4741 with Myo3 SH3 in *MYO3*, Myo5 SH3 in *MYO3*, Myo3 SH3 in *MYO5*, and Myo5 SH3 in *MYO5* and used the primers to generate the two overlapping amplicons both bearing the mutation of interest. The first reaction, which was amplified from the 5’ flanking homology arm before the start of the SH3 domain coding sequence up to the mutated amplicon overlap, used oligonucleotides 302 and 304 as the forward primers and reverse primers from reverse primers present in 234-301. In the second PCR reaction, amplification was done from the mutated overlap to the 3’ homology arm after the SH3 domain coding sequence, using the forward primers present between 234-301 as forward primers and oligonucleotides 303 and 305 as reverse primers. Both resulting amplicons are then diluted 1:20 and used as the template for a second reaction using oligonucleotides 302, 303 for the *MYO5* locus and 304, 305 for the *MYO3* locus to reconstitute the full-length SH3 domain sequence.

We then performed one-by-one CRISPR knock-ins using the same reaction setup as for the DMS library but using the fusion PCR products as donor DNA. Transformants (four per mutant) were used to inoculate 1.2 ml YPD cultures, grown overnight, and then streaked on YPD. Two colonies from each streak were resuspended in 200 μl water and spotted on YPD as well as YPD + G418 and YPD + NAT to check for pCAS loss and the presence of the *NATMX4* cassette to ensure the DHFR[1,2] tag was still present. SH3 domain integration was verified by PCR using oligonucleotides 306, 307 and 308 on quick DNA extractions^65^. PCR-positive strains were then validated by Sanger sequencing using oligonucleotide 308 as the sequencing oligonucleotide. We succeeded in reconstructing all 60 of the selected 60 mutants.

### CRISPR-Cas9 knock-in of chimeric variants at the native locus

Chimeric variants of divergent substitutions between the two SH3 domains with single and double amino substitutions and the wildtype sequence with 20bp homology arms on either side were synthesized as oligonucleotide pools (Integrated DNA Technologies Ltd) for insertion at the native loci. The genomic insertion of the synthesized libraries was performed by targeting the stuffer sequence in the *MYO3* and *MYO5* SH3-deleted strains. *MYO3* and *MYO5* SH3-deleted yeast competent cells were co-transformed with pCAS (Addgene plasmid 60847) containing the stuffer-specific gRNA and *Streptococcus pyogenes* Cas9^29, 45, 50^ with the PCR amplified *MYO3* and *MYO5* SH3 chimeric variant synthesized libraries acting as the donor DNA, respectively. We targeted ∼1000 colonies on each plate, which were retrieved by washing with sterile YPD. Glycerol stocks of yeast cells were kept for each position (2 different stocks corresponding to the chimeric variants of Myo3 SH3 in *MYO3* and Myo5 SH3 in *MYO5*). The stocks were validated using MiSeq Reagent Kit v3 on an Illumina MiSeq for 600 cycles for paired-end 300 sequencing (IBIS sequencing platform, Université Laval, Québec).

### Liquid DHFR-PCA with deep mutational scanning variant libraries

Bulk competition liquid DHFR-PCA experiments were performed using the four yeast strains with the SH3 single-position mutant libraries with various prey strains in three independent biological replicates. Twelve overnight cultures (three replicates for each domain-locus combination) were started in YPD + NAT using the master pool containing each mutated position. Three independent overnight precultures were started for each prey strain tagged with DHFR F[3] by picking individual colonies from streaks on YPD + HPH plates. Then, the master pools and prey strains tagged with DHFR F[3] were mixed in YPD (2:1 ratio) for mating and incubated for 8 h at 30 °C with shaking at 400 rpm^45^. Diploid cells were selected for the first time by transferring the mixture in a YPD + NAT + HPH (OD 0.5) culture for 16 h at 30 °C with shaking at 400 rpm. The next day, the first diploid selection was transferred in SC complete pH 6.0 + NAT + HPH at 0.5 OD, and the culture was grown for 24 h at 30 °C with shaking at 400 rpm. A fraction of the cells (5U OD) was collected for Illumina sequencing at this time point (first DHFR-PCA experiment time point without PCA selection, reference condition S2), following which the optical density of the diploid cells was read using a TECAN Infinite F200 Pro and diluted to 0.1 OD in 15 mL of PCA selection media without agar in a 50 mL tube. Tubes were incubated at 30 °C for 72 h (second-time point of the DHFR-PCA experiment, first PCA selection). The cells from the first PCA step were grown for a second PCA selection step using the same procedures (final DHFR-PCA experiment time point, condition MTX2). A fraction of the cells (5U OD) was collected for Illumina sequencing. Finally, the genomic DNA of the cells (5U OD) was extracted using MasterPure Yeast DNA Purification Kit (Biosearch Technologies) before and after two cycles of PCA selection (S2 and MTX2). The surviving SH3 mutants in each experiment were detected by Illumina sequencing, as described below.

### Liquid DHFR-PCA with chimeric variant libraries

For the chimera experiment, bulk competition liquid DHFR-PCA was performed using the two yeast strains with the SH3 chimeric variant libraries with various prey strains in three independent biological replicates. The six master pools (three replicates for *MYO3* and three replicates for *MYO5*) were thawed and grown overnight in liquid YPD. Three independent overnight precultures were started for each prey strain tagged with DHFR F[3] by picking individual colonies from streaks on YPD + HPH plates. Then, the master pools and prey strains tagged with DHFR F[3] were mixed in YPD (2:1) for mating and incubated for 8 h at 30 °C with shaking at 250 rpm. Diploid cells were selected for the first time by transferring the mixture in a YPD + NAT + HPH (OD 0.5) culture for 16 h at 30 °C with shaking at 250 rpm. The next day, the first diploid selection was transferred in SC complete pH 6.0 + NAT + HPH at 0.5 OD, and the culture was grown for 24 h at 30 °C with shaking at 250 rpm. A fraction of the cells (5U OD) was collected for Illumina sequencing at this time point (first DHFR-PCA experiment time point without PCA selection, reference condition S2), following which the optical density of the diploid cells was read using a TECAN Infinite F200 Pro and diluted to 0.1 OD in 15 mL of PCA selection media without agar in a 50 mL tube. Tubes were incubated at 30 °C for 72 h (second-time point of the DHFR-PCA experiment, first PCA selection). The cells from the first PCA step were grown for a second PCA selection step using the same procedures (final DHFR-PCA experiment time point, condition MTX2). A fraction of the cells (5U OD) was collected for Illumina sequencing. Finally, the genomic DNA of the cells (5U OD) was extracted using MasterPure Yeast DNA Purification Kit (Biosearch Technologies) before and after two cycles of PCA selection (S2 and MTX2). The surviving SH3 mutants in each experiment were detected by Illumina sequencing, as described below.

### Low-throughput liquid DHFR-PCA of independently reconstructed SH3 variants

Low-throughput liquid DHFR-PCA experiments were performed to validate the protein-protein interaction scores obtained from the high-throughput bulk competition. Mating was performed in liquid YPD by combining the reconstructed bait strain mutant and prey strain in a 1:1 ratio, and diploid selection was performed by spotting on solid YPD + NAT + HPH plates with agar in three independent biological replicates. Following diploid selection, diploid cells were inoculated in a 96 v-shaped well plate using an SC complete medium with MSG at pH 6.0 with NAT and HPH. After 24h of growth, OD was read using a TECAN Infinite F200 Pro, and dilutions to 1 OD in sterile nanopore water were prepared. Cells were then diluted to 0.1 OD by combining 25 µl of cells at 1 OD and 225 µl of PCA selection media without agar in a 96-well plate. Plates were incubated at 30 °C in a TECAN Infinite M Nano, and OD was monitored every 15 min for 72 h.

### Deep mutational scanning libraries sequencing

Sequencing of yeast genomic DNA was performed at two different time points during the liquid DHFR-PCA experiments: After diploid selection of the bait–prey strains (reference condition S2) and following the two PCA selection rounds (condition MTX2). Genomic DNA was extracted from samples at the two timepoints of the PCA experiment using MasterPure Yeast DNA Purification Kit (Biosearch Technologies).

The libraries for sequencing the deep mutational scanning experiments were prepared by three successive rounds of PCR. The first PCR was performed with primers to amplify SH3 domains from saturation mutagenesis mini preps (4.5 ng of plasmid) or on genomic DNA extracted from yeast (90 ng of genomic DNA) (PCR program: 3 min at 98 °C, 20 cycles: 30s at 98°C,15s at 60°C and 30s at 72°C, final elongation 1min at 72°C) using oligonucleotides 169, 170 for *MYO3* locus and 171, 172 for *MYO5* locus. The second PCR was performed to increase diversity in the libraries by adding row and column barcodes (oligonucleotides 173-197) for identification in a 96-well plate (PCR program: 3 min at 98°C, 15 cycles: 30s at 98°C, 15s at 60°C and 30s at 72°C, final elongation 1 min at 72 °C). The first PCR served as a template for the second one (2.25 μl of 1/ 2500 dilution). After the second PCR, 2 μl of the products were run on a 1.5% agarose gel, and band intensity was estimated using Image Lab (BioRad Laboratories). The PCR products were mixed based on their intensity on an agarose gel to roughly equal amounts of each in the final library. Mixed PCRs were purified on magnetic beads and quantified using a NanoDrop (ThermoFisher). The third PCR was performed on 0.0045 ng of the purified pool from the second PCR to add plate barcodes (oligonucleotides 198-210) and Illumina adapters (PCR program: 3 min at 98 °C, 15 cycles: 30 s at 98°C, 15s at 61°C and 35s at 72°C, final elongation 1min 30s at 72°C). Each reaction for the third PCR was performed in four replicates, combined, and purified on magnetic beads. After purification, libraries were quantified using a NanoDrop (ThermoFisher). Equal amounts of each library were combined and sent to the Genomic Analysis Platform (CHUL, Quebec, Canada) for paired-end 250 bp sequencing on a NovaSeq (Illumina).

### Chimeric libraries sequencing

Yeast genomic DNA extraction of the libraries of the *MYO3* and *MYO5* SH3 chimeric variants was performed using a standard phenol/chloroform extraction protocol. Sequencing of yeast genomic DNA was performed at two different time points during the liquid DHFR-PCA experiments: After diploid selection of the bait–prey strains (reference condition S2) and following the two PCA selection rounds (condition MTX2).

The libraries for sequencing the chimera experiments were prepared by two successive rounds of PCR. The first PCR was performed with primers to amplify SH3 domains on genomic DNA extracted from yeast (90 ng of genomic DNA) (PCR program: 3 min at 98 °C, 20 cycles: 30s at 98°C,15s at 60°C and 30s at 72°C, final elongation 1min at 72°C) using primers with homology to the flanking regions of the Myo3 SH3 at the *MYO3* locus and Myo5 SH3 at the *MYO5* locus (oligonucleotides 211-218). The PCR products were purified on magnetic beads and quantified using a NanoDrop (ThermoFisher). The second PCR was performed on 0.0045 ng of the purified product from the first PCR to add Nextera barcodes and Illumina adapters (PCR program: 3 min at 98 °C, 15 cycles: 30 s at 98°C, 15s at 61°C and 35s at 72°C, final elongation 1min 30s at 72°C) using oligonucleotides 219-233. Each reaction for the second PCR was performed in two replicates, combined, and purified on magnetic beads. After purification, libraries were quantified using a NanoDrop (ThermoFisher). Equal amounts of each library were combined and sent to the Genomic Analysis Platform (CHUL, Quebec, Canada) for pair-end 250 bp sequencing on a NovaSeq (Illumina).

### Analysis of deep sequencing data for deep mutational scanning experiments

Quality control of the NovaSeq sequencing data was performed using FastQC version 0.11.4^66^. Trimmomatic version 0.39^67^ was used to select reads with a minimal length from the raw data (MINLEN parameter: 250) and trim them to a final length (CROP parameter: 225). Selected reads were aligned using Bowtie version 1.3.0^68^ to the plate, row, and column barcodes to demultiplex sequences from each pool (bait variant library + prey + time point). The remaining paired reads were merged with Pandaseq version 2.11^69^, and identical sequences were aggregated using vsearch version 2.15.1^70^ and aligned to the appropriate SH3 amplicon (Myo3 SH3 in *MYO3*, Myo5 SH3 in *MYO3*, Myo3 SH3 in *MYO5*, and Myo5 SH3 in *MYO5*). Next, we filtered the alignments to keep only the sequences that had an exact match to the wild-type SH3 sequence or mutations at a single codon of the SH3 sequence. Reads with mutations at multiple codons were discarded, and we retained between 70-80% of merged reads following these filters.

For computing the log fold-change (F) between before and after methotrexate selection, we added one to all variant counts to calculate fold-change for variants whose counts dropped to zero after selection. These variant read counts were then divided by total read counts for each pool to obtain variant frequencies of that pool. The fold-change of a variant was calculated by comparing the variant frequencies between reference condition (S2) and after methotrexate selection (MTX2). We discarded all codon-level variants which had a read count below ten reads at S2 for at least one biological replicate. Most codon-level variants were well covered at the reference condition (S2), with median read counts between 130-170 in each pool, resulting in a very high coverage for amino acid variants. The distribution of log fold-changes for all experiments was bimodal, with one peak corresponding to the distribution of synonymous codon variants of wild-type SH3 sequence (wild-type-like interaction) and the other corresponding to the distribution of stop-codon variants (non-functional interaction). Hence, we calculated the codon-level functional score by scaling the codon-level log fold changes in each biological replicate between the median of synonymous codon variants of wild-type SH3 sequence (1) and the median of all stop-codon variants (0). Then, we calculated the amino acid variant functional score (ΔF) for each biological replicate as the median of functional scores of the synonymous codon variants for each amino acid variant. Finally, we calculated the mean functional score (ΔF) for each amino acid variant as the mean of the functional scores from three biological replicates.

### Analysis of deep sequencing data for chimera experiments

We obtained demultiplexed reads from the Novaseq sequencing run for the chimera experiments due to the use of Nextera indices. The NovaSeq sequencing data quality control was performed using FastQC version 0.11.4^66^. Trimmomatic version 0.39^67^ was used to select reads with a minimal length from the raw data (MINLEN parameter: 210) and trimmed with the Nextera adapter command ILLUMINACLIP:NexteraPE-PE.fa:2:30:10:2:True using a sliding window approach such that mean PHRED score in the window was greater than 33. The retained paired reads were merged with Pandaseq version 2.11^69^, and identical sequences were aggregated using vsearch version 2.15.1^70^ and aligned to the appropriate SH3 amplicon (Myo3 SH3 in *MYO3* and Myo5 SH3 in *MYO5*). Next, we filtered the alignments to keep only the sequences that had an exact match to the wild-type SH3 sequence or synthesized chimeric mutations in the SH3 sequence. Reads with mutations at other codons which were not part of the synthesized library were discarded, and we retained between 70-80% of merged reads following these filters.

For computing the log fold-change between before and after methotrexate selection, we added one to all variant counts to calculate fold-change for variants whose counts dropped to zero after selection. These variant read counts were then divided by total read counts for each pool to obtain variant frequencies of that pool. The fold-change of a variant was calculated by comparing the variant frequencies between reference condition (S2) and after methotrexate selection (MTX2). We discarded all variants which had a read count below ten reads at S2 for at least one biological replicate. Most variants were well covered at the reference condition (S2), with median read counts varying between 2000-3000 in each pool, resulting in a very high coverage for chimeric variants. Since this was a small-scale bulk competition assay, we obtained a unimodal distribution of log fold changes. Next, we calculated the functional score for each chimeric variant (*F_C_*) for each biological replicate as the difference in the log fold-change of the chimeric variant and the median of all variants in a pool. Finally, we calculated each chimeric variant’s mean functional score (*F_C_*) as the arithmetic mean of the functional scores from three biological replicates.

### Expression level measurement by flow cytometry

To determine promoter strength for each paralogous locus, we replaced the complete coding sequence of *MYO3* and *MYO5* with eGFP. First, we replaced the coding sequence from start to stop codon with a hygromycin (HPH) resistance marker using a standard lithium acetate transformation protocol^63^. Once those strains were constructed, we co-transformed them with a pCAS plasmid (Addgene) targeting the HPH marker and the eGFP sequence with homology arms for each locus^50^. Proper replacement of the HPH marker with the eGFP was confirmed by PCR (primers 322, 323 for *MYO3* and 324, 325 for *MYO5*). To measure the eGFP signal, 8 clones of each locus were grown overnight in liquid SC complete MSG pH 6.0 at 30°C at 250 rpm. The following morning, the cultures were diluted at 0.1 OD/ml in fresh SC complete MSG pH 6.0 and grown until ∼0.5 OD/ml. Once the appropriate OD was reached, cells were diluted 10x in sterile water in a 96 well plate (Greiner). For each clone, 5000 events were acquired with a Guava EasyCyte instrument (Cytek Bio). We calculated the *log*_2_ median fluorescence intensity for each clone from the 5000 measurements obtained from the flow cytometer to quantify the promoter activity at each locus.

### Ancestral sequence reconstruction

We used a recently inferred ML phylogeny for type-1 fungal myosins^44^ with Microsporidia as an outgroup to reconstruct the ancestral sequences of the C-terminal SH3 domain pre-duplication. Most probable ancestral SH3 domain sequences were reconstructed on this ML phylogeny FAST-ML (v3.11)^71, 72^. The evolutionary trajectory of the sequence changes was determined from the amino acid differences between reconstructed sequences on successive ancestral nodes on the lineage from the common ancestor of Fungi (ancFungi) to the duplication node of the type-1 myosins (AncDuplication). Ancestral states are amino acid states not present in AncDuplication that occurred in at least one ancestral node on the lineage from ancFungi to AncDuplication. The maximal posterior probability for an ancestral amino acid state is determined by taking the highest posterior probability of reconstruction for that state at any ancestral node in the evolutionary trajectory.

### Classification of binding and non-binding variants

Next, we used mean functional scores to classify the strength with which each library variant binds to various binding partners using a nonparametric comparison to distributions of nonsense mutants. A variant was classified as binding if its mean functional score was significantly greater than that of nonsense variants contained in the library. To determine whether a missense variant’s functional score is significantly greater than the distribution of nonsense variants, we generated 10,000 bootstrap replicates from the functional score of nonsense variants. For each variant, we calculated an empirical p-value for the null hypothesis that a variant is non-binding as the proportion of bootstrap replicates with a greater mean functional score than the functional score of that variant. Variants were assigned as binding if the null hypothesis could be rejected at a 1% false discovery rate (using the Benjamini-Hochberg correction method) or non-binding if the null hypothesis could not be rejected. This classification method was followed for each replicate, and a variant was classified as active for a pair of binding partners only when it was classified as binding or non-binding in all three replicates. This conservative approach allowed us to reduce false positives instead of choosing a hard threshold.

### Data and code availability

Raw sequencing files for all assays have been deposited on the NCBI SRA (accession number PRJNA1079703). All oligonucleotides used in this study are listed in supplementary_file_1.xlsx. Log fold changes for all deep mutational scanning experiments are listed in supplementary_file_2.xlsx. Functional scores for all deep mutational scanning experiments are listed in supplementary_file_3.xlsx. Functional scores for all chimera experiments are listed in supplementary_file_4.xlsx. Phenotypic landscapes for all deep mutational scanning experiments are listed in supplementary_file_5.pdf. Histograms of read count distribution for all deep mutational scanning experiments are listed in supplementary_file_6.pdf. Histograms of read count distribution for all chimera experiments are listed in supplementary_file_7.pdf. Posterior probabilities of reconstructed ancestral states for the ancestral SH3 domains of the Myo3/Myo5 phylogeny are listed in supplementary_file_8.xlsx. Flow cytometry data for the promoter activity measurement are listed in supplementary_file_9.xlsx. All strains used in this study are listed in supplementary_file_10.xlsx. All plasmids used in this study are listed in supplementary_file_11.xlsx. All culture media recipes used in this study are listed in supplementary_file_12.xlsx. Data from all independent validation experiments are listed in supplementary_file_13.xlsx. All the scripts needed to reproduce the figures in the paper are in https://github.com/Landrylab/Dibyachintan_et_al_2024.

## Author Contributions

Conceptualization: S.D., A.K.D., D.B., C.R.L.; Experiments: S.D., A.K.D., P.L.; Data analysis: S.D., D.B.; Development of methodology: S.D., A.K.D., U.D.; Writing – original draft: S.D., C.R.L.; Writing – review and editing: S.D., A.K.D., D.B., P.L., U.D., C.R.L.; Funding acquisition: C.R.L.

## Acknowledgement

We thank Angel Cisneros, Francois Rouleau and Phillipe Despres for insightful discussions and feedback on the manuscript. All statistical analyses were performed with scipy v1.9.1 on Python 3.9. All figures were created with Python 3.9 and Adobe Illustrator. All molecular graphics and analyses performed with UCSF ChimeraX, developed by the Resource for Biocomputing, Visualization, and Informatics at the University of California, San Francisco, with support from National Institutes of Health R01-GM129325 and the Office of Cyber Infrastructure and Computational Biology, National Institute of Allergy and Infectious Diseases.

---

**Supplementary Figure 1:**
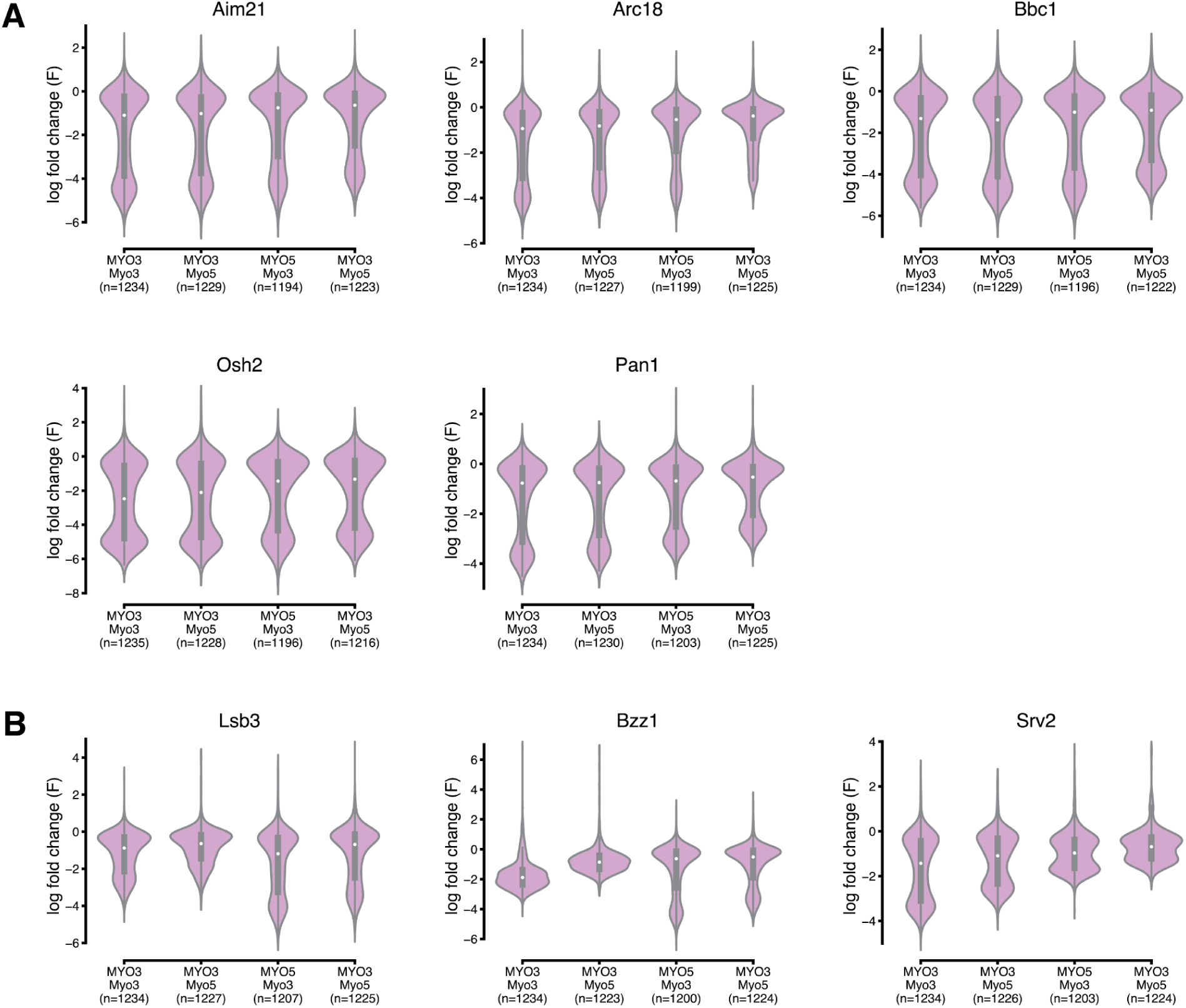
**(A)** Distribution of log fold-changes (F) of mutations for interaction phenotypes where the dynamic ranges are similar across SH3 domain-loci combinations (locus in all caps in the first row and SH3 domain in the second row) **(B)** Distribution of log fold-changes (F) of mutations for phenotypes where the dynamic ranges are dependent on the locus (locus in all caps in the first row and SH3 domain in the second row).

**Supplementary Figure 2:**
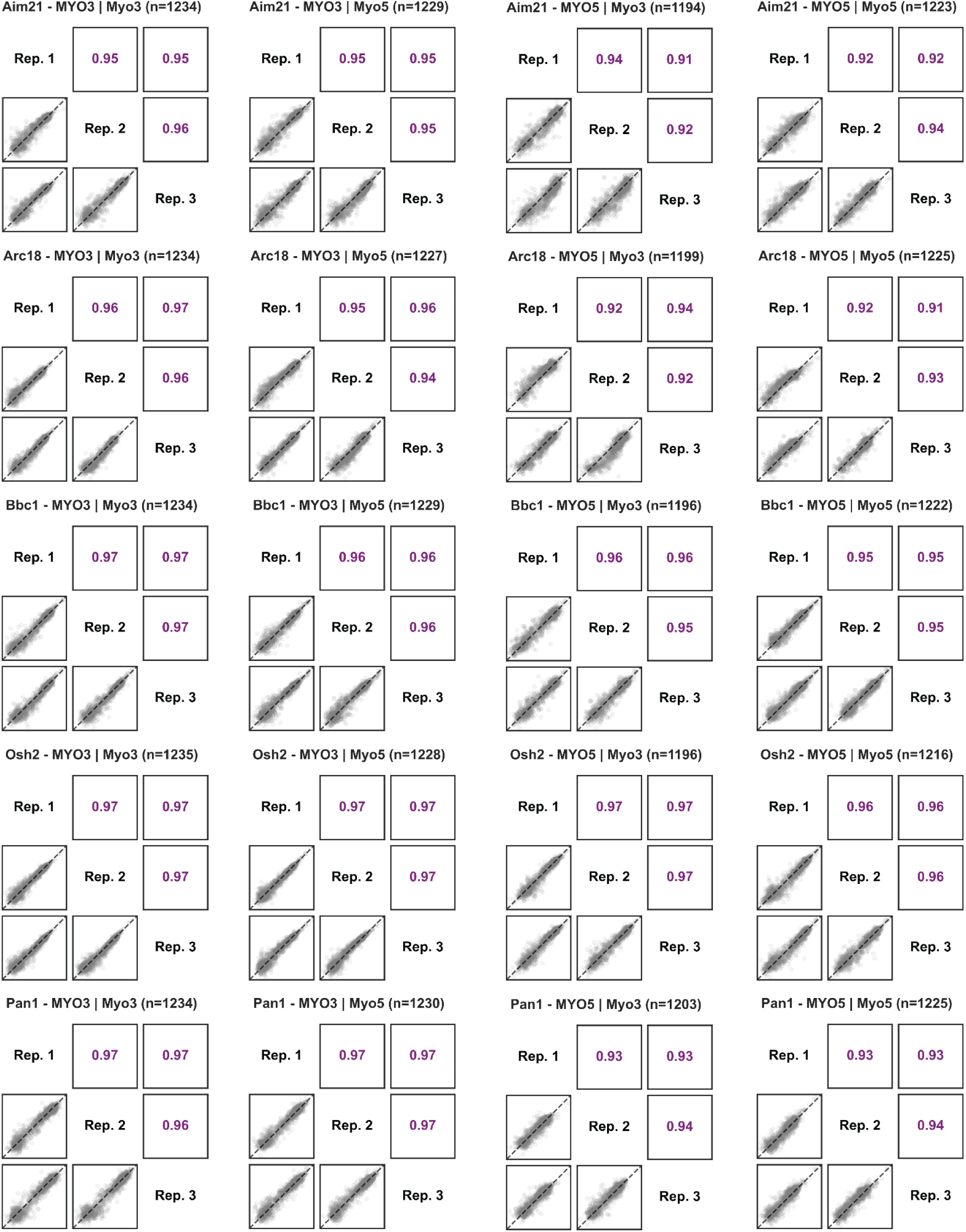
Pearson correlation between the three biological replicates for the functional scores (ΔF) of mutations introduced in the four domain-loci combinations for multiple phenotypes.

**Supplementary Figure 3:**
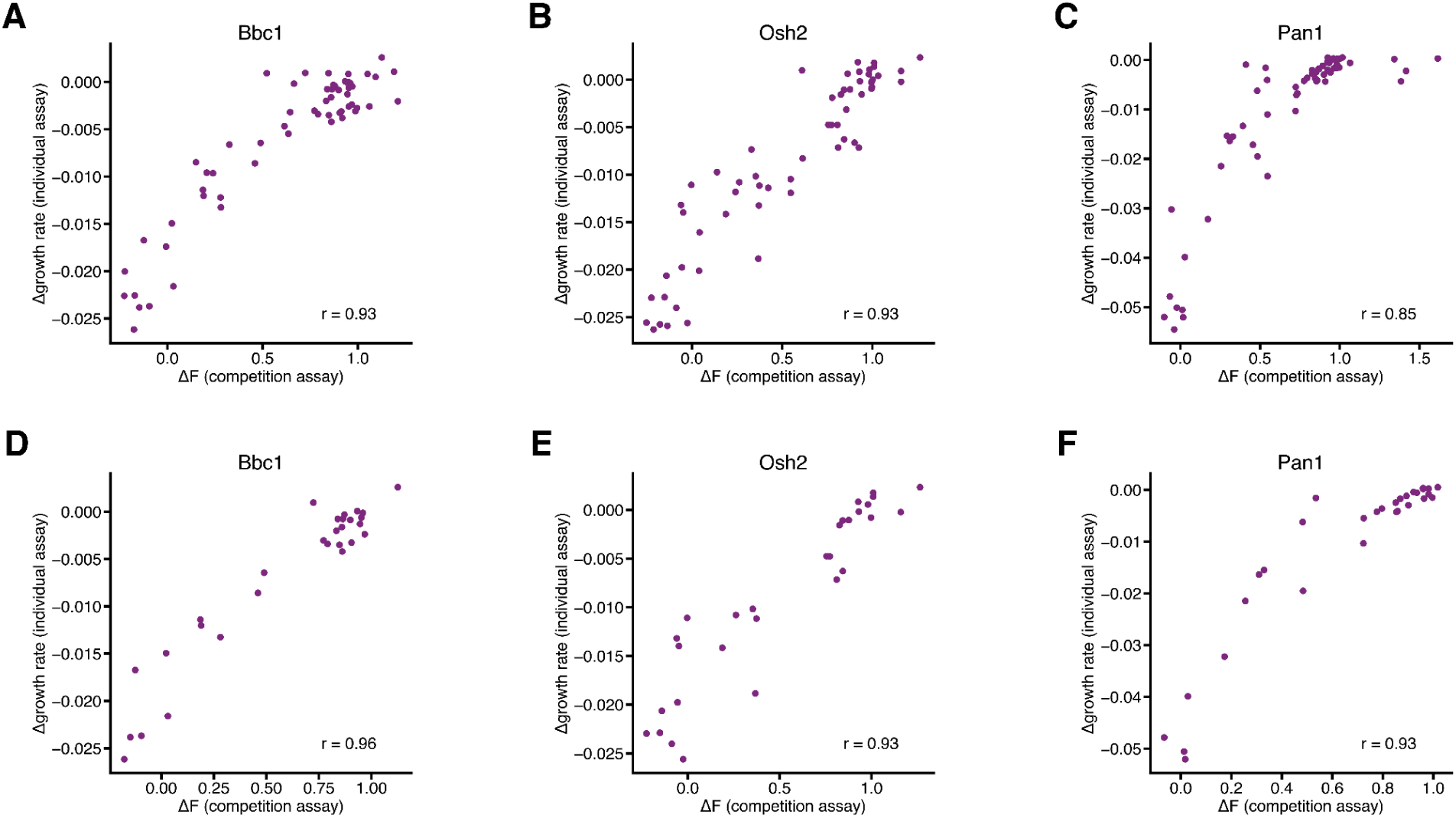
Pearson correlation between functional scores (n=3 biological replicates) in the high-throughput bulk competition assay (x-axis) with the low-throughput validation growth rates with reference to the wild-type protein (n=3 biological replicates) for 60 independently reconstructed mutants in all four domain-loci backgrounds (y-axis) for **(A)** Bbc1 phenotype, **(B)** Osh2 phenotype, and **(C)** Pan1 phenotype. Pearson correlation between functional scores in the high-throughput bulk competition assay (x-axis) with the low-throughput validation growth rates for independently reconstructed 30 mutants in the MYO3 locus for both domains (y-axis) for **(D)** Bbc1 phenotype, **(E)** Osh2 phenotype, and **(F)** Pan1 phenotype.

**Supplementary Figure 4:**
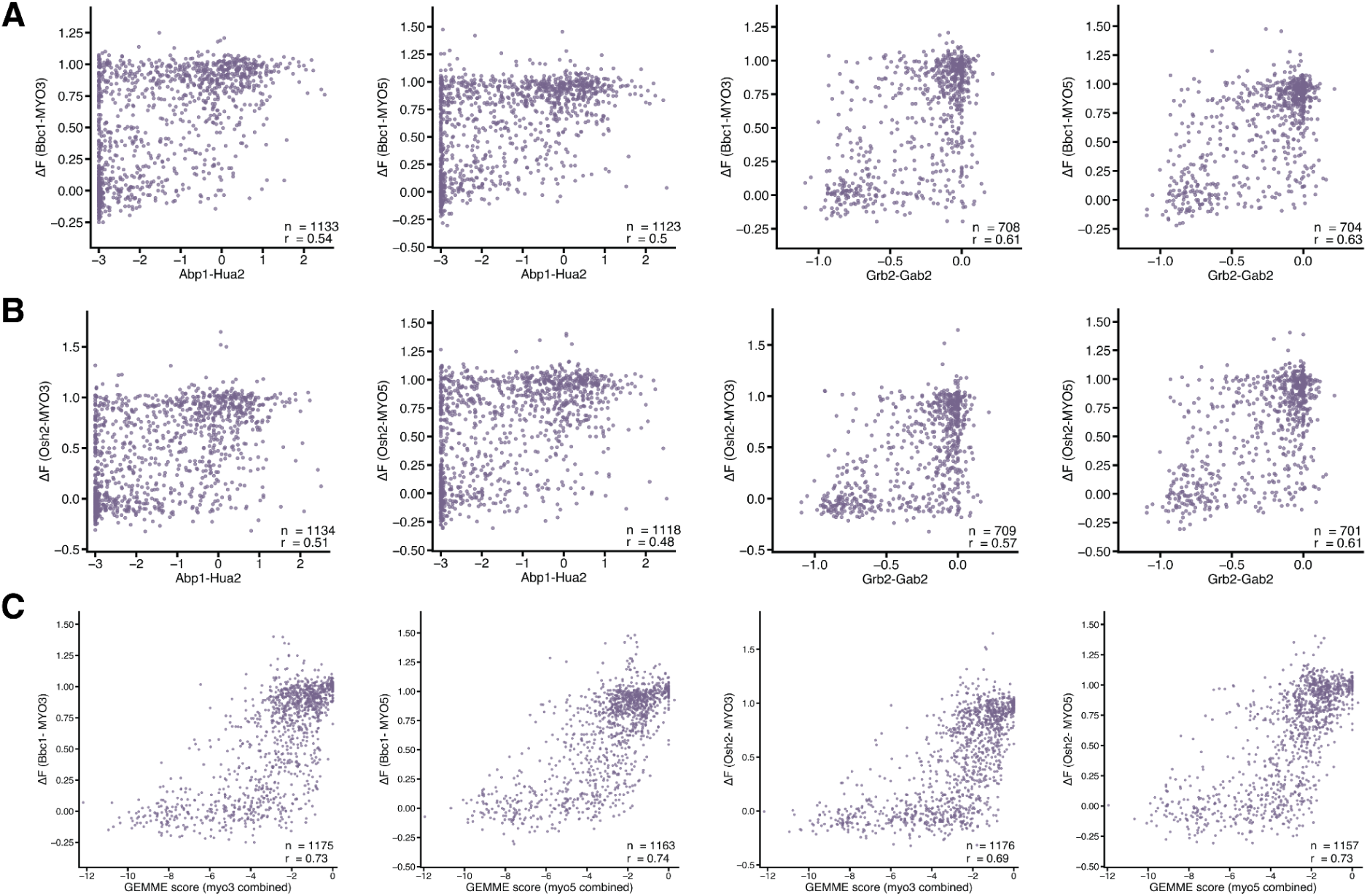
**(A)** Pearson correlation between our experiments involving Bbc1 phenotype with published deep mutational scanning studies involving SH3 domain-mediated interactions for other SH3 domains: Abp1 SH3 - Hua2^29^ (sequence identity between Abp1 and Myo3 SH3 domains = 22%, RMSD between Abp1 (PDB: 1JO8) and Myo3 (PDB: 1RUW) SH3 domains = 1.75) and Grb2 SH3 - Gab2^52^ (sequence identity between Grb2 and Myo3 SH3 domains = 21%, RMSD between Grb2 (PDB: 1JO8) and Myo3 (PDB: 1RUW) SH3 domains = 1.67). Abp1 SH3 is a yeast SH3 domain, and Grb2 SH3 is a human SH3 domain. **(B)** Pearson correlation between our experiments involving Osh2 phenotype with published deep mutational scanning studies involving SH3 domain-mediated interactions: Abp1 SH3 - Hua2^29^ and Grb2 SH3 - Gab2^52^. **(C)** Pearson correlation between our experiments involving Bbc1 and Osh2 phenotype with combined evolutionary models (epistatic and independent) using GEMME^54^ for both Myo3 and Myo5 SH3 domains.

**Supplementary Figure 5:**
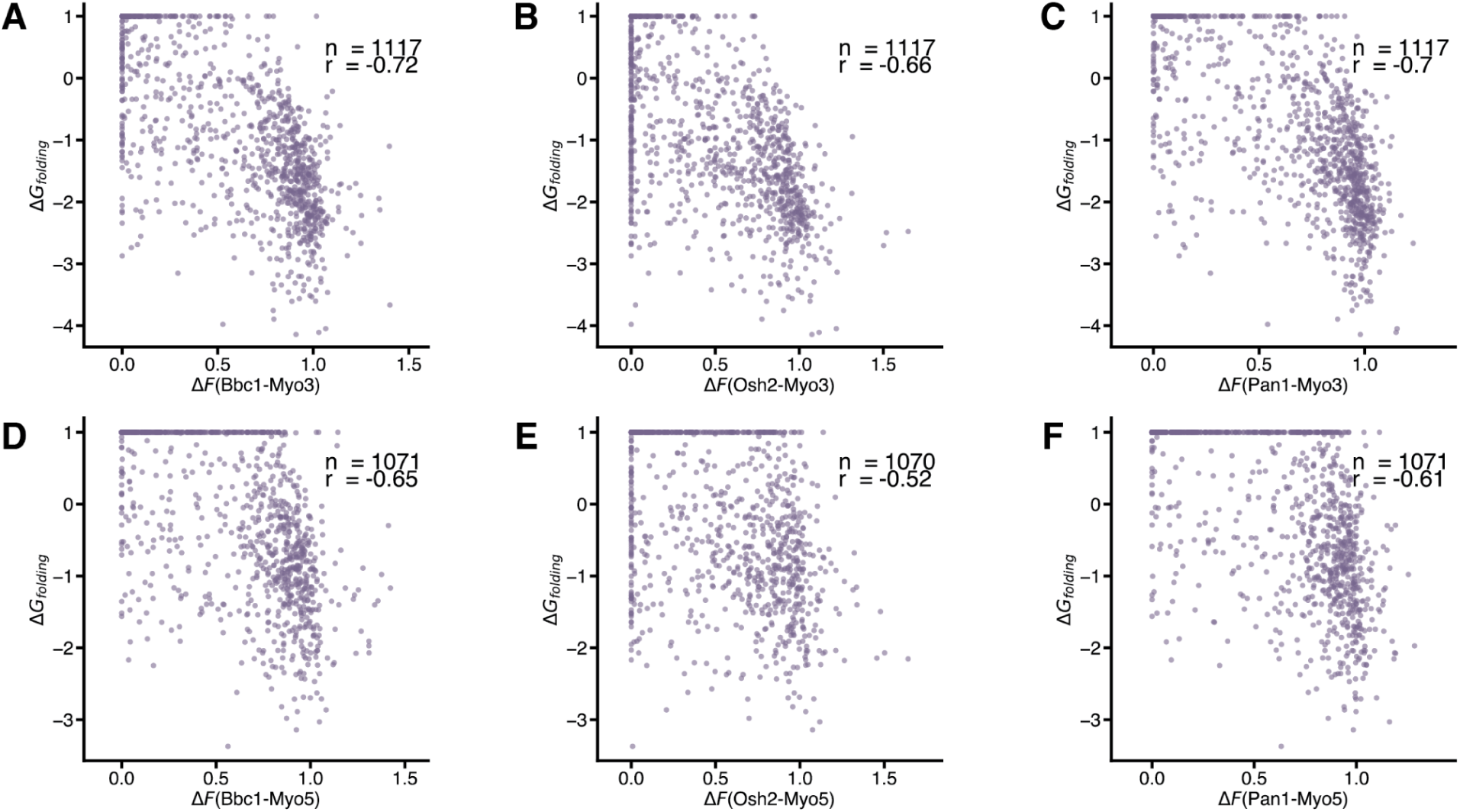
Pearson correlation between functional scores from our experiments at the *MYO3* locus and the folding stability Gibbs Free Energy obtained experimentally^53^ for **(A)** Bbc1-Myo3 SH3 interaction and Myo3 SH3 folding stability, **(B)** Osh2-Myo3 SH3 interaction and Myo3 SH3 folding stability, **(C)** Pan1-Myo3 SH3 interaction and Myo3 SH3 folding stability, **(D)** Bbc1-Myo5 SH3 interaction and Myo5 SH3 folding stability, **(E)** Osh2-Myo5 SH3 interaction and Myo5 SH3 folding stability, and **(F)** Pan1-Myo5 SH3 interaction and Myo5 SH3 folding stability.

**Supplementary Figure 6:**
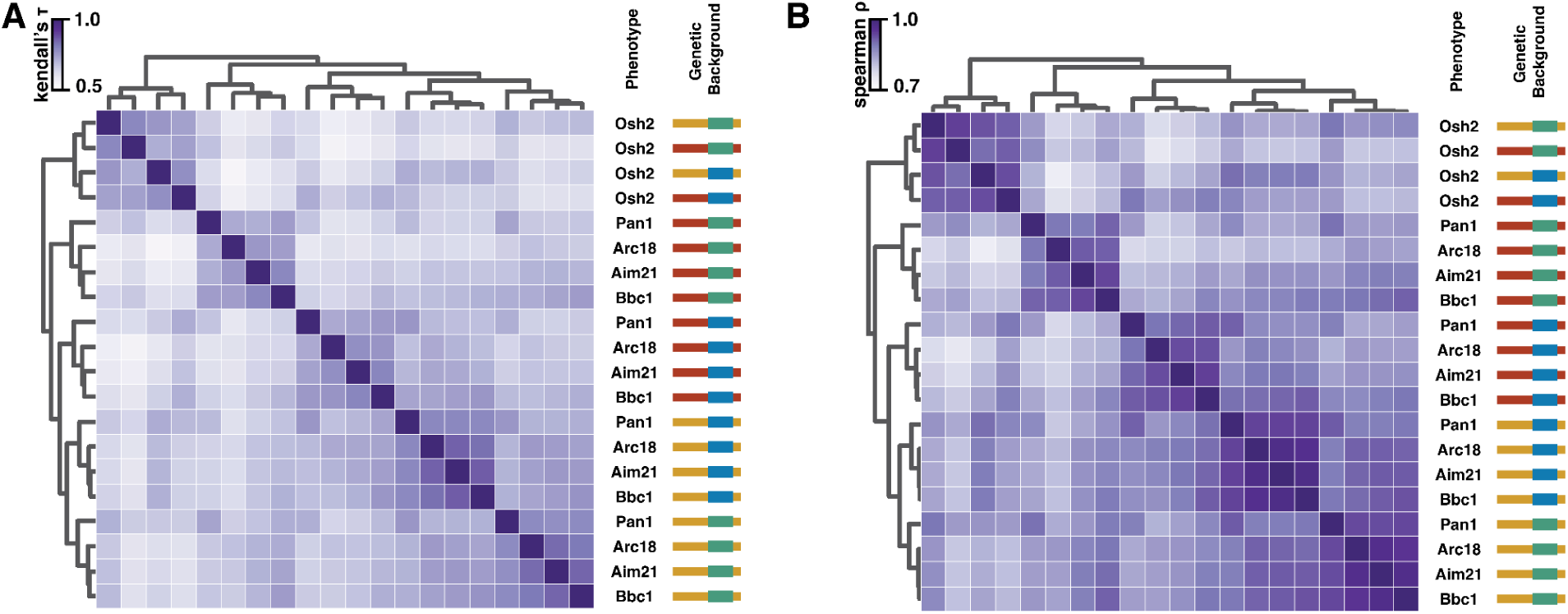
Hierarchical clustering of functional scores (ΔF) for five binding phenotypes across the four domain-loci combinations using **(A)** Kendall-tau correlation coefficients between functional scores of any two experiments and **(B)** Spearman correlation coefficients between functional scores of any two experiments.

**Supplementary Figure 7:**
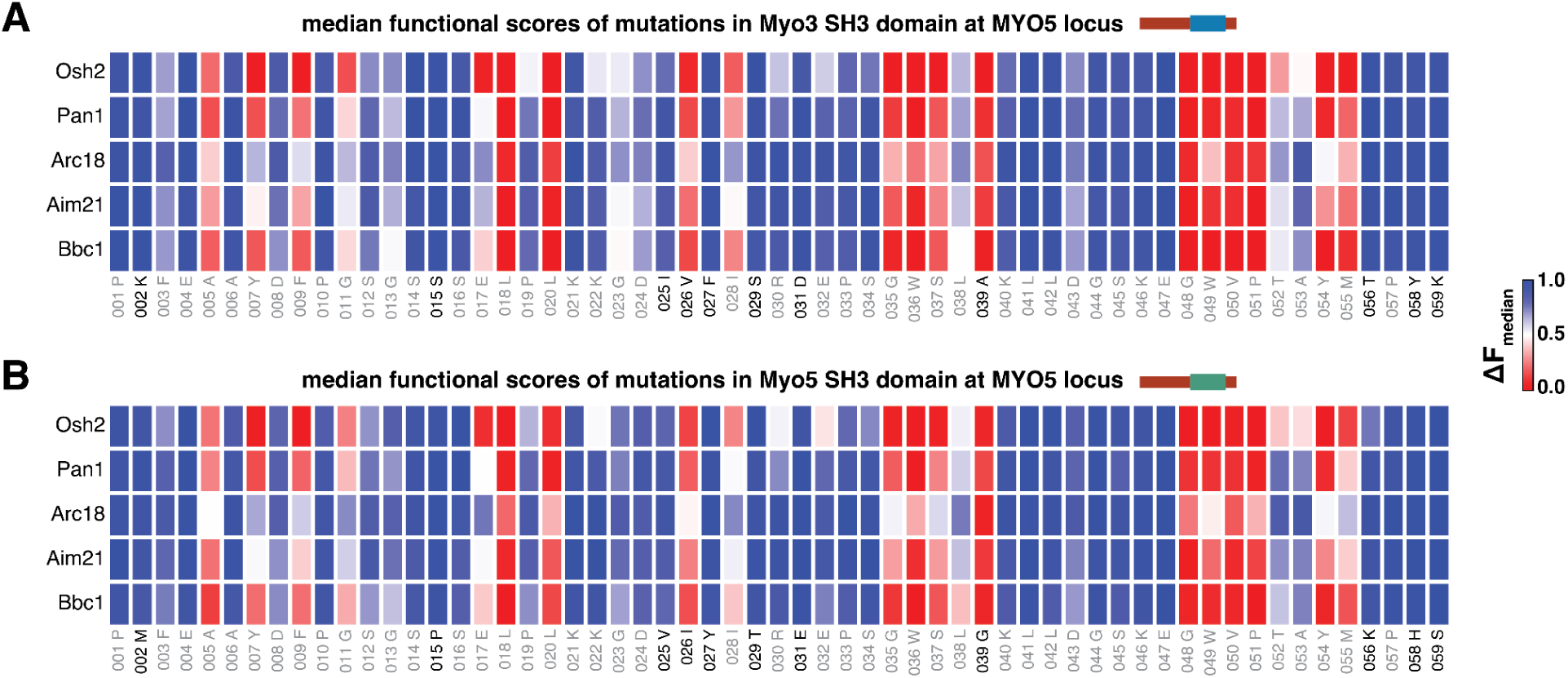
Median functional scores of all 59 sites for five phenotypes at the *MYO5* locus for the **(A)** Myo3 SH3 domain and **(B)** Myo5 SH3 domain.

**Supplementary Figure 8:**
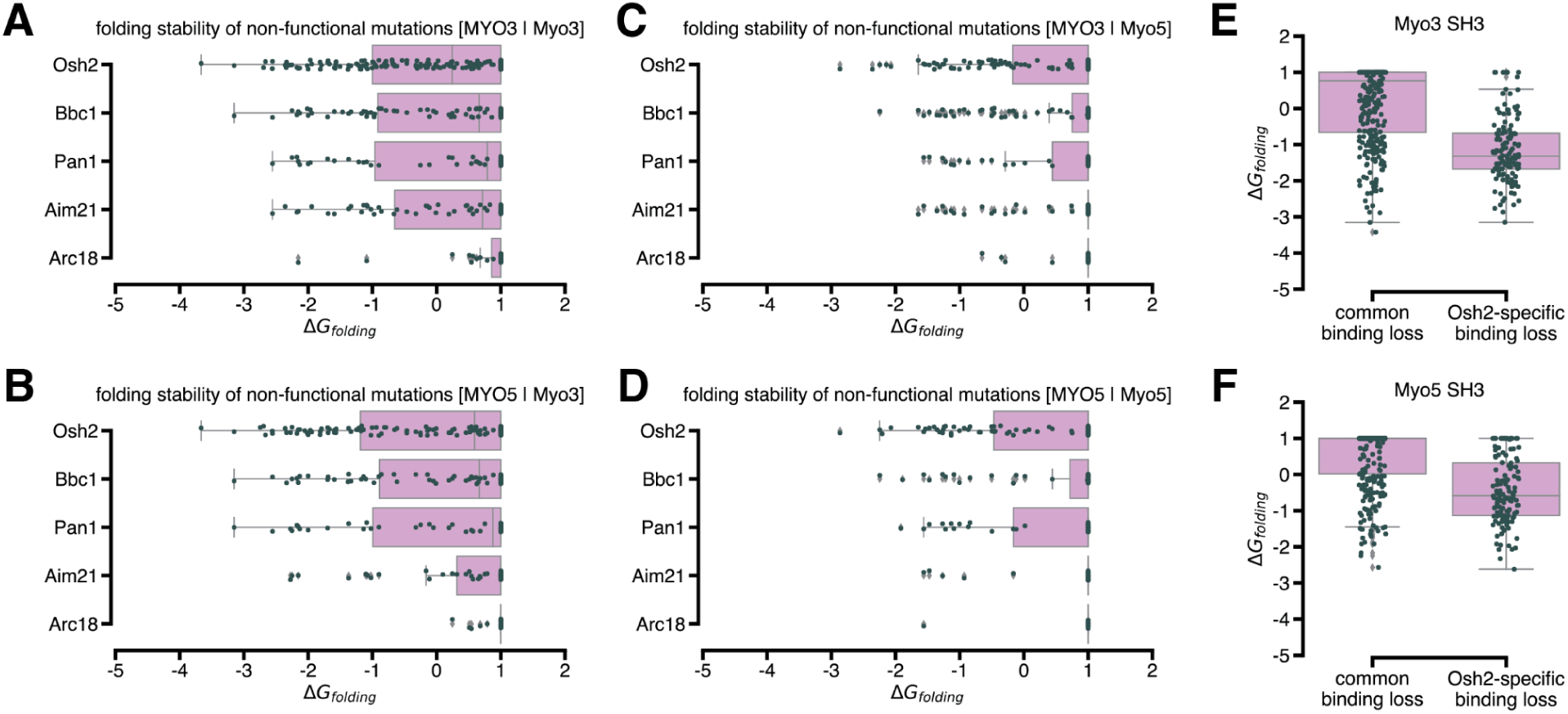
Folding stability Gibbs Free Energy^53^ of non-functional mutations in each phenotype for **(A)** Myo3 SH3 at the *MYO3* locus, **(B)** Myo3 SH3 at the *MYO5* locus, **(C)** Myo5 SH3 at the *MYO3* locus, and **(D)** Myo5 SH3 at the *MYO5* locus. Folding stability Gibbs Free Energy^53^ for mutations at positions which cause binding loss for all interaction partners compared to those at positions which cause binding loss only for Osh2 at the *MYO3* locus in **(E)** Myo3 SH3 (p<0.0001, two-sided Mann-Whitney U test)and **(F)** Myo5 SH3 (p<0.0001, two-sided Mann-Whitney U test).

**Supplementary Figure 9:**
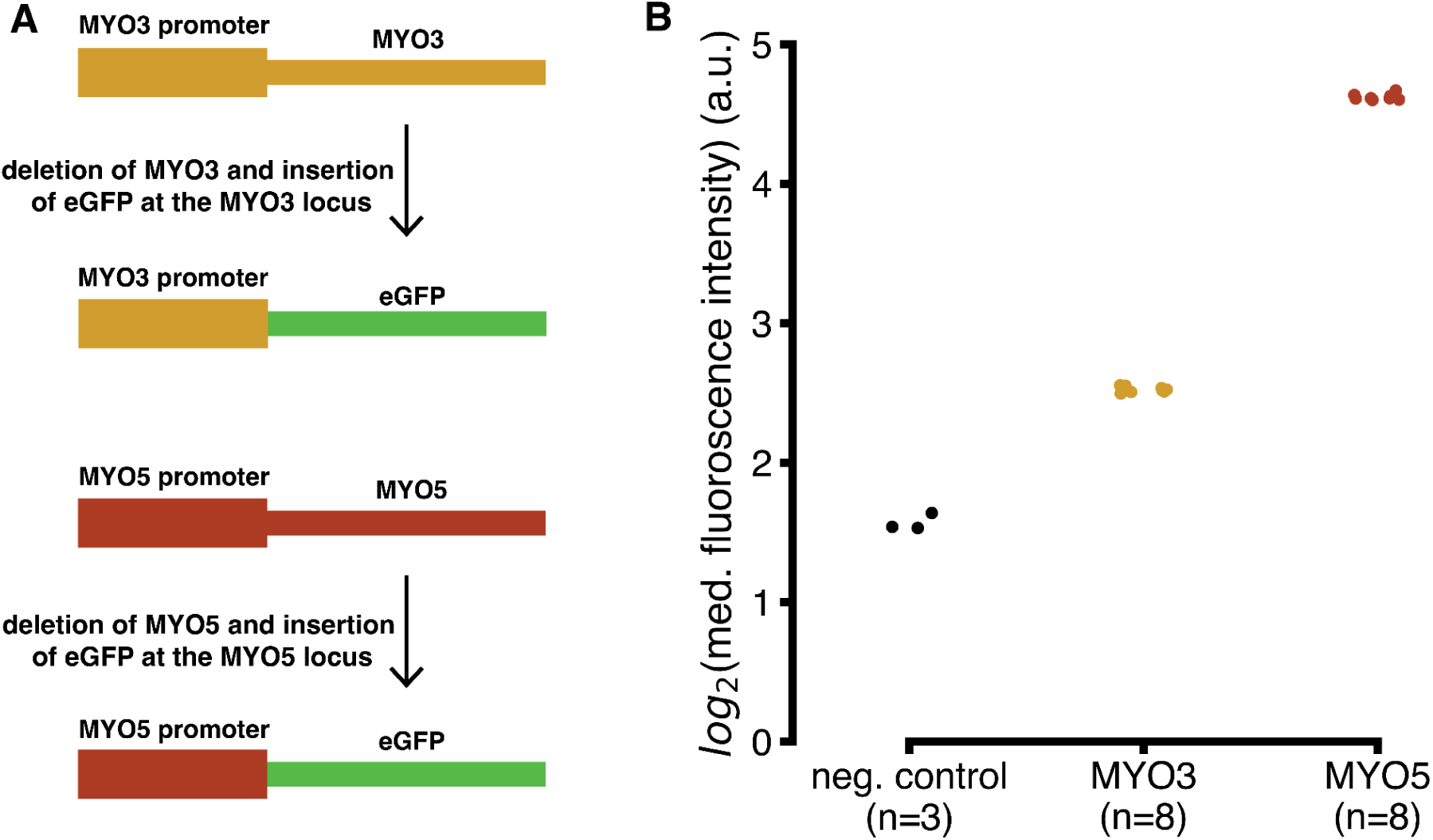
**(A)** Construction of the eGFP strains for measuring the promoter activity of the MYO3 and MYO5 loci. The entire coding sequence was replaced with the coding sequence of eGFP **(B)** log2 of median fluorescence intensity measured using a flow cytometer for negative control (n=3 biological replicates, no eGFP), MYO3 promoter-eGFP (n=8 biological replicates) and MYO5 promoter-eGFP (n=8 biological replicates) (p < 0.0001, two-sided independent samples t-test).

**Supplementary Figure 10:**
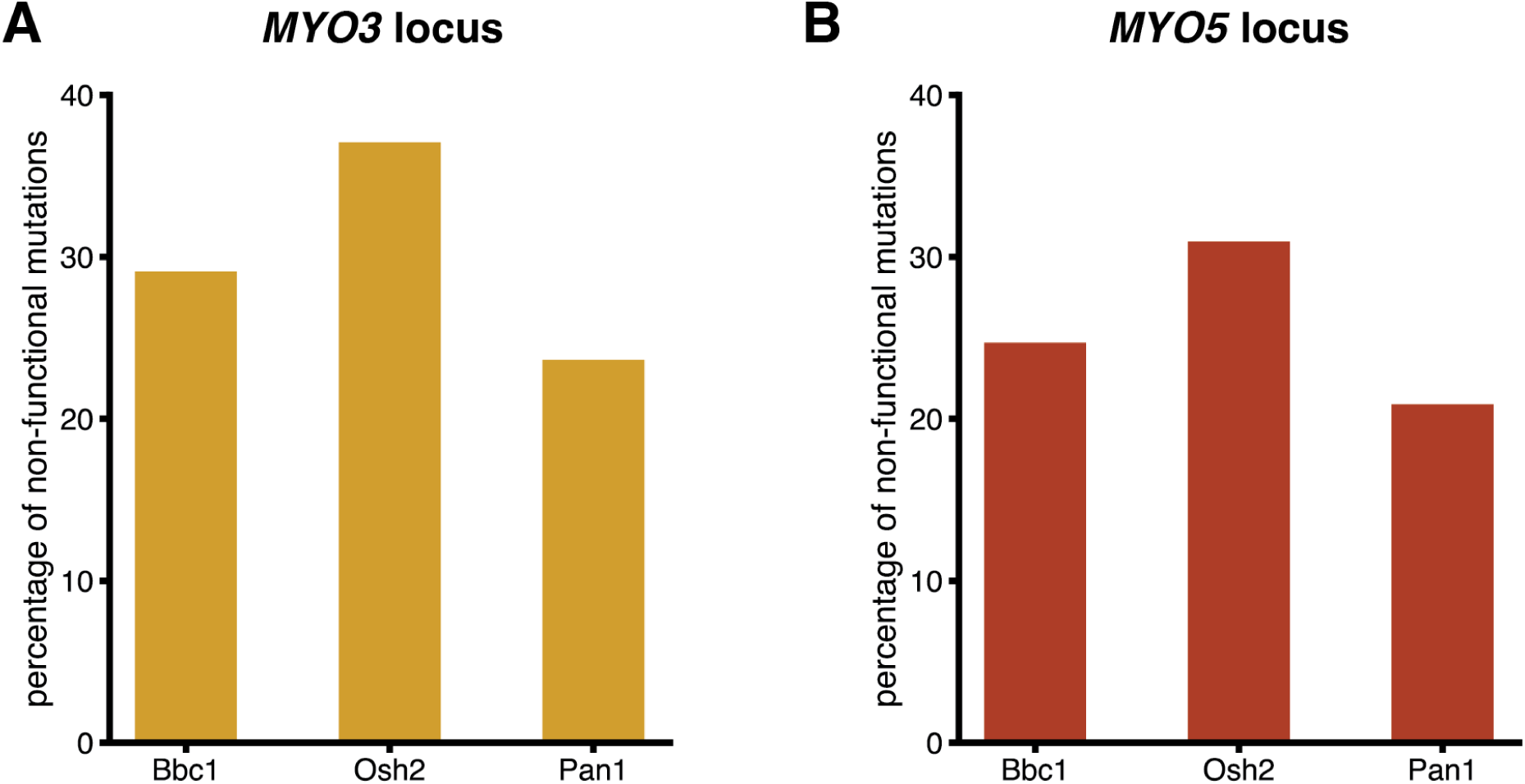
Percentage of mutations that are non-functional when introduced in both paralogous domains for any binding phenotype at the **(A)** *MYO3* locus (Bbc1 = 29%, Osh2 = 37%, Pan1 = 24%), **(B)** *MYO5* locus (Bbc1 = 25%, Osh2 = 31%, Pan1 = 21%).

**Supplementary Figure 11:**
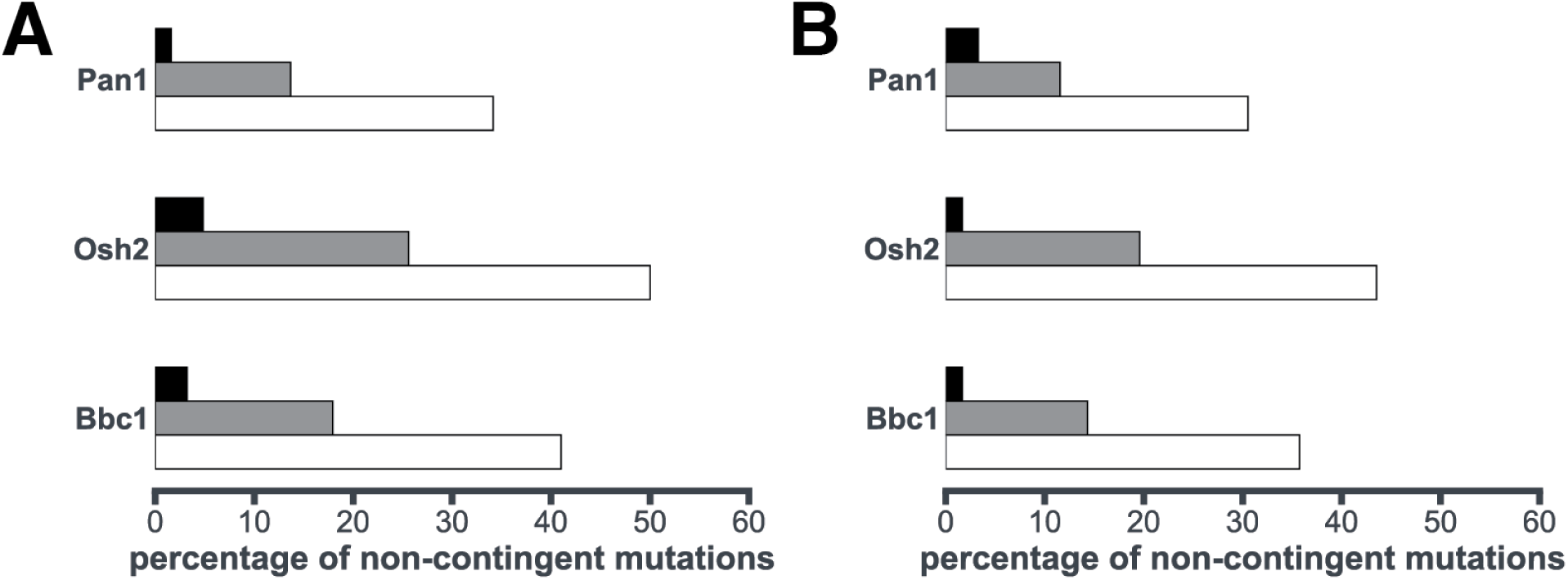
**(A)** Percentage of mutations that are non-functional in both domains in each evolutionary category at the *MYO3* locus. All pairwise comparisons are statistically significant (p < 0.05, Fisher’s exact test) [same as Figure 3E]. **(B)** Percentage of mutations that are non-functional in both domains in each evolutionary category at the *MYO5* locus. All pairwise comparisons are statistically significant (p < 0.05, Fisher’s exact test)

**Supplementary Figure 12:**
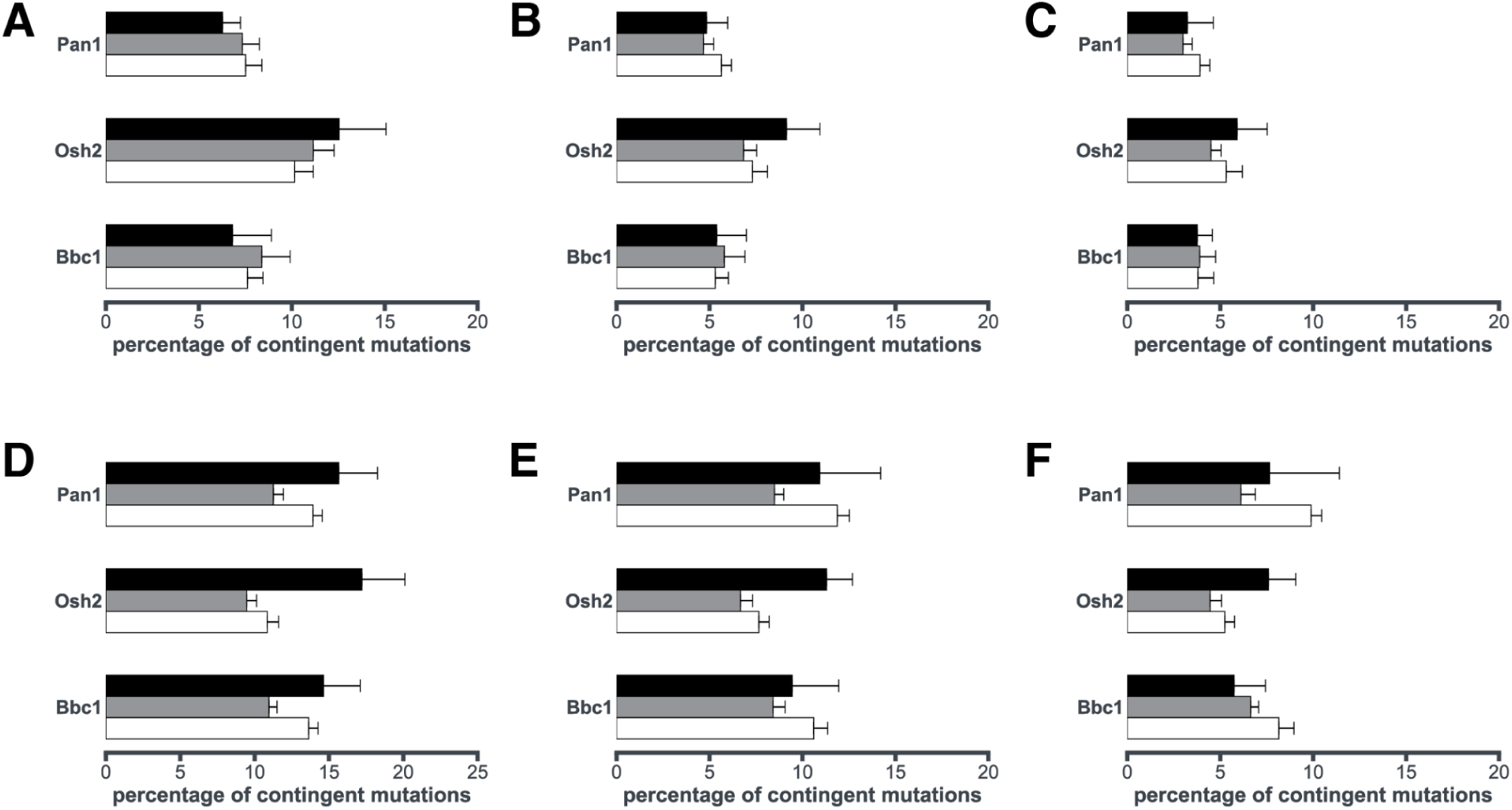
Percentage of mutations that are contingent in each evolutionary category at the *MYO3* locus for **(A)** (|ΔΔF|>0.20), all pairwise comparisons are statistically not significant (p > 0.1, Fisher’s exact test) [same as Figure 3F], **(B)** (|ΔΔF|>0.25), all pairwise comparisons are statistically not significant (p > 0.1, Fisher’s exact test), and **(C)** (|ΔΔF|>0.30), all pairwise comparisons are statistically not significant (p > 0.1, Fisher’s exact test). Percentage of mutations that are contingent in each evolutionary category at the *MYO5* locus for **(D)** (|ΔΔF|>0.20), all pairwise comparisons are statistically not significant (p > 0.1, Fisher’s exact test), **(E)** (|ΔΔF|>0.25), all pairwise comparisons are statistically not significant (p > 0.1, Fisher’s exact test), and **(F)** (|ΔΔF|>0.30), all pairwise comparisons are statistically not significant (p > 0.1, Fisher’s exact test).

**Supplementary Figure 13:**
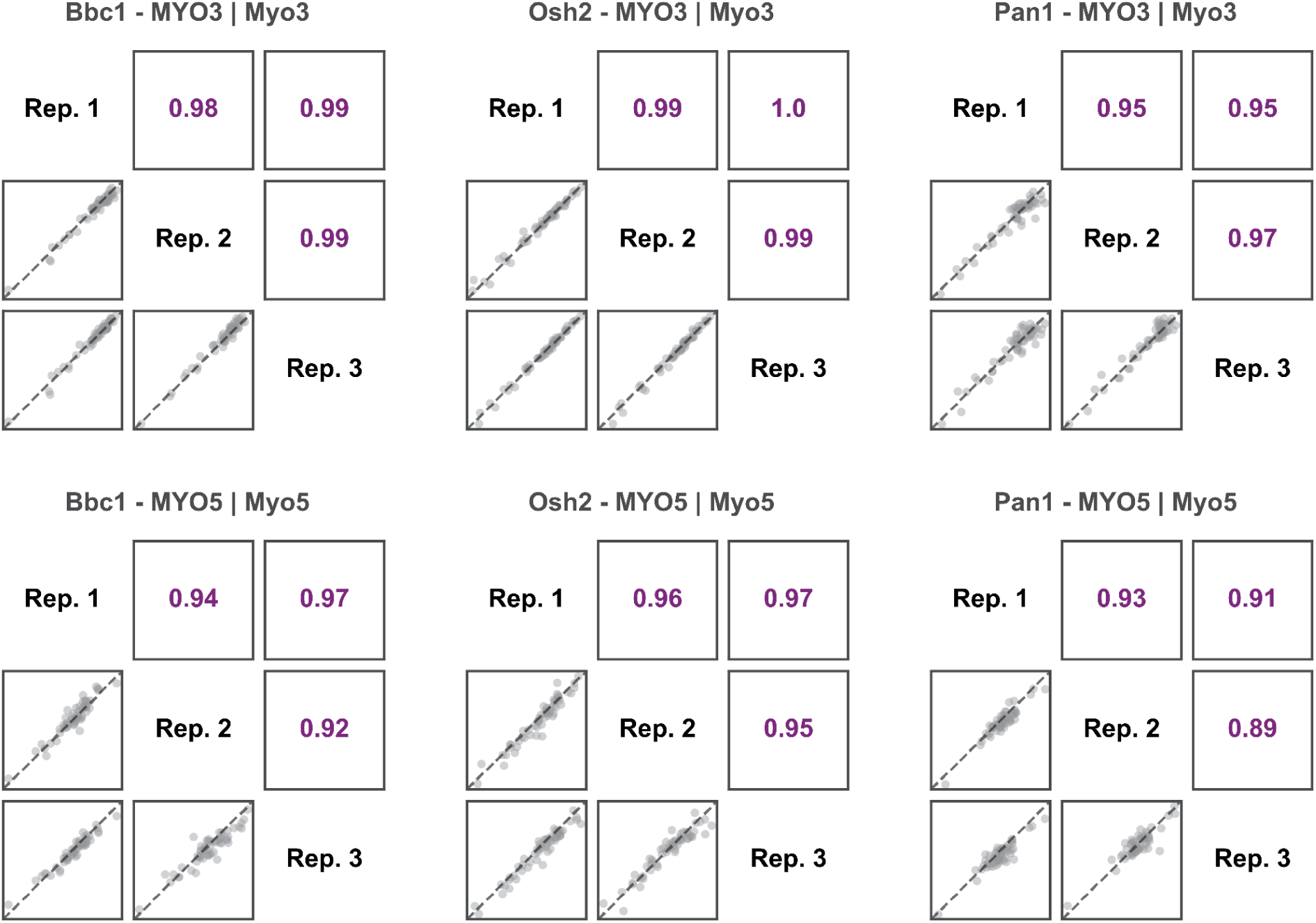
Pearson correlation between the three biological replicates for the functional scores of chimeric substitutions (n=67 for each) introduced in Myo3 SH3 domain at *MYO3* locus and Myo5 SH3 domain at *MYO5* locus for Bbc1, Osh2 and Pan1 phenotypes.

**Supplementary Figure 14:**
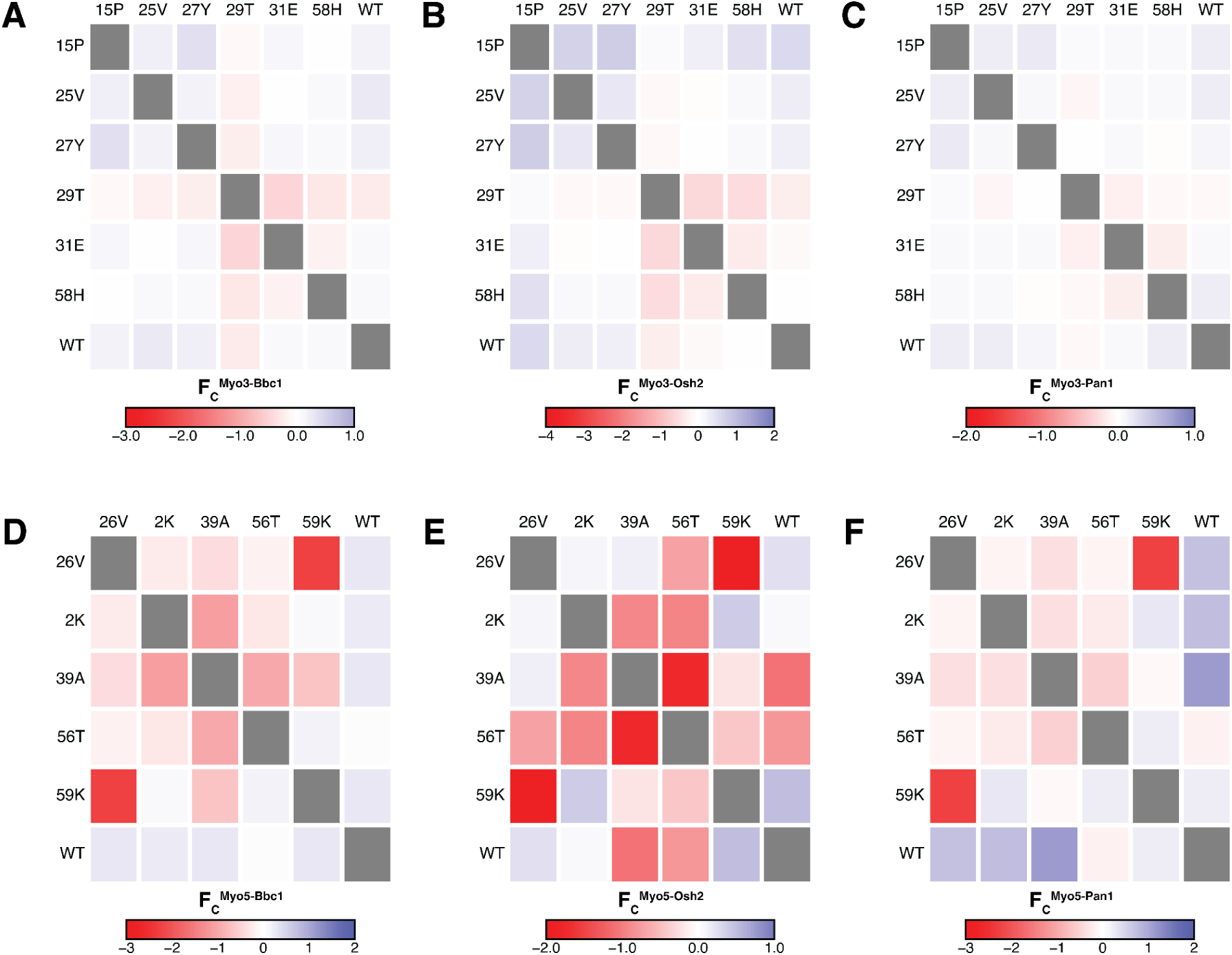
Functional scores (n=3 biological replicates) of chimeric mutants for ancestral states introduced in pairs in the Myo3 SH3 domain for **(A)** Bbc1 phenotype, **(B)** Osh2 phenotype, and **(C)** Pan1 phenotype. Functional scores (n=3 biological replicates) of chimeric mutants for ancestral states introduced in pairs in the Myo5 SH3 domain for **(D)** Bbc1 phenotype, **(E)** Osh2 phenotype, and **(F)** Pan1 phenotype.

**Supplementary Figure 15:**
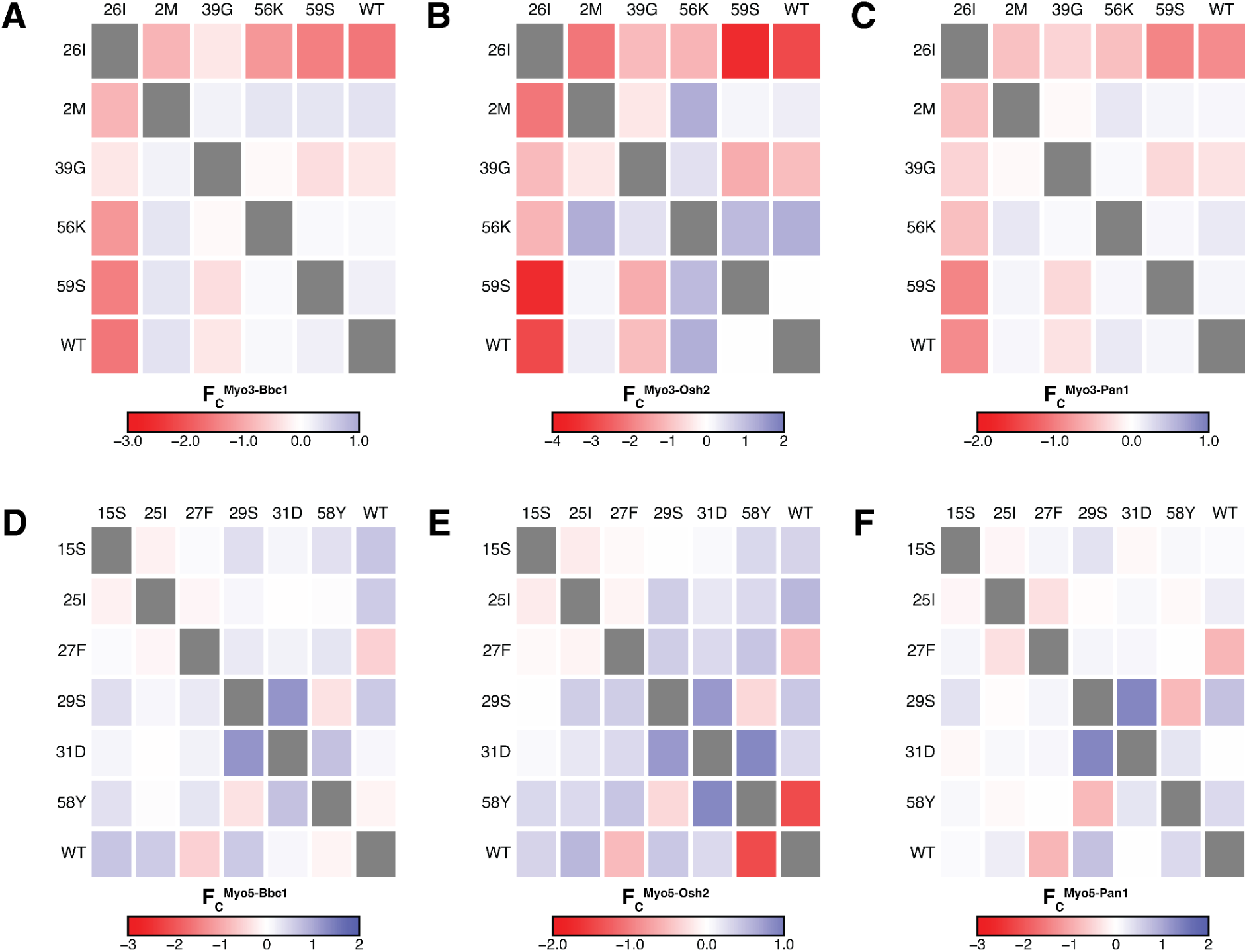
Functional scores (n=3 biological replicates) of chimeric mutants for derived states introduced in pairs in the Myo3 SH3 domain for **(A)** Bbc1 phenotype, **(B)** Osh2 phenotype, and **(C)** Pan1 phenotype. Functional scores (n=3 biological replicates) of chimeric mutants for derived states introduced in pairs in the Myo5 SH3 domain for **(D)** Bbc1 phenotype, **(E)** Osh2 phenotype, and **(F)** Pan1 phenotype.

**Supplementary Figure 16:**
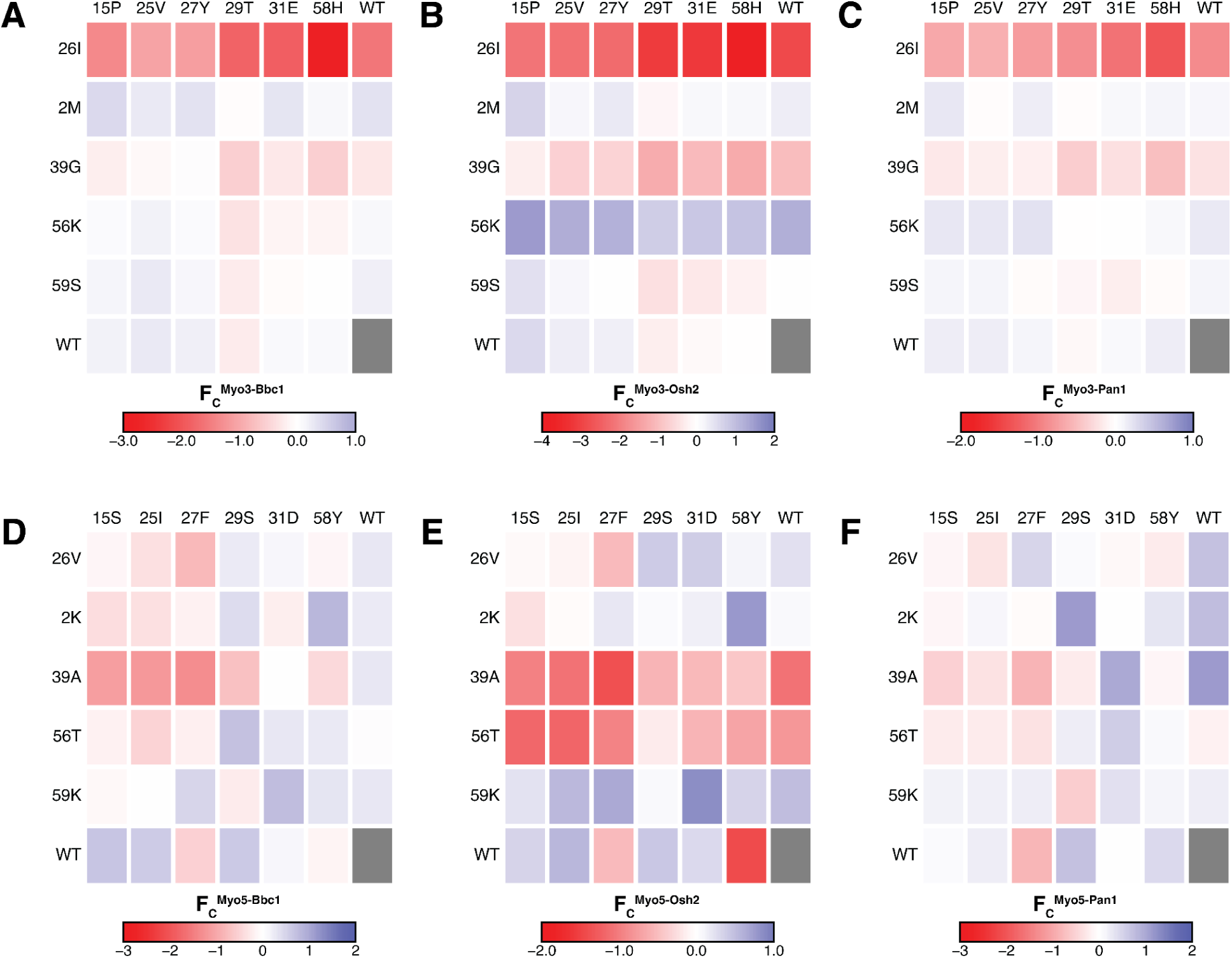
Functional scores (n=3 biological replicates) of chimeric mutants for an ancestral state and a derived state introduced together in the Myo3 SH3 domain for **(A)** Bbc1 phenotype, **(B)** Osh2 phenotype, and **(C)** Pan1 phenotype.Functional scores (n=3 biological replicates) of chimeric mutants for an ancestral state and a derived state introduced together in the Myo5 SH3 domain for **(D)** Bbc1 phenotype, **(E)** Osh2 phenotype, and **(F)** Pan1 phenotype.

**Supplementary Figure 17:**
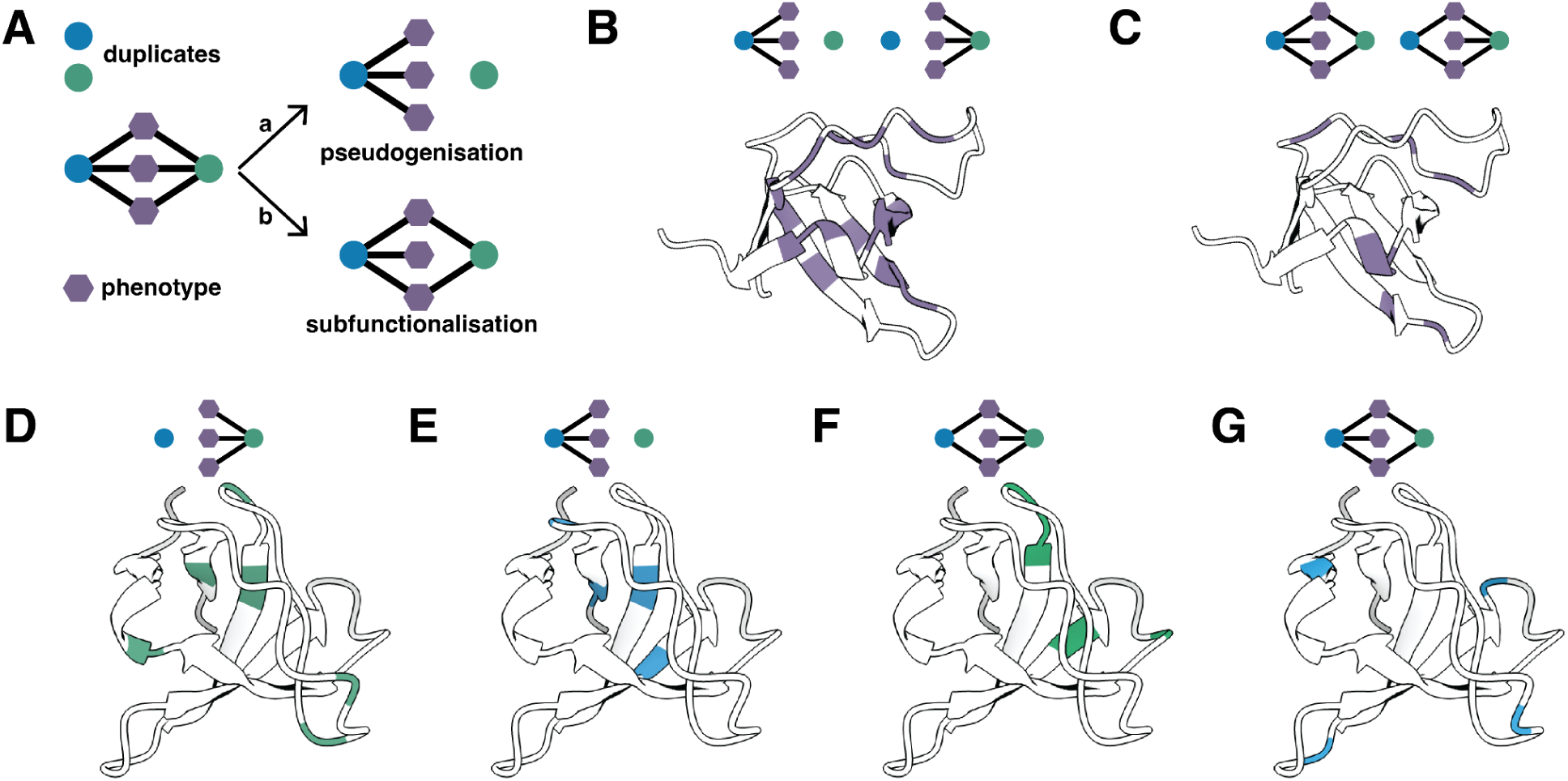
**(A)** Long-term fates of gene duplicates in a protein interaction network: **(a)** pseudogenisation and **(b)** subfunctionalisation. **(B)** The marked positions (PDB: 1RUW) mark the enrichment of mutations, which can lead to pseudogenisation irrespective of the duplicate. **(C)** The marked positions on the structure mark the enrichment of mutations, which can lead to subfunctionalisation irrespective of the duplicate. **(D)** The marked positions mark the enrichment of mutations, which can lead to pseudogenisation when introduced in the Myo3 SH3 domain but not the Myo5 SH3 domain. **(E)** The marked positions mark the enrichment of mutations, which can lead to pseudogenisation when introduced in the Myo5 SH3 domain but not the Myo3 SH3 domain. **(F)** The marked positions mark the enrichment of mutations, which can lead to subfunctionalisation when introduced in the Myo3 SH3 domain but not the Myo5 SH3 domain. **(G)** The marked positions mark the enrichment of mutations, which can lead to subfunctionalisation when introduced in the Myo5 SH3 domain but not the Myo3 SH3 domain.

