## Supplemental File 5 for "Cryptic genetic variation shapes the fate of gene duplicates in a protein interaction network"

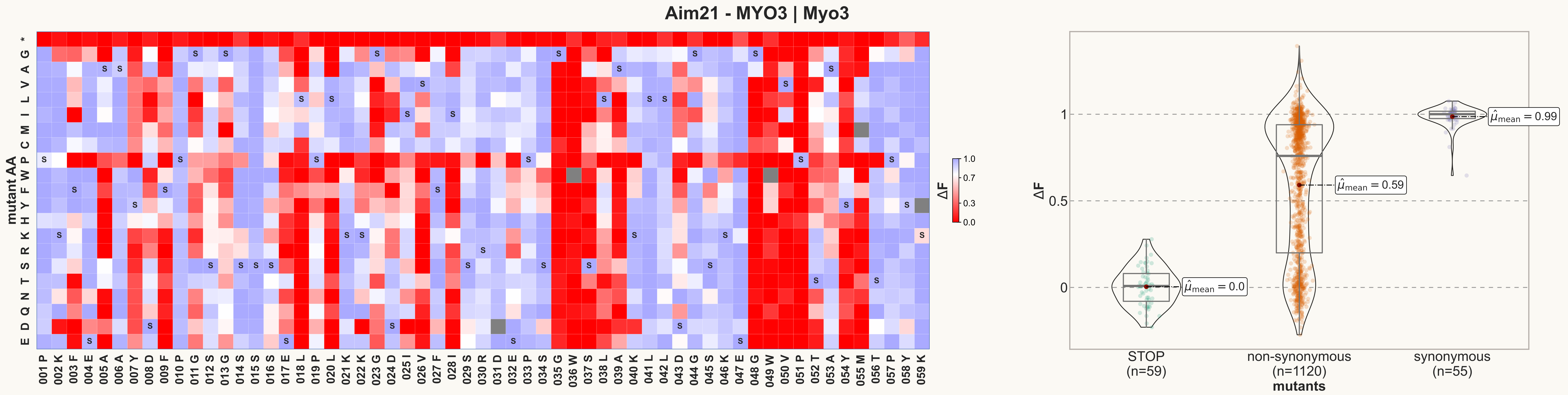

Aim21 - MYO3 | Myo3
mutant AA
ΔF
STOP
(n=59)
non-synonymous
(n=1120)
synonymous
(n=55)
mutants
$\hat{\mu}_{mean} = 0.0$
$\hat{\mu}_{mean} = 0.59$
$\hat{\mu}_{mean} = 0.99$

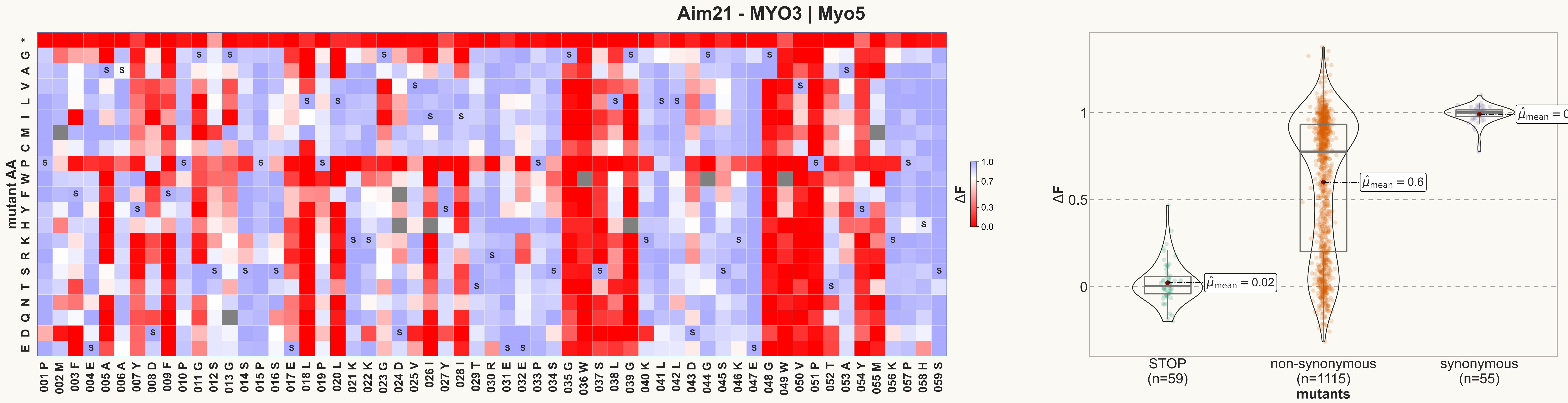

Aim21 - MYO3 | Myo5
mutant AA
ΔF
1.0
0.7
0.3
0.0
STOP
(n=59)
non-synonymous
(n=1115)
synonymous
(n=55)
mutants
$\hat{\mu}_{mean} = 0.02$
$\hat{\mu}_{mean} = 0.6$
$\hat{\mu}_{mean} = 0.99$

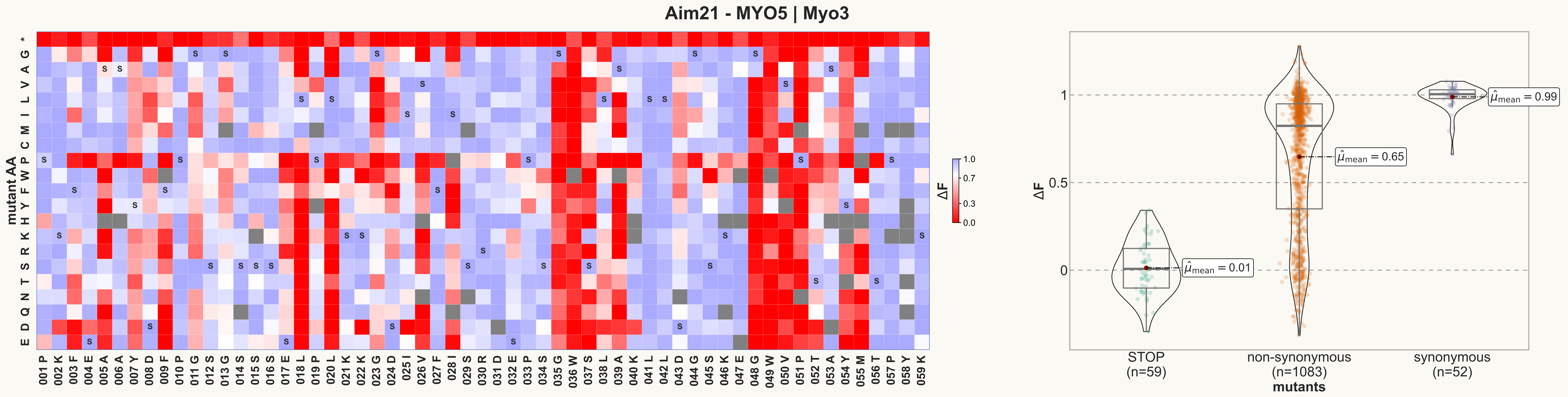
Aim21 - MYO5 | Myo3
mutant AA
* G A V L I M C P W F Y H K R S T N Q D E
001 P 002 K 003 F 004 E 005 A 006 A 007 Y 008 D 009 F 010 P 011 G 012 S 013 G 014 S 015 S 016 S 017 E 018 L 019 P 020 L 021 K 022 K 023 G 024 D 025 I 026 V 027 F 028 I 029 S 030 R 031 D 032 E 033 P 034 S 035 G 036 W 037 S 038 L 039 A 040 K 041 L 042 L 043 D 044 G 045 S 046 K 047 E 048 G 049 W 050 V 051 P 052 T 053 A 054 Y 055 M 056 T 057 P 058 Y 059 K
ΔF
1.0
0.7
0.3
0.0
1
0.5
0
$\hat{\mu}_{mean} = 0.01$
$\hat{\mu}_{mean} = 0.65$
$\hat{\mu}_{mean} = 0.99$
STOP (n=59)
non-synonymous (n=1083)
synonymous (n=52)
mutants

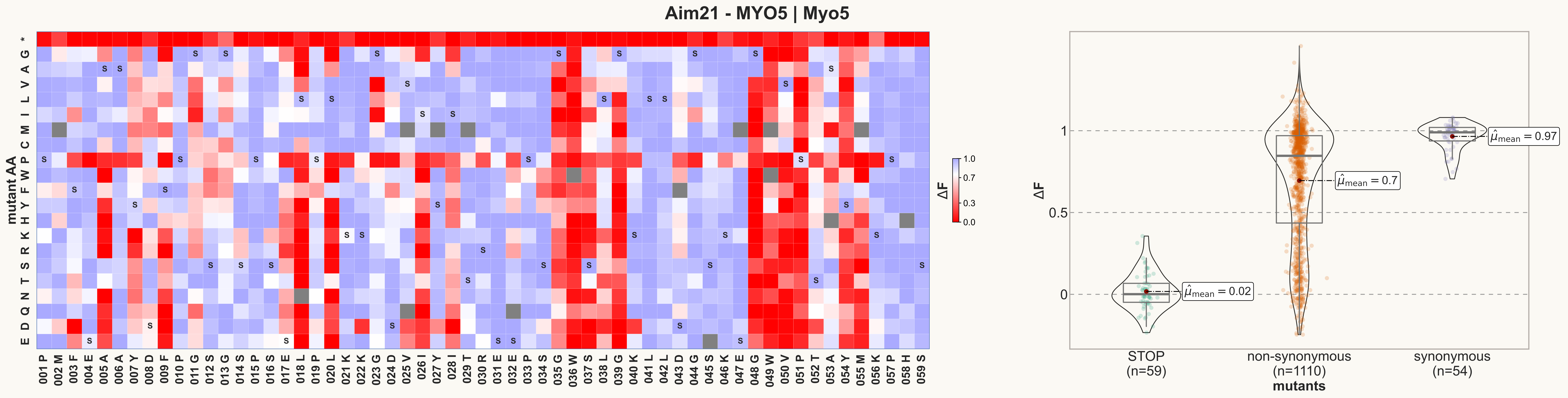
Aim21 - MYO5 | Myo5
mutant AA
ΔF
1.0
0.7
0.3
0.0
001 P
002 M
003 F
004 E
005 A
006 A
007 Y
008 D
009 F
010 P
011 G
012 S
013 G
014 S
015 P
016 S
017 E
018 L
019 P
020 L
021 K
022 K
023 G
024 D
025 V
026 I
027 Y
028 I
029 T
030 R
031 E
032 E
033 P
034 S
035 G
036 W
037 S
038 L
039 G
040 K
041 L
042 L
043 D
044 G
045 S
046 K
047 E
048 G
049 W
050 V
051 P
052 T
053 A
054 Y
055 M
056 K
057 P
058 H
059 S
$\hat{\mu}_{mean} = 0.02$
$\hat{\mu}_{mean} = 0.7$
$\hat{\mu}_{mean} = 0.97$
STOP
(n=59)
non-synonymous
(n=1110)
synonymous
(n=54)
mutants

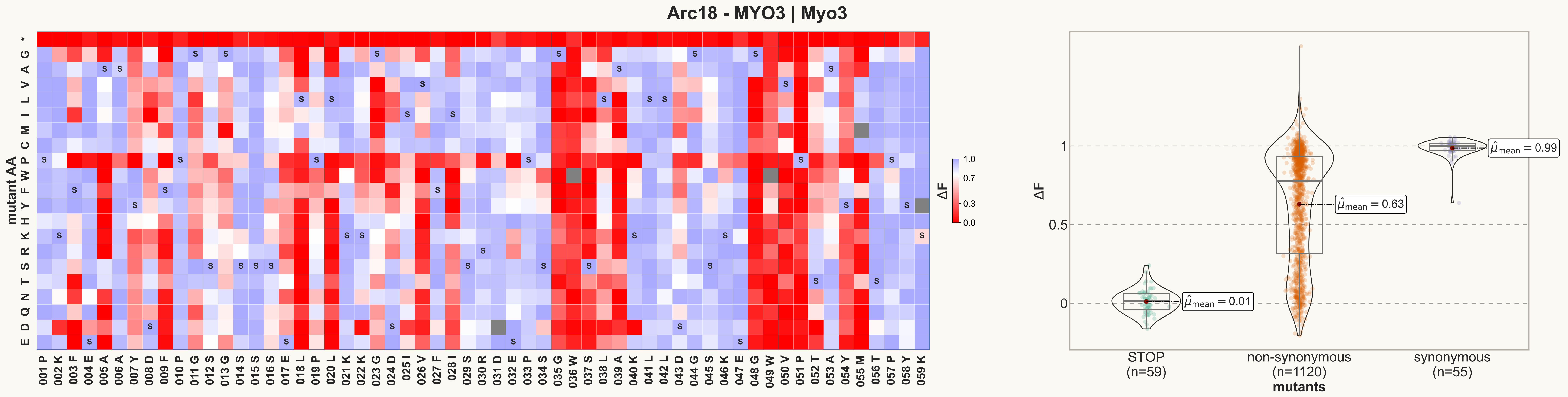
Arc18 - MYO3 | Myo3
mutant AA
* G A V L I M C P W F Y H K R S T N Q D E
001 P 002 K 003 F 004 E 005 A 006 A 007 Y 008 D 009 F 010 P 011 G 012 S 013 G 014 S 015 S 016 S 017 E 018 L 019 P 020 L 021 K 022 K 023 G 024 D 025 I 026 V 027 F 028 I 029 S 030 R 031 D 032 E 033 P 034 S 035 G 036 W 037 S 038 L 039 A 040 K 041 L 042 L 043 D 044 G 045 S 046 K 047 E 048 G 049 W 050 V 051 P 052 T 053 A 054 Y 055 M 056 T 057 P 058 Y 059 K
ΔF
1.0
0.7
0.3
0.0
ΔF
1
0.5
0
$\hat{\mu}_{\mathrm{mean}} = 0.01$
$\hat{\mu}_{\mathrm{mean}} = 0.63$
$\hat{\mu}_{\mathrm{mean}} = 0.99$
STOP
(n=59)
non-synonymous
(n=1120)
synonymous
(n=55)
mutants

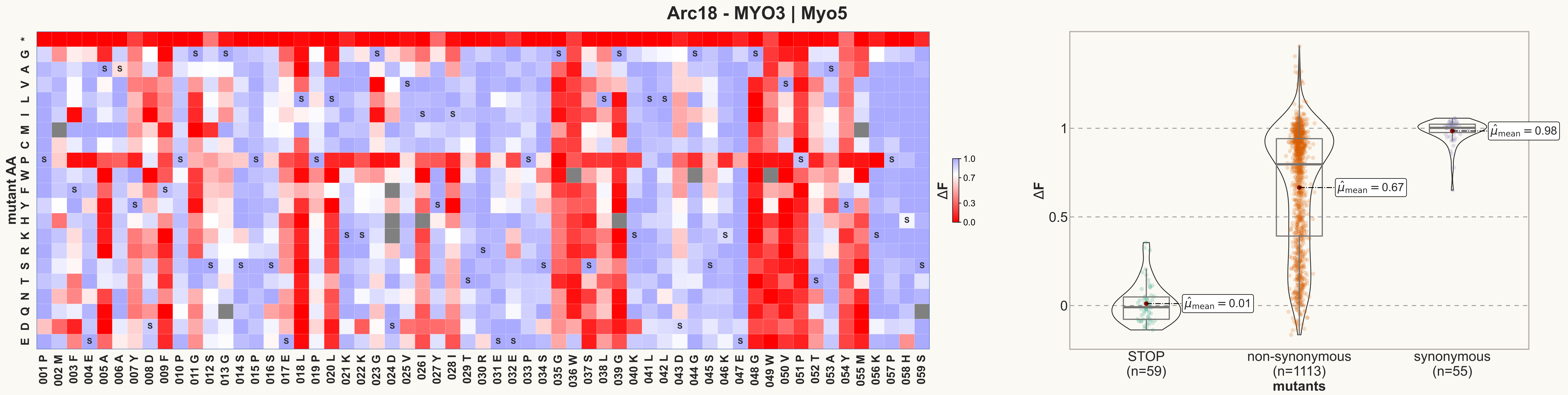
Arc18 - MYO3 | Myo5
mutant AA
ΔF
STOP
(n=59)
non-synonymous
(n=1113)
synonymous
(n=55)
mutants
$\hat{\mu}_{mean}$ = 0.01
$\hat{\mu}_{mean}$ = 0.67
$\hat{\mu}_{mean}$ = 0.98

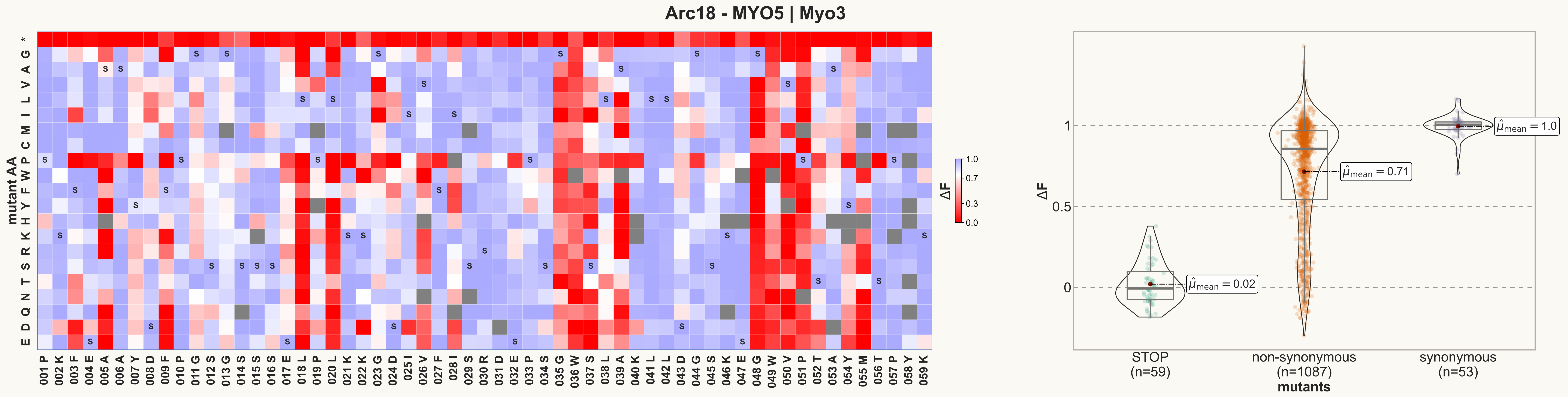

Arc18 - MYO5 | Myo3
mutant AA
* G A V L I M C P W F Y H K R S T N Q D E
001 P 002 K 003 F 004 E 005 A 006 A 007 Y 008 D 009 F 010 P 011 G 012 S 013 G 014 S 015 S 016 S 017 E 018 L 019 P 020 L 021 K 022 K 023 G 024 D 025 I 026 V 027 F 028 I 029 S 030 R 031 D 032 E 033 P 034 S 035 G 036 W 037 S 038 L 039 A 040 K 041 L 042 L 043 D 044 G 045 S 046 K 047 E 048 G 049 W 050 V 051 P 052 T 053 A 054 Y 055 M 056 T 057 P 058 Y 059 K
ΔF
1.0
0.7
0.3
0.0
ΔF
1
0.5
0
$\hat{\mu}_{mean} = 0.02$
$\hat{\mu}_{mean} = 0.71$
$\hat{\mu}_{mean} = 1.0$
STOP (n=59)
non-synonymous (n=1087)
synonymous (n=53)
mutants

### Arc18 - MYO5 | Myo5

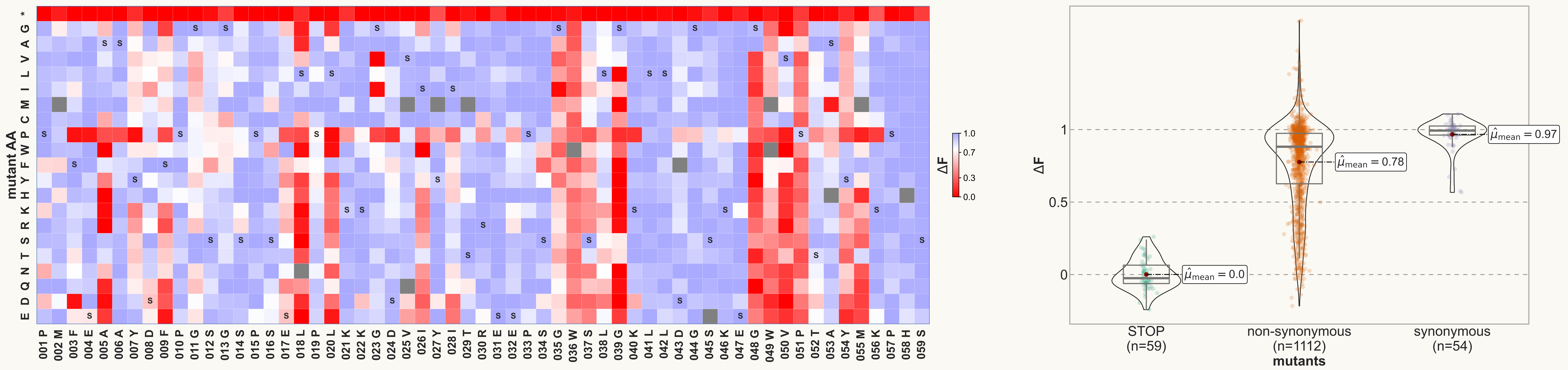

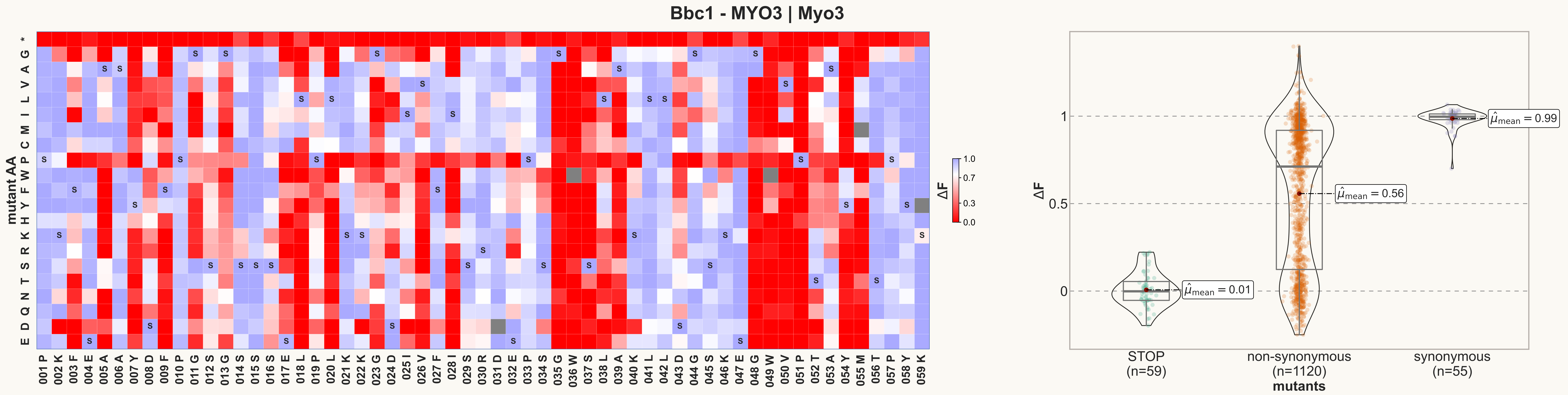
Bbc1 - MYO3 | Myo3
mutant AA
* G A V L I M C P W F Y H K R S T N Q D E
001 P 002 K 003 F 004 E 005 A 006 A 007 Y 008 D 009 F 010 P 011 G 012 S 013 G 014 S 015 S 016 S 017 E 018 L 019 P 020 L 021 K 022 K 023 G 024 D 025 I 026 V 027 F 028 I 029 S 030 R 031 D 032 E 033 P 034 S 035 G 036 W 037 S 038 L 039 A 040 K 041 L 042 L 043 D 044 G 045 S 046 K 047 E 048 G 049 W 050 V 051 P 052 T 053 A 054 Y 055 M 056 T 057 P 058 Y 059 K
ΔF
1.0
0.7
0.3
0.0
ΔF
1
0.5
0
$\hat{\mu}_{mean} = 0.99$
$\hat{\mu}_{mean} = 0.56$
$\hat{\mu}_{mean} = 0.01$
STOP (n=59)
non-synonymous (n=1120)
synonymous (n=55)
mutants

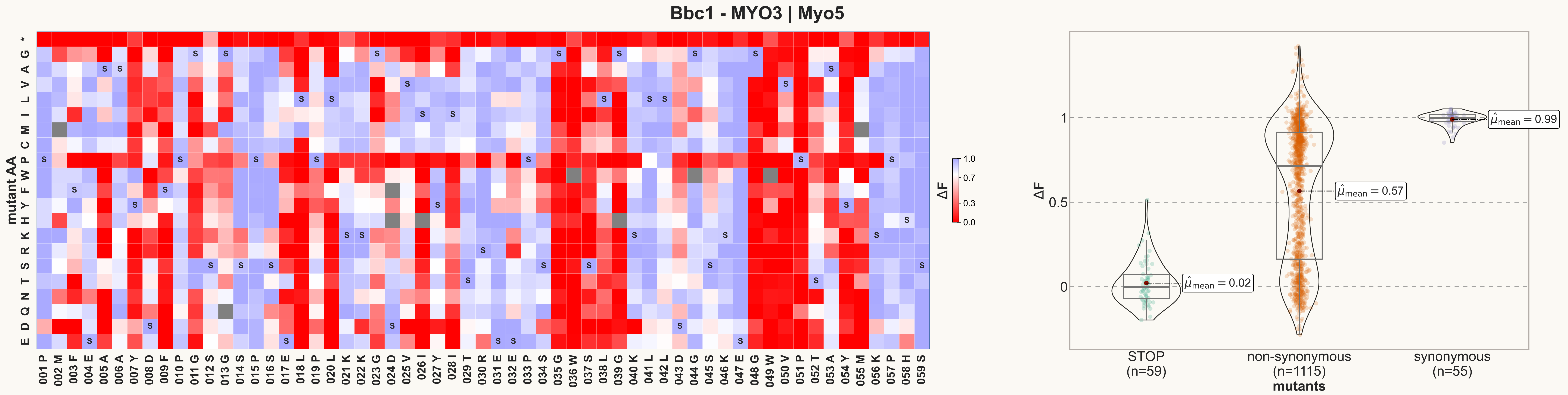

Bbc1 - MYO3 | Myo5
mutant AA
* G A V L I M C P W F Y H K R S T N Q D E
001 P 002 M 003 F 004 E 005 A 006 A 007 Y 008 D 009 F 010 P 011 G 012 S 013 G 014 S 015 P 016 S 017 E 018 L 019 P 020 L 021 K 022 K 023 G 024 D 025 V 026 I 027 Y 028 I 029 T 030 R 031 E 032 E 033 P 034 S 035 G 036 W 037 S 038 L 039 G 040 K 041 L 042 L 043 D 044 G 045 S 046 K 047 E 048 G 049 W 050 V 051 P 052 T 053 A 054 Y 055 M 056 K 057 P 058 H 059 S
ΔF
1.0
0.7
0.3
0.0
ΔF
1
0.5
0
$\hat{\mu}_{mean} = 0.02$
$\hat{\mu}_{mean} = 0.57$
$\hat{\mu}_{mean} = 0.99$
STOP (n=59)
non-synonymous (n=1115)
synonymous (n=55)
mutants

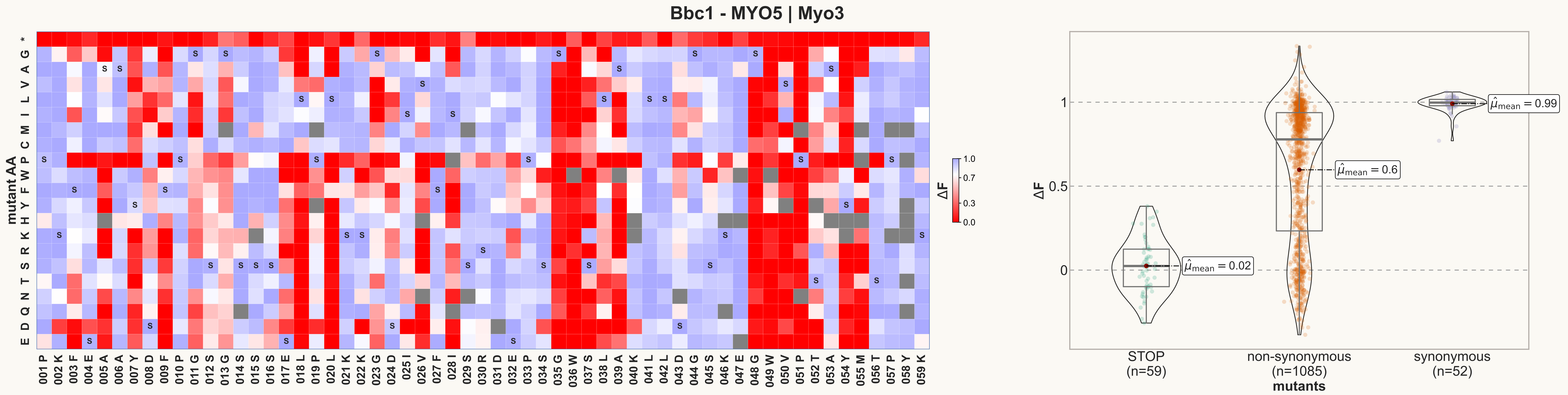
Bbc1 - MYO5 | Myo3
mutant AA
* G A V L I M C P W F Y H K R S T N Q D E
001 P 002 K 003 F 004 E 005 A 006 A 007 Y 008 D 009 F 010 P 011 G 012 S 013 G 014 S 015 S 016 S 017 E 018 L 019 P 020 L 021 K 022 K 023 G 024 D 025 I 026 V 027 F 028 I 029 S 030 R 031 D 032 E 033 P 034 S 035 G 036 W 037 S 038 L 039 A 040 K 041 L 042 L 043 D 044 G 045 S 046 K 047 E 048 G 049 W 050 V 051 P 052 T 053 A 054 Y 055 M 056 T 057 P 058 Y 059 K
ΔF
1.0
0.7
0.3
0.0
$\hat{\mu}_{mean} = 0.02$
$\hat{\mu}_{mean} = 0.6$
$\hat{\mu}_{mean} = 0.99$
1
0.5
0
STOP (n=59)
non-synonymous (n=1085)
synonymous (n=52)
mutants

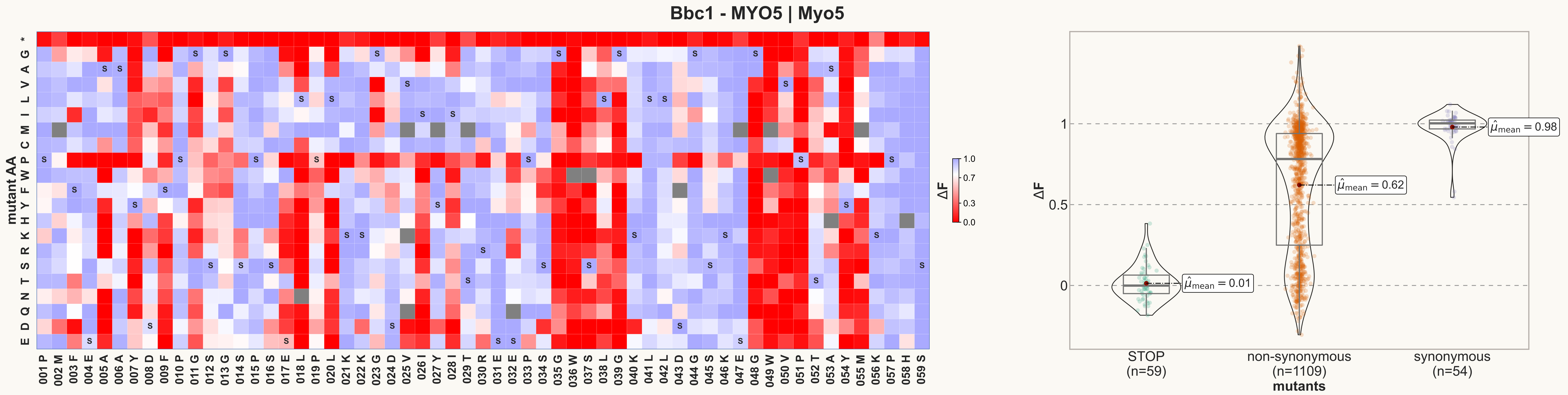

Bbc1 - MYO5 | Myo5
mutant AA
ΔF
mutants
STOP (n=59)
non-synonymous (n=1109)
synonymous (n=54)
$\hat{\mu}_{mean} = 0.01$
$\hat{\mu}_{mean} = 0.62$
$\hat{\mu}_{mean} = 0.98$

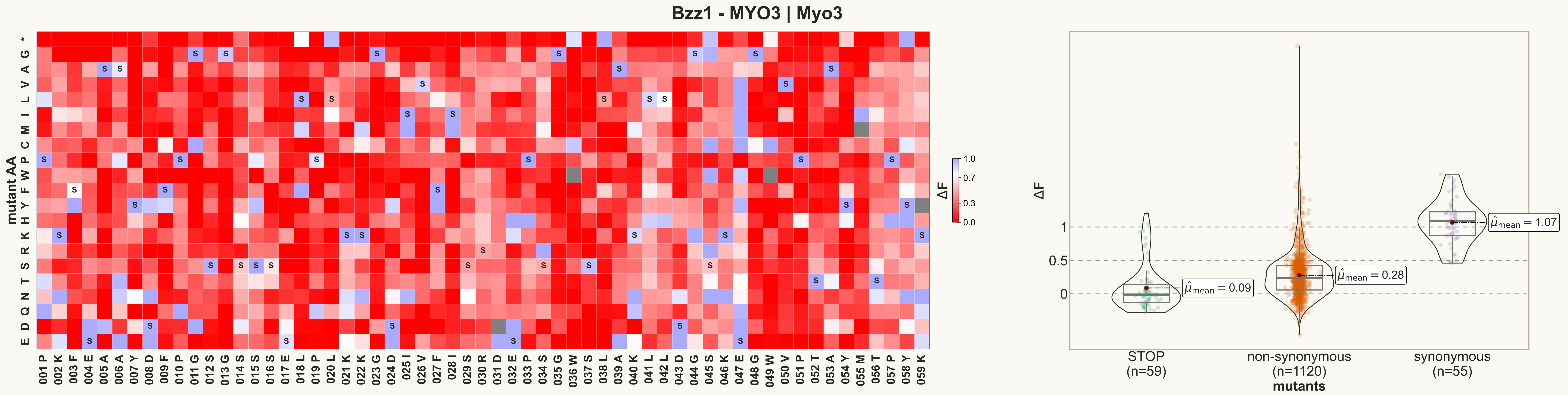

Bzz1 - MYO3 | Myo3
mutant AA
ΔF
STOP (n=59)
non-synonymous (n=1120)
synonymous (n=55)
mutants
$\hat{\mu}_{mean} = 0.09$
$\hat{\mu}_{mean} = 0.28$
$\hat{\mu}_{mean} = 1.07$

#### Bzz1 - MYO3 | Myo5

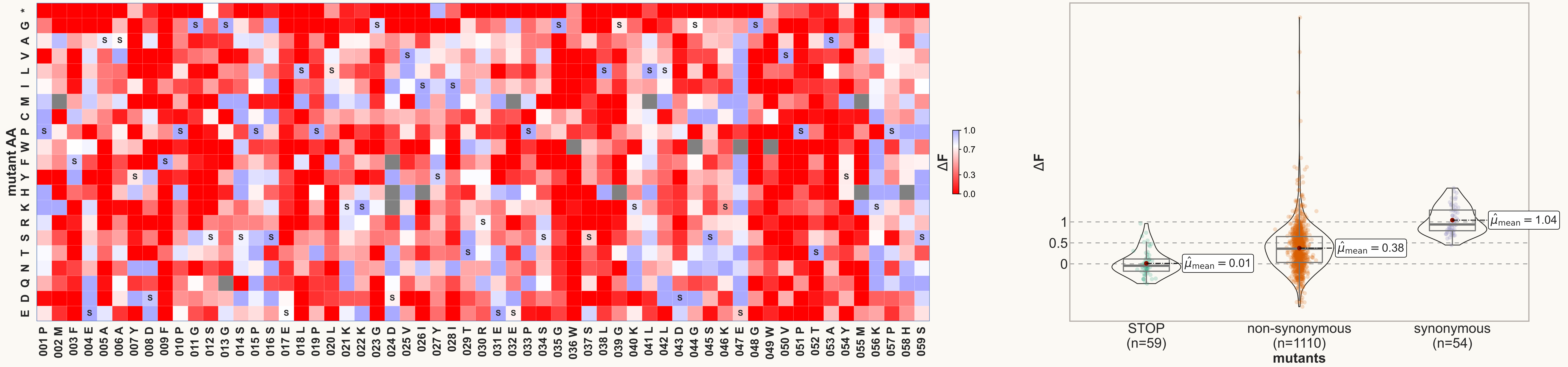

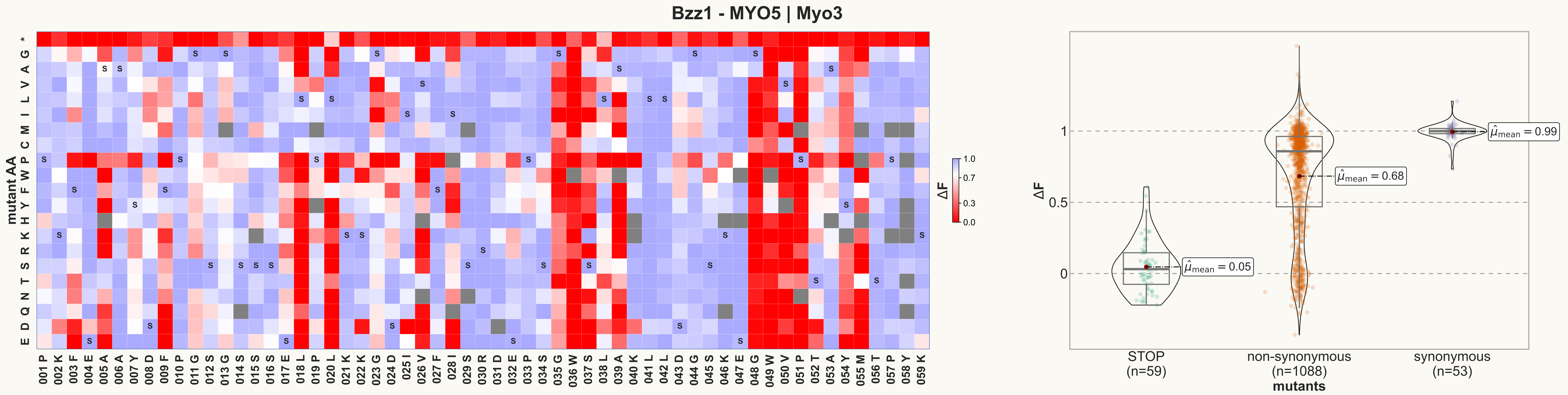

Bzz1 - MYO5 | Myo3
mutant AA
ΔF
1.0
0.7
0.3
0.0
STOP
(n=59)
non-synonymous
(n=1088)
synonymous
(n=53)
mutants
$\hat{\mu}_{mean} = 0.05$
$\hat{\mu}_{mean} = 0.68$
$\hat{\mu}_{mean} = 0.99$

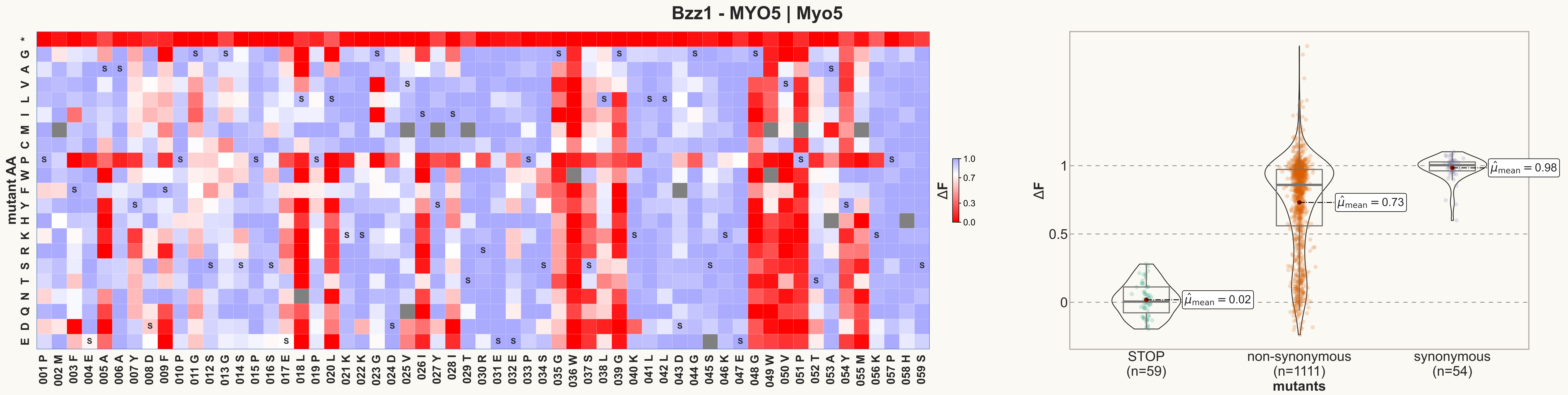

Bzz1 - MYO5 | Myo5
mutant AA
* G A V L I M C P W F Y H K R S T N Q D E
001 P 002 M 003 F 004 E 005 A 006 A 007 Y 008 D 009 F 010 P 011 G 012 S 013 G 014 S 015 P 016 S 017 E 018 L 019 P 020 L 021 K 022 K 023 G 024 D 025 V 026 I 027 Y 028 I 029 T 030 R 031 E 032 E 033 P 034 S 035 G 036 W 037 S 038 L 039 G 040 K 041 L 042 L 043 D 044 G 045 S 046 K 047 E 048 G 049 W 050 V 051 P 052 T 053 A 054 Y 055 M 056 K 057 P 058 H 059 S
ΔF
1.0
0.7
0.3
0.0
ΔF
1
0.5
0
$\hat{\mu}_{mean}$ = 0.02
$\hat{\mu}_{mean}$ = 0.73
$\hat{\mu}_{mean}$ = 0.98
STOP (n=59)
non-synonymous (n=1111)
synonymous (n=54)
mutants

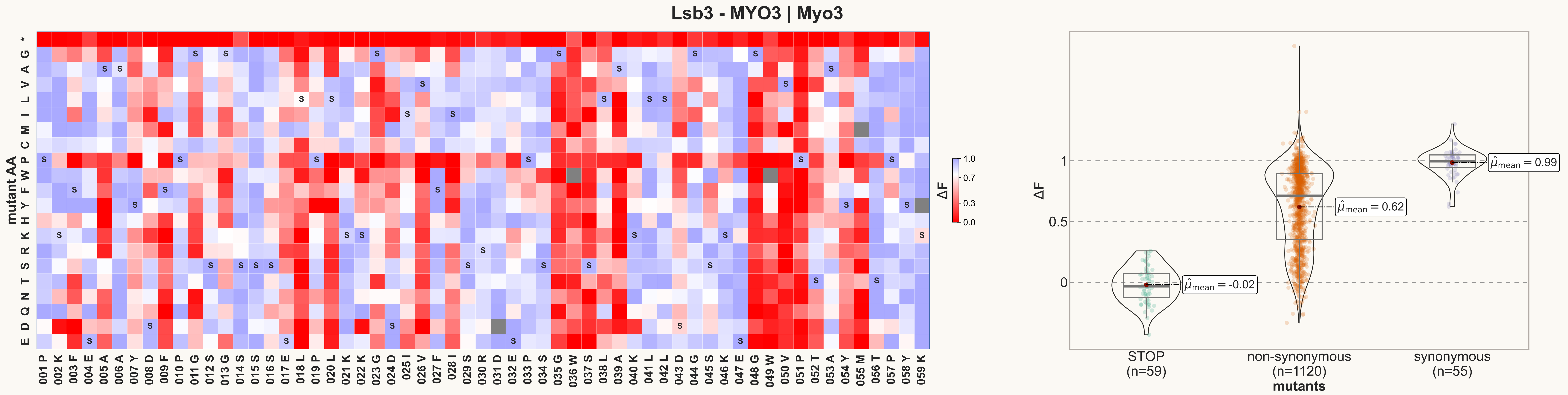
Lsb3 - MYO3 | Myo3
mutant AA
ΔF
mutants
STOP (n=59)
non-synonymous (n=1120)
synonymous (n=55)
$\hat{\mu}_{mean}$ = -0.02
$\hat{\mu}_{mean}$ = 0.62
$\hat{\mu}_{mean}$ = 0.99

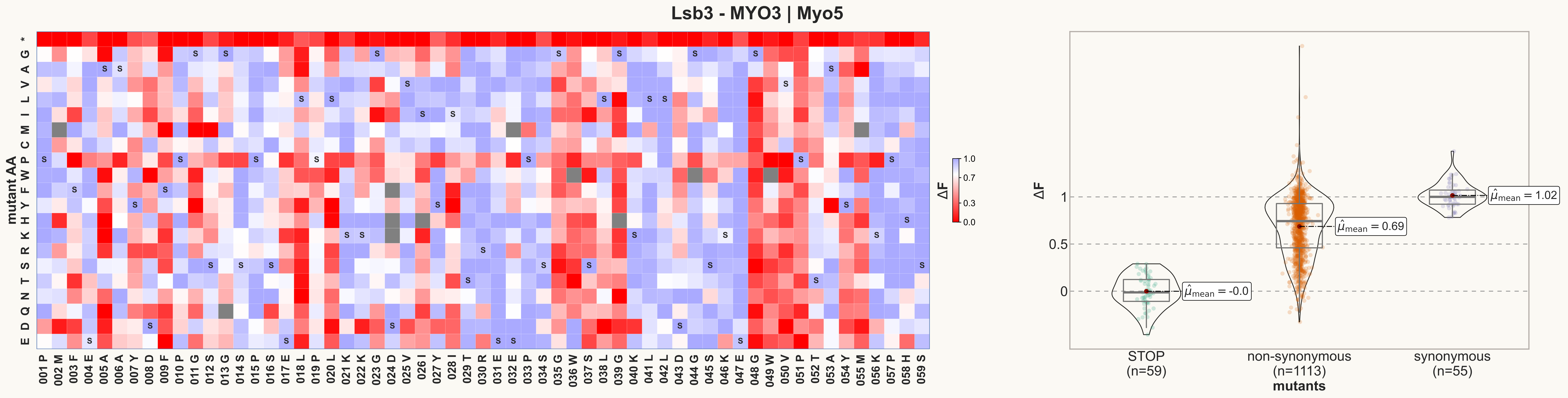

Lsb3 - MYO3 | Myo5
mutant AA
* G A V L I M C P W F Y H K R S T N Q D E
001 P 002 M 003 F 004 E 005 A 006 A 007 Y 008 D 009 F 010 P 011 G 012 S 013 G 014 S 015 P 016 S 017 E 018 L 019 P 020 L 021 K 022 K 023 G 024 D 025 V 026 I 027 Y 028 I 029 T 030 R 031 E 032 E 033 P 034 S 035 G 036 W 037 S 038 L 039 G 040 K 041 L 042 L 043 D 044 G 045 S 046 K 047 E 048 G 049 W 050 V 051 P 052 T 053 A 054 Y 055 M 056 K 057 P 058 H 059 S
ΔF
1.0
0.7
0.3
0.0
ΔF
1
0.5
0
$\hat{\mu}_{mean}$ = -0.0
$\hat{\mu}_{mean}$ = 0.69
$\hat{\mu}_{mean}$ = 1.02
STOP (n=59)
non-synonymous (n=1113)
synonymous (n=55)
mutants

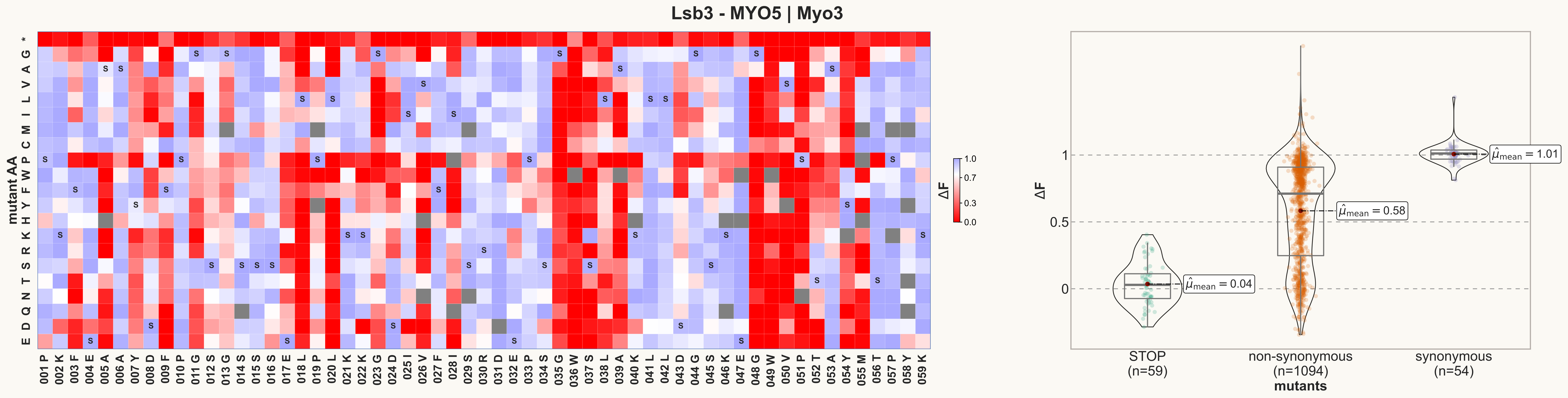
Lsb3 - MYO5 | Myo3
mutant AA
* G A V L I M C P W F Y H K R S T N Q D E
001 P 002 K 003 F 004 E 005 A 006 A 007 Y 008 D 009 F 010 P 011 G 012 S 013 G 014 S 015 S 016 S 017 E 018 L 019 P 020 L 021 K 022 K 023 G 024 D 025 I 026 V 027 F 028 I 029 S 030 R 031 D 032 E 033 P 034 S 035 G 036 W 037 S 038 L 039 A 040 K 041 L 042 L 043 D 044 G 045 S 046 K 047 E 048 G 049 W 050 V 051 P 052 T 053 A 054 Y 055 M 056 T 057 P 058 Y 059 K
ΔF
1.0
0.7
0.3
0.0
ΔF
1
0.5
0
$\hat{\mu}_{mean} = 0.04$
$\hat{\mu}_{mean} = 0.58$
$\hat{\mu}_{mean} = 1.01$
STOP (n=59)
non-synonymous (n=1094)
synonymous (n=54)
mutants

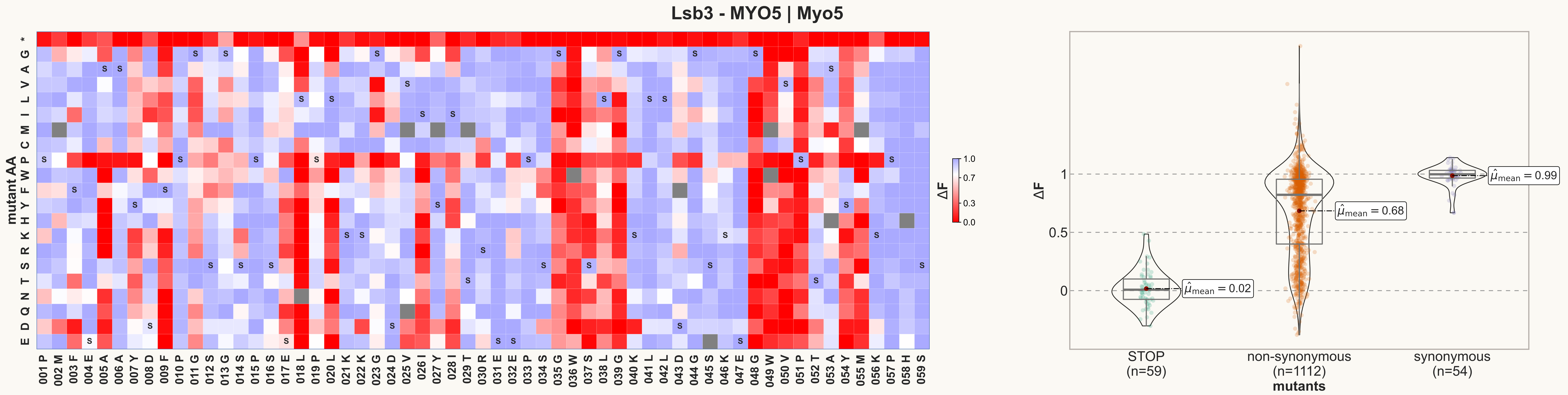

Lsb3 - MYO5 | Myo5
mutant AA
* G A V L I M C P W F Y H K R S T N Q D E
001 P 002 M 003 F 004 E 005 A 006 A 007 Y 008 D 009 F 010 P 011 G 012 S 013 G 014 S 015 P 016 S 017 E 018 L 019 P 020 L 021 K 022 K 023 G 024 D 025 V 026 I 027 Y 028 I 029 T 030 R 031 E 032 E 033 P 034 S 035 G 036 W 037 S 038 L 039 G 040 K 041 L 042 L 043 D 044 G 045 S 046 K 047 E 048 G 049 W 050 V 051 P 052 T 053 A 054 Y 055 M 056 K 057 P 058 H 059 S
ΔF
1.0
0.7
0.3
0.0
ΔF
1
0.5
0
$\hat{\mu}_{mean} = 0.02$
$\hat{\mu}_{mean} = 0.68$
$\hat{\mu}_{mean} = 0.99$
STOP (n=59)
non-synonymous (n=1112)
synonymous (n=54)
mutants

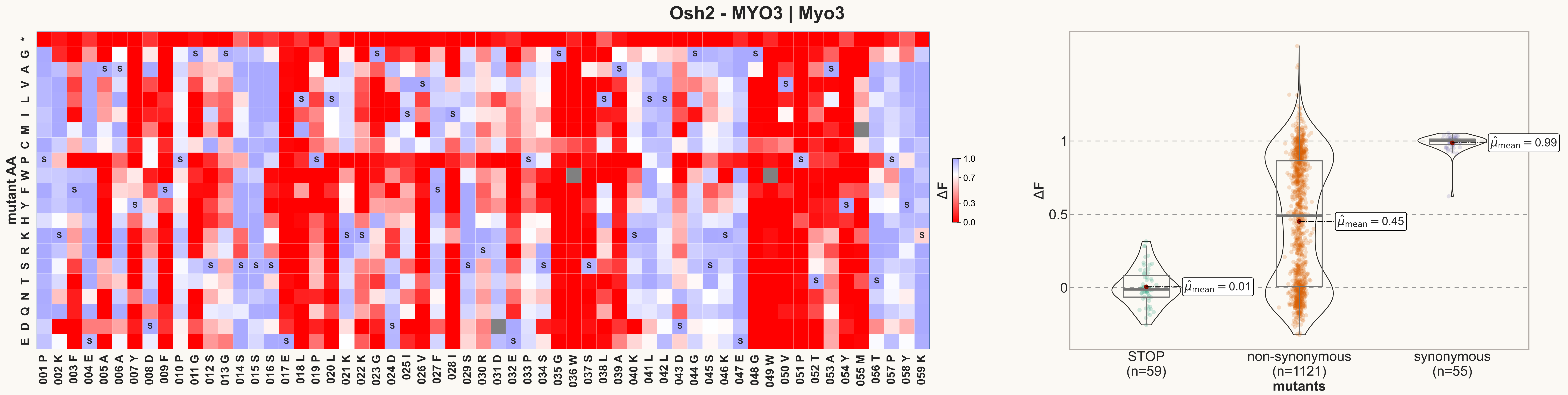
Osh2 - MYO3 | Myo3
mutant AA
ΔF
STOP
(n=59)
non-synonymous
(n=1121)
synonymous
(n=55)
mutants
$\hat{\mu}_{mean} = 0.01$
$\hat{\mu}_{mean} = 0.45$
$\hat{\mu}_{mean} = 0.99$

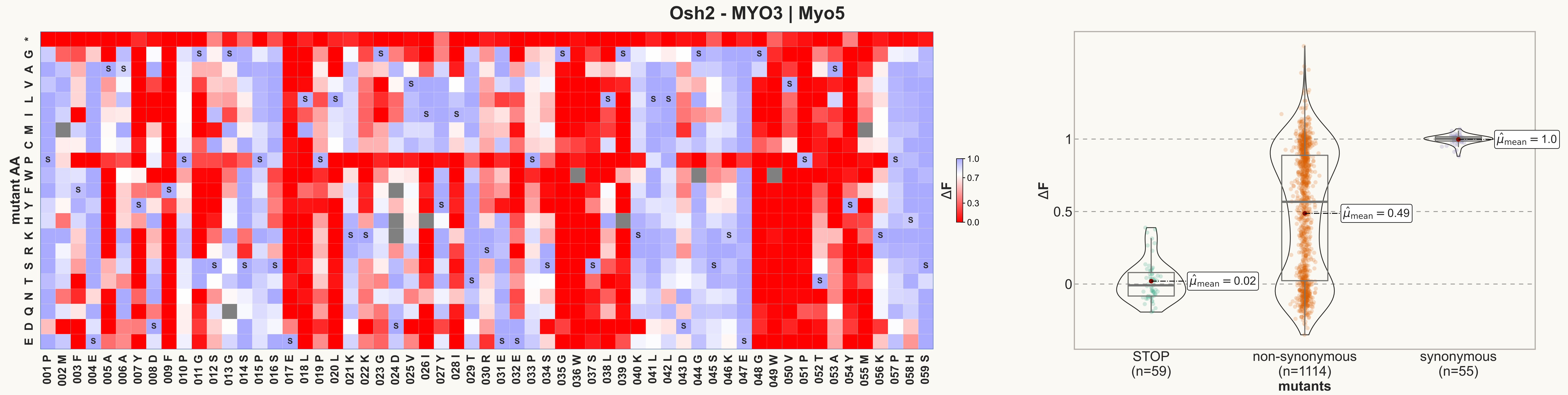

Osh2 - MYO3 | Myo5
mutant AA
ΔF
1.0
0.7
0.3
0.0
$\hat{\mu}_{mean}$ = 0.02
$\hat{\mu}_{mean}$ = 0.49
$\hat{\mu}_{mean}$ = 1.0
STOP
(n=59)
non-synonymous
(n=1114)
synonymous
(n=55)
mutants

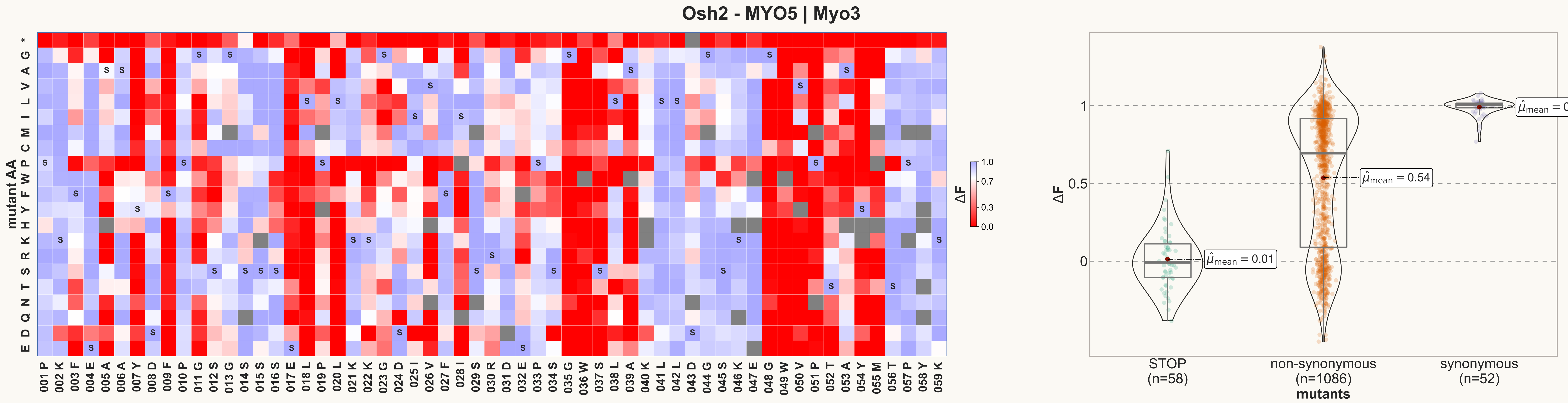

Osh2 - MYO5 | Myo3
mutant AA
ΔF
STOP (n=58)
non-synonymous (n=1086)
synonymous (n=52)
mutants
$\hat{\mu}_{mean} = 0.01$
$\hat{\mu}_{mean} = 0.54$
$\hat{\mu}_{mean} = 0.99$

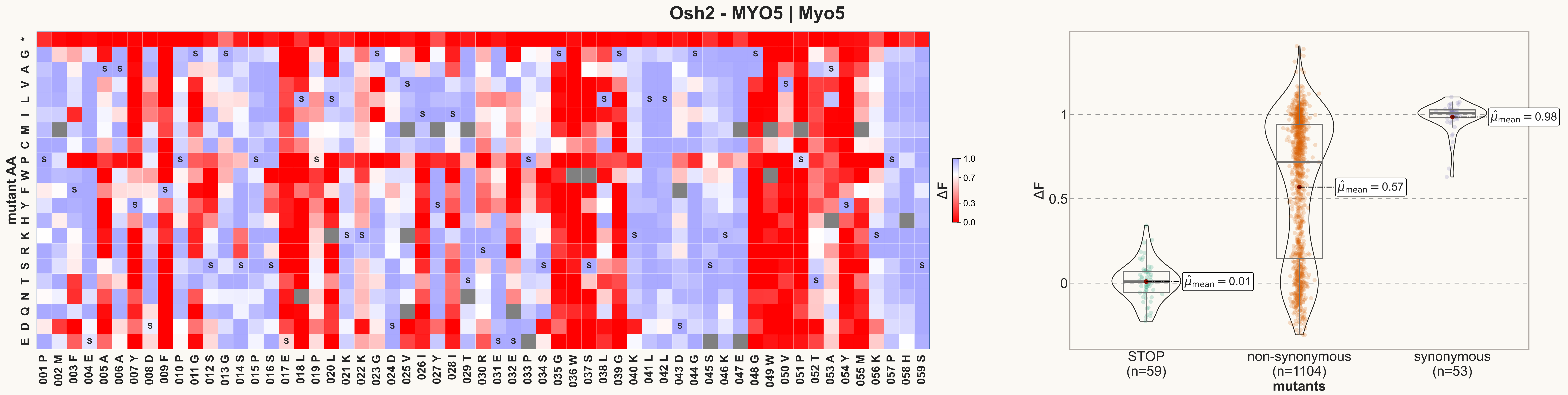

Osh2 - MYO5 | Myo5
mutant AA
* G A V L I M C P W F Y H K R S N T Q D E
001 P 002 M 003 F 004 E 005 A 006 A 007 Y 008 D 009 F 010 P 011 G 012 S 013 G 014 S 015 P 016 S 017 E 018 L 019 P 020 L 021 K 022 K 023 G 024 D 025 V 026 I 027 Y 028 I 029 T 030 R 031 E 032 E 033 P 034 S 035 G 036 W 037 S 038 L 039 G 040 K 041 L 042 L 043 D 044 G 045 S 046 K 047 E 048 G 049 W 050 V 051 P 052 T 053 A 054 Y 055 M 056 K 057 P 058 H 059 S
ΔF
1.0
0.7
0.3
0.0
ΔF
1
0.5
0
$\hat{\mu}_{mean} = 0.01$
$\hat{\mu}_{mean} = 0.57$
$\hat{\mu}_{mean} = 0.98$
STOP (n=59)
non-synonymous (n=1104)
synonymous (n=53)
mutants

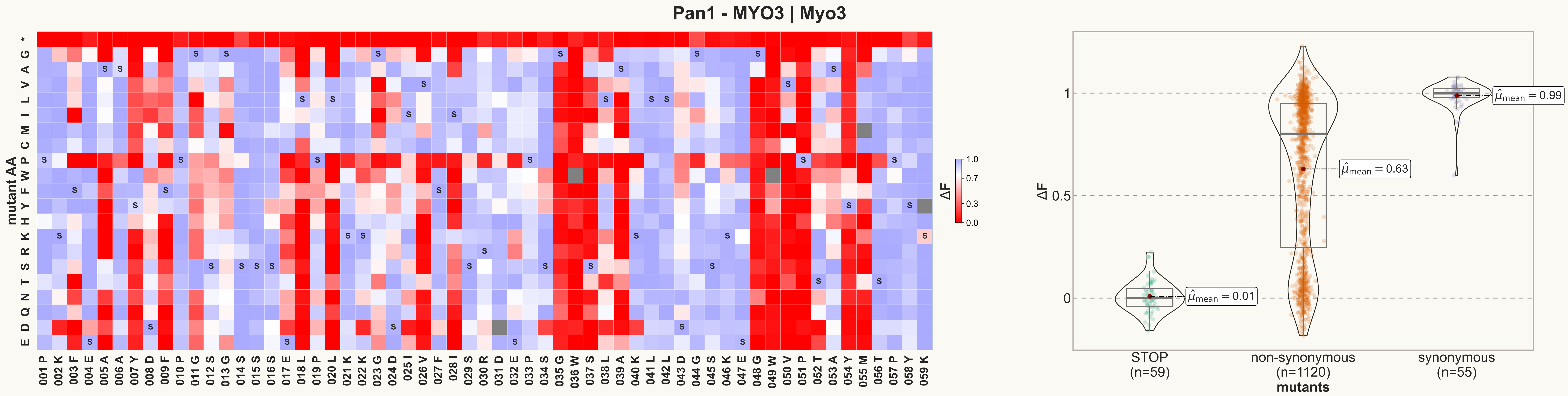
Pan1 - MYO3 | Myo3
mutant AA
* G A V L I M C P W F Y H K R S T N Q D E
001 P 002 K 003 F 004 E 005 A 006 A 007 Y 008 D 009 F 010 P 011 G 012 S 013 G 014 S 015 S 016 S 017 E 018 L 019 P 020 L 021 K 022 K 023 G 024 D 025 I 026 V 027 F 028 I 029 S 030 R 031 D 032 E 033 P 034 S 035 G 036 W 037 S 038 L 039 A 040 K 041 L 042 L 043 D 044 G 045 S 046 K 047 E 048 G 049 W 050 V 051 P 052 T 053 A 054 Y 055 M 056 T 057 P 058 Y 059 K
ΔF
1.0
0.7
0.3
0.0
ΔF
1
0.5
0
$\hat{\mu}_{mean} = 0.01$
$\hat{\mu}_{mean} = 0.63$
$\hat{\mu}_{mean} = 0.99$
STOP (n=59)
non-synonymous (n=1120)
synonymous (n=55)
mutants

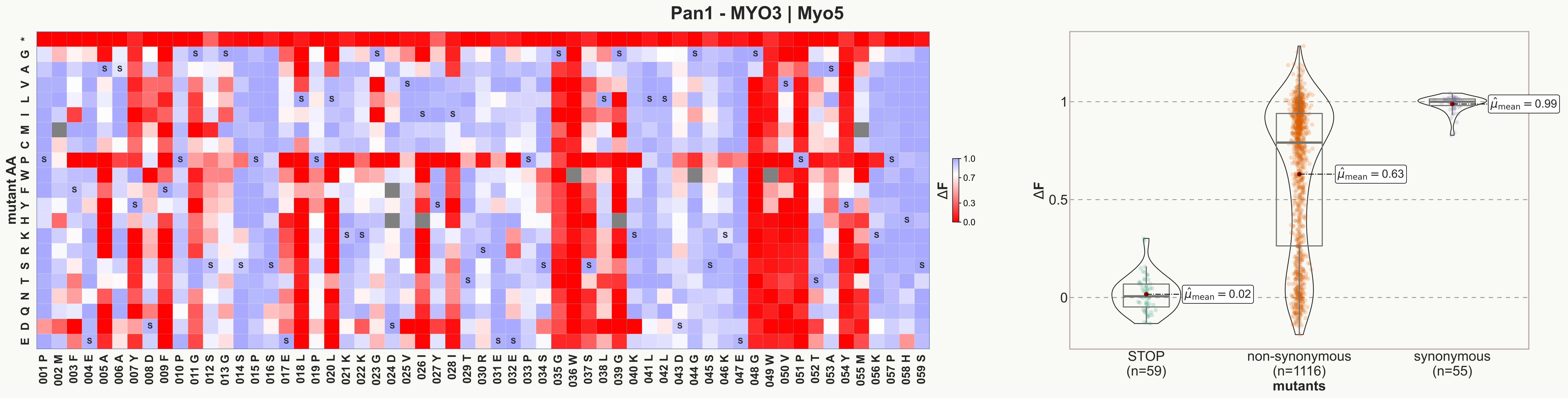
Pan1 - MYO3 | Myo5
mutant AA
* G A V L I M C P W F Y H K R S T N Q D E
001 P
002 M
003 F
004 E
005 A
006 A
007 Y
008 D
009 F
010 P
011 G
012 S
013 G
014 S
015 P
016 S
017 E
018 L
019 P
020 L
021 K
022 K
023 G
024 D
025 V
026 I
027 Y
028 I
029 T
030 R
031 E
032 E
033 P
034 S
035 G
036 W
037 S
038 L
039 G
040 K
041 L
042 L
043 D
044 G
045 S
046 K
047 E
048 G
049 W
050 V
051 P
052 T
053 A
054 Y
055 M
056 K
057 P
058 H
059 S
ΔF
1.0
0.7
0.3
0.0
ΔF
1
0.5
0
$\hat{\mu}_{mean}$ = 0.02
$\hat{\mu}_{mean}$ = 0.63
$\hat{\mu}_{mean}$ = 0.99
STOP
(n=59)
non-synonymous
(n=1116)
synonymous
(n=55)
mutants

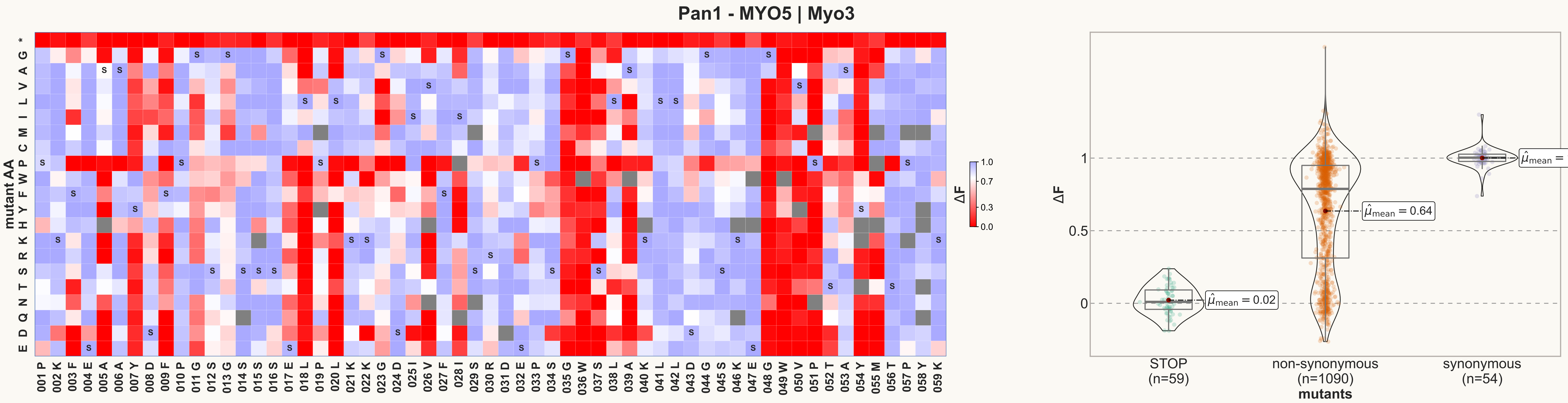
Pan1 - MYO5 | Myo3
mutant AA
* G A V L I M C P W F Y H K R S T N Q D E
001 P 002 K 003 F 004 E 005 A 006 A 007 Y 008 D 009 F 010 P 011 G 012 S 013 G 014 S 015 S 016 S 017 E 018 L 019 P 020 L 021 K 022 K 023 G 024 D 025 I 026 V 027 F 028 I 029 S 030 R 031 D 032 E 033 P 034 S 035 G 036 W 037 S 038 L 039 A 040 K 041 L 042 L 043 D 044 G 045 S 046 K 047 E 048 G 049 W 050 V 051 P 052 T 053 A 054 Y 055 M 056 T 057 P 058 Y 059 K
ΔF
1.0
0.7
0.3
0.0
ΔF
1
0.5
0
$\hat{\mu}_{mean}$ = 0.02
$\hat{\mu}_{mean}$ = 0.64
$\hat{\mu}_{mean}$ = 1.0
STOP (n=59)
non-synonymous (n=1090)
synonymous (n=54)
mutants

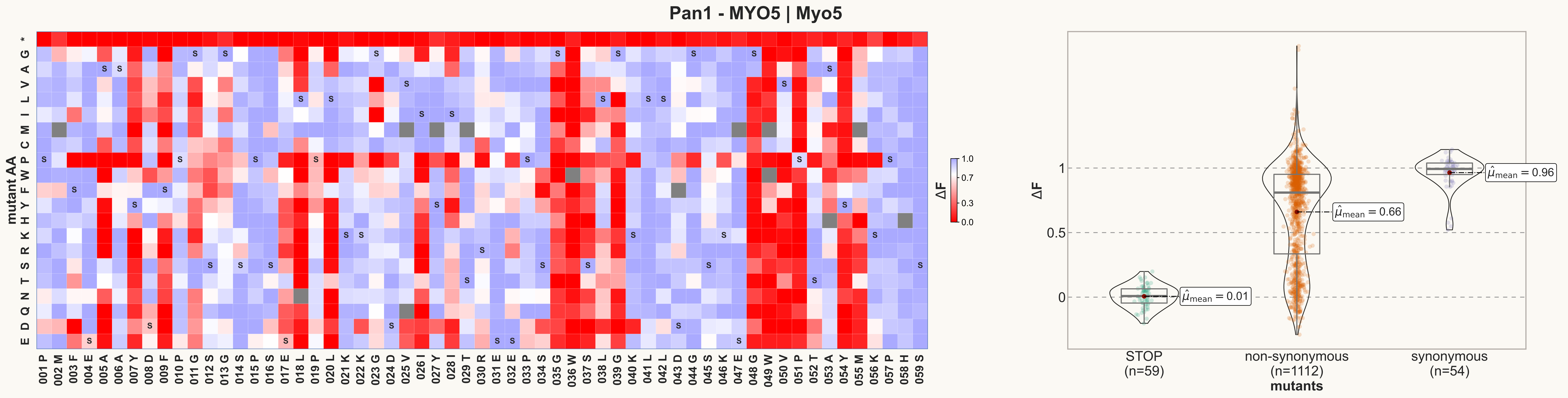
Pan1 - MYO5 | Myo5
mutant AA
* G A V L I M C P W F Y H K R S T N Q D E
001 P 002 M 003 F 004 E 005 A 006 A 007 Y 008 D 009 F 010 P 011 G 012 S 013 G 014 S 015 P 016 S 017 E 018 L 019 P 020 L 021 K 022 K 023 G 024 D 025 V 026 I 027 Y 028 I 029 T 030 R 031 E 032 E 033 P 034 S 035 G 036 W 037 S 038 L 039 G 040 K 041 L 042 L 043 D 044 G 045 S 046 K 047 E 048 G 049 W 050 V 051 P 052 T 053 A 054 Y 055 M 056 K 057 P 058 H 059 S
ΔF
1.0
0.7
0.3
0.0
ΔF
1
0.5
0
$\hat{\mu}_{mean} = 0.01$
$\hat{\mu}_{mean} = 0.66$
$\hat{\mu}_{mean} = 0.96$
STOP (n=59)
non-synonymous (n=1112)
synonymous (n=54)
mutants

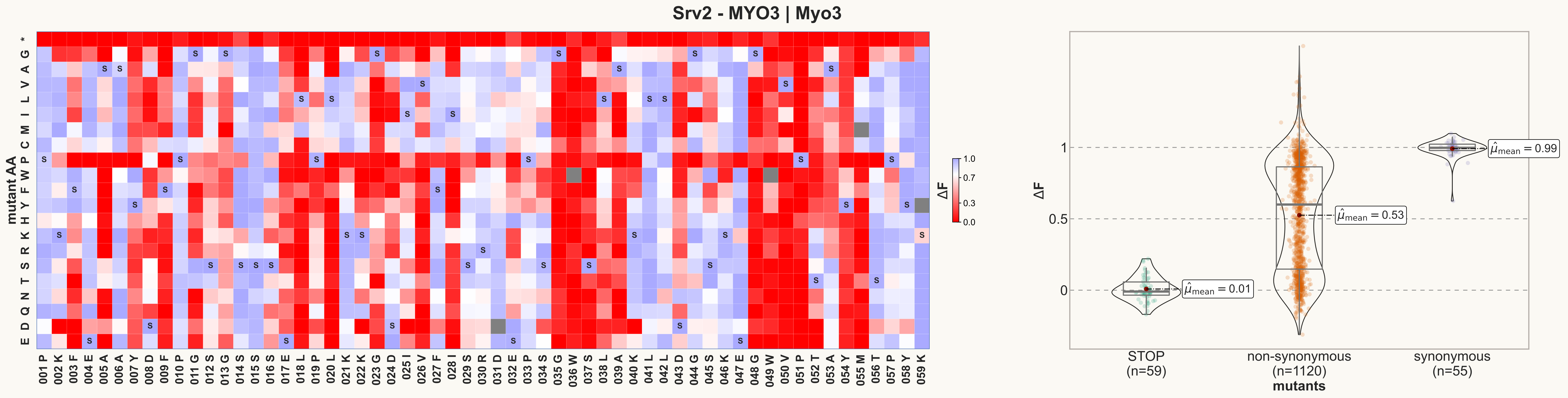
Srv2 - MYO3 | Myo3
mutant AA
* G A V L I M C P W F Y H K R S T N Q D E
001 P 002 K 003 F 004 E 005 A 006 A 007 Y 008 D 009 F 010 P 011 G 012 S 013 G 014 S 015 S 016 S 017 E 018 L 019 P 020 L 021 K 022 K 023 G 024 D 025 I 026 V 027 F 028 I 029 S 030 R 031 D 032 E 033 P 034 S 035 G 036 W 037 S 038 L 039 A 040 K 041 L 042 L 043 D 044 G 045 S 046 K 047 E 048 G 049 W 050 V 051 P 052 T 053 A 054 Y 055 M 056 T 057 P 058 Y 059 K
ΔF
1.0
0.7
0.3
0.0
ΔF
1
0.5
0
$\hat{\mu}_{mean} = 0.01$
$\hat{\mu}_{mean} = 0.53$
$\hat{\mu}_{mean} = 0.99$
STOP (n=59)
non-synonymous (n=1120)
synonymous (n=55)
mutants

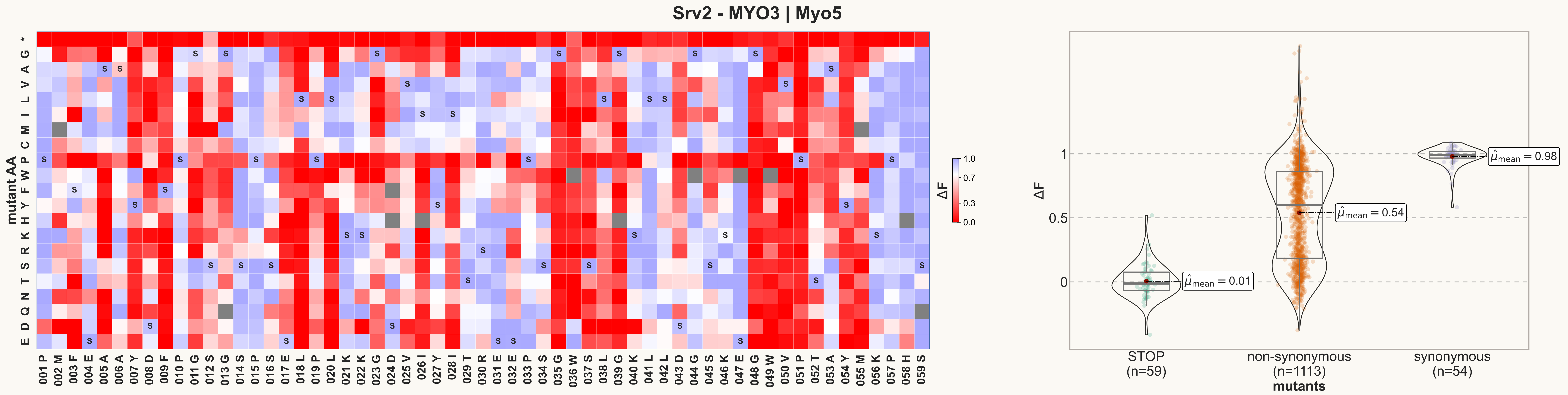
Srv2 - MYO3 | Myo5
mutant AA
ΔF
STOP (n=59)
non-synonymous (n=1113)
synonymous (n=54)
mutants
$\hat{\mu}_{mean} = 0.01$
$\hat{\mu}_{mean} = 0.54$
$\hat{\mu}_{mean} = 0.98$

**Srv2 - MYO5 | Myo3**

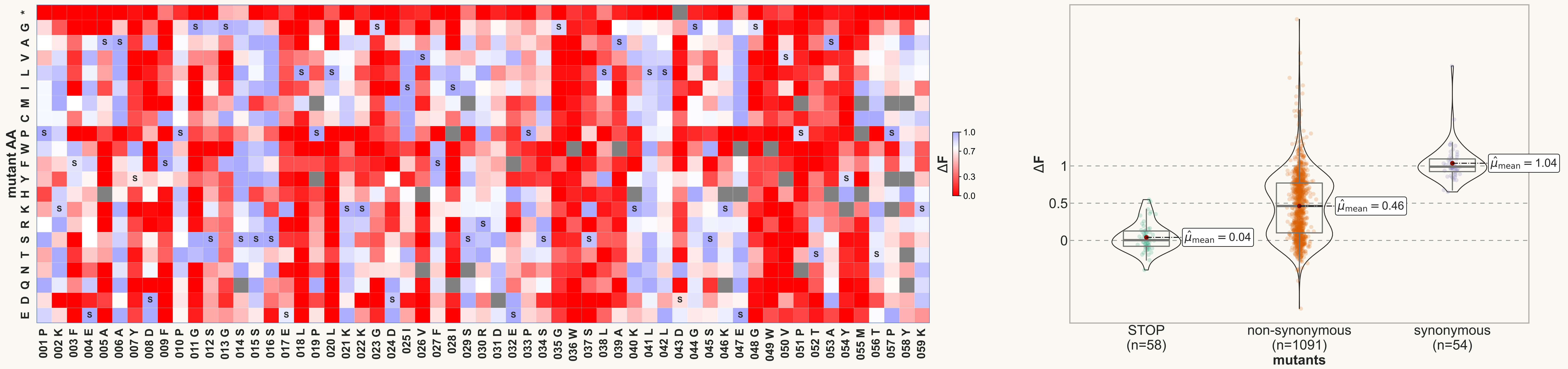

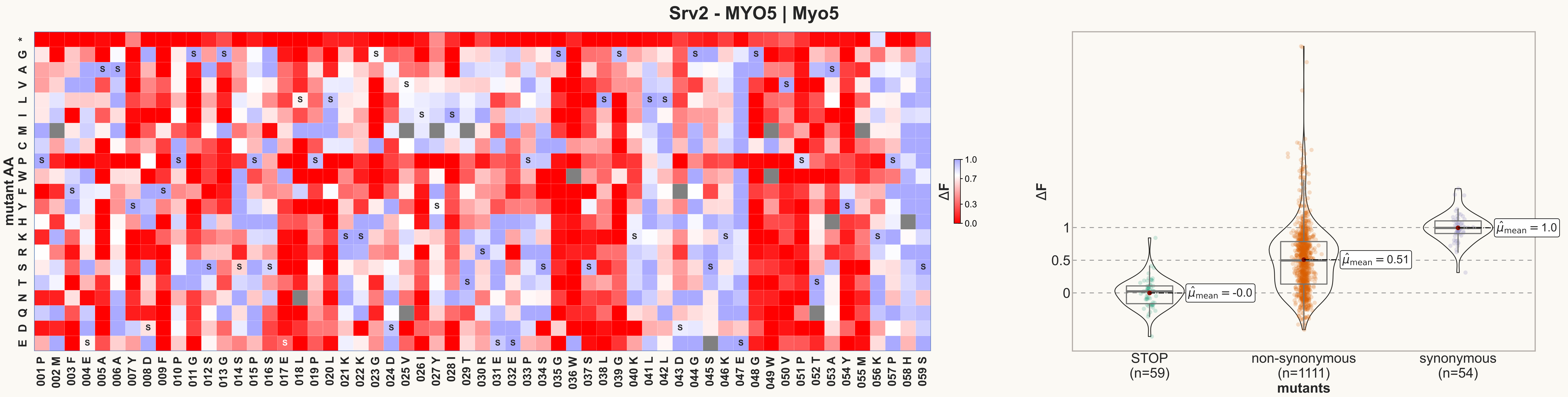

Srv2 - MYO5 | Myo5
mutant AA
ΔF
STOP
(n=59)
non-synonymous
(n=1111)
synonymous
(n=54)
mutants
$\hat{\mu}_{mean}$ = -0.0
$\hat{\mu}_{mean}$ = 0.51
$\hat{\mu}_{mean}$ = 1.0
