## Supplemental File 6 for "Cryptic genetic variation shapes the fate of gene duplicates in a protein interaction network"

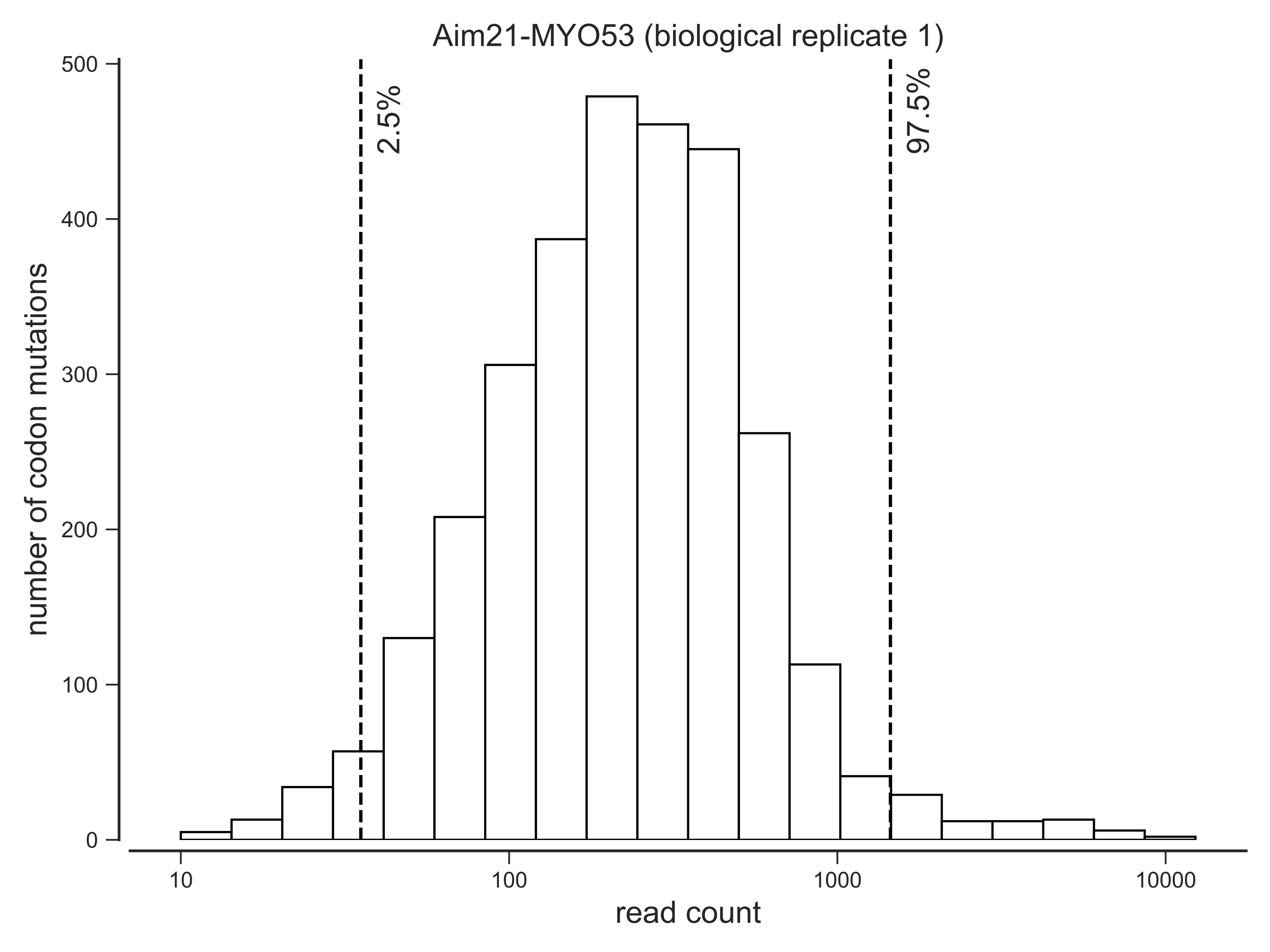
Aim21-MYO53 (biological replicate 1)
number of codon mutations
read count
2.5%
97.5%
0
100
200
300
400
500
10
100
1000
10000

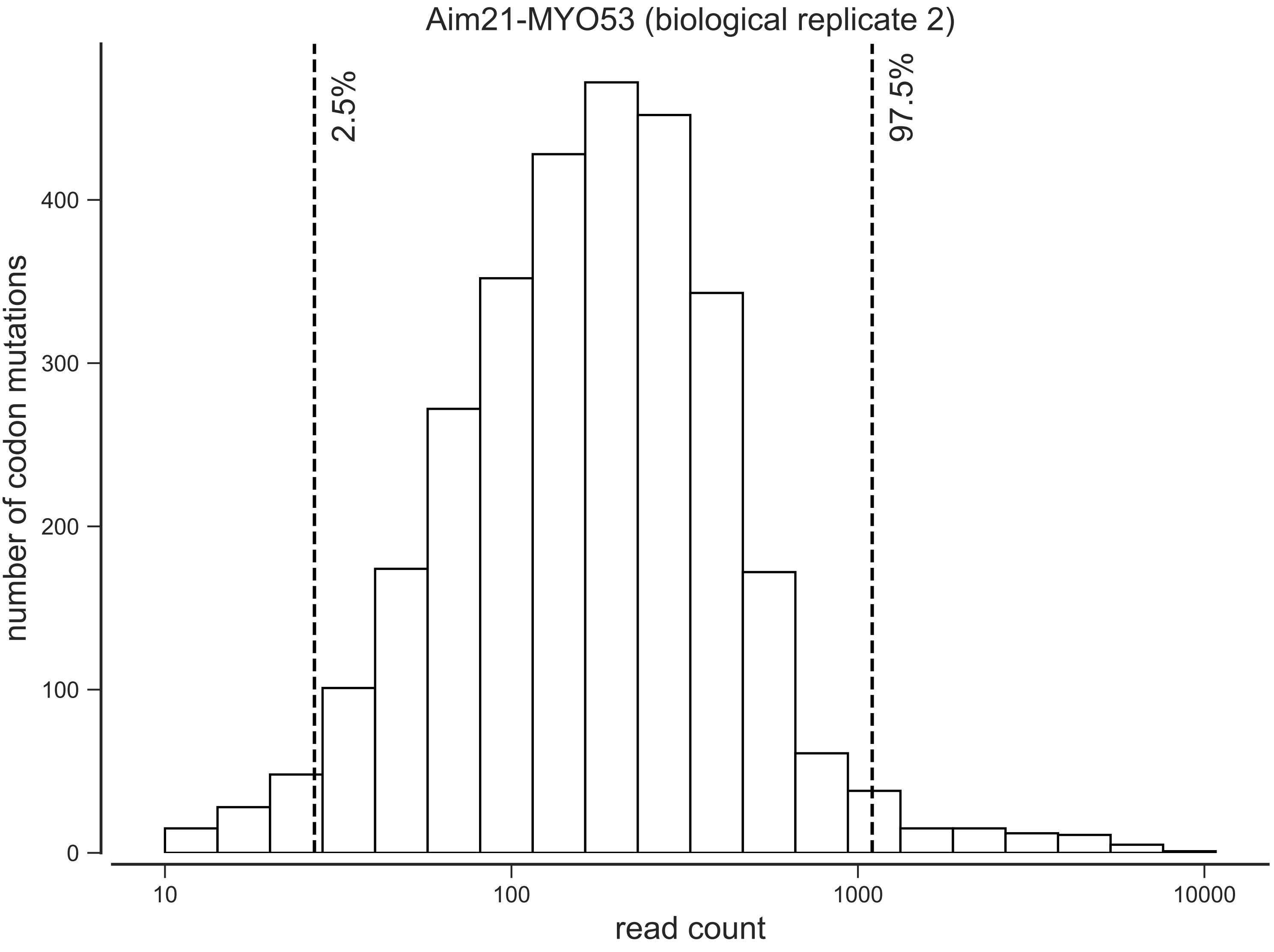

Aim21-MYO53 (biological replicate 2)
2.5%
97.5%
number of codon mutations
0
100
200
300
400
10
100
1000
10000
read count

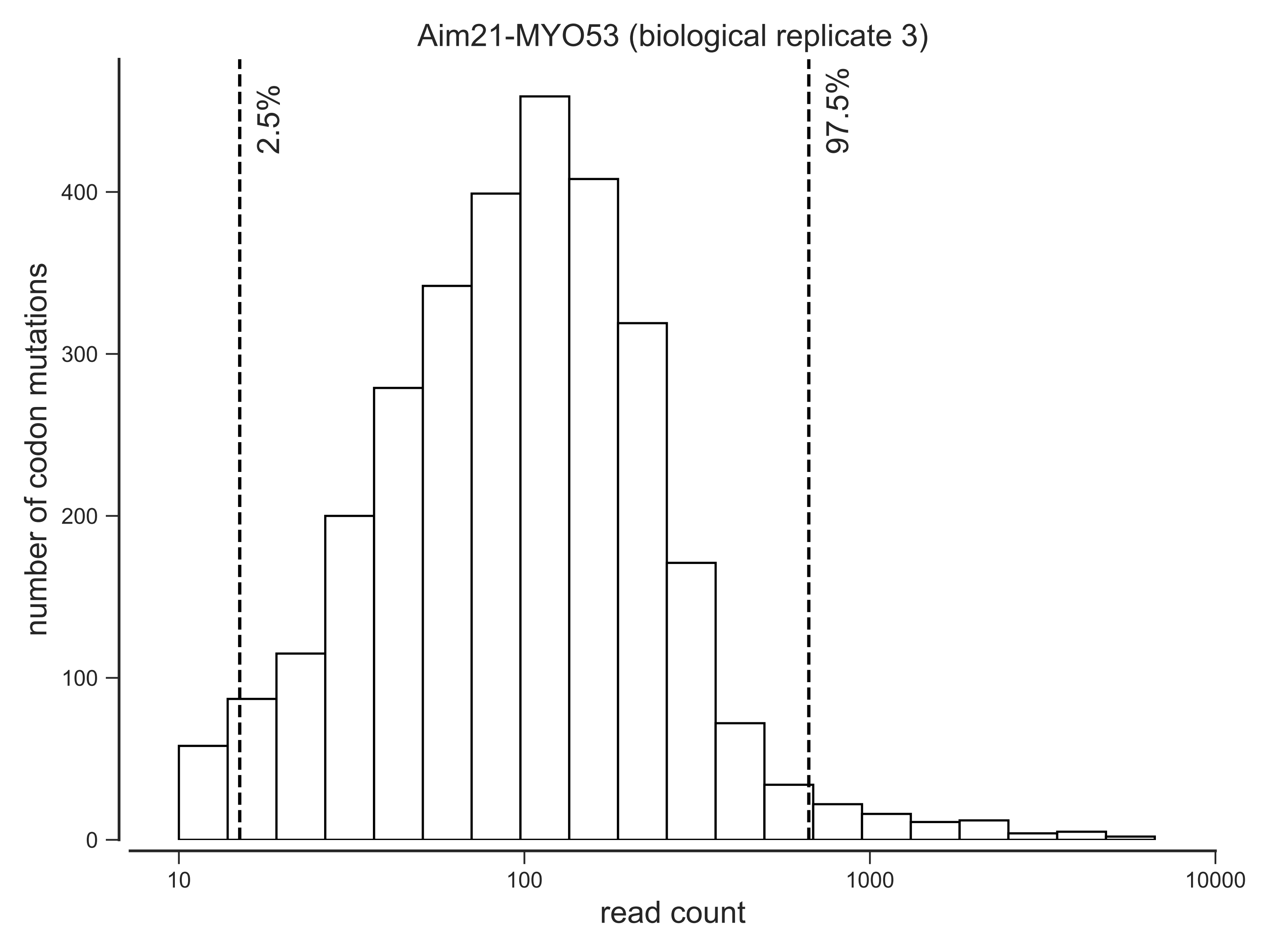

Aim21-MYO53 (biological replicate 3)
2.5%
97.5%
number of codon mutations
0
100
200
300
400
10
100
1000
10000
read count

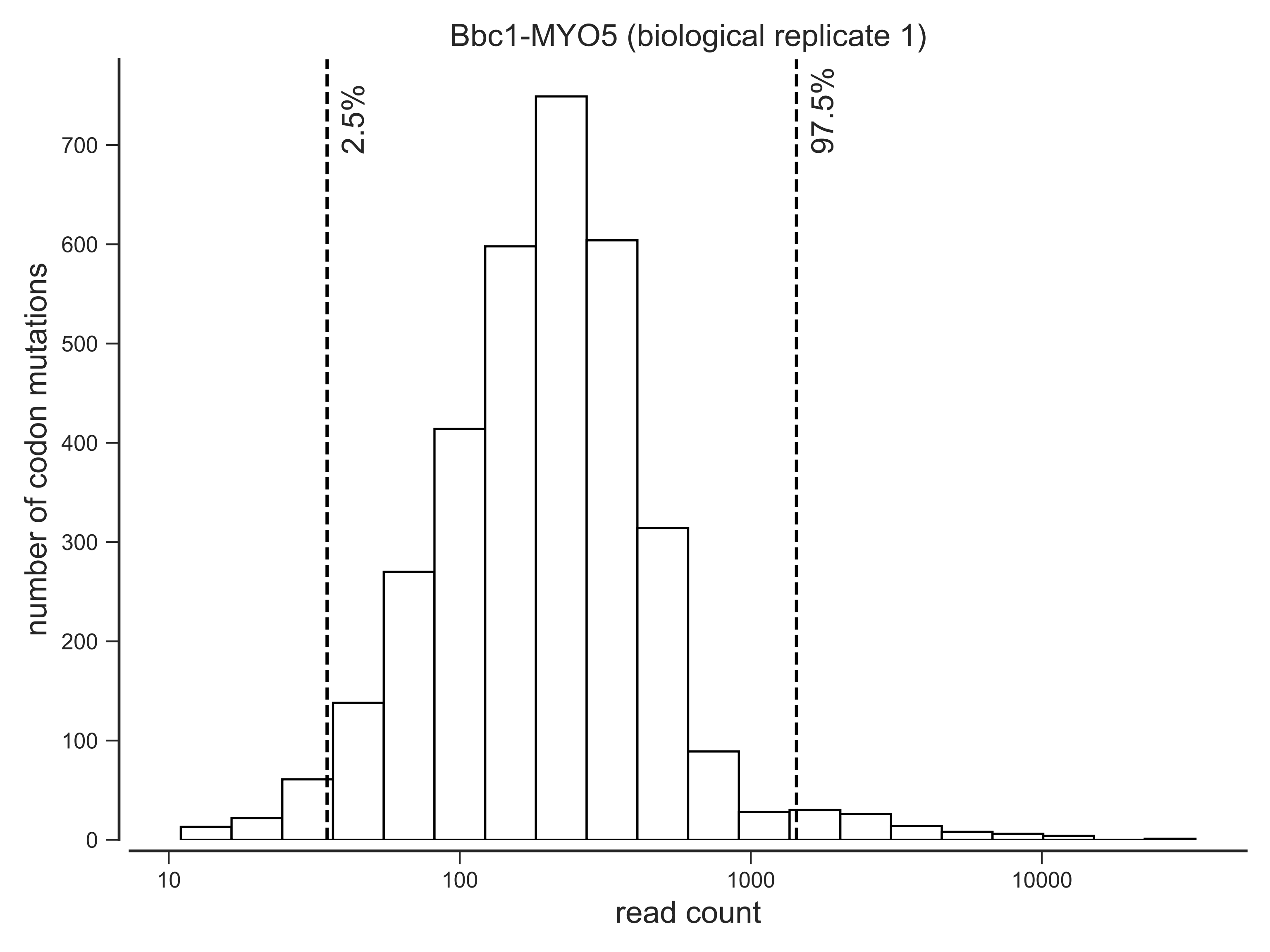

Bbc1-MYO5 (biological replicate 1)
2.5%
97.5%
0
100
200
300
400
500
600
700
number of codon mutations
10
100
1000
10000
read count

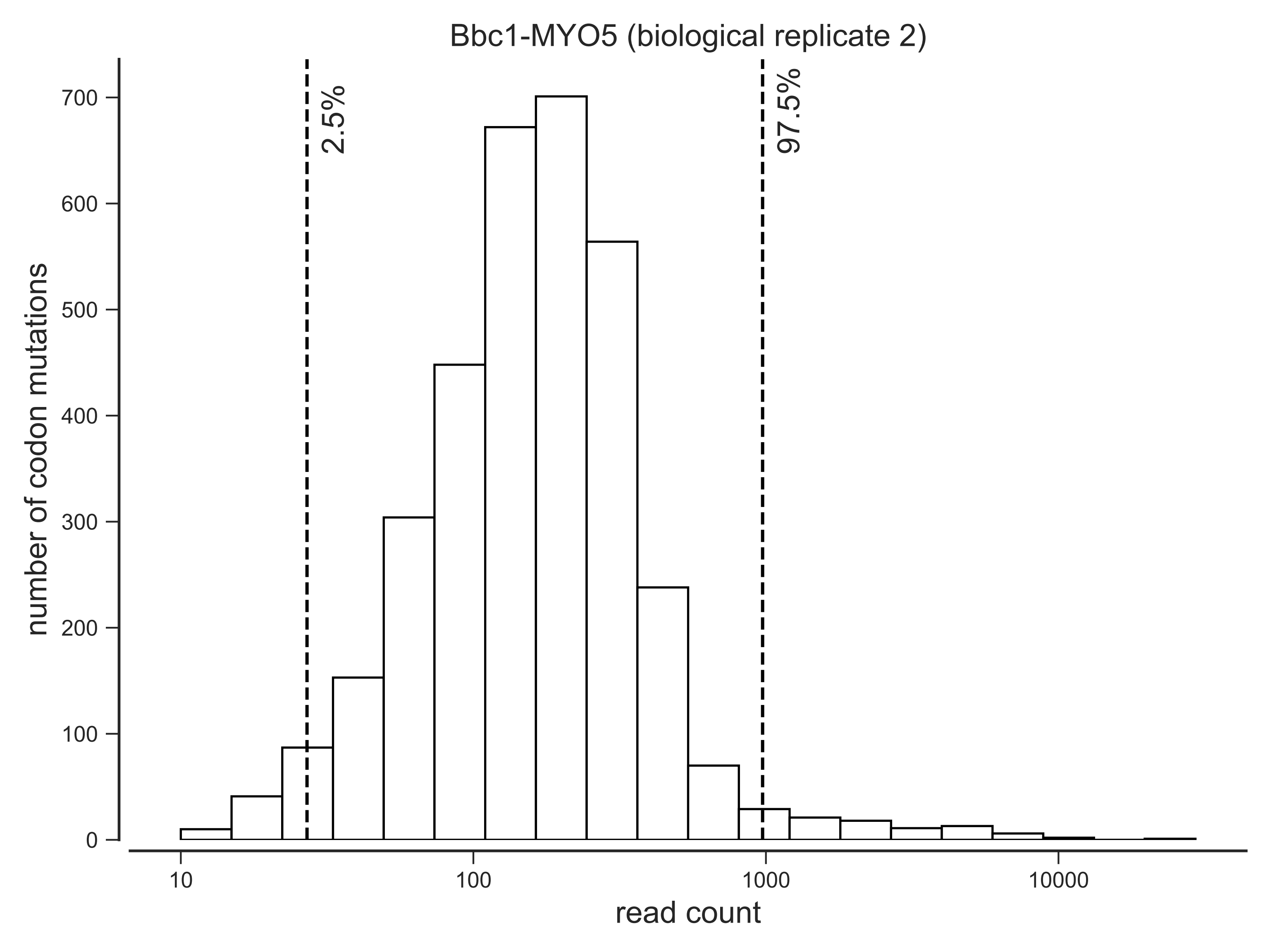
Bbc1-MYO5 (biological replicate 2)
2.5%
97.5%
number of codon mutations
0
100
200
300
400
500
600
700
10
100
1000
10000
read count

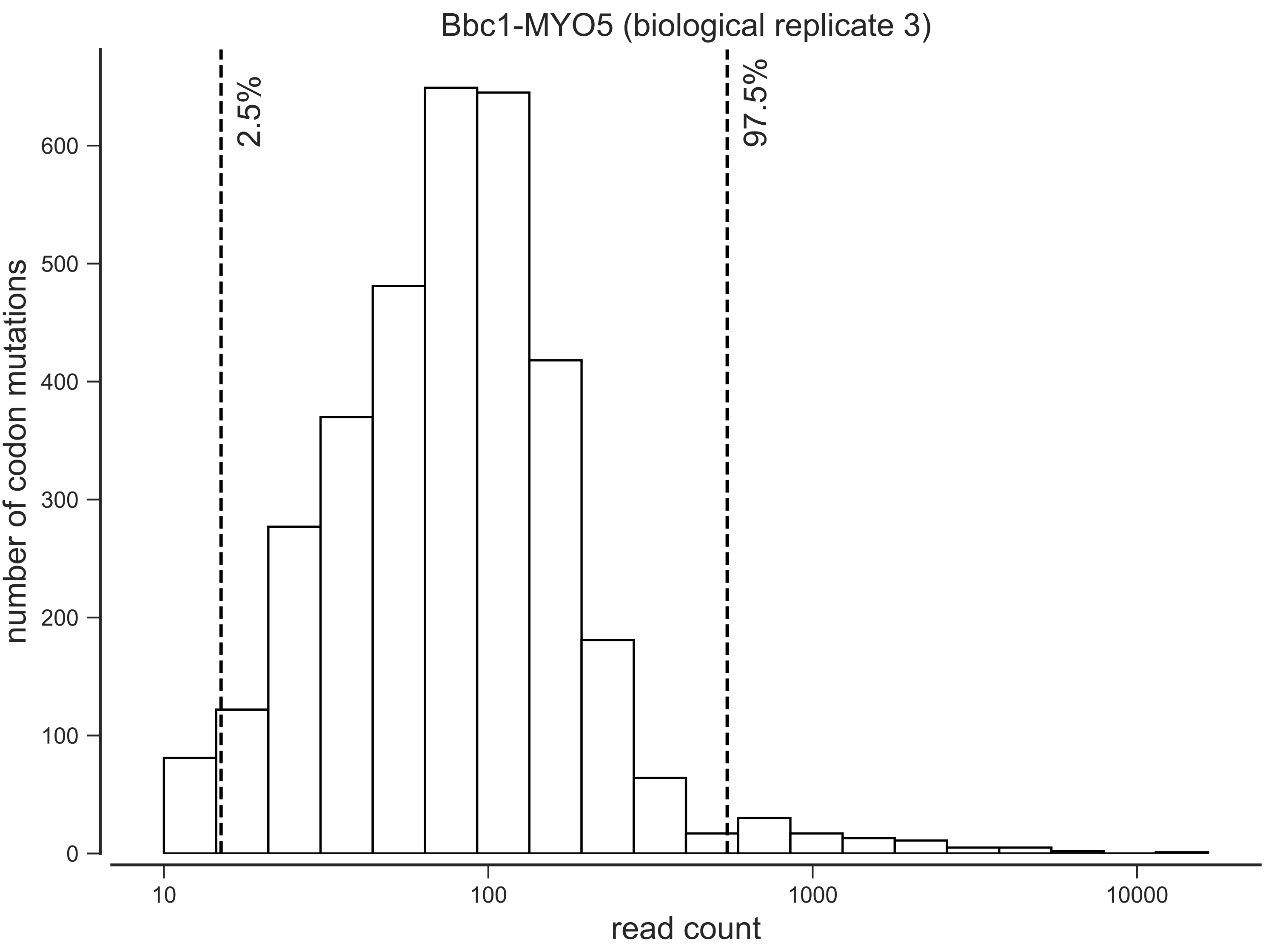

Bbc1-MYO5 (biological replicate 3)
2.5%
97.5%
number of codon mutations
0
100
200
300
400
500
600
10
100
1000
10000
read count

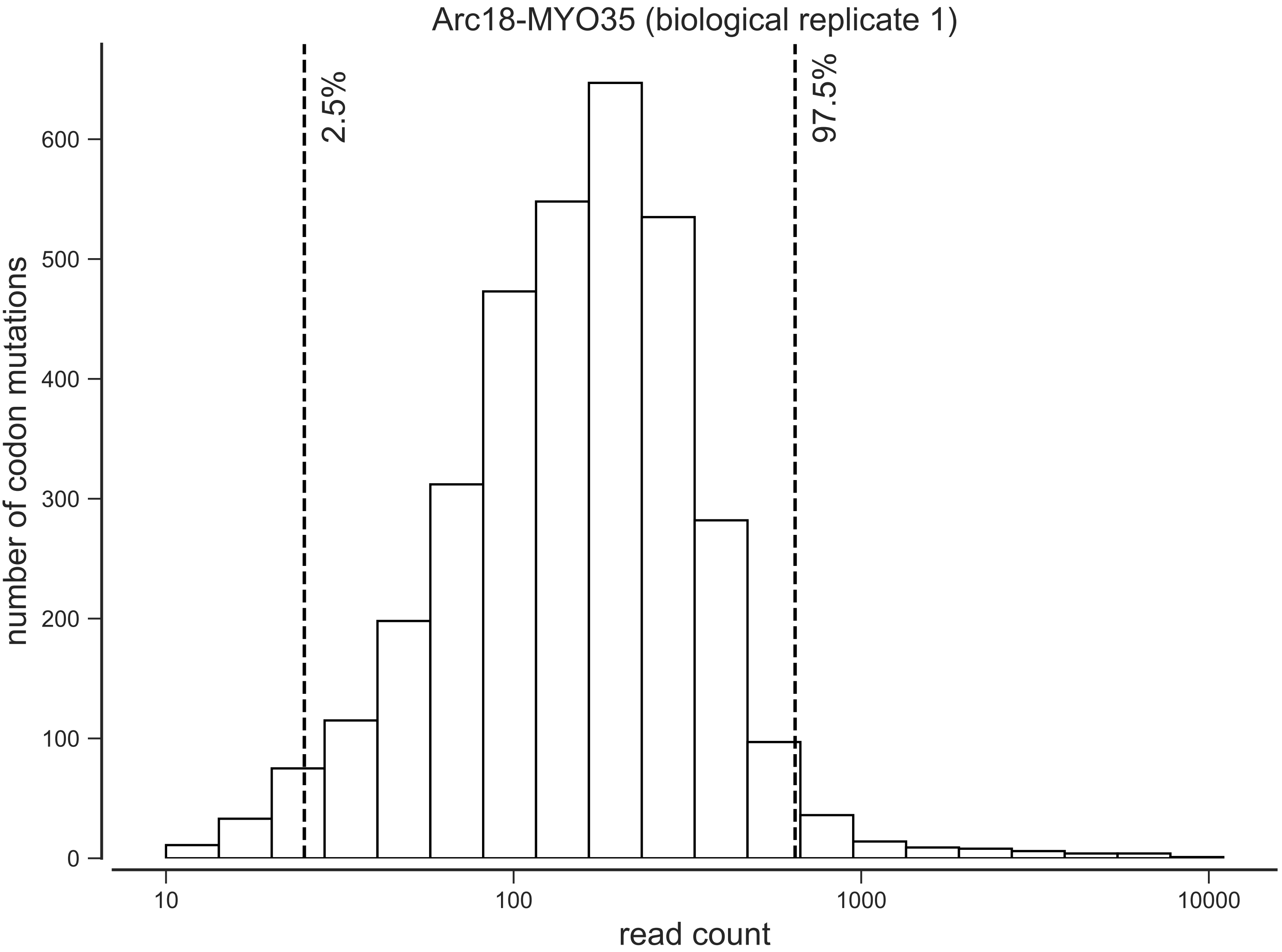

Arc18-MYO35 (biological replicate 1)
2.5%
97.5%
number of codon mutations
0
100
200
300
400
500
600
10
100
1000
10000
read count

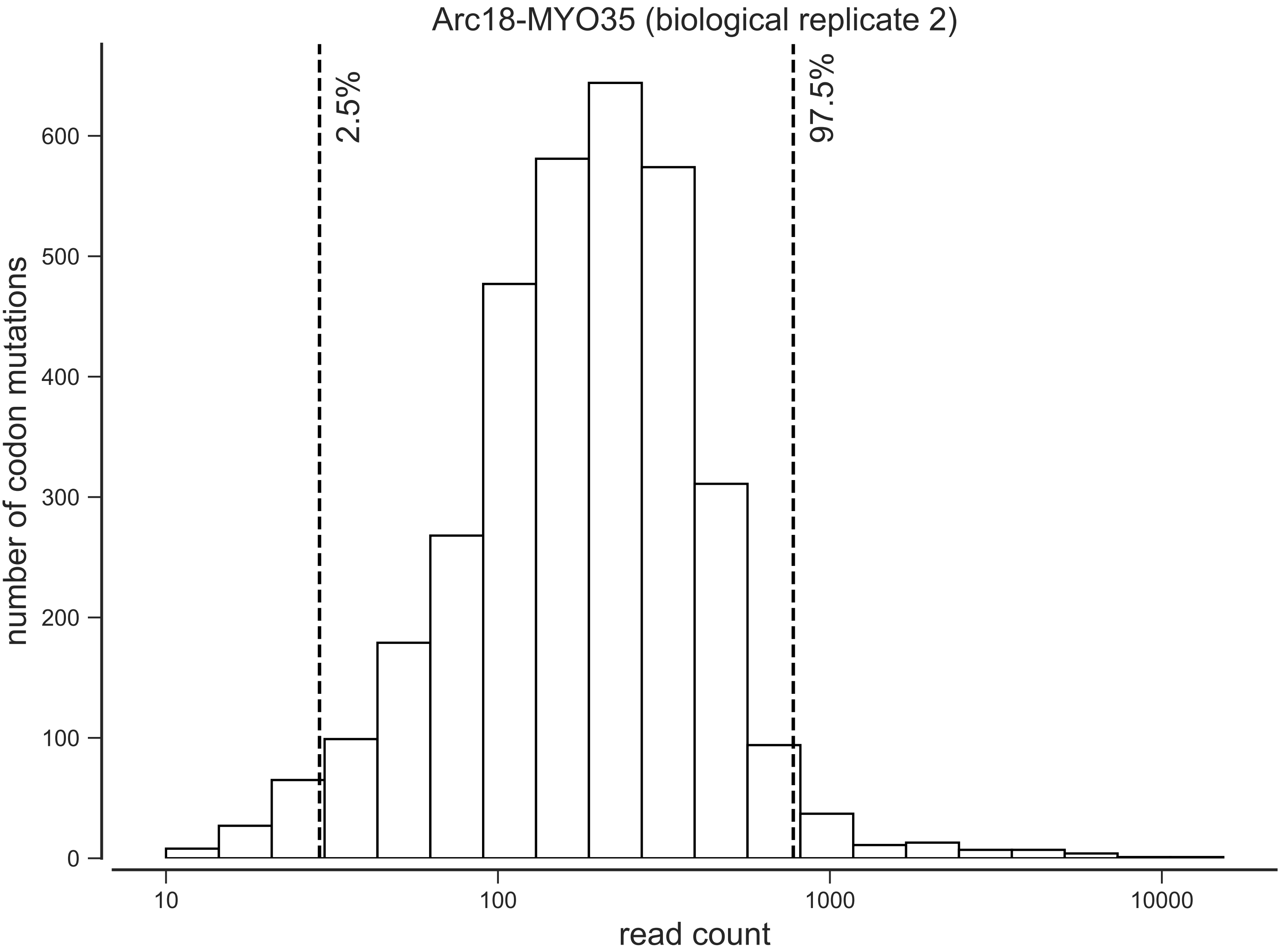
Arc18-MYO35 (biological replicate 2)
2.5%
97.5%
number of codon mutations
0
100
200
300
400
500
600
10
100
1000
10000
read count

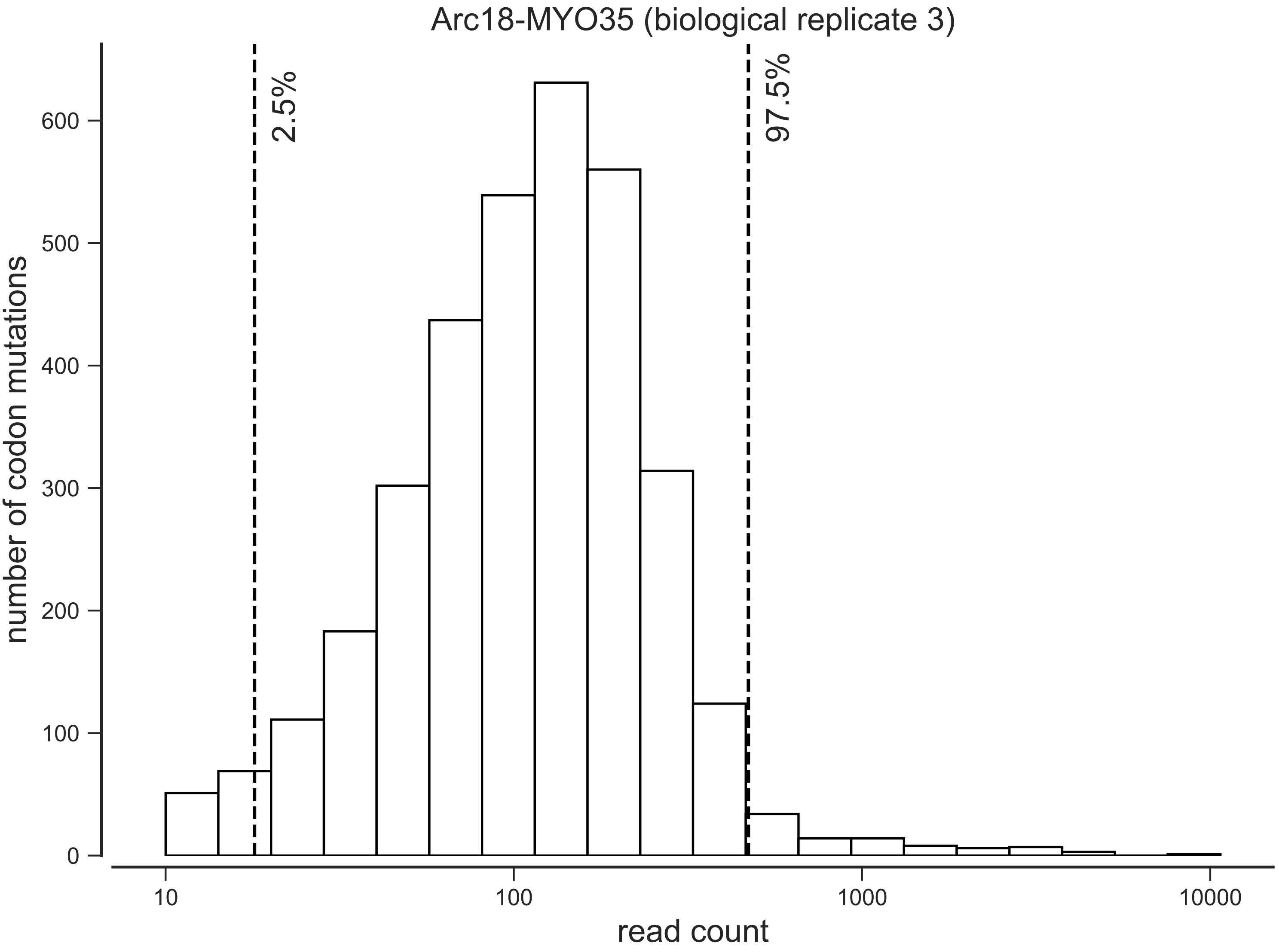

Arc18-MYO35 (biological replicate 3)
2.5%
97.5%
number of codon mutations
0
100
200
300
400
500
600
10
100
1000
10000
read count

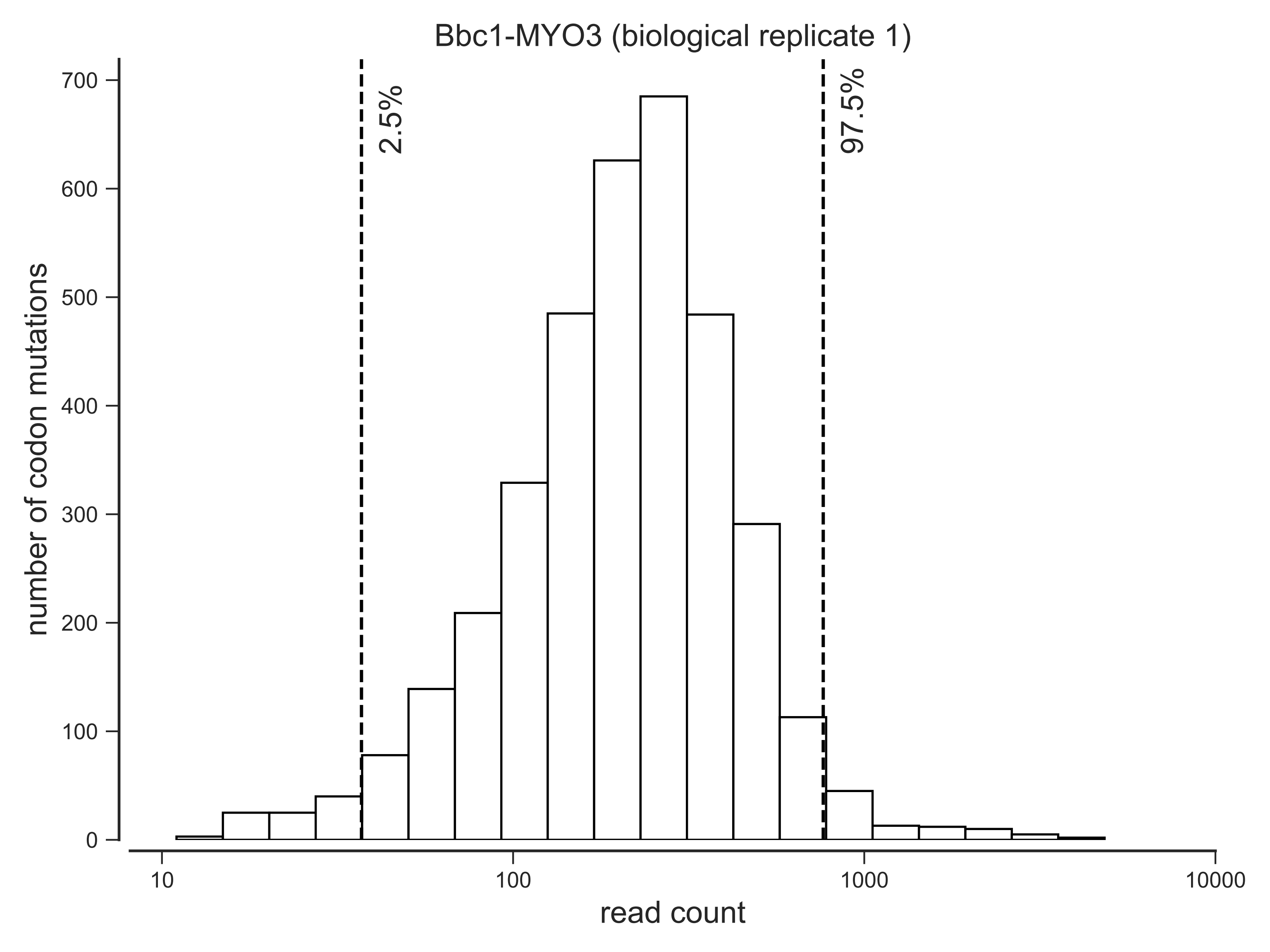
Bbc1-MYO3 (biological replicate 1)
2.5%
97.5%
number of codon mutations
0
100
200
300
400
500
600
700
10
100
1000
10000
read count

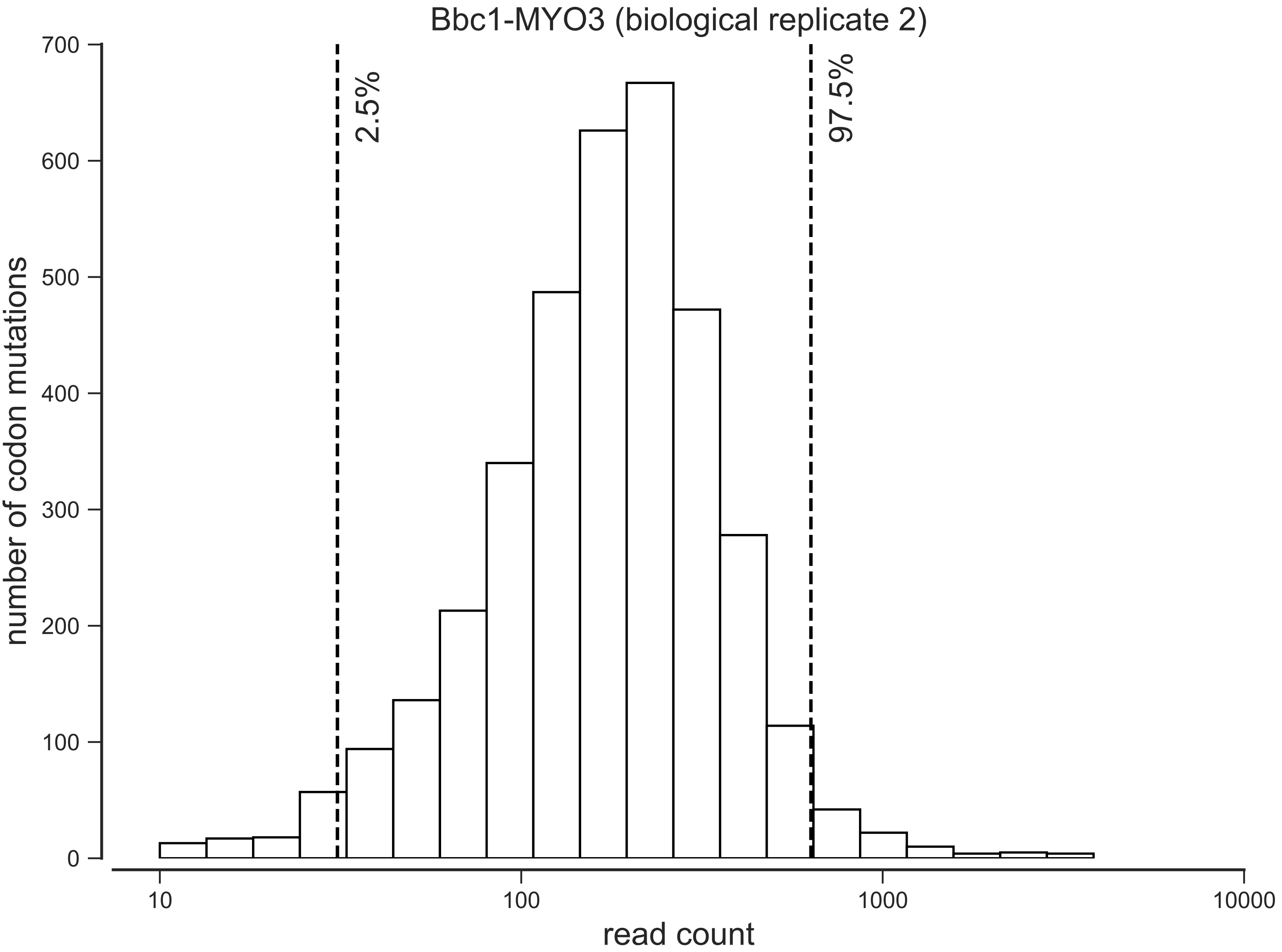

Bbc1-MYO3 (biological replicate 2)
2.5%
97.5%
number of codon mutations
read count
0
100
200
300
400
500
600
700
10
100
1000
10000

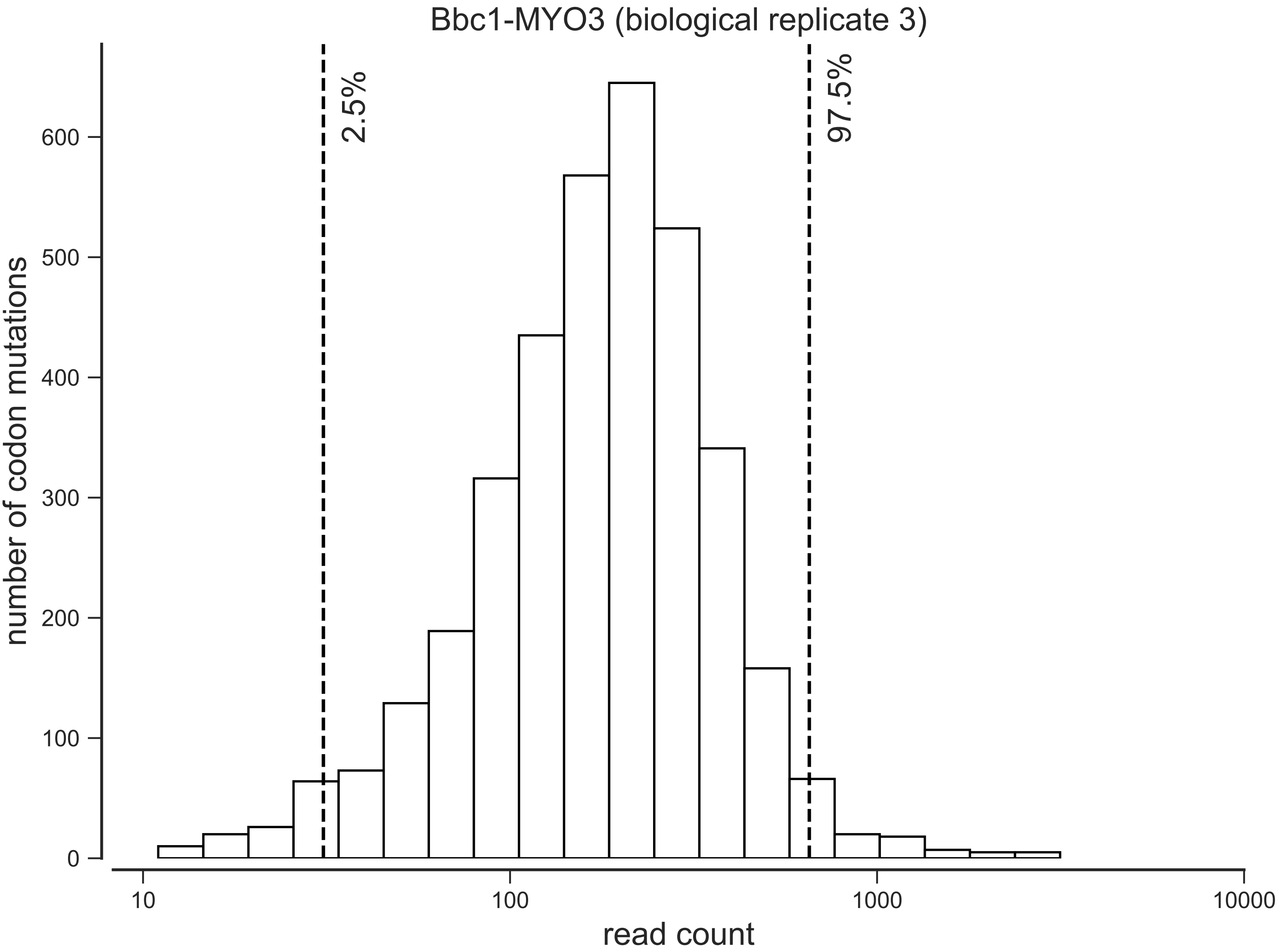

Bbc1-MYO3 (biological replicate 3)
2.5%
97.5%
number of codon mutations
0
100
200
300
400
500
600
10
100
1000
10000
read count

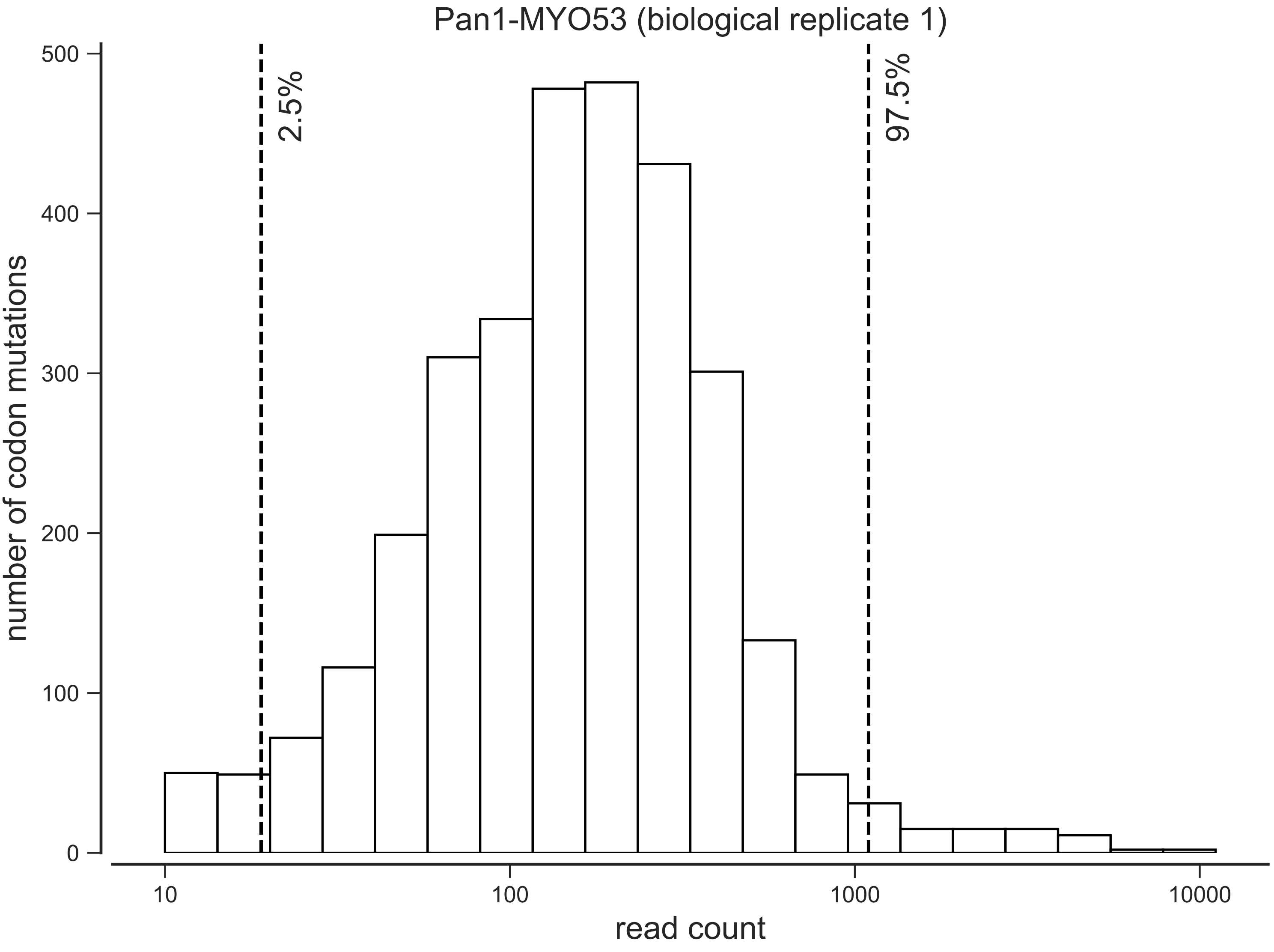

Pan1-MYO53 (biological replicate 1)
number of codon mutations
read count
2.5%
97.5%
0
100
200
300
400
500
10
100
1000
10000

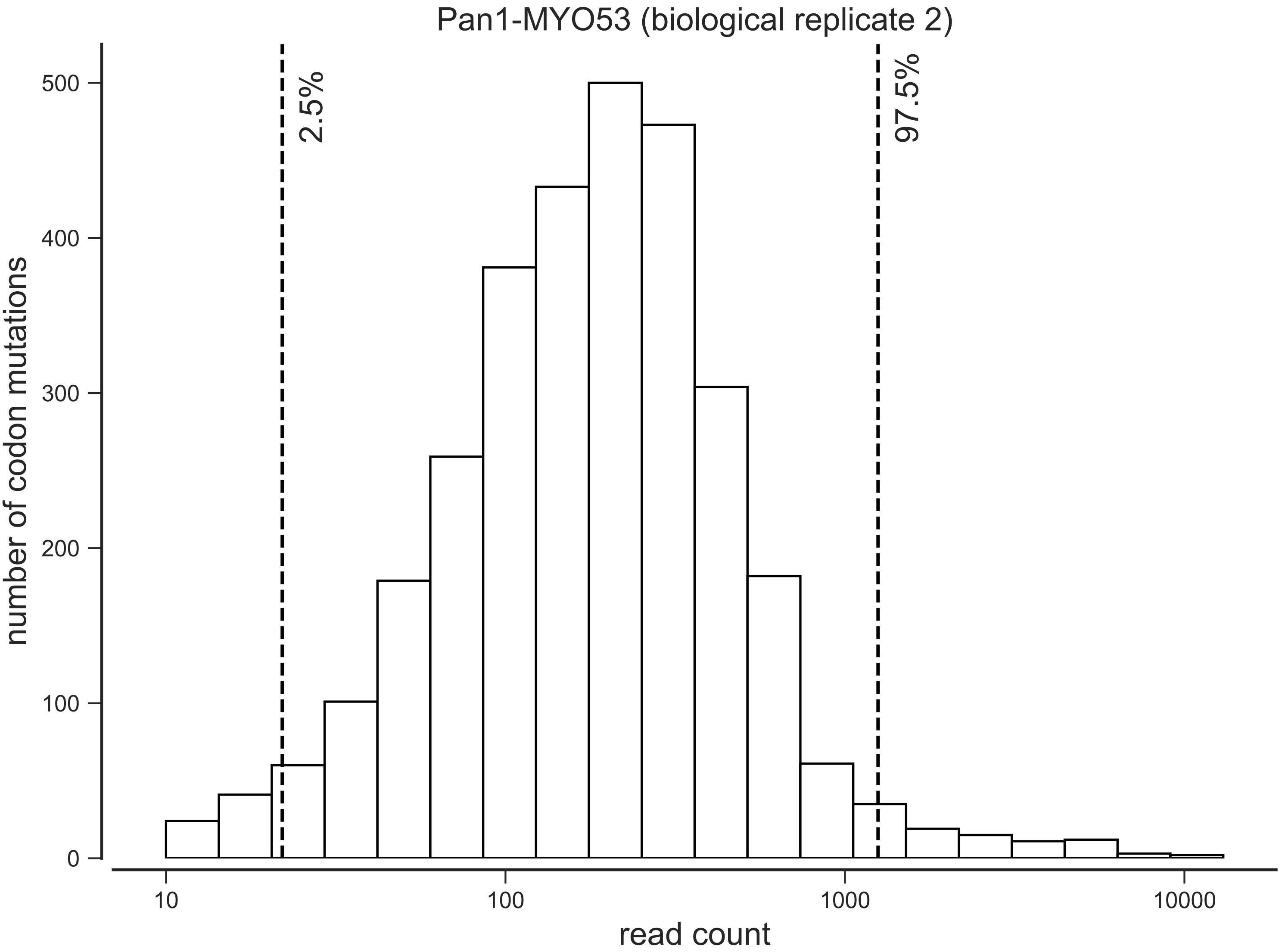
Pan1-MYO53 (biological replicate 2)
2.5%
97.5%
number of codon mutations
0
100
200
300
400
500
10
100
1000
10000
read count

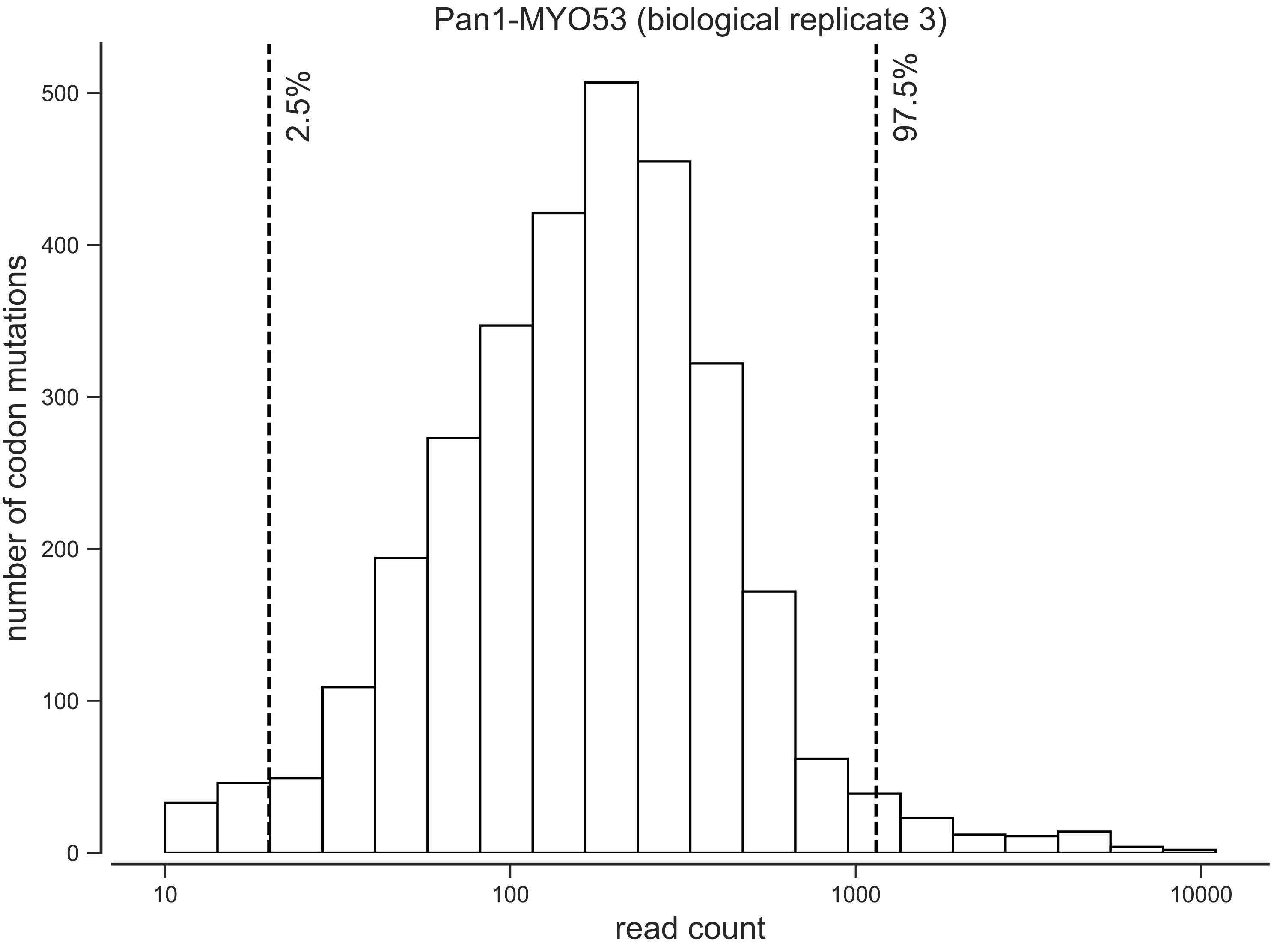

Pan1-MYO53 (biological replicate 3)
2.5%
97.5%
number of codon mutations
0
100
200
300
400
500
10
100
1000
10000
read count

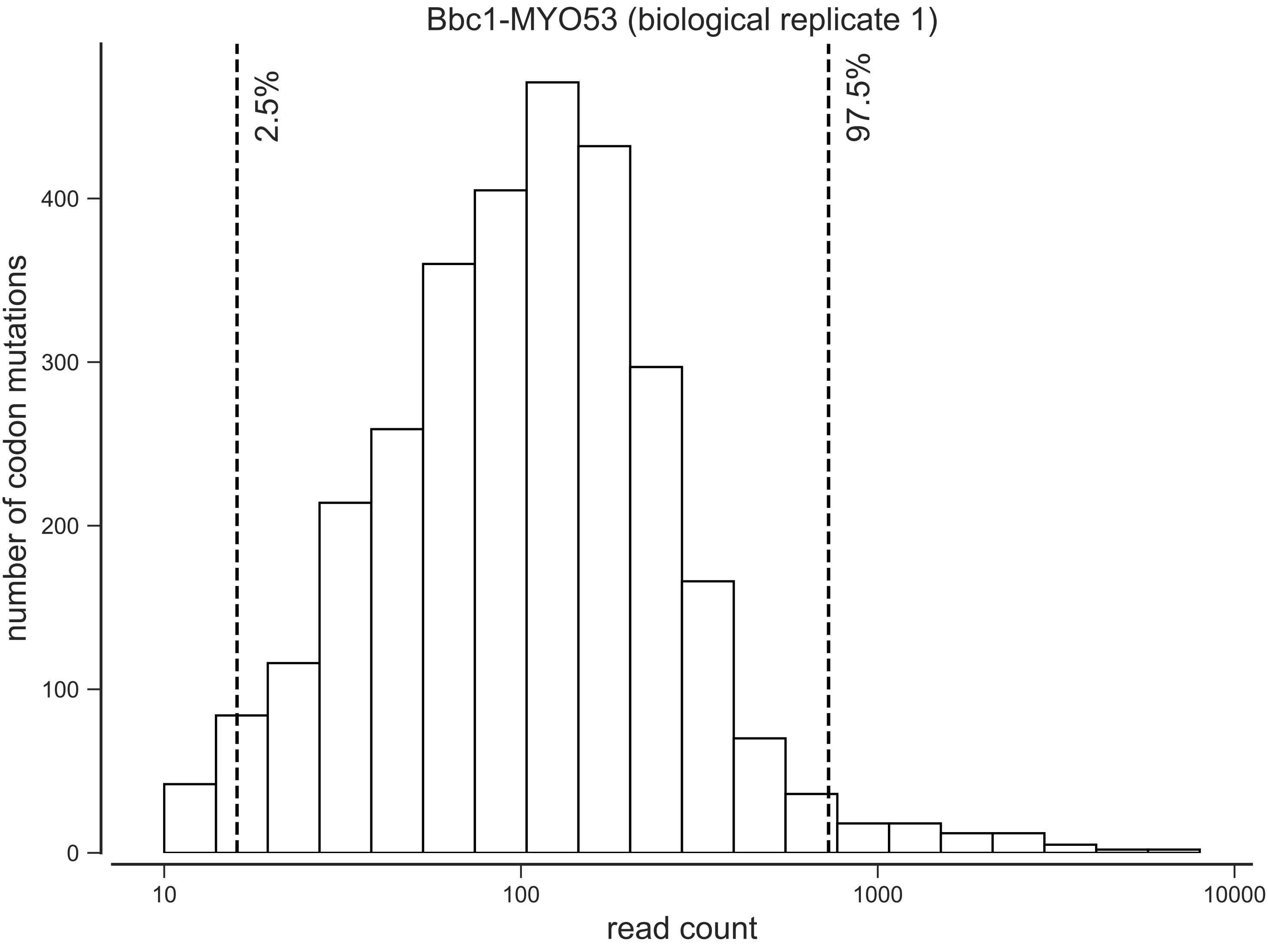

Bbc1-MYO53 (biological replicate 1)
2.5%
97.5%
number of codon mutations
0
100
200
300
400
10
100
1000
10000
read count

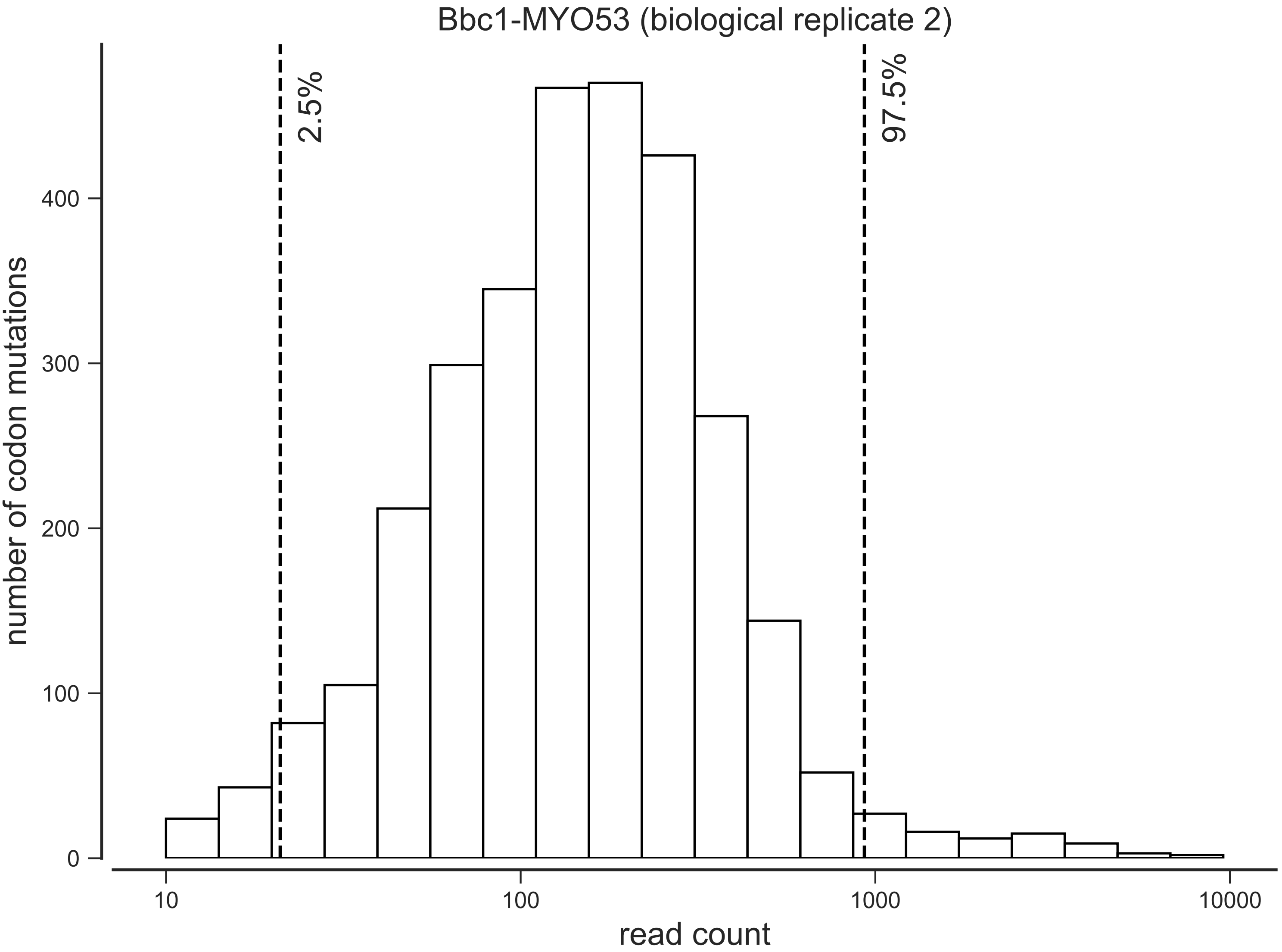

Bbc1-MYO53 (biological replicate 2)
2.5%
97.5%
number of codon mutations
0
100
200
300
400
10
100
1000
10000
read count

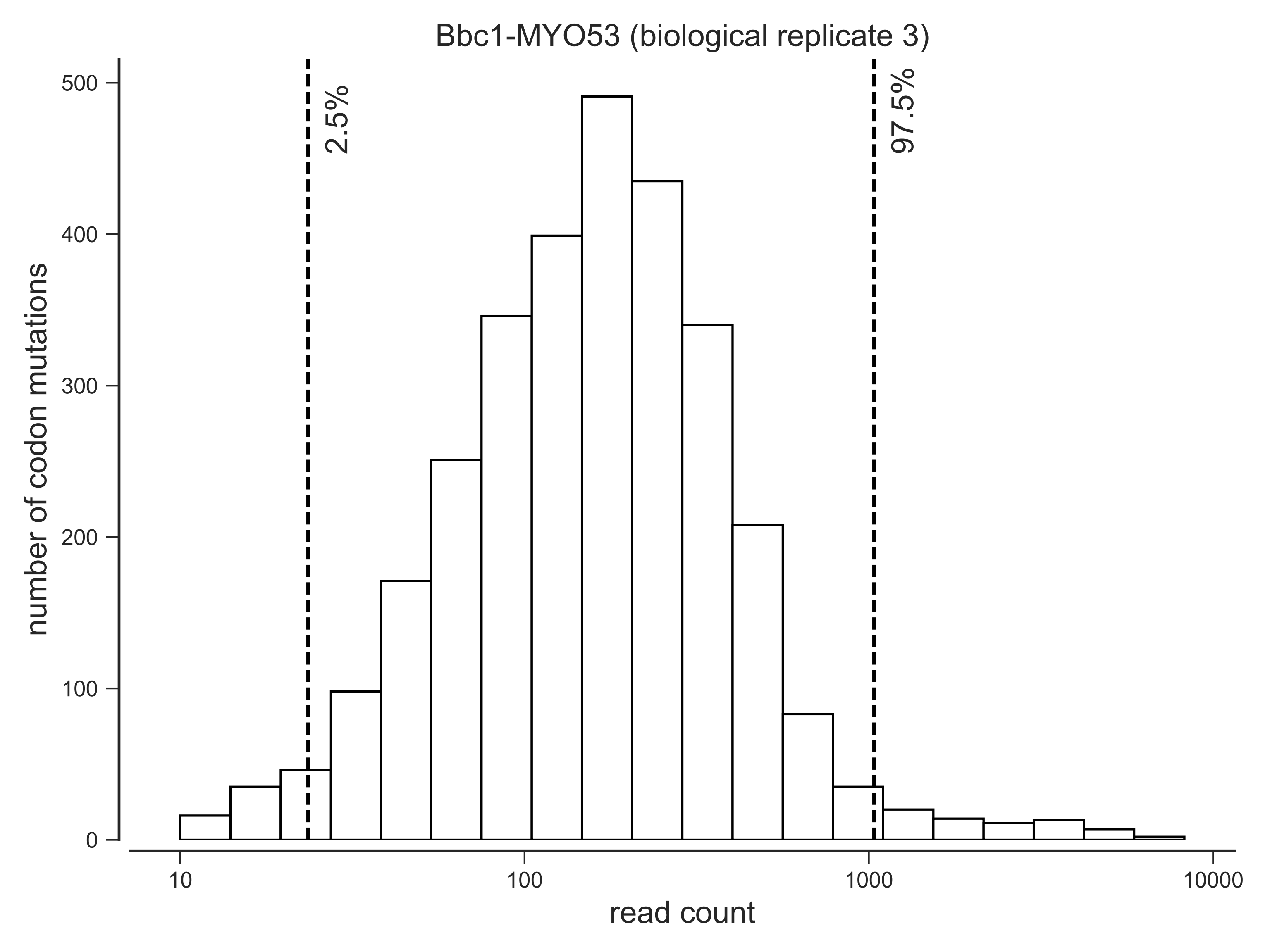

Bbc1-MYO53 (biological replicate 3)
number of codon mutations
read count
2.5%
97.5%
0
100
200
300
400
500
10
100
1000
10000

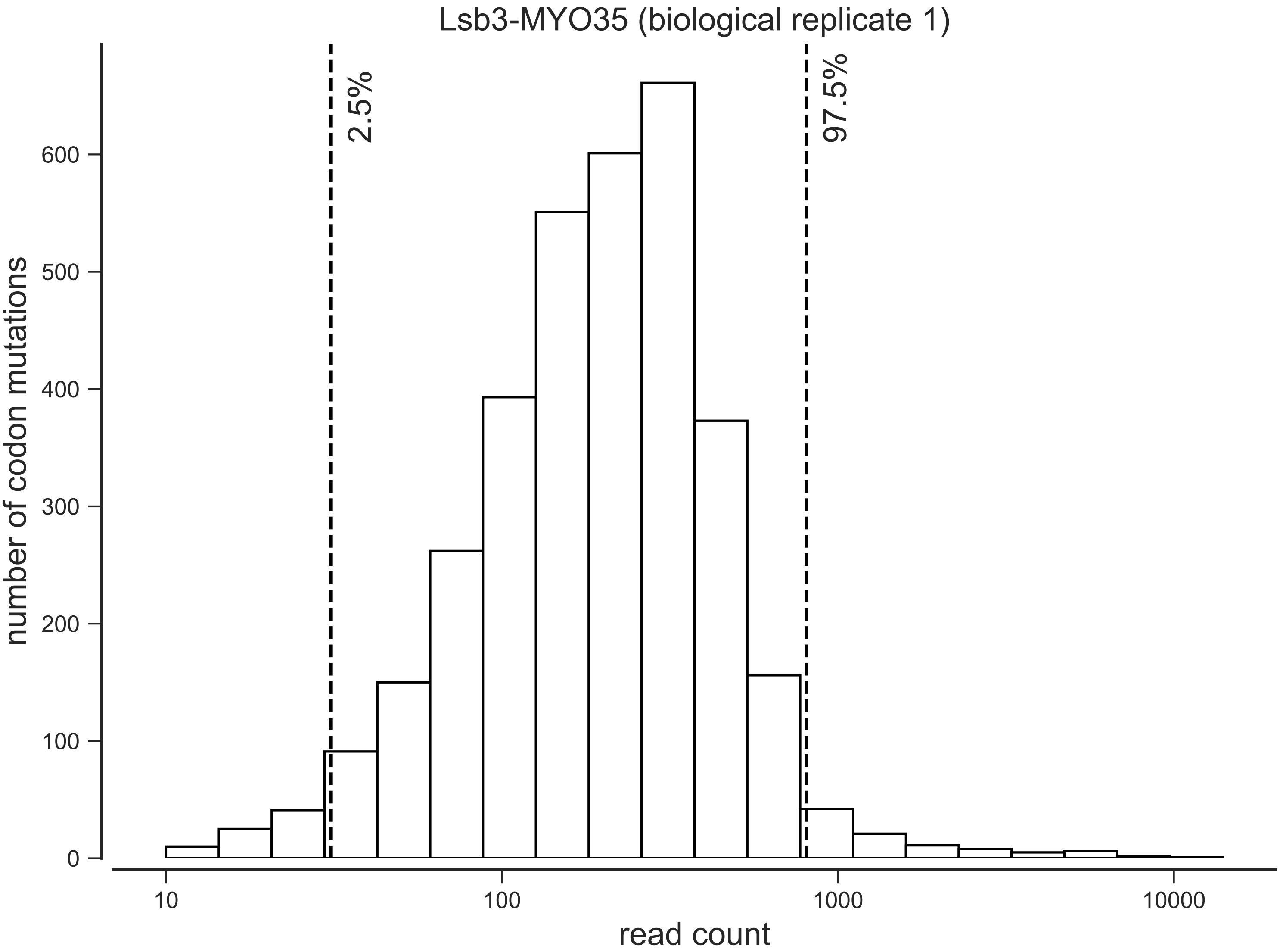

Lsb3-MYO35 (biological replicate 1)
2.5%
97.5%
number of codon mutations
0
100
200
300
400
500
600
10
100
1000
10000
read count

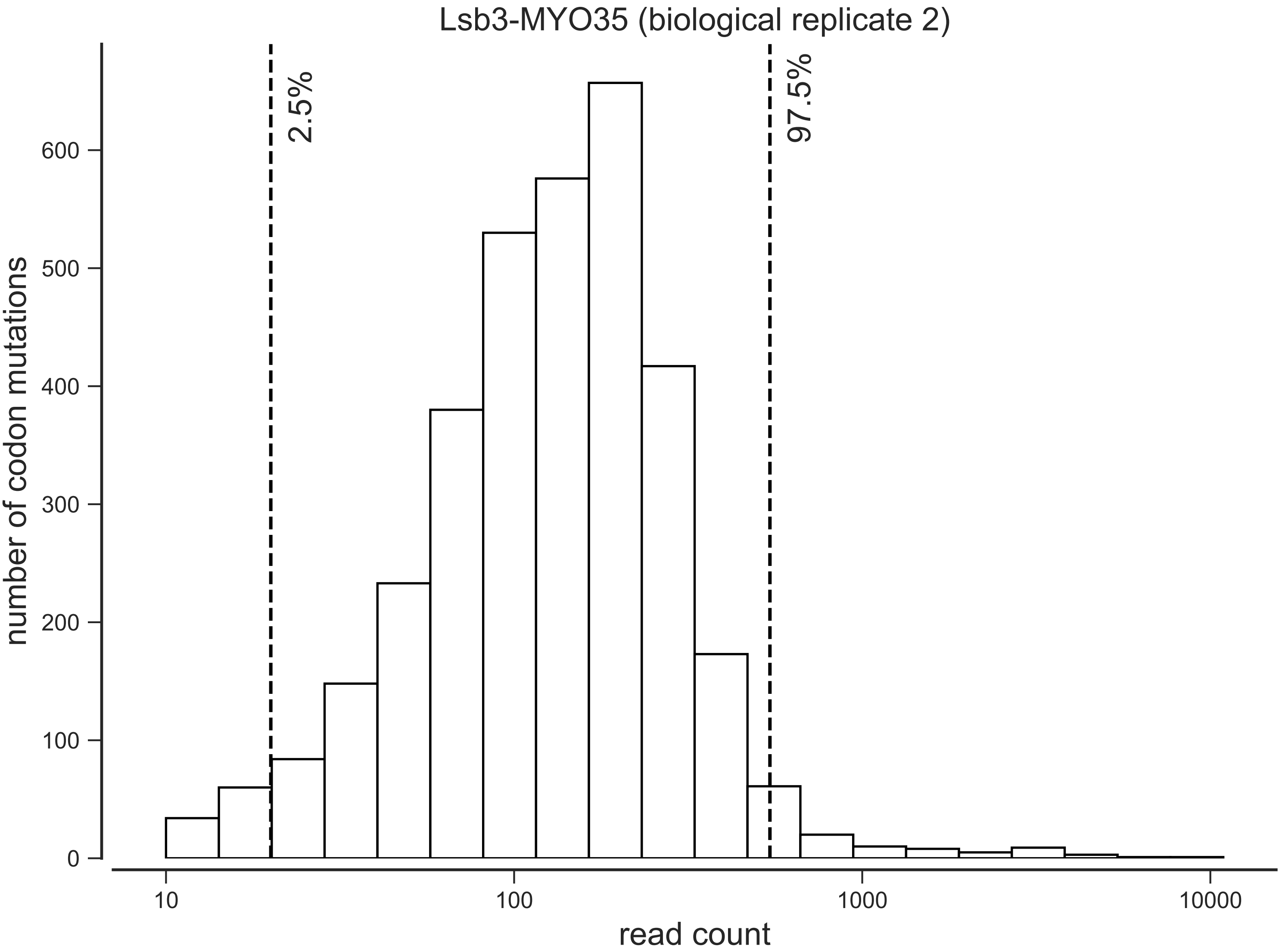

Lsb3-MYO35 (biological replicate 2)
2.5%
97.5%
number of codon mutations
0
100
200
300
400
500
600
10
100
1000
10000
read count

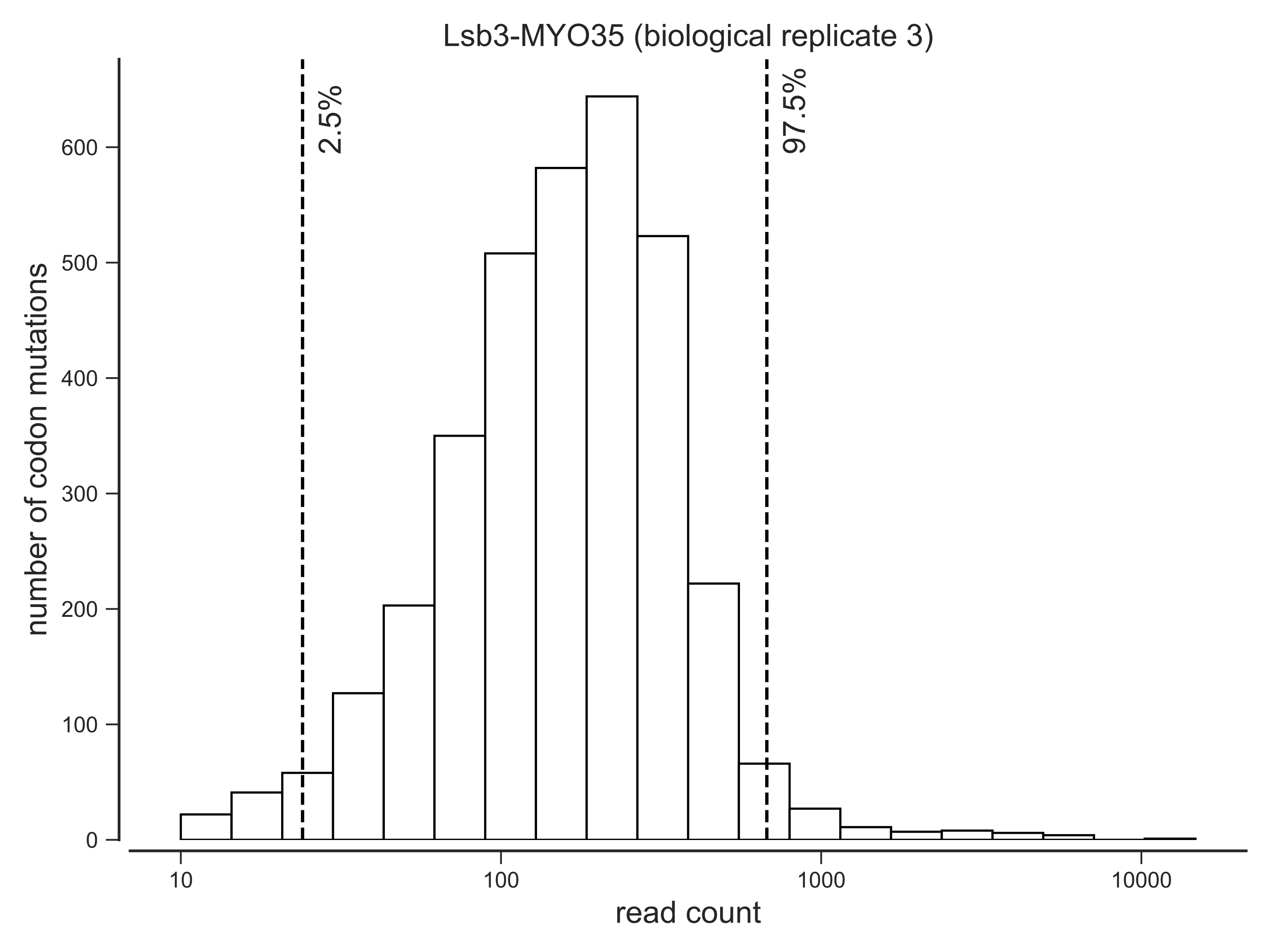

Lsb3-MYO35 (biological replicate 3)
2.5%
97.5%
number of codon mutations
0
100
200
300
400
500
600
10
100
1000
10000
read count

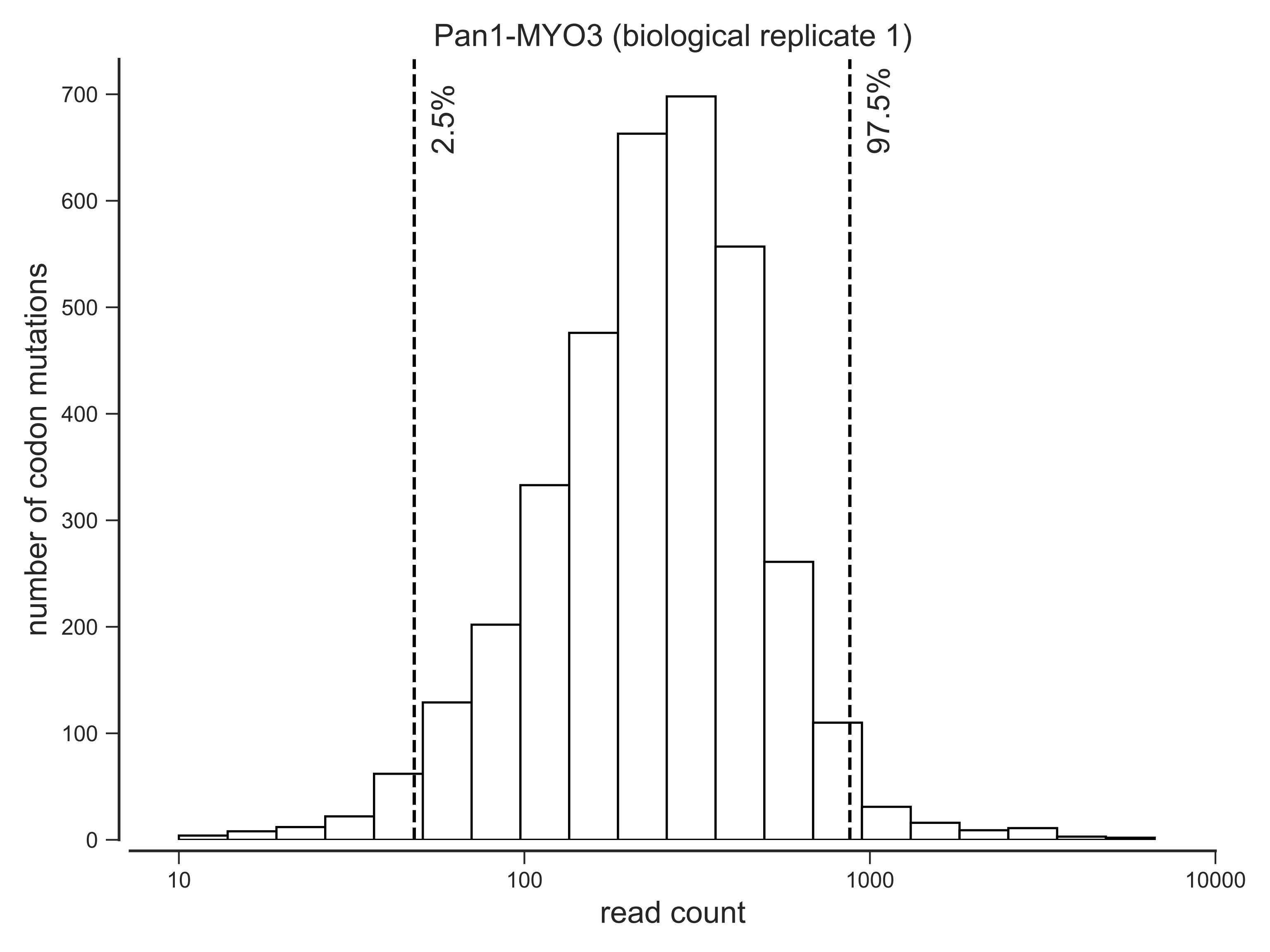
Pan1-MYO3 (biological replicate 1)
2.5%
97.5%
number of codon mutations
0
100
200
300
400
500
600
700
10
100
1000
10000
read count

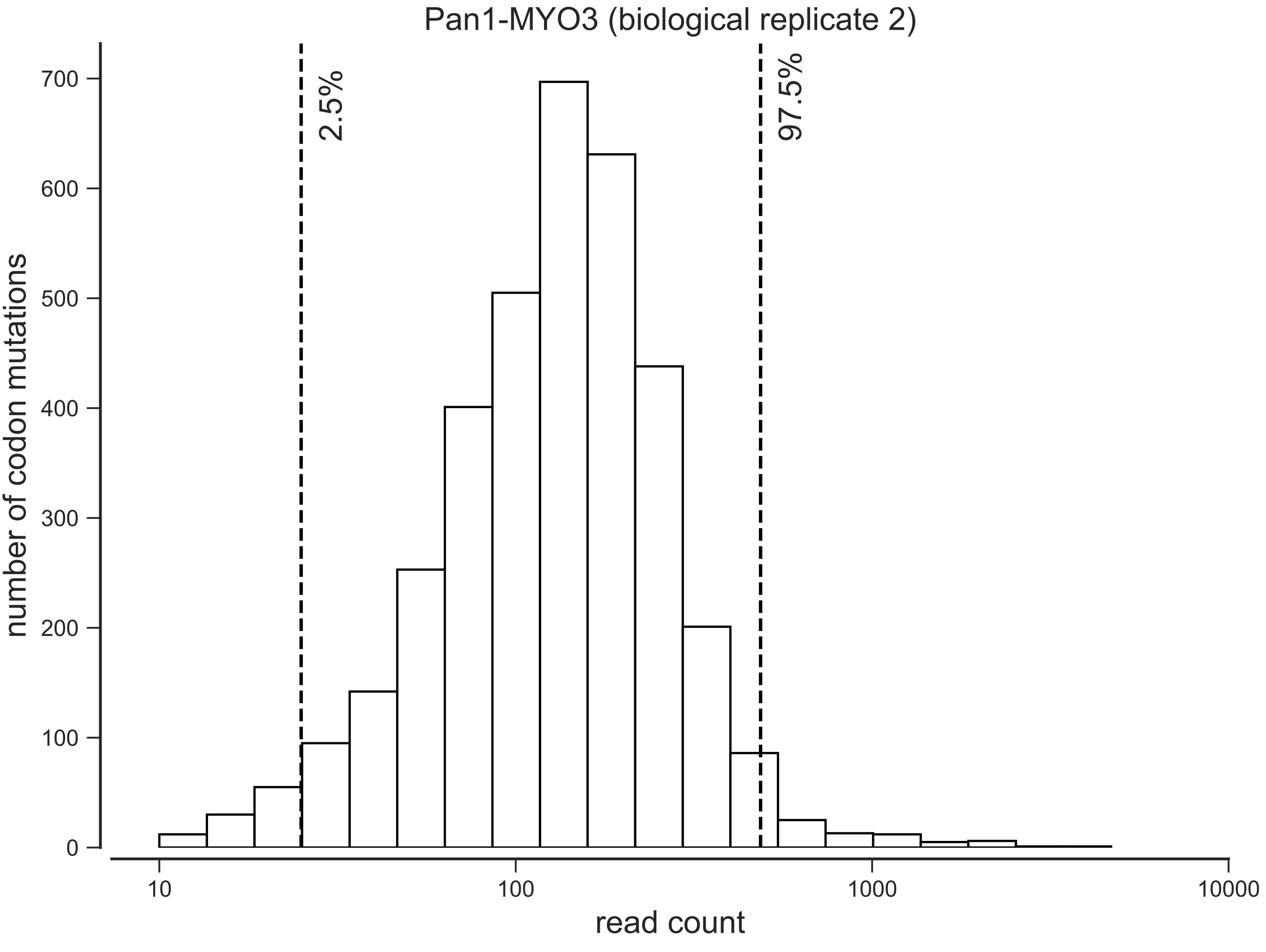

Pan1-MYO3 (biological replicate 2)
2.5%
97.5%
number of codon mutations
0
100
200
300
400
500
600
700
10
100
1000
10000
read count

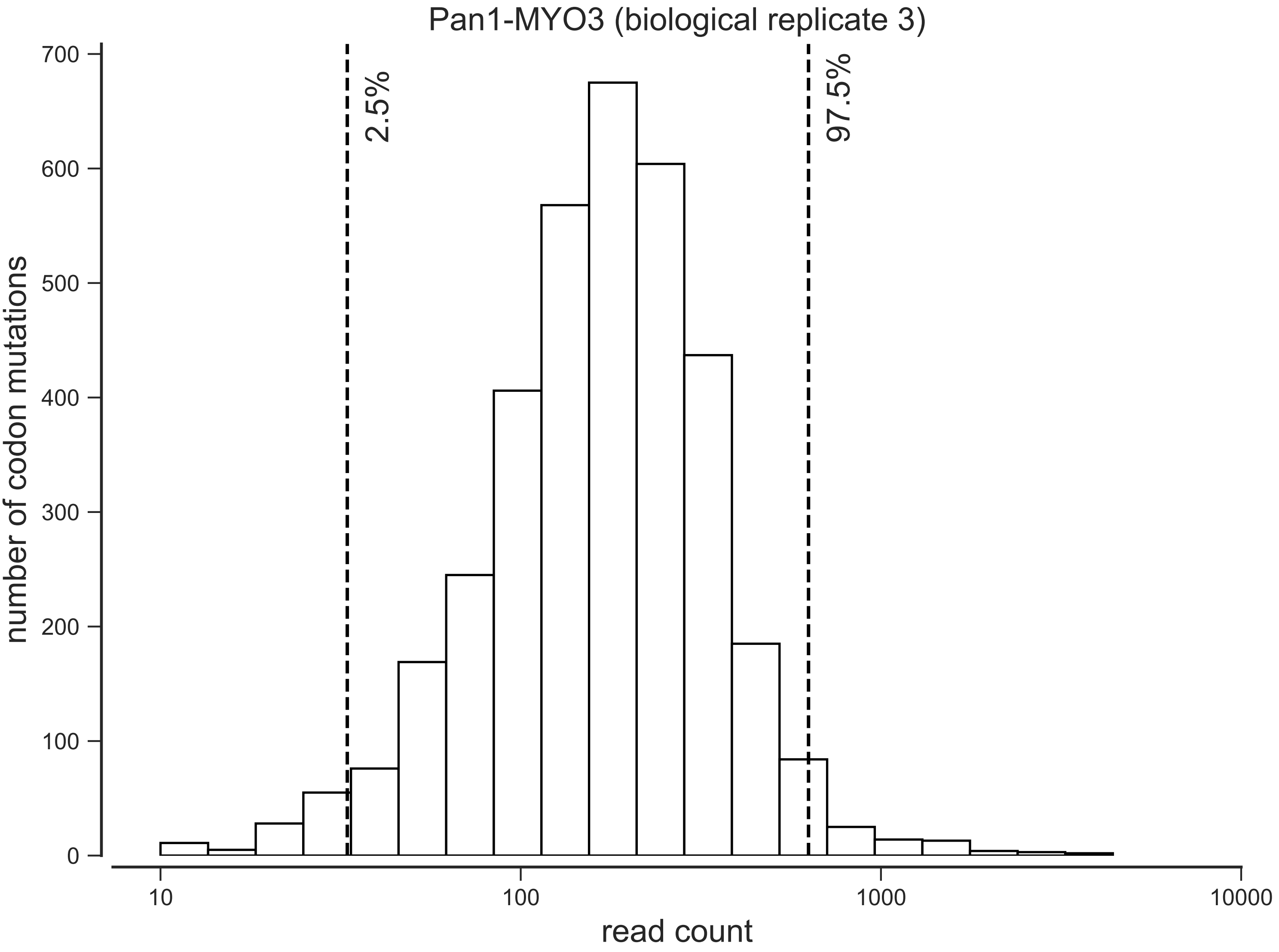
Pan1-MYO3 (biological replicate 3)
number of codon mutations
0
100
200
300
400
500
600
700
2.5%
97.5%
10
100
1000
10000
read count

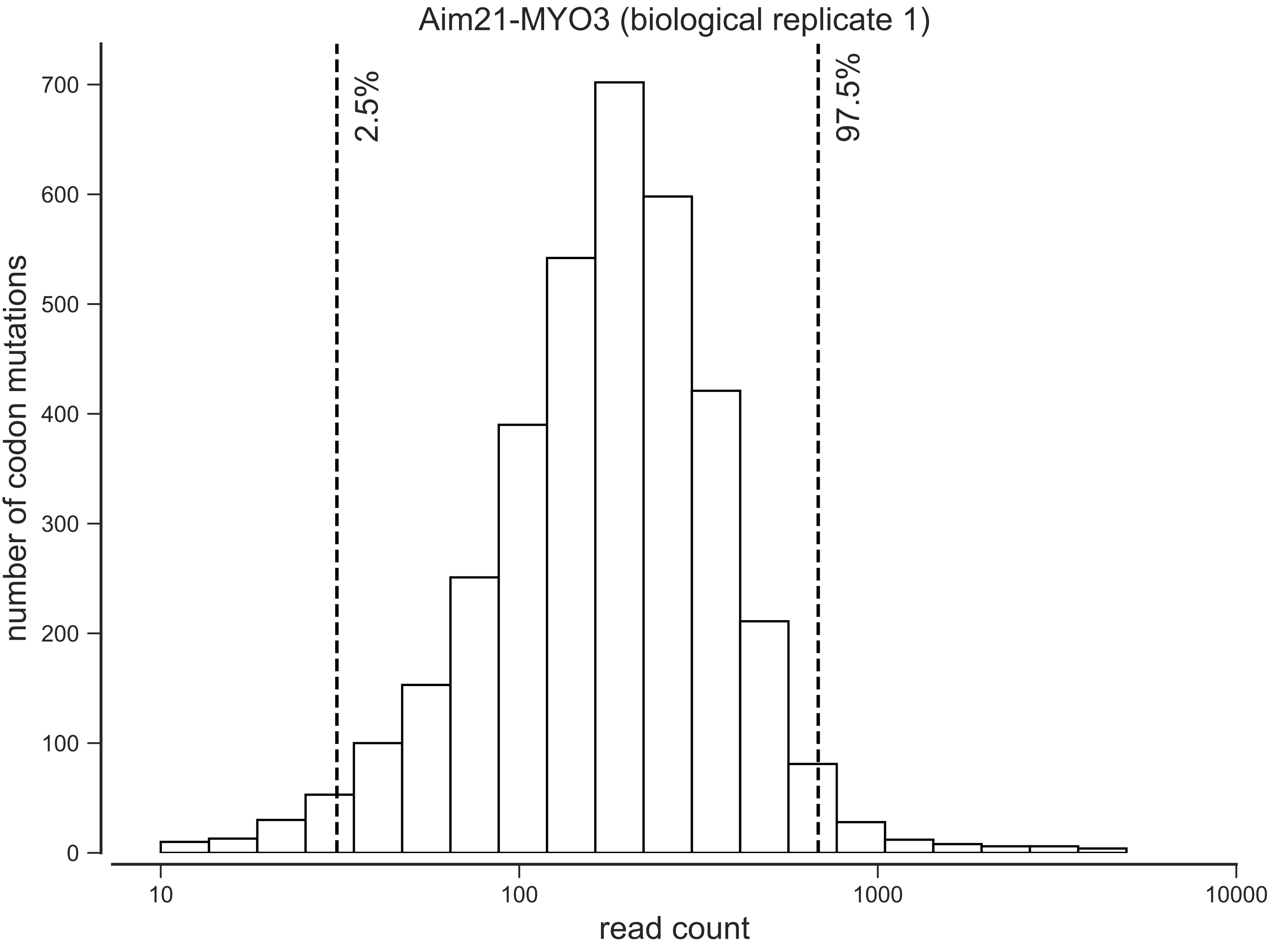

Aim21-MYO3 (biological replicate 1)
2.5%
97.5%
number of codon mutations
700
600
500
400
300
200
100
0
10
100
1000
10000
read count

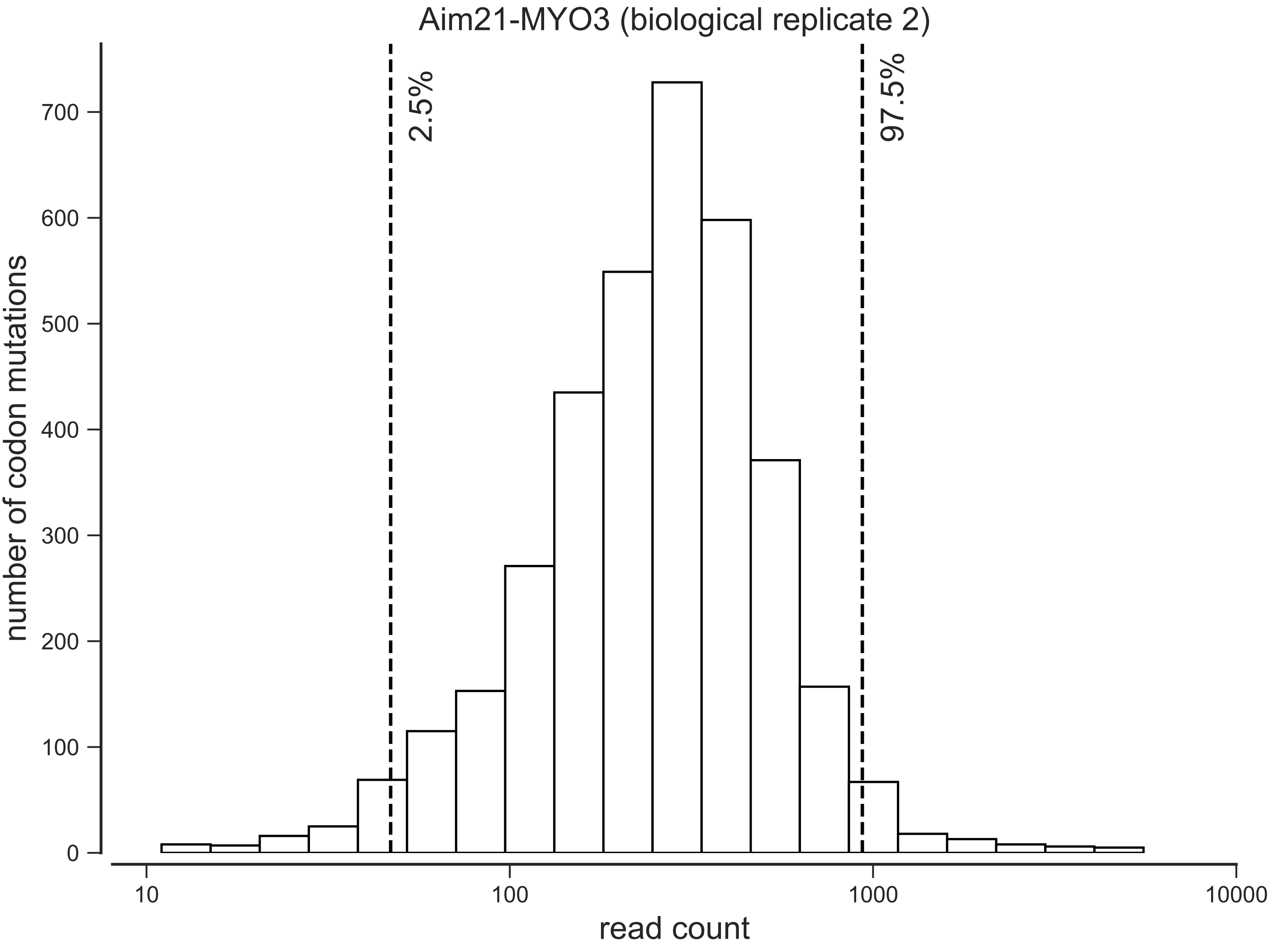
Aim21-MYO3 (biological replicate 2)
2.5%
97.5%
number of codon mutations
0
100
200
300
400
500
600
700
10
100
1000
10000
read count

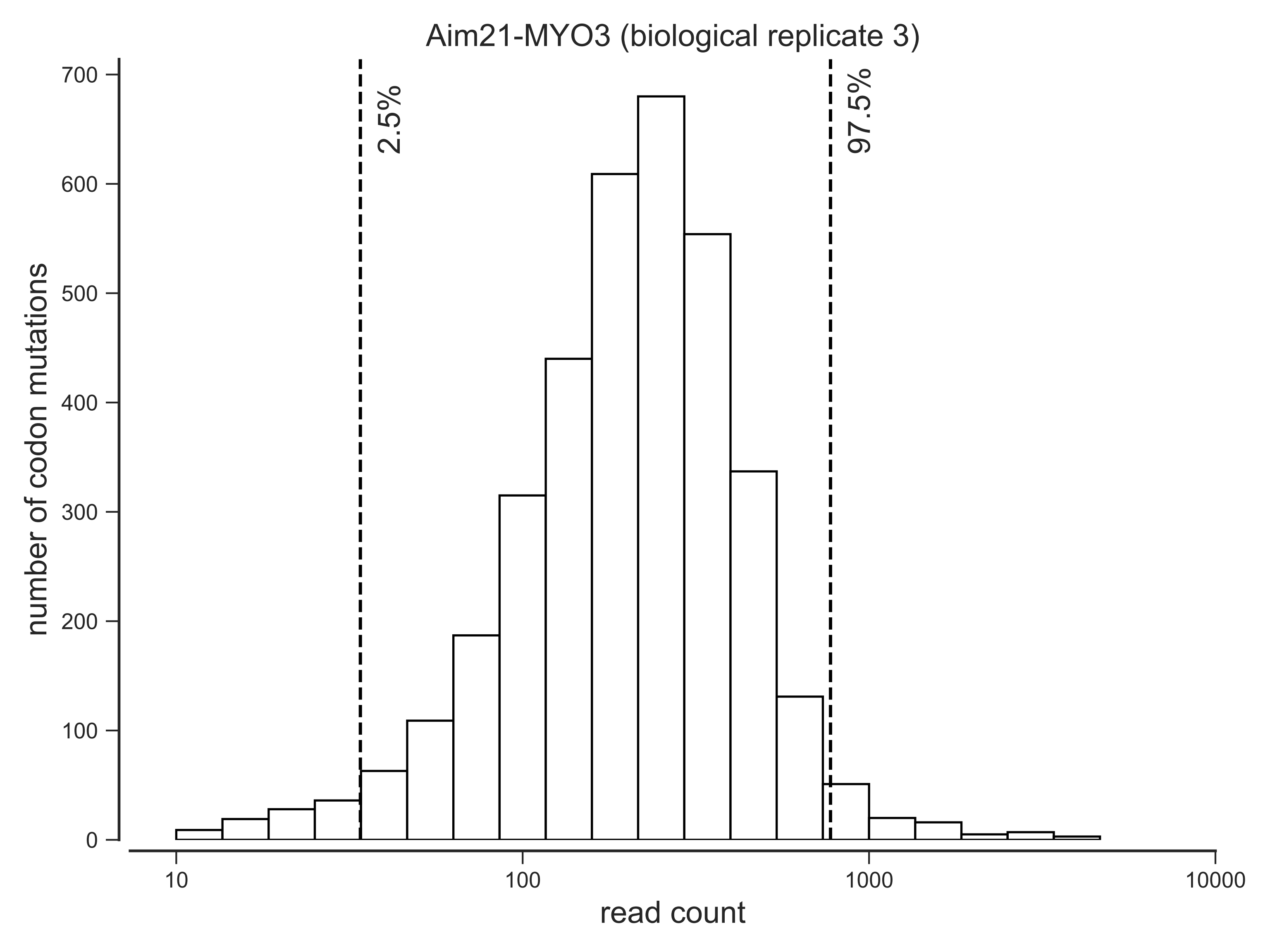

Aim21-MYO3 (biological replicate 3)
2.5%
97.5%
number of codon mutations
read count
0
100
200
300
400
500
600
700
10
100
1000
10000

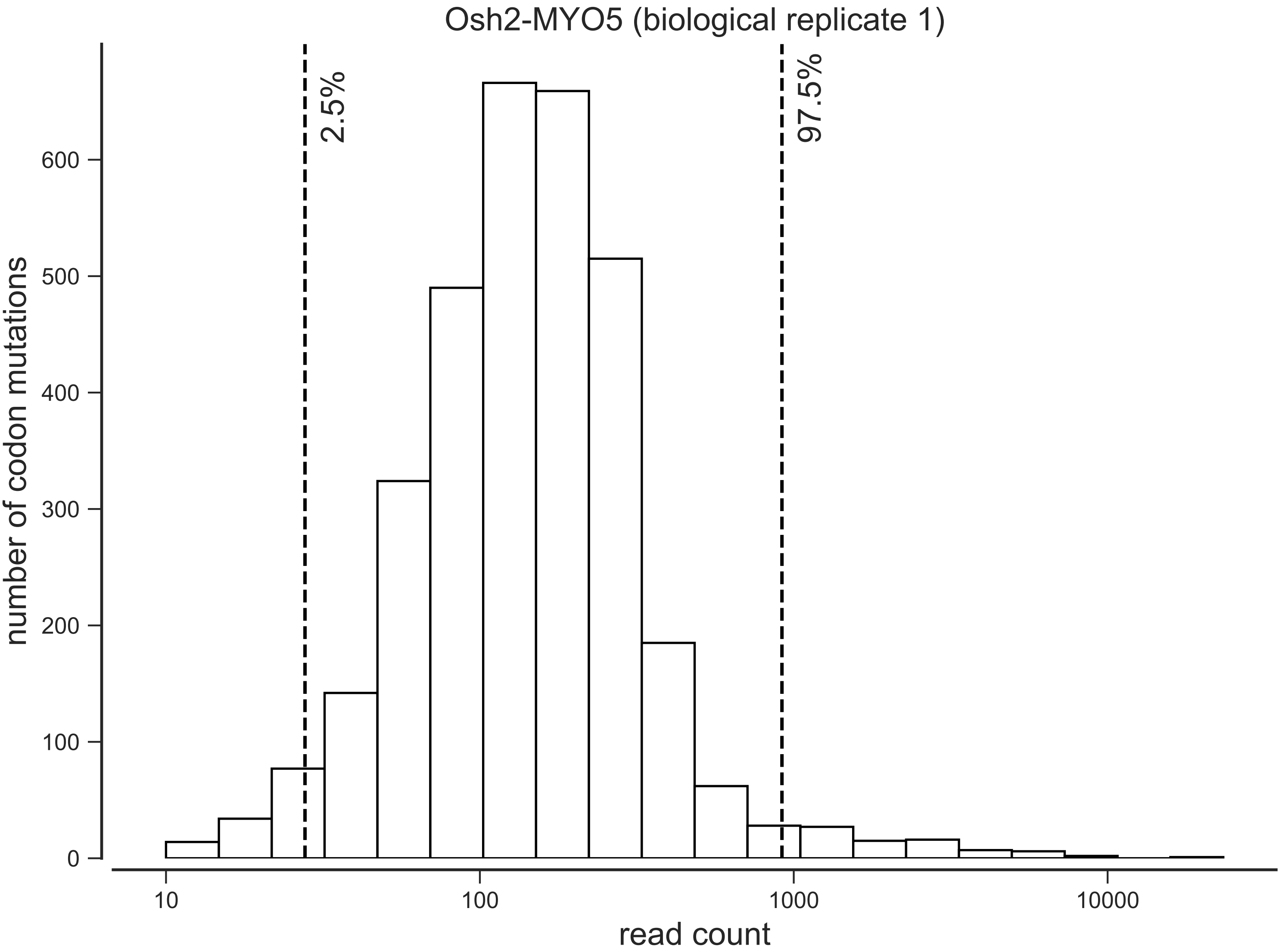
Osh2-MYO5 (biological replicate 1)
2.5%
97.5%
number of codon mutations
0
100
200
300
400
500
600
10
100
1000
10000
read count

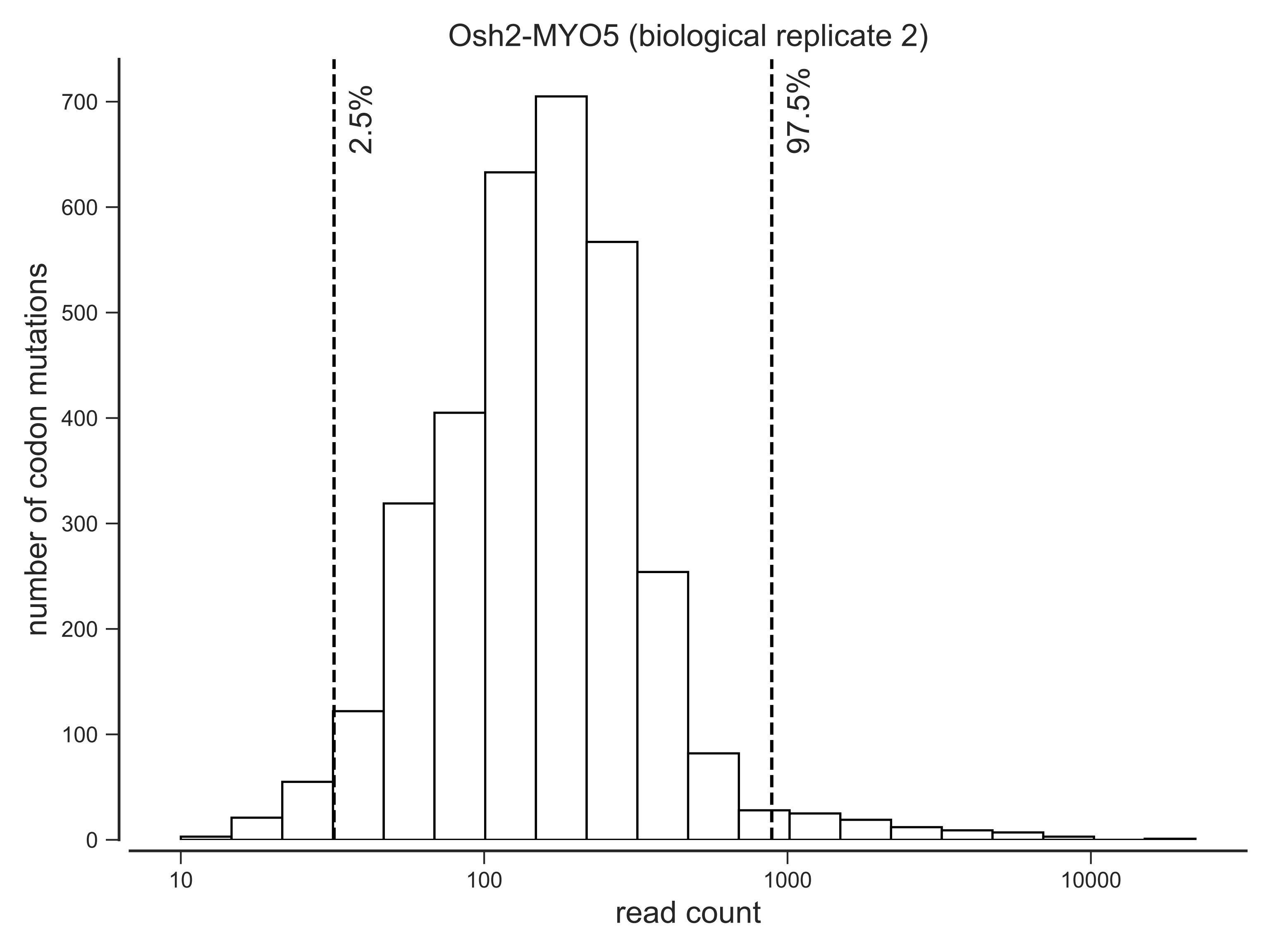

Osh2-MYO5 (biological replicate 2)
number of codon mutations
read count
2.5%
97.5%
0
100
200
300
400
500
600
700
10
100
1000
10000

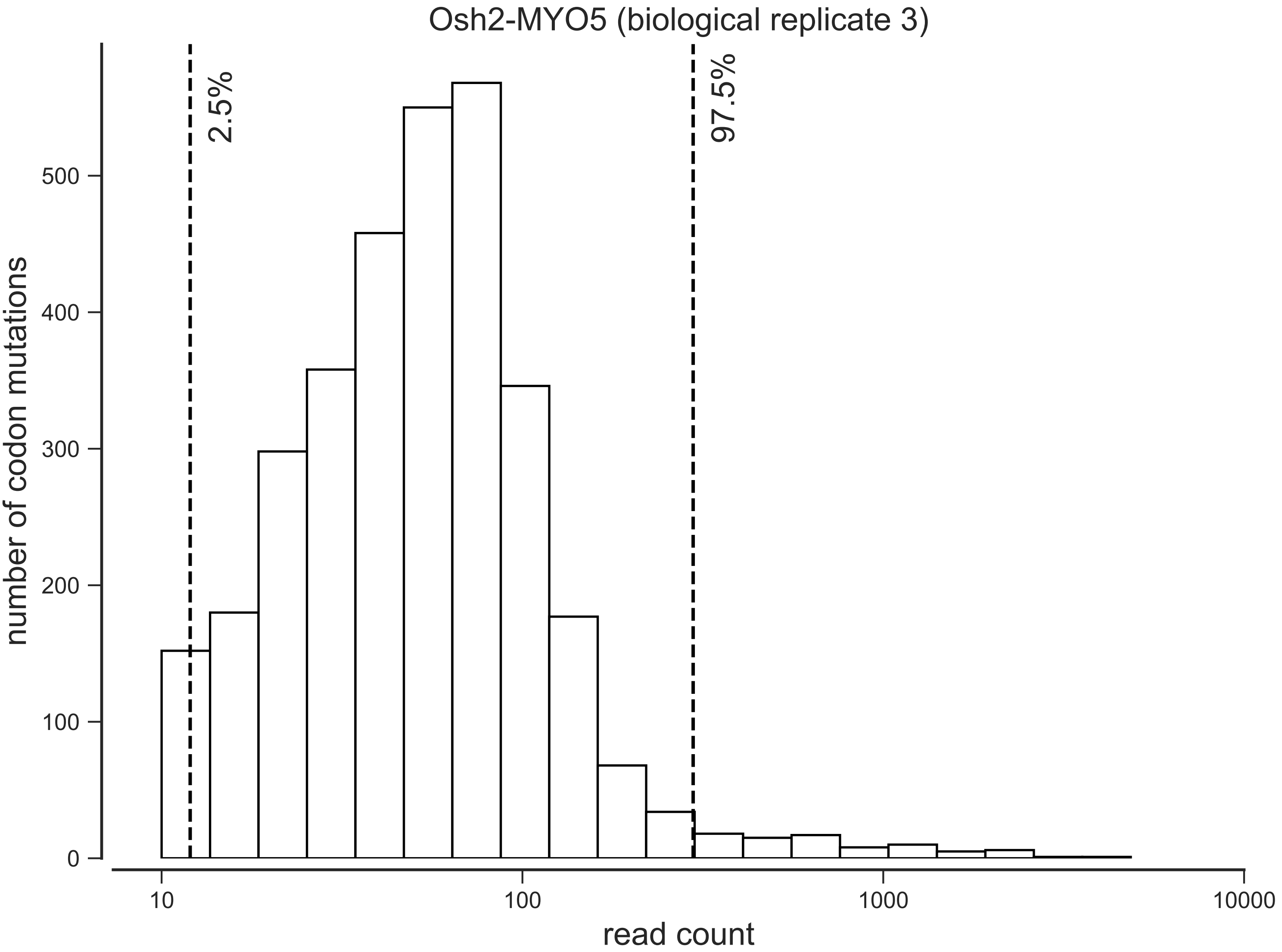

Osh2-MYO5 (biological replicate 3)
2.5%
97.5%
0
100
200
300
400
500
number of codon mutations
10
100
1000
10000
read count

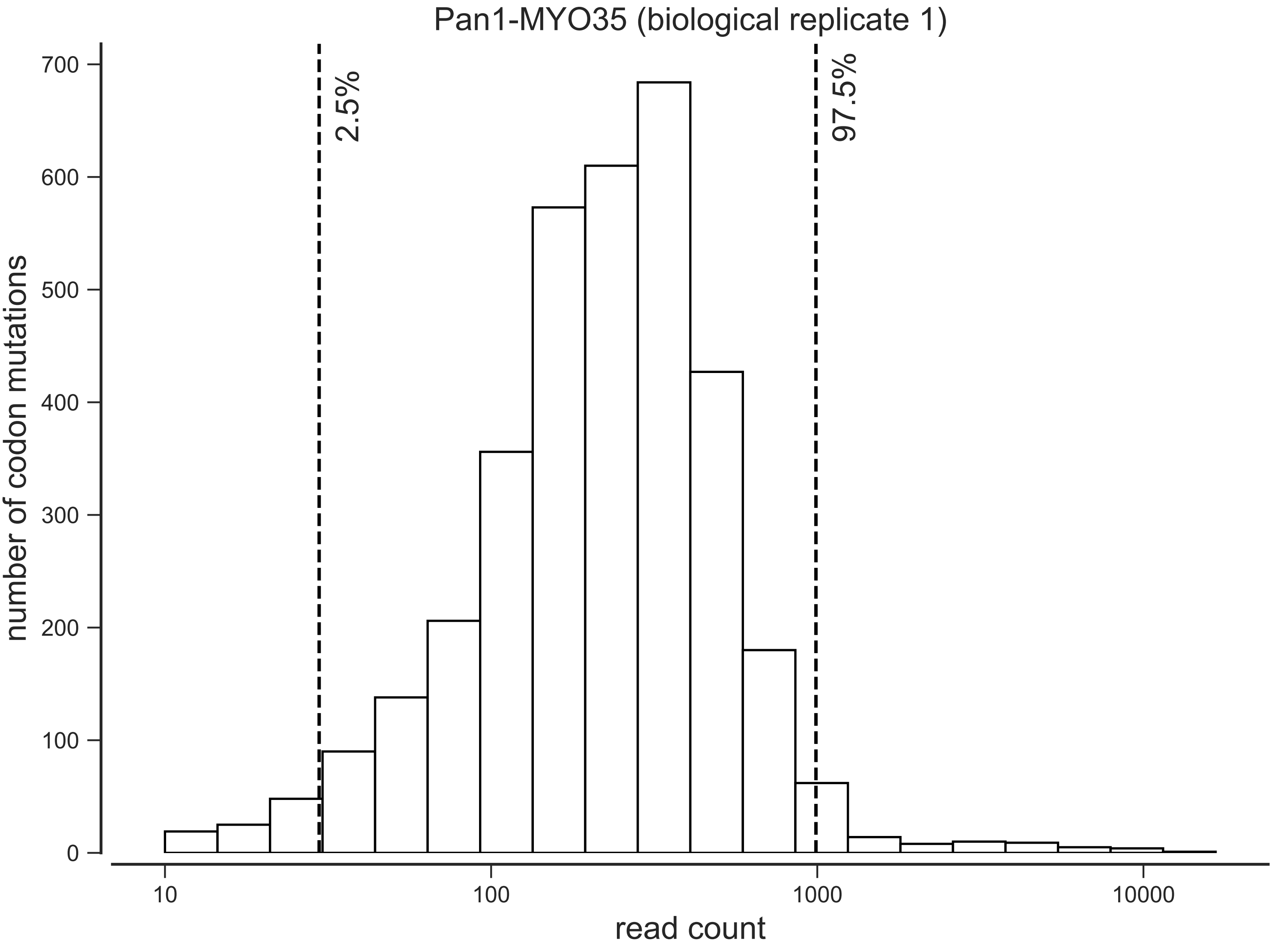
Pan1-MYO35 (biological replicate 1)
2.5%
97.5%
number of codon mutations
read count
0
100
200
300
400
500
600
700
10
100
1000
10000

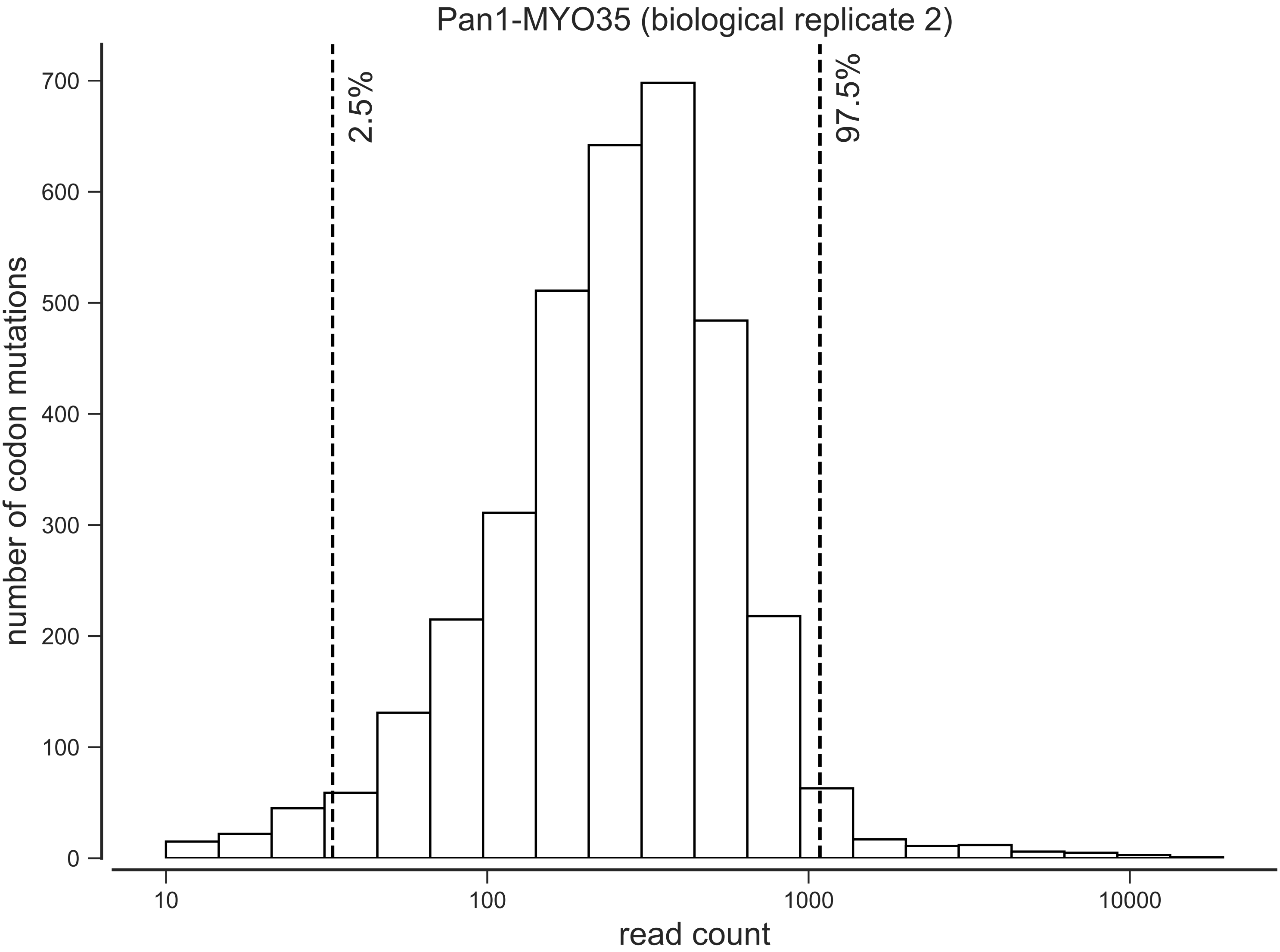

Pan1-MYO35 (biological replicate 2)
2.5%
97.5%
number of codon mutations
read count
0
100
200
300
400
500
600
700
10
100
1000
10000

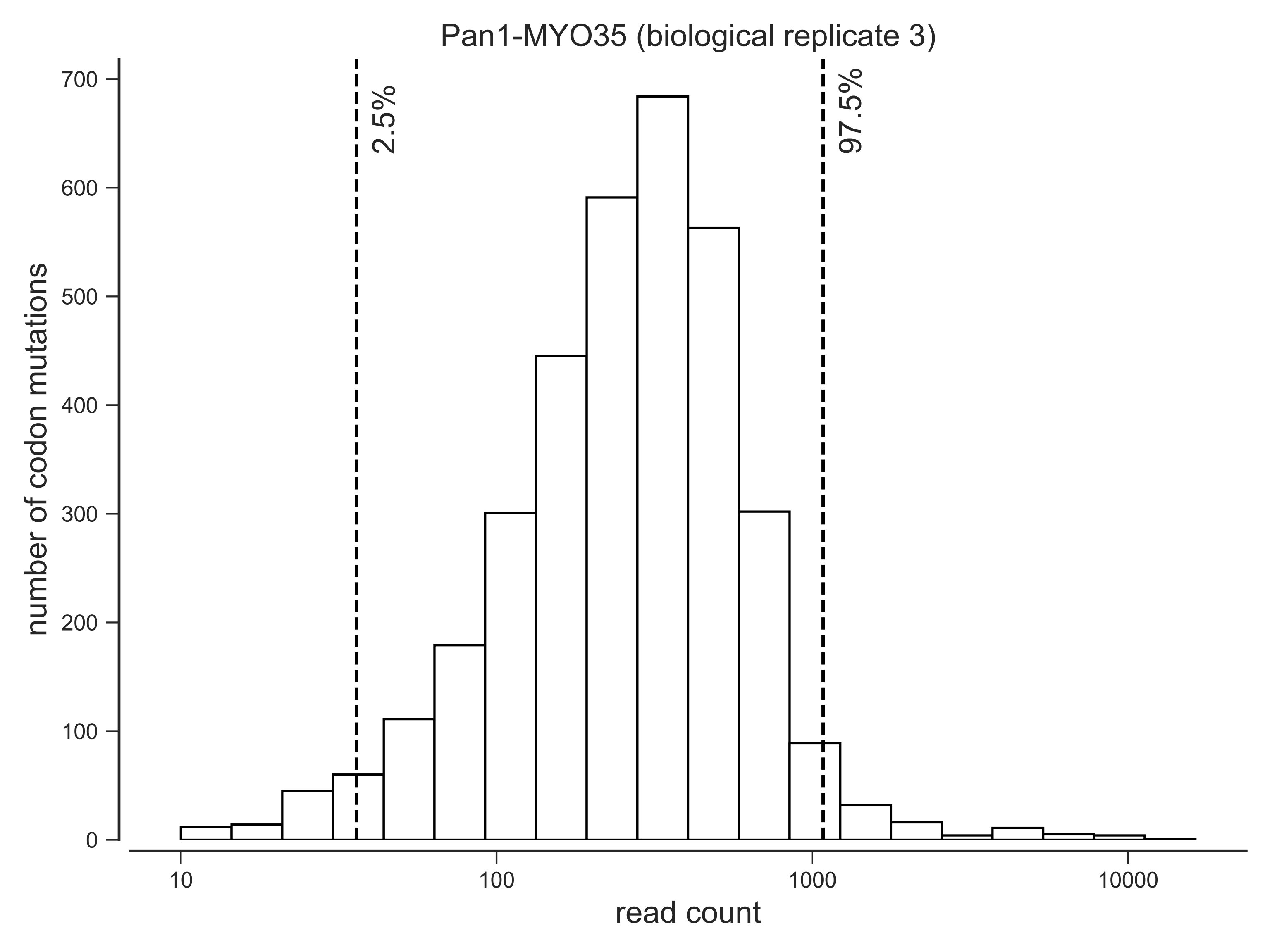

Pan1-MYO35 (biological replicate 3)
2.5%
97.5%
number of codon mutations
0
100
200
300
400
500
600
700
10
100
1000
10000
read count

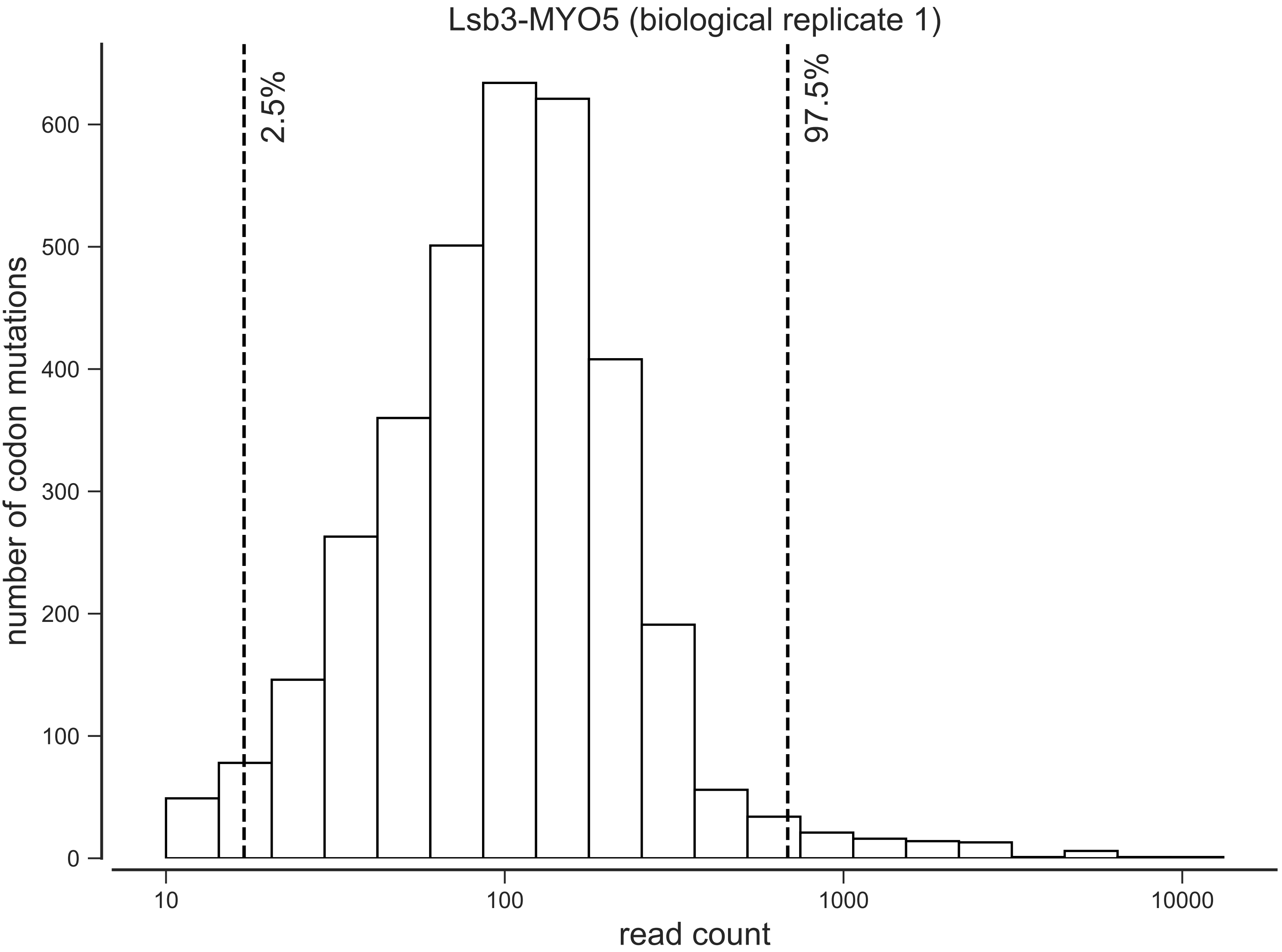
Lsb3-MYO5 (biological replicate 1)
2.5%
97.5%
number of codon mutations
0
100
200
300
400
500
600
10
100
1000
10000
read count

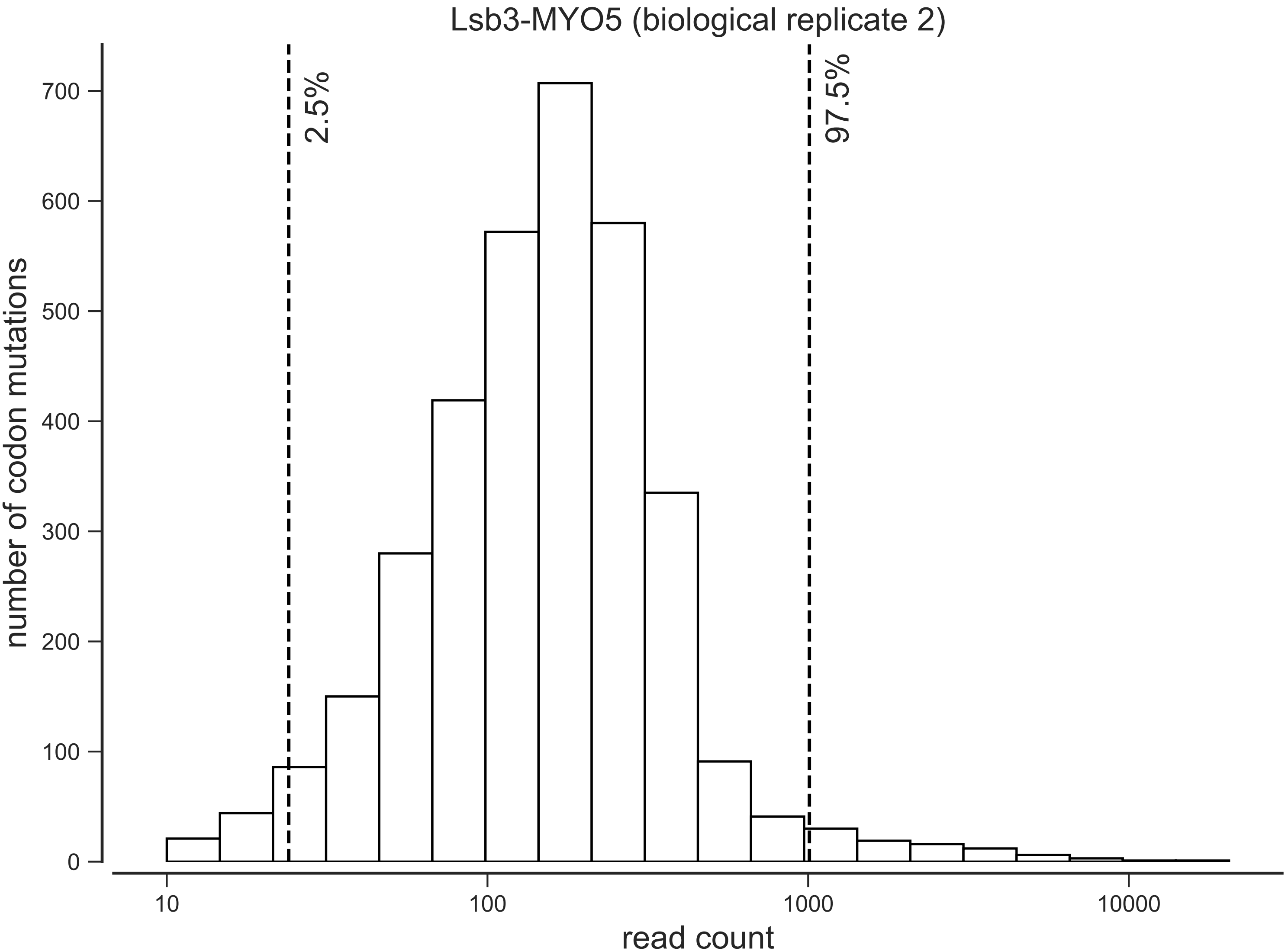

Lsb3-MYO5 (biological replicate 2)
2.5%
97.5%
number of codon mutations
0
100
200
300
400
500
600
700
10
100
1000
10000
read count

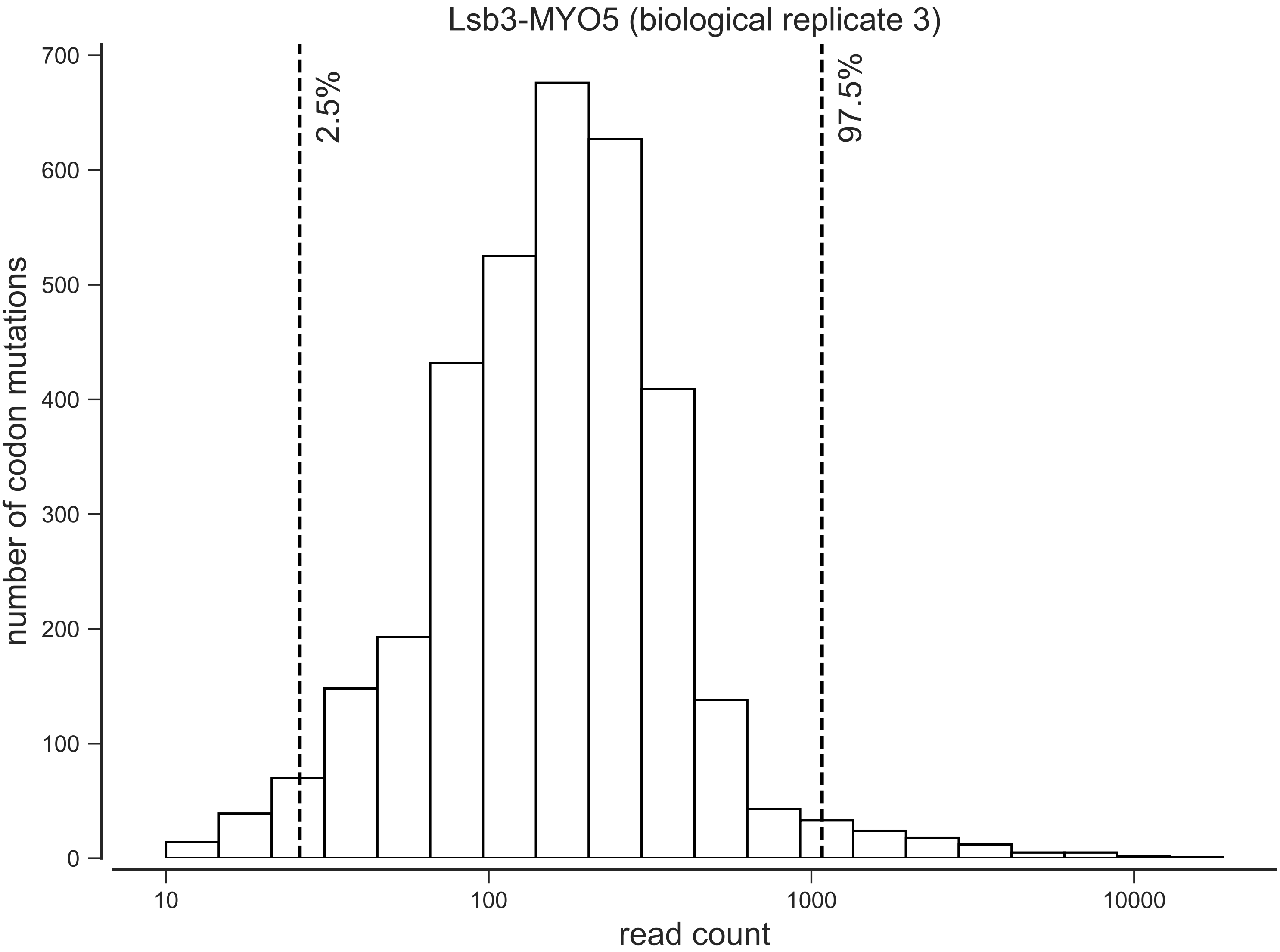
Lsb3-MYO5 (biological replicate 3)
2.5%
97.5%
number of codon mutations
0
100
200
300
400
500
600
700
10
100
1000
10000
read count

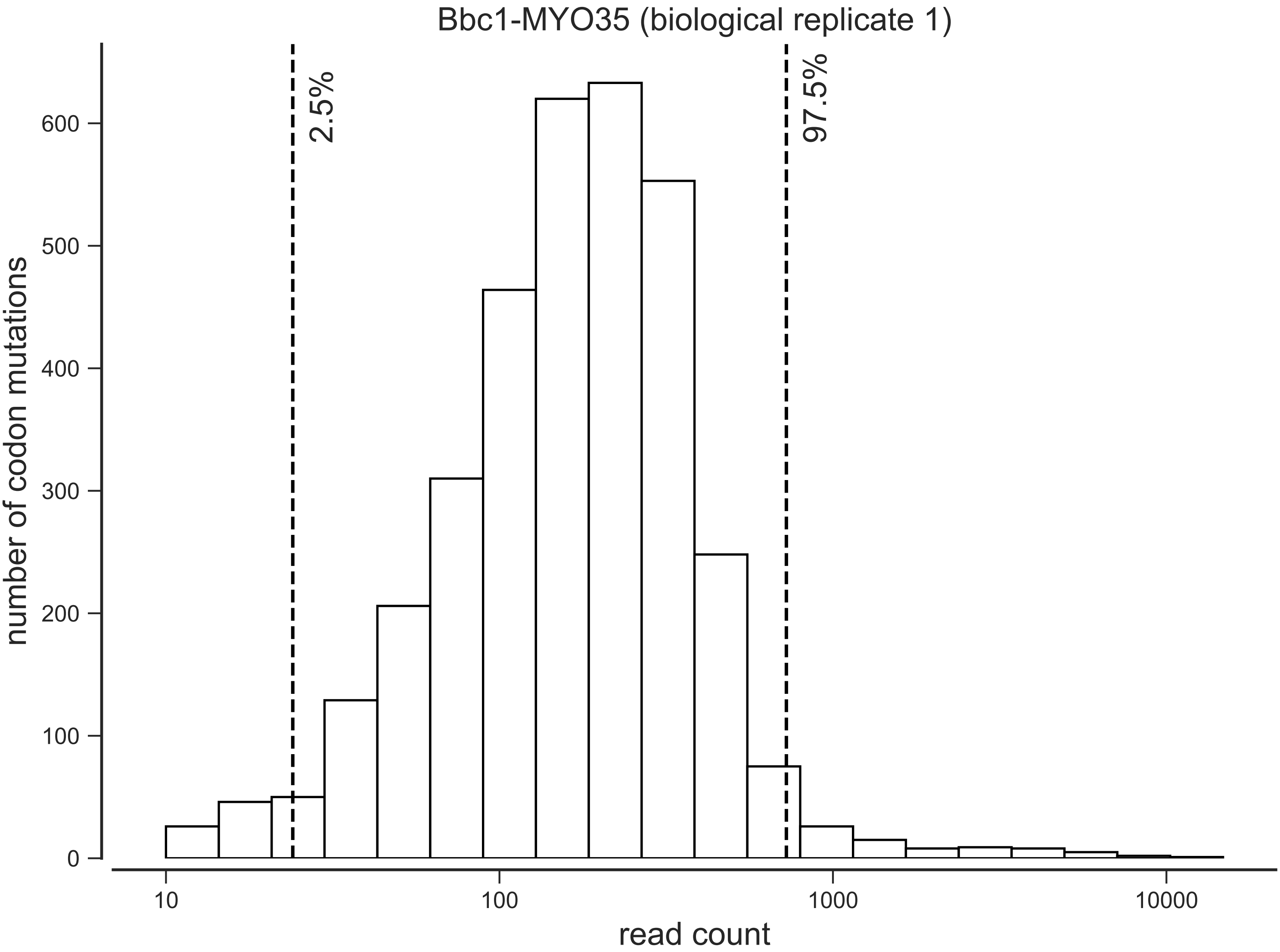
Bbc1-MYO35 (biological replicate 1)
2.5%
97.5%
number of codon mutations
0
100
200
300
400
500
600
10
100
1000
10000
read count

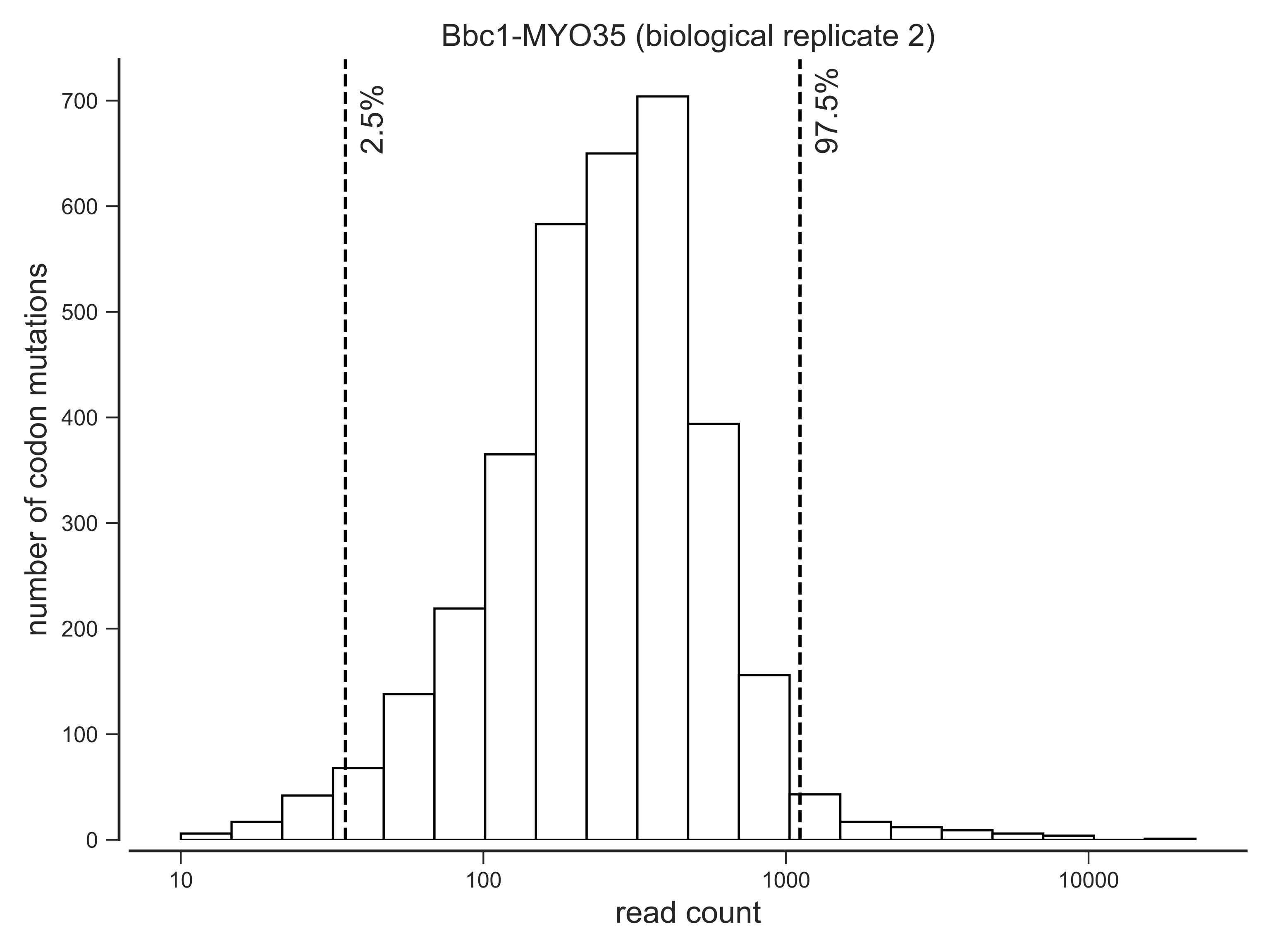

Bbc1-MYO35 (biological replicate 2)
2.5%
97.5%
number of codon mutations
0
100
200
300
400
500
600
700
10
100
1000
10000
read count

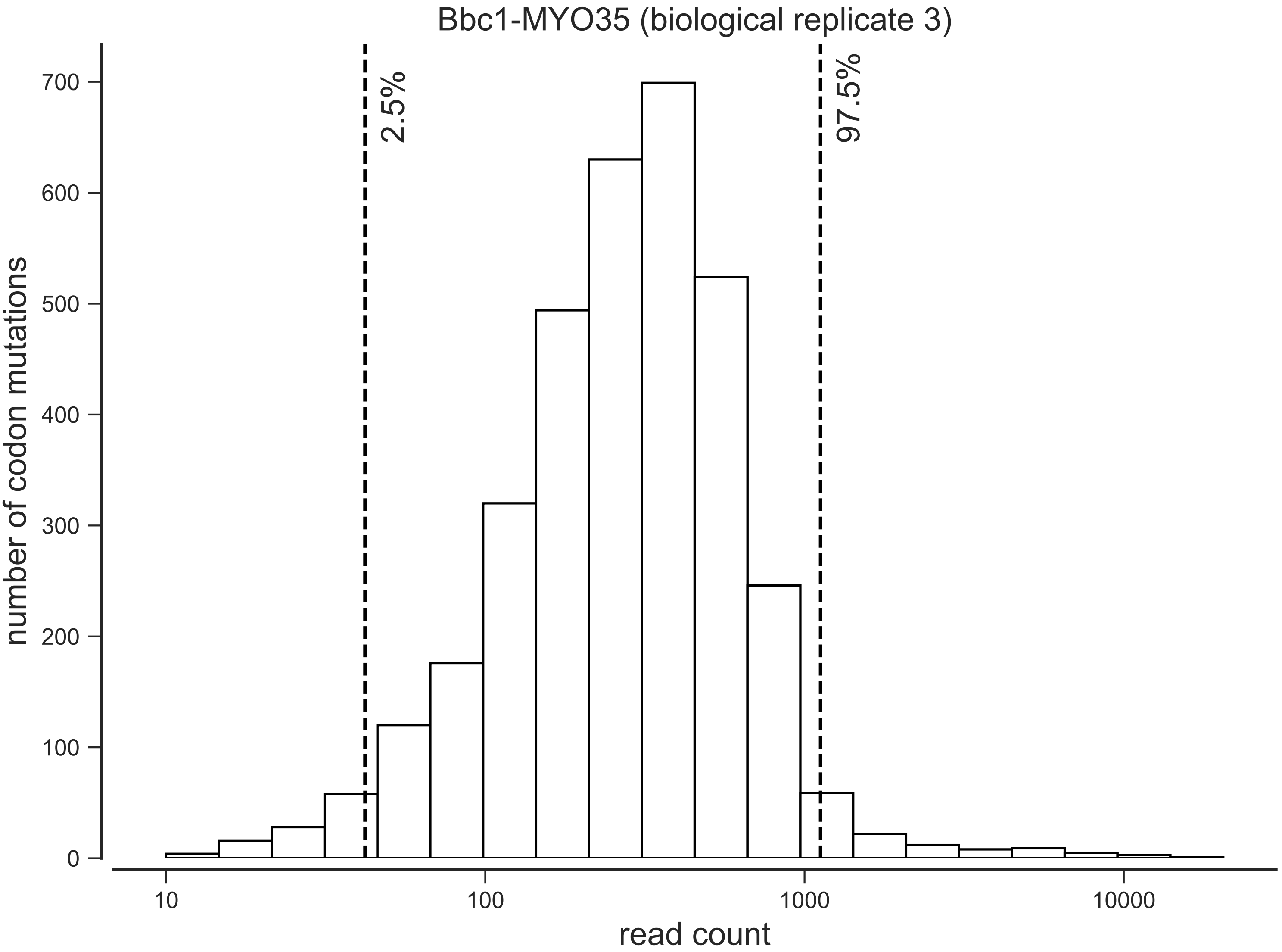

Bbc1-MYO35 (biological replicate 3)
2.5%
97.5%
number of codon mutations
0
100
200
300
400
500
600
700
10
100
1000
10000
read count

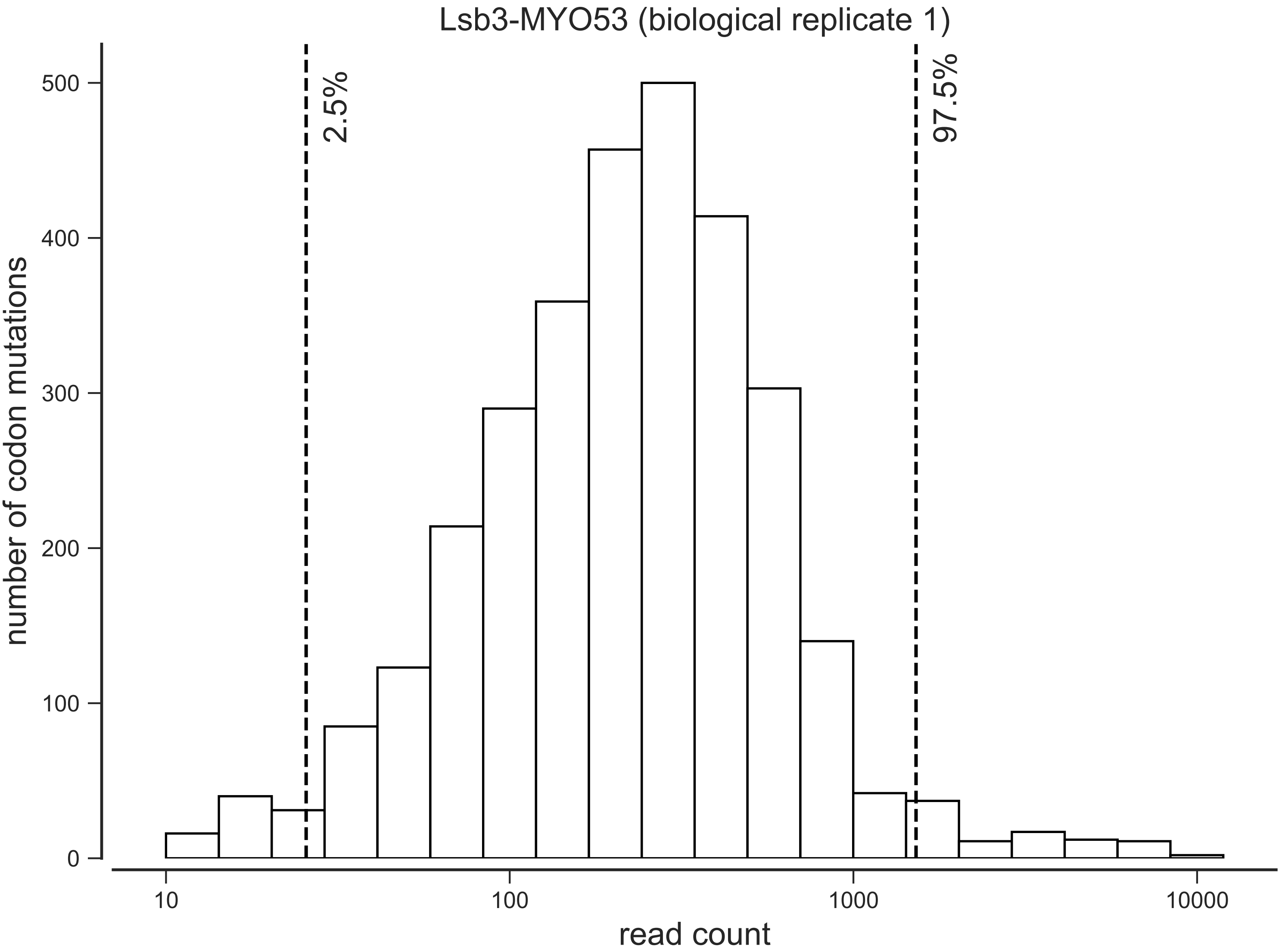
Lsb3-MYO53 (biological replicate 1)
2.5%
97.5%
number of codon mutations
0
100
200
300
400
500
10
100
1000
10000
read count

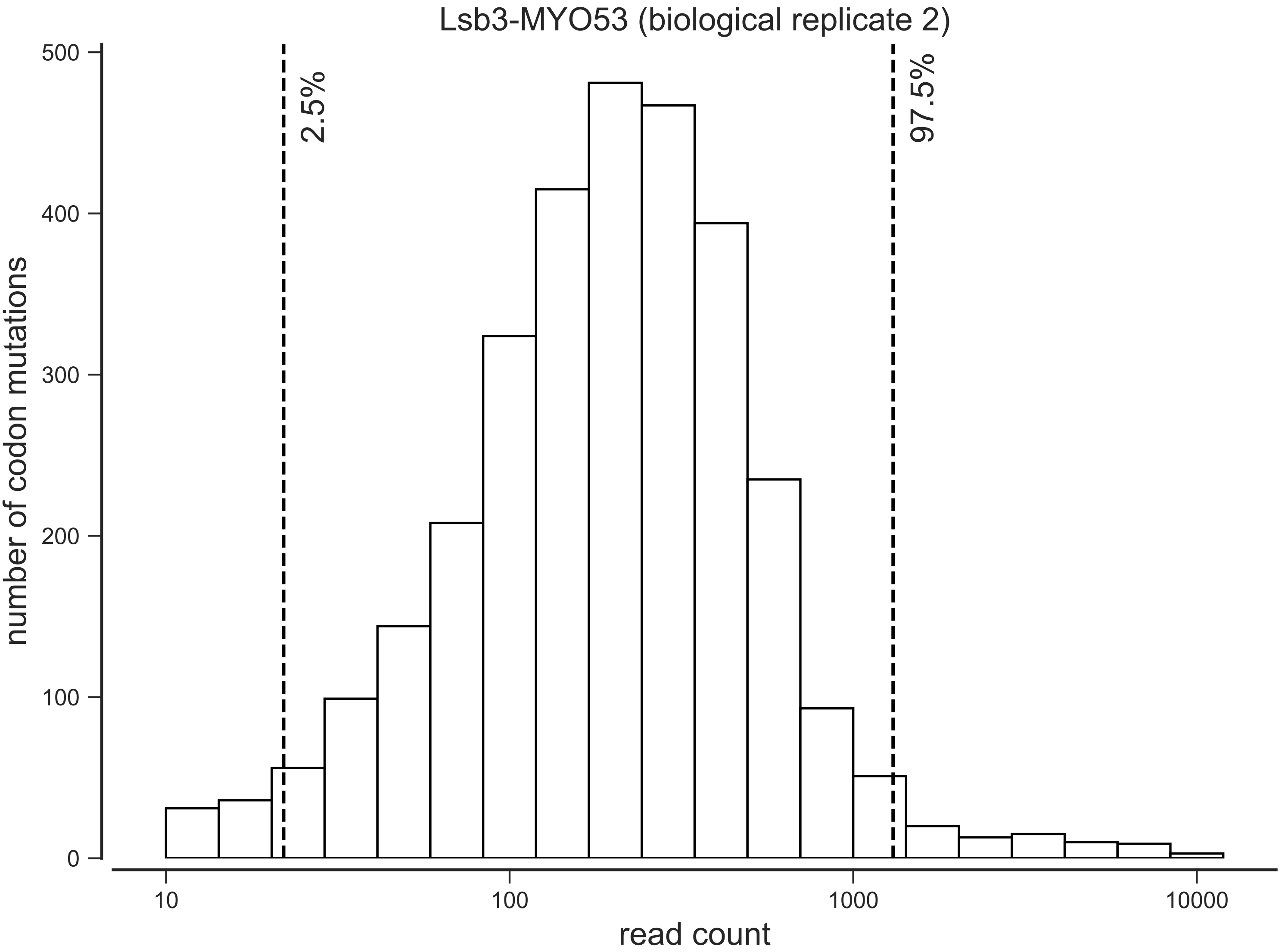

Lsb3-MYO53 (biological replicate 2)
2.5%
97.5%
number of codon mutations
0
100
200
300
400
500
10
100
1000
10000
read count

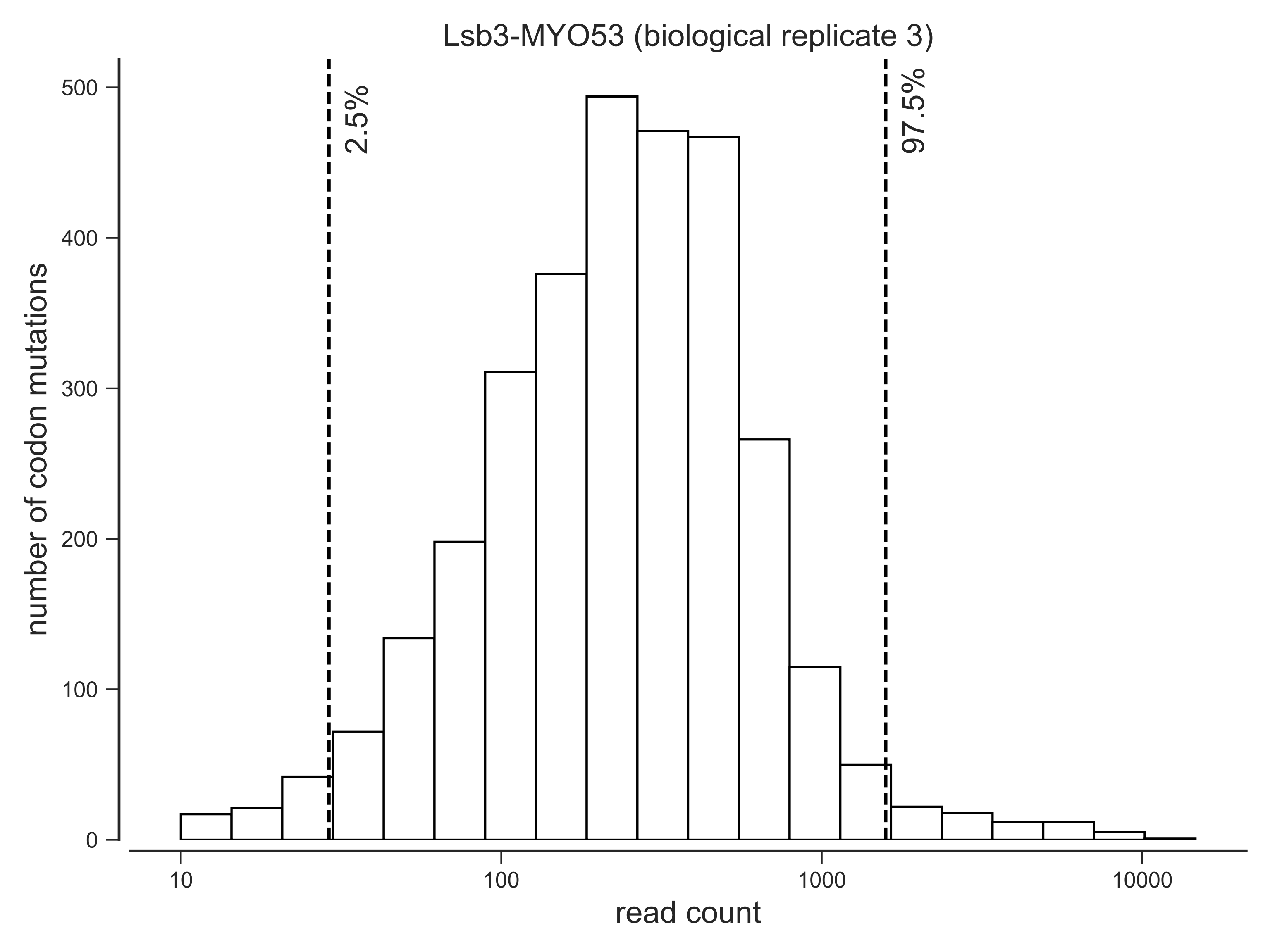
Lsb3-MYO53 (biological replicate 3)
2.5%
97.5%
number of codon mutations
0
100
200
300
400
500
10
100
1000
10000
read count

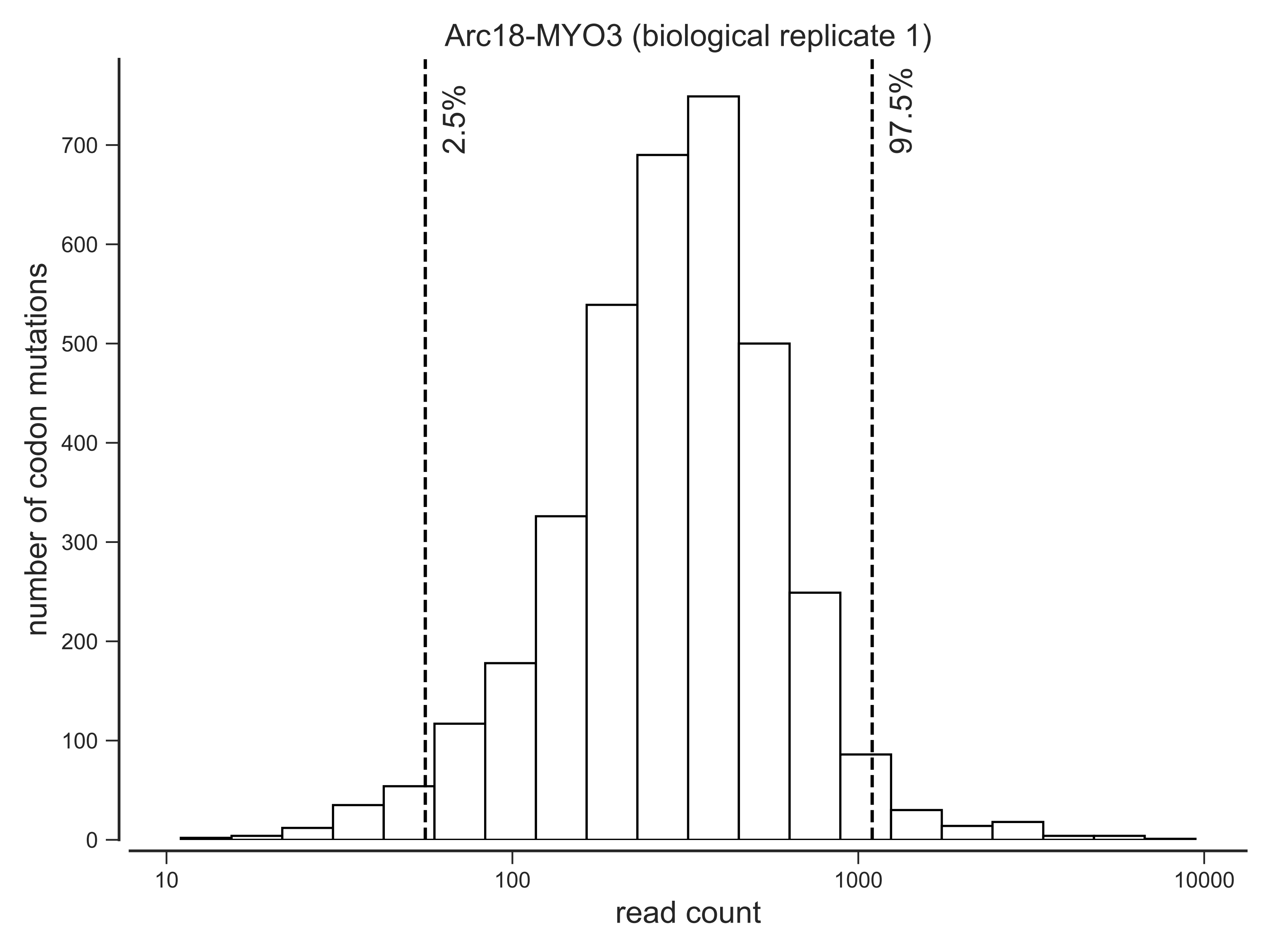

Arc18-MYO3 (biological replicate 1)
2.5%
97.5%
number of codon mutations
0
100
200
300
400
500
600
700
10
100
1000
10000
read count

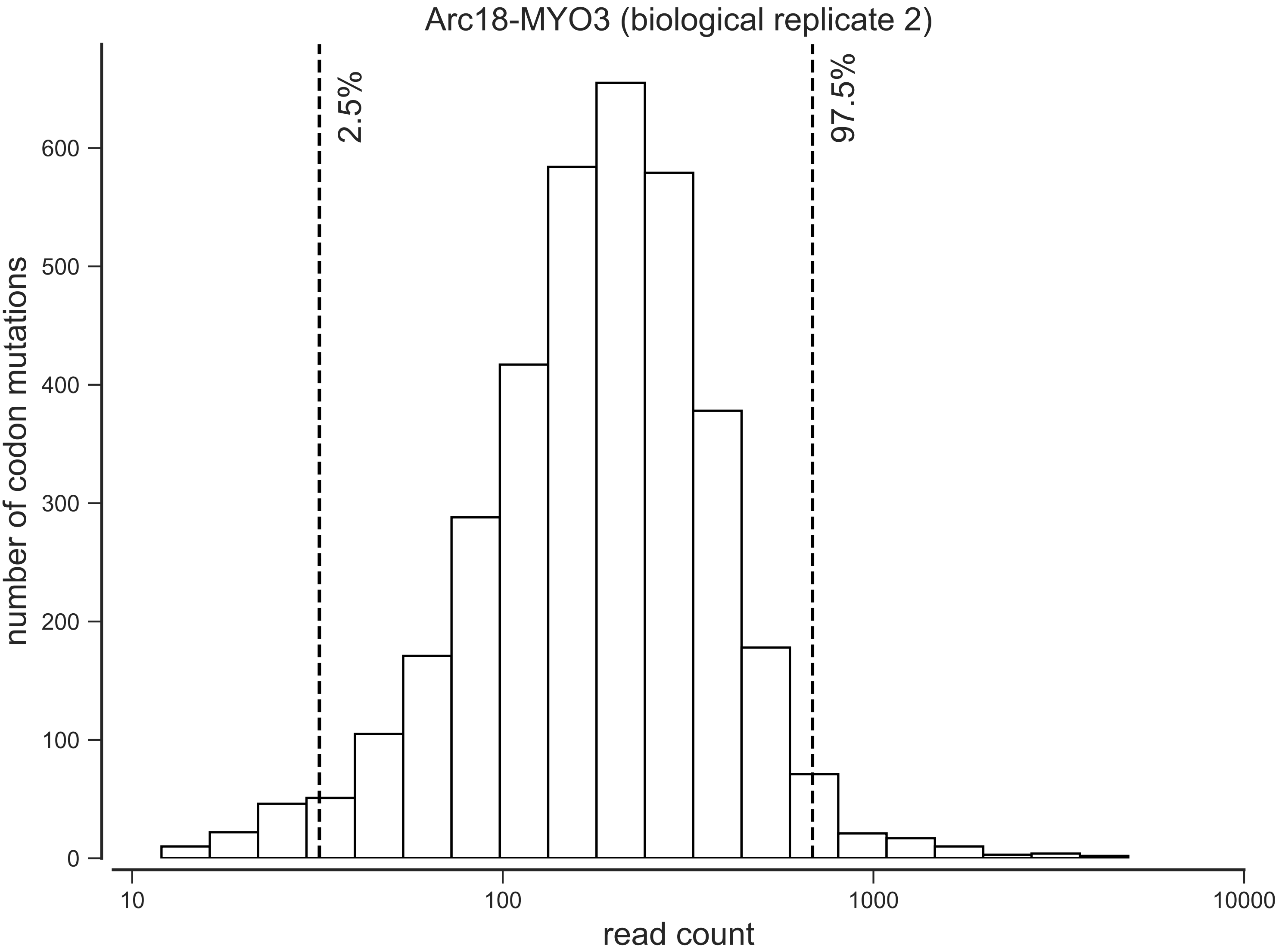

Arc18-MYO3 (biological replicate 2)
2.5%
97.5%
number of codon mutations
0
100
200
300
400
500
600
10
100
1000
10000
read count

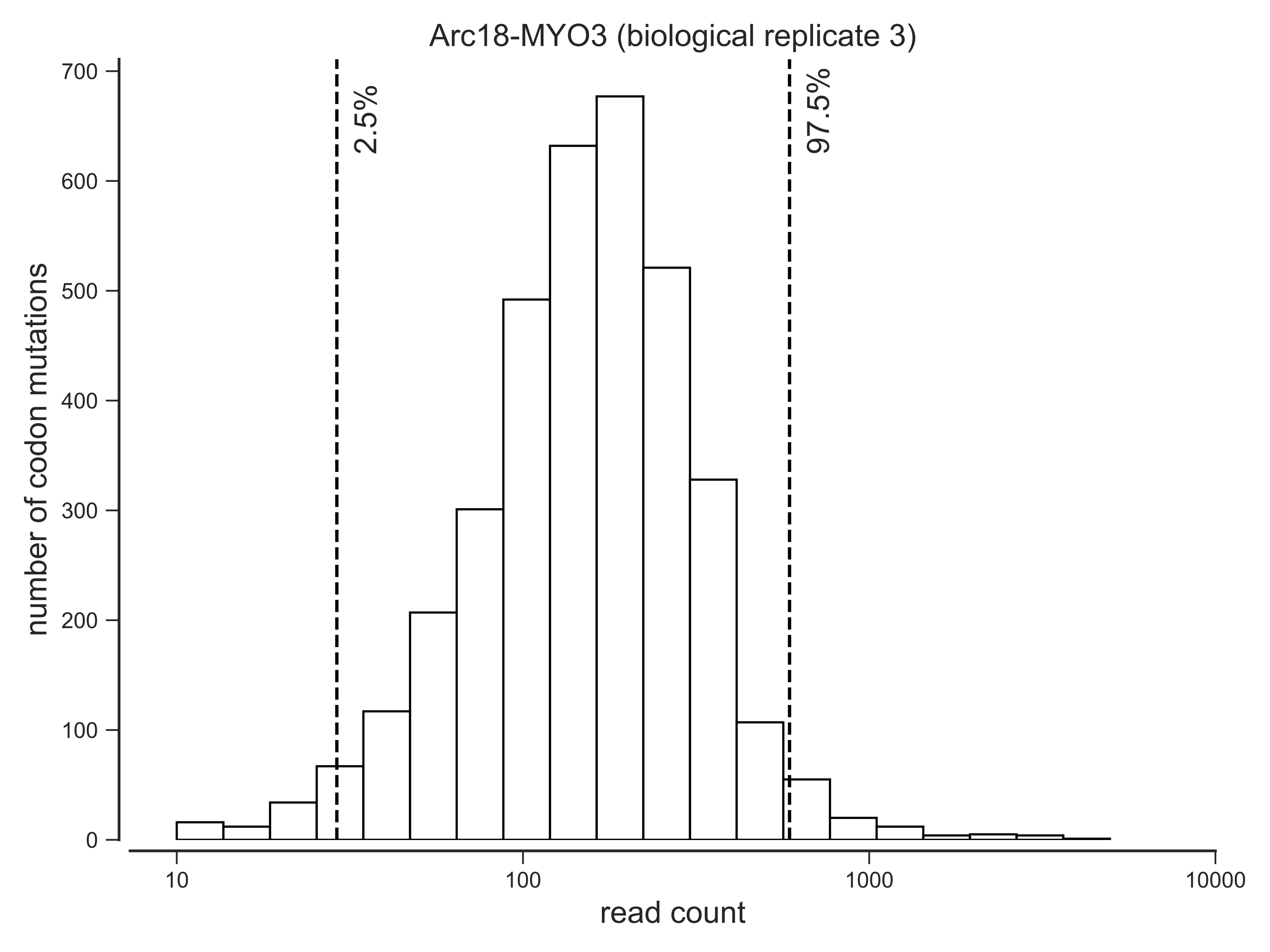

Arc18-MYO3 (biological replicate 3)
number of codon mutations
read count
2.5%
97.5%
0
100
200
300
400
500
600
700
10
100
1000
10000

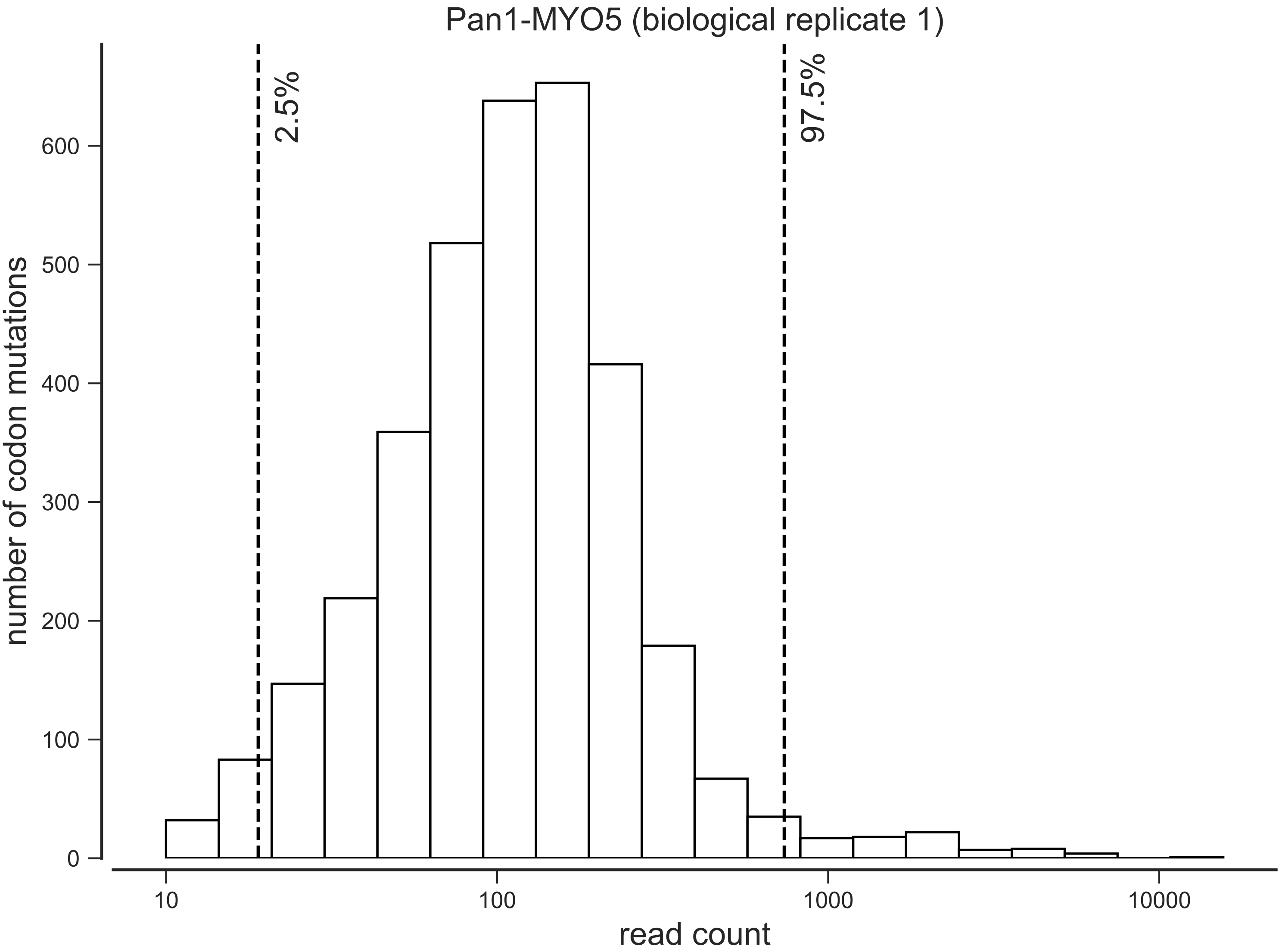

Pan1-MYO5 (biological replicate 1)
2.5%
97.5%
number of codon mutations
0
100
200
300
400
500
600
10
100
1000
10000
read count

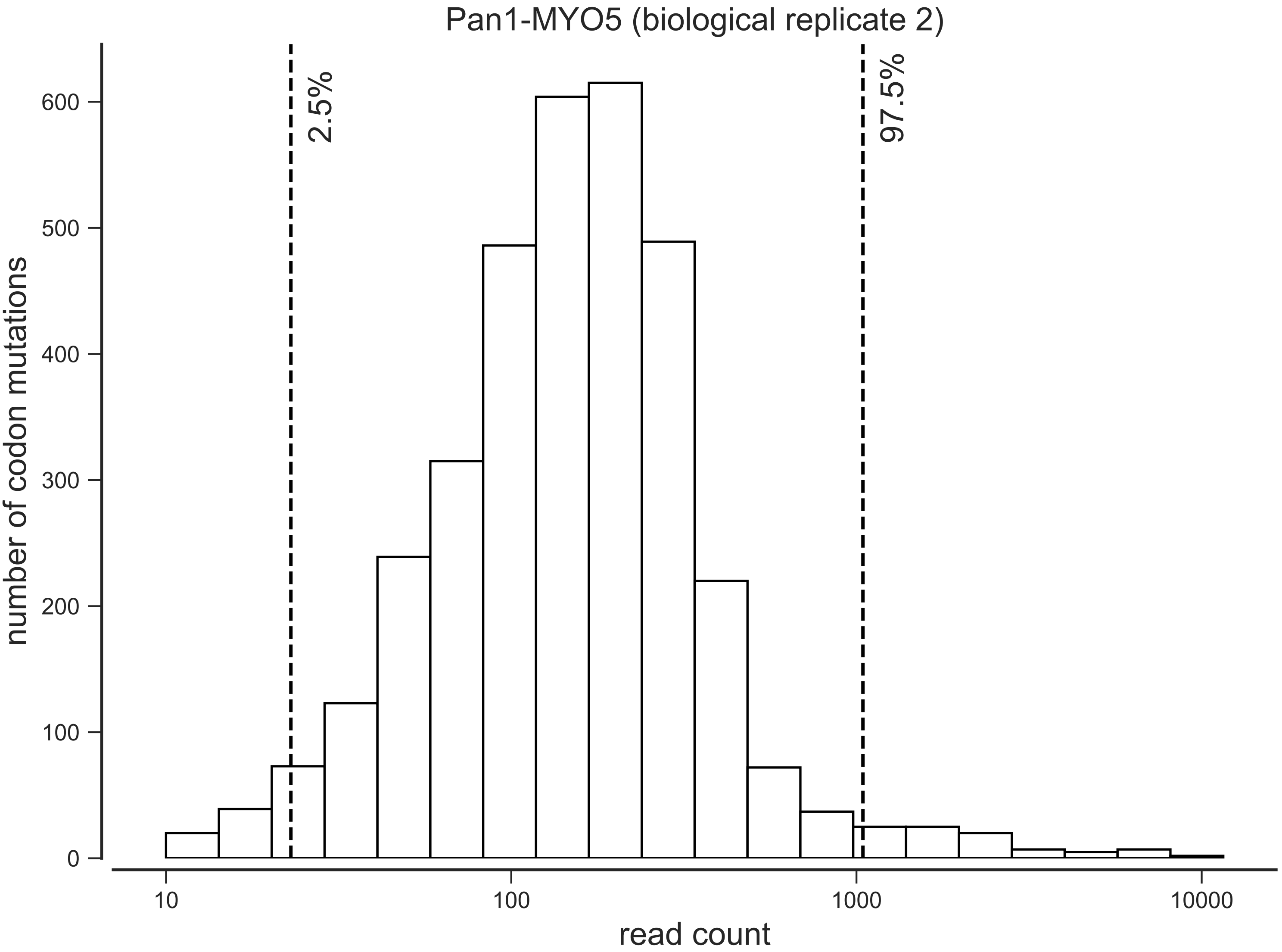

Pan1-MYO5 (biological replicate 2)
2.5%
97.5%
number of codon mutations
0
100
200
300
400
500
600
10
100
1000
10000
read count

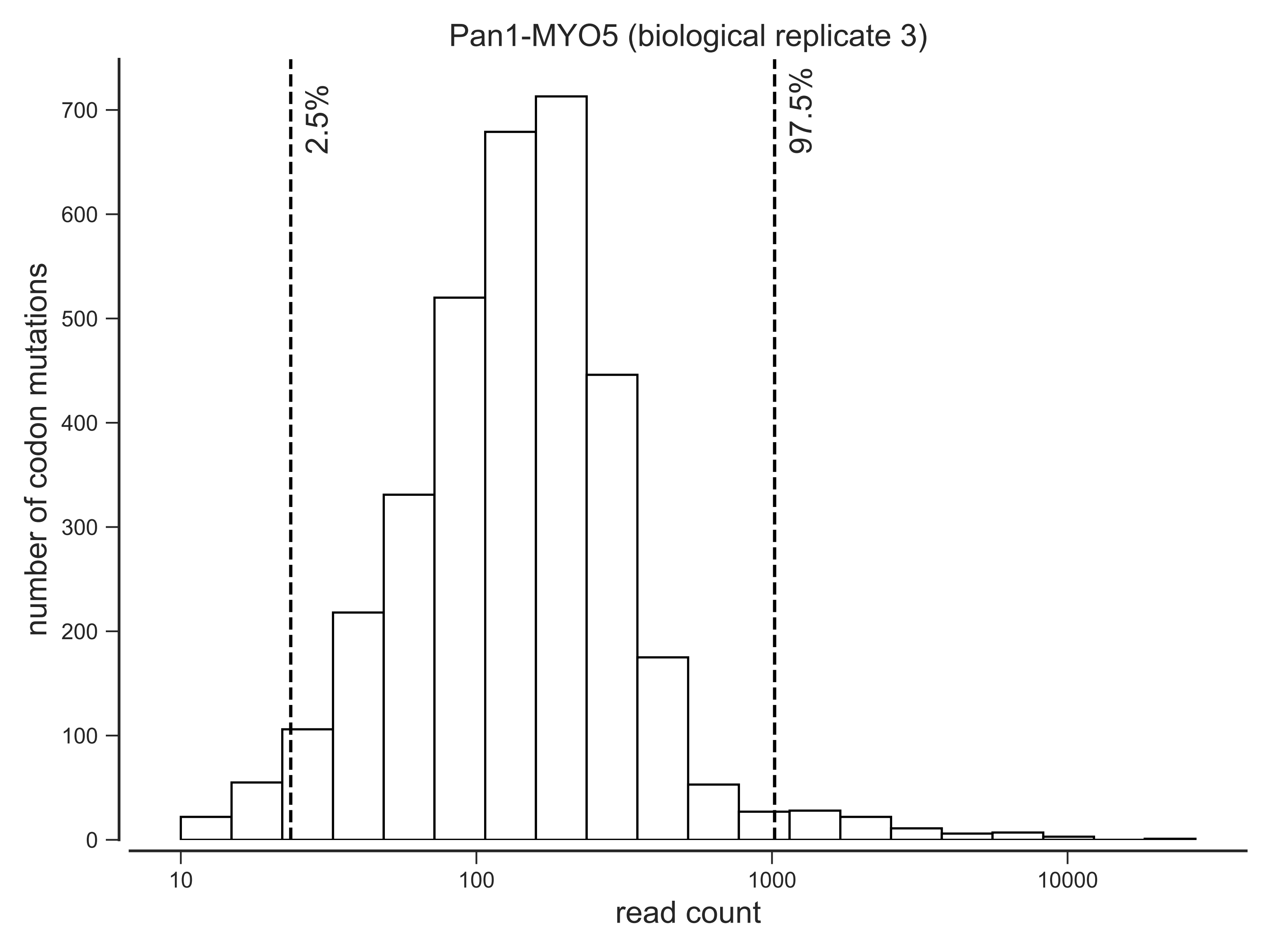

Pan1-MYO5 (biological replicate 3)
2.5%
97.5%
number of codon mutations
0
100
200
300
400
500
600
700
10
100
1000
10000
read count

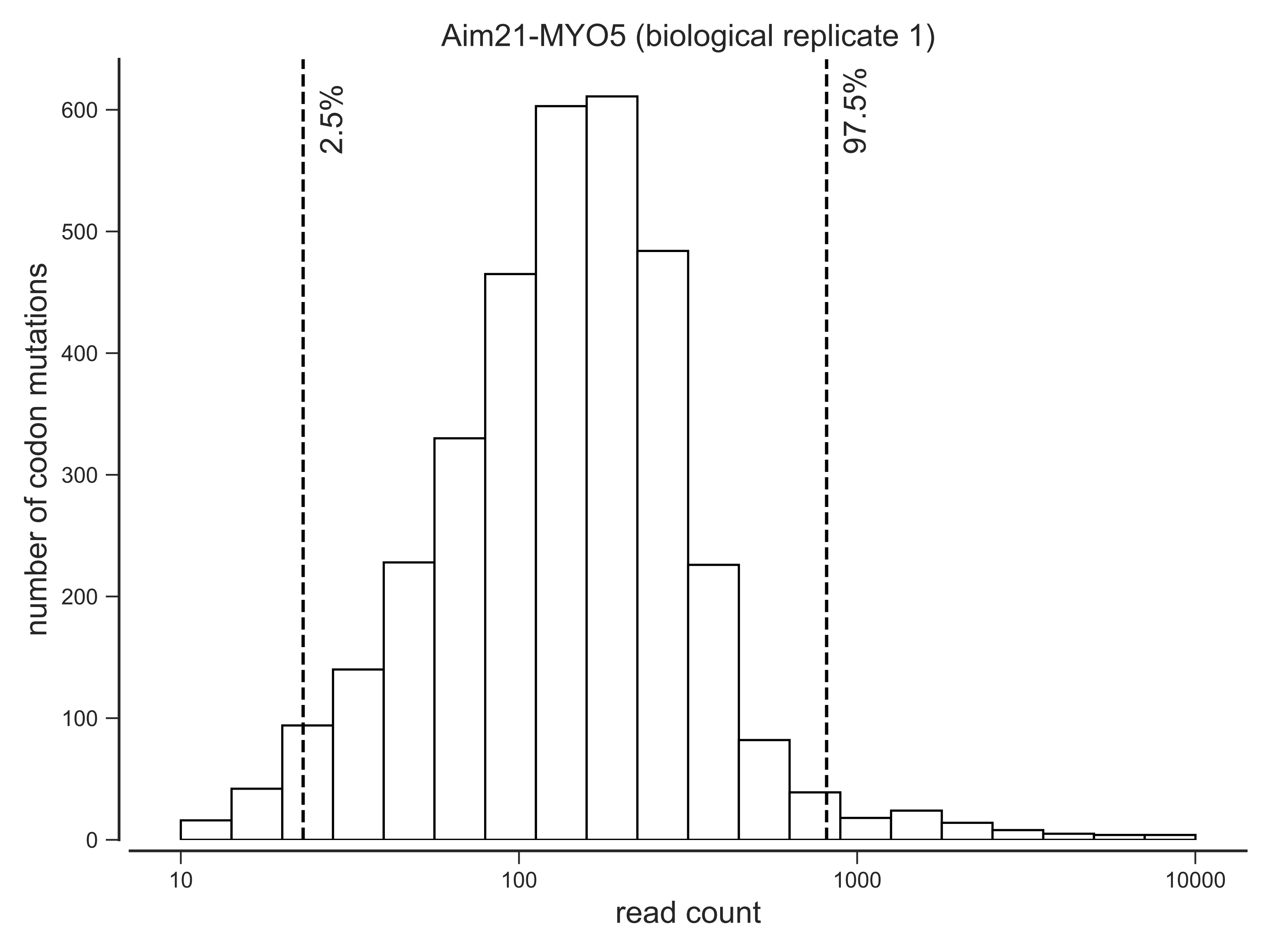

Aim21-MYO5 (biological replicate 1)
2.5%
97.5%
number of codon mutations
0
100
200
300
400
500
600
10
100
1000
10000
read count

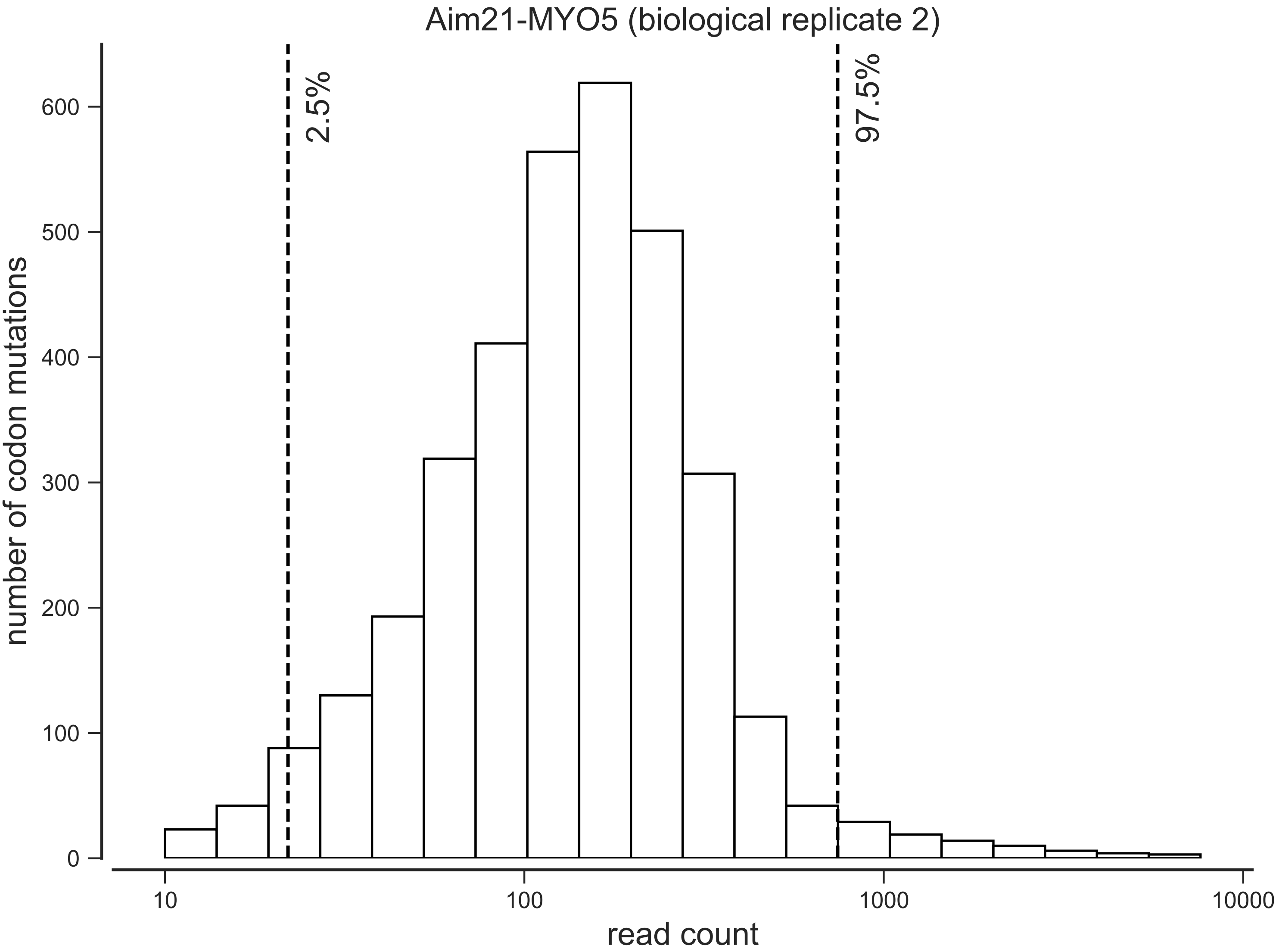
Aim21-MYO5 (biological replicate 2)
2.5%
97.5%
number of codon mutations
0
100
200
300
400
500
600
10
100
1000
10000
read count

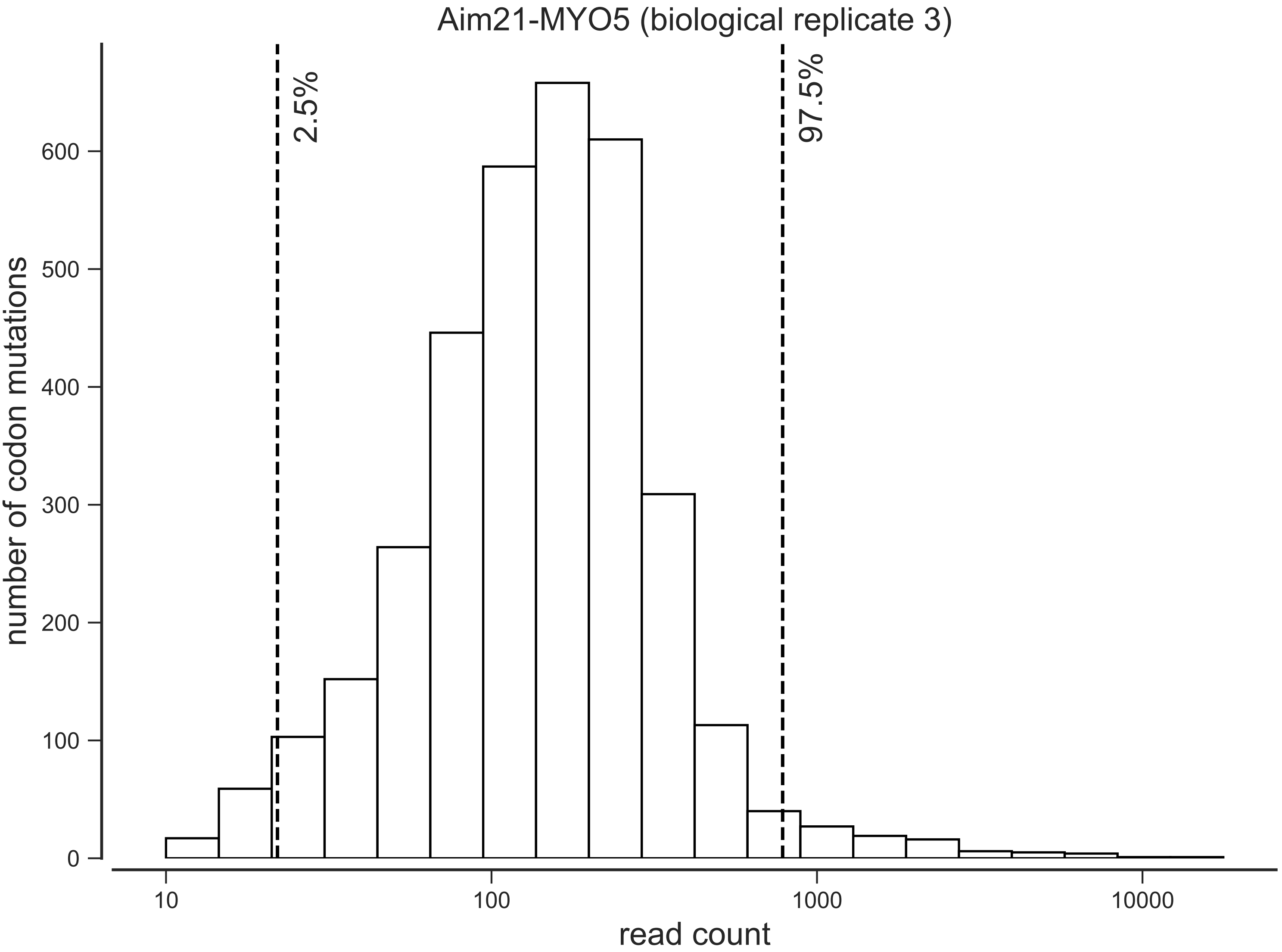

Aim21-MYO5 (biological replicate 3)
2.5%
97.5%
number of codon mutations
0
100
200
300
400
500
600
10
100
1000
10000
read count

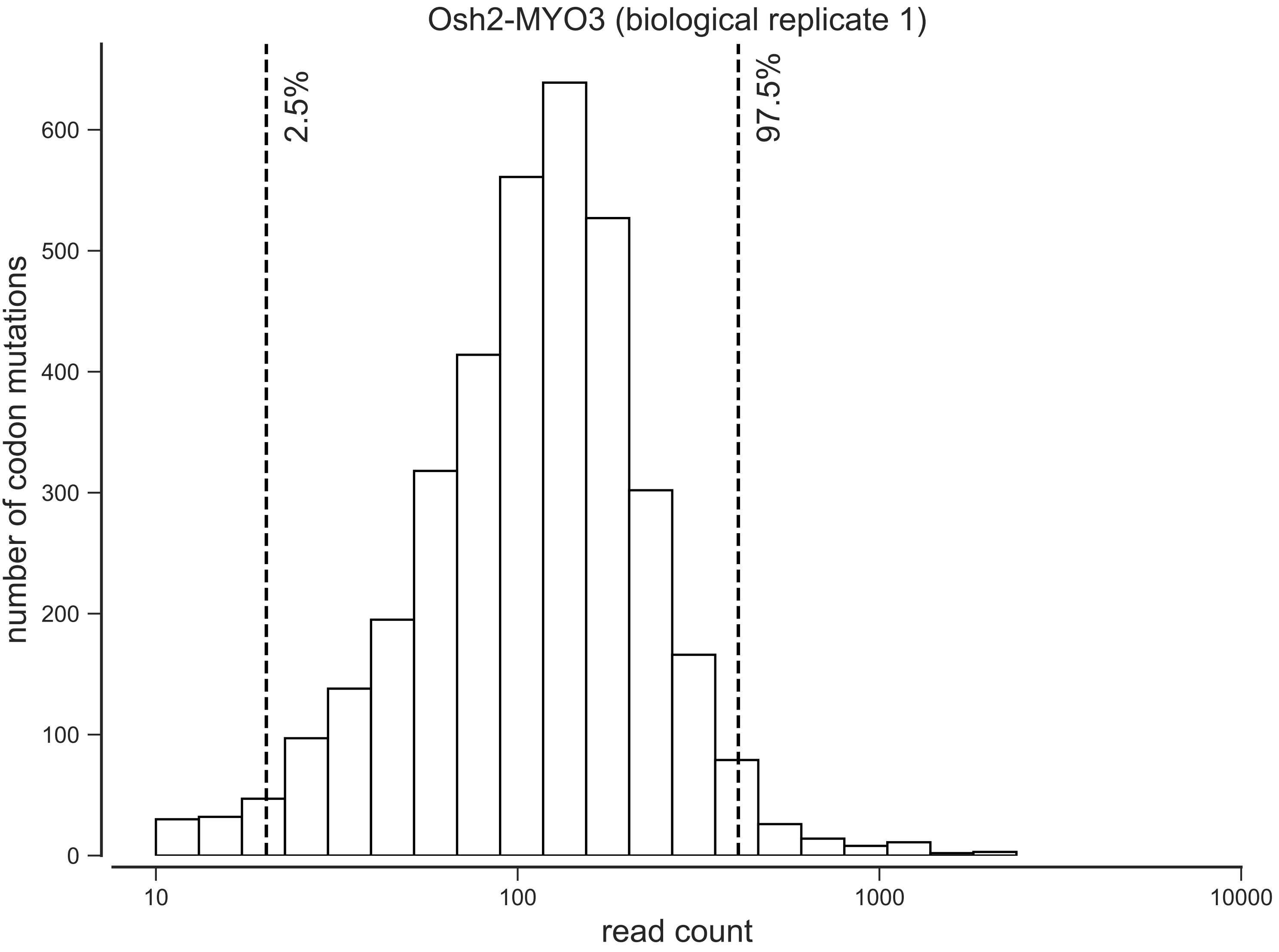

Osh2-MYO3 (biological replicate 1)
2.5%
97.5%
number of codon mutations
0
100
200
300
400
500
600
10
100
1000
10000
read count

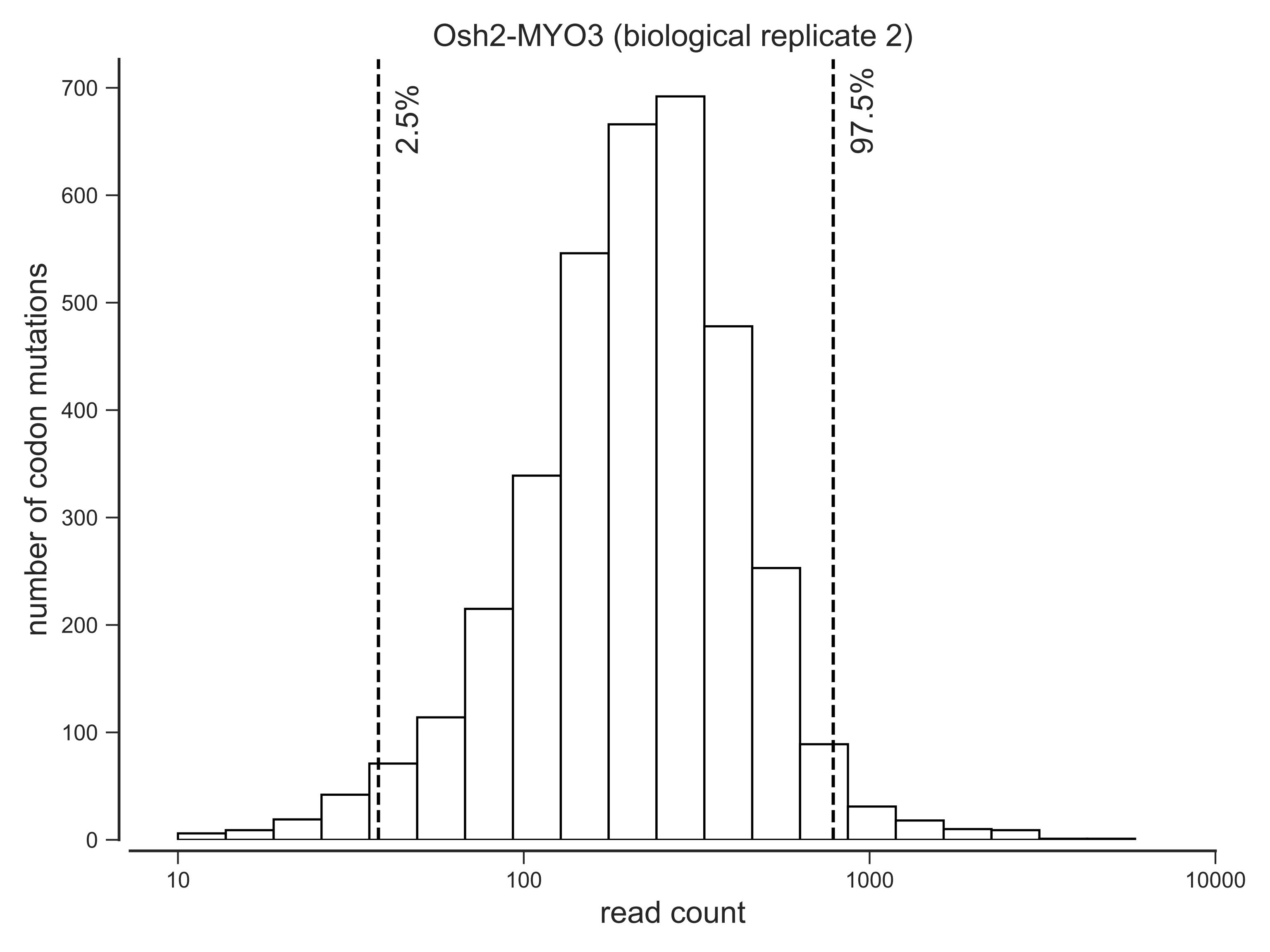
Osh2-MYO3 (biological replicate 2)
2.5%
97.5%
number of codon mutations
0
100
200
300
400
500
600
700
10
100
1000
10000
read count

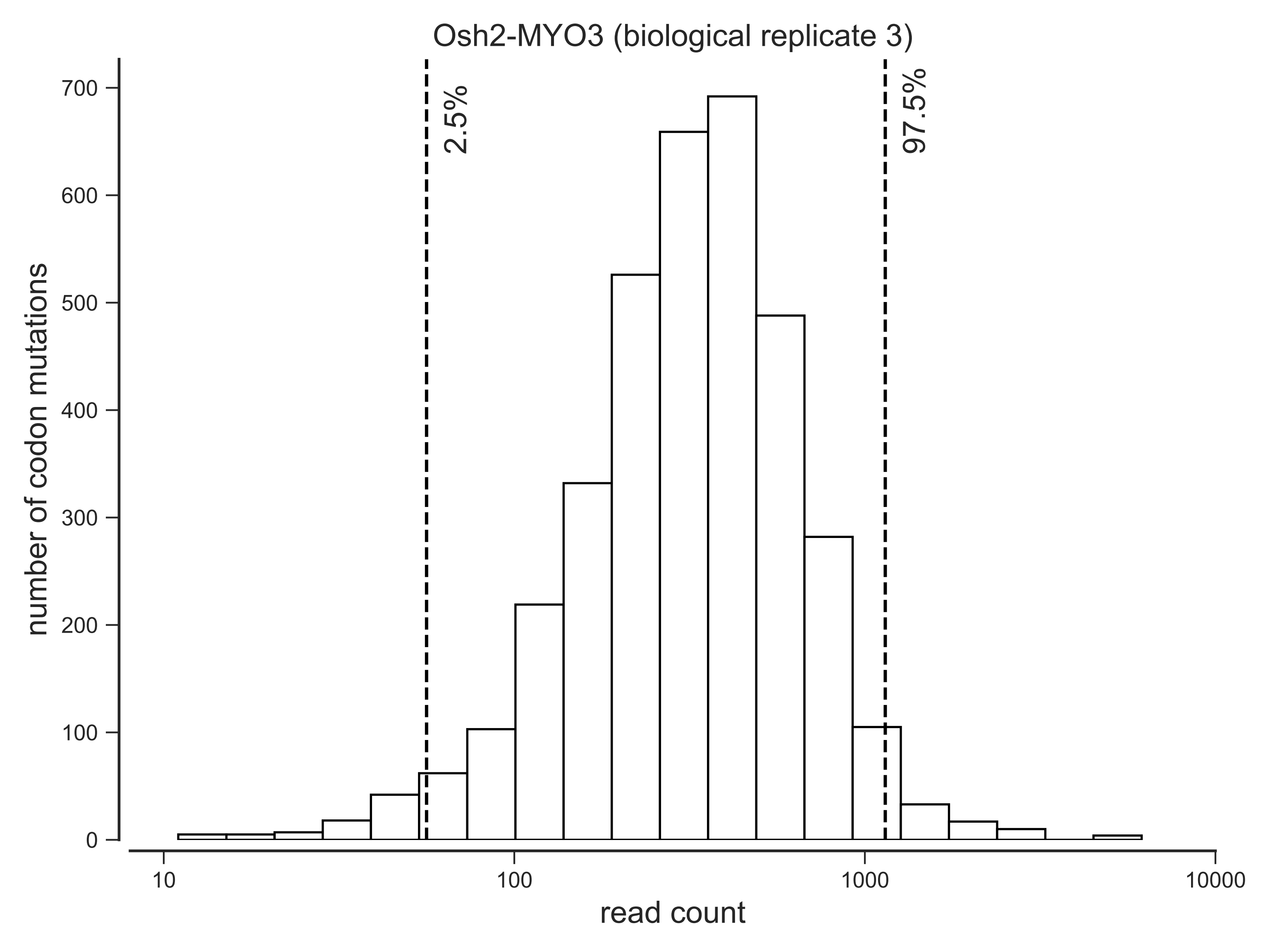

Osh2-MYO3 (biological replicate 3)
number of codon mutations
read count
2.5%
97.5%
0
100
200
300
400
500
600
700
10
100
1000
10000

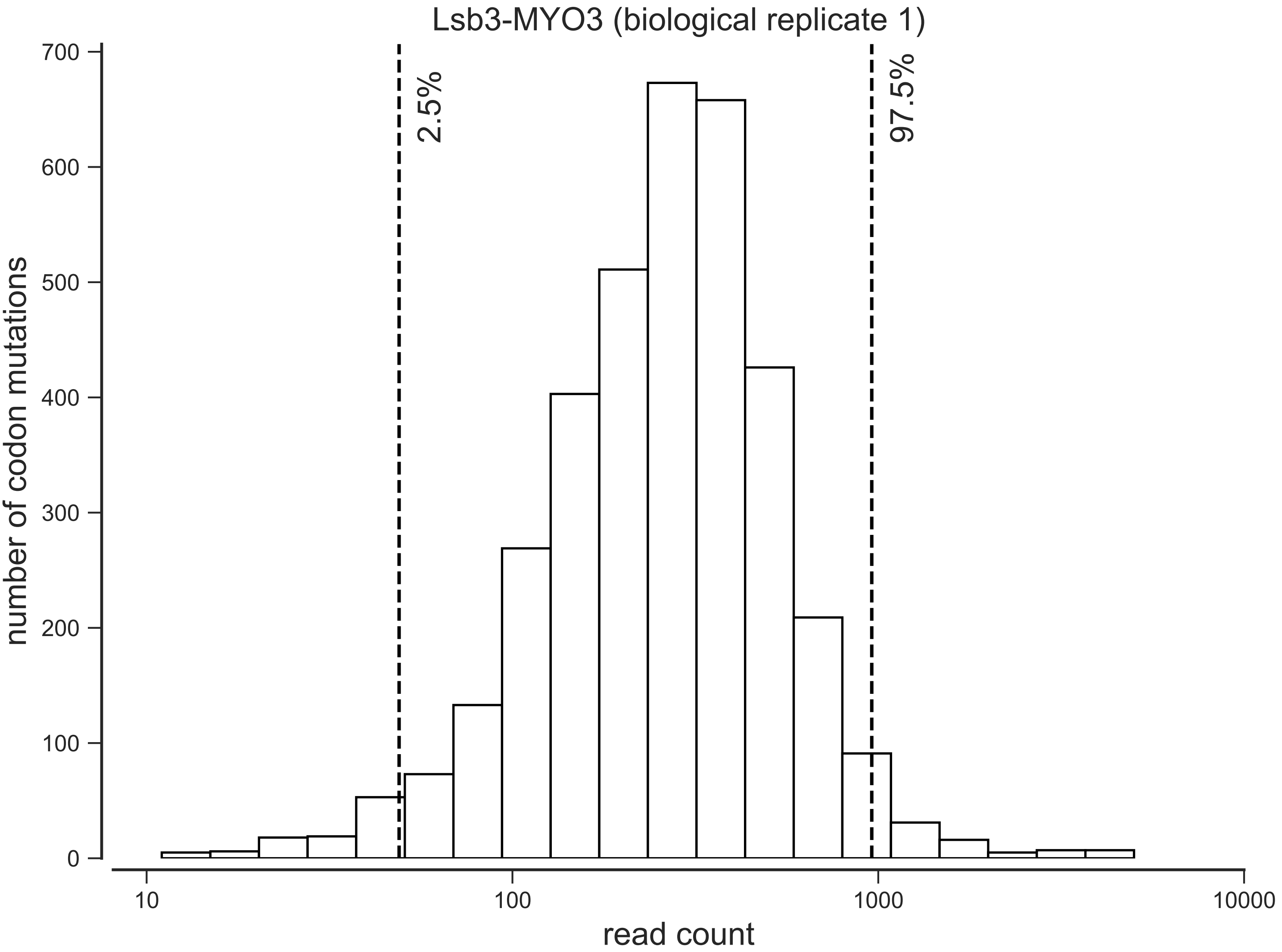

Lsb3-MYO3 (biological replicate 1)
2.5%
97.5%
number of codon mutations
read count
0
100
200
300
400
500
600
700
10
100
1000
10000

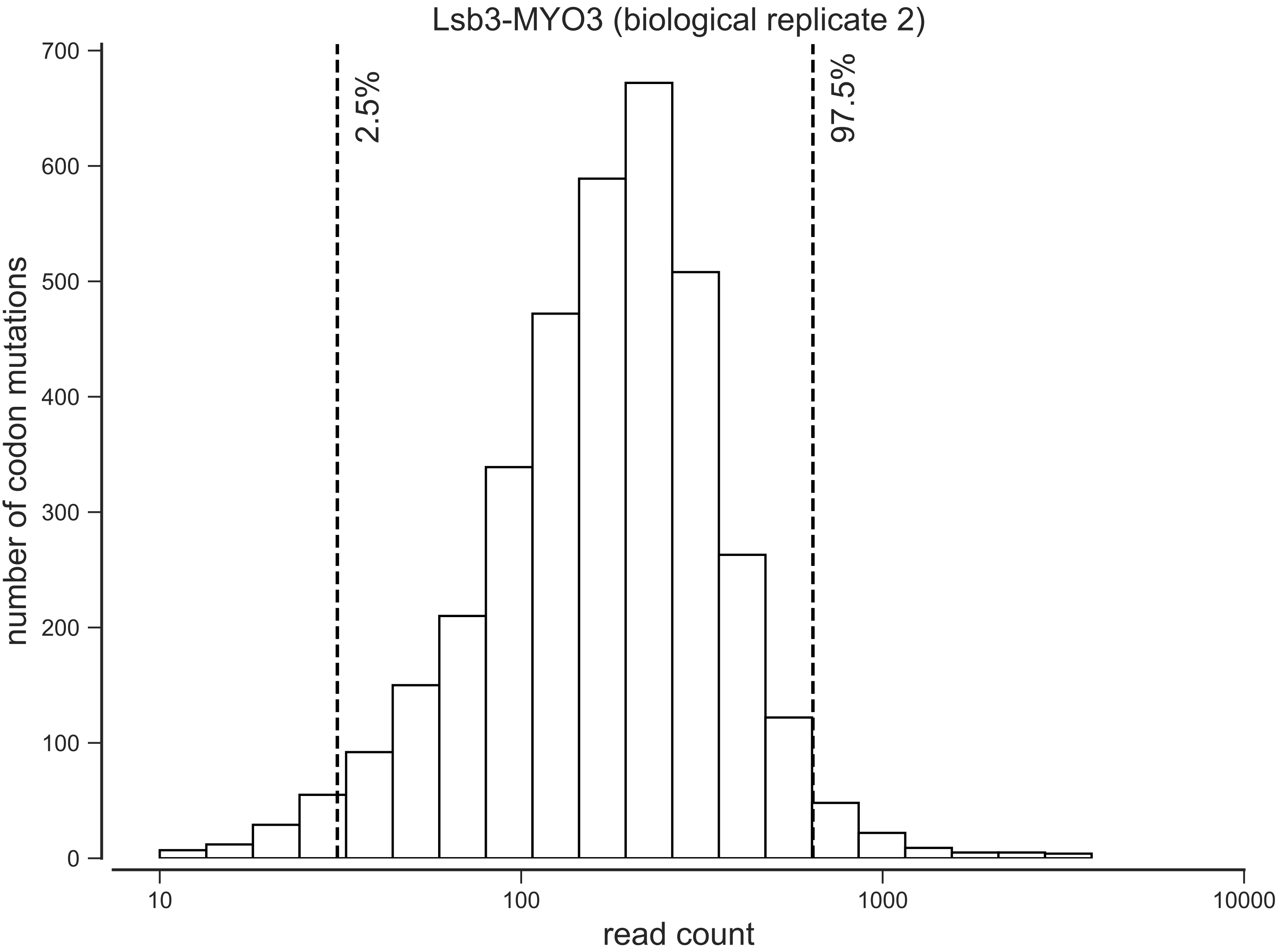

Lsb3-MYO3 (biological replicate 2)
2.5%
97.5%
number of codon mutations
0
100
200
300
400
500
600
700
10
100
1000
10000
read count

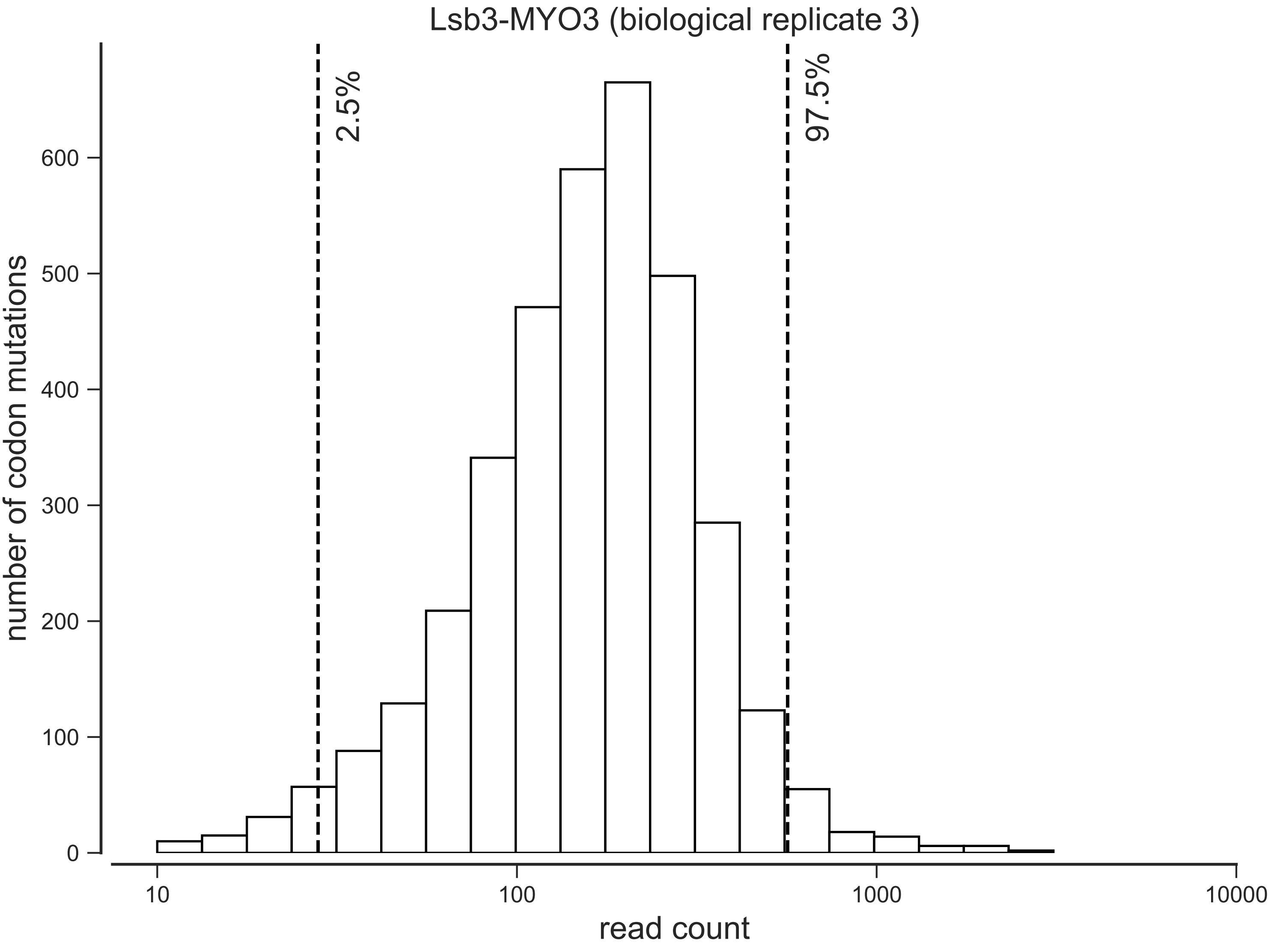
Lsb3-MYO3 (biological replicate 3)
2.5%
97.5%
number of codon mutations
0
100
200
300
400
500
600
10
100
1000
10000
read count

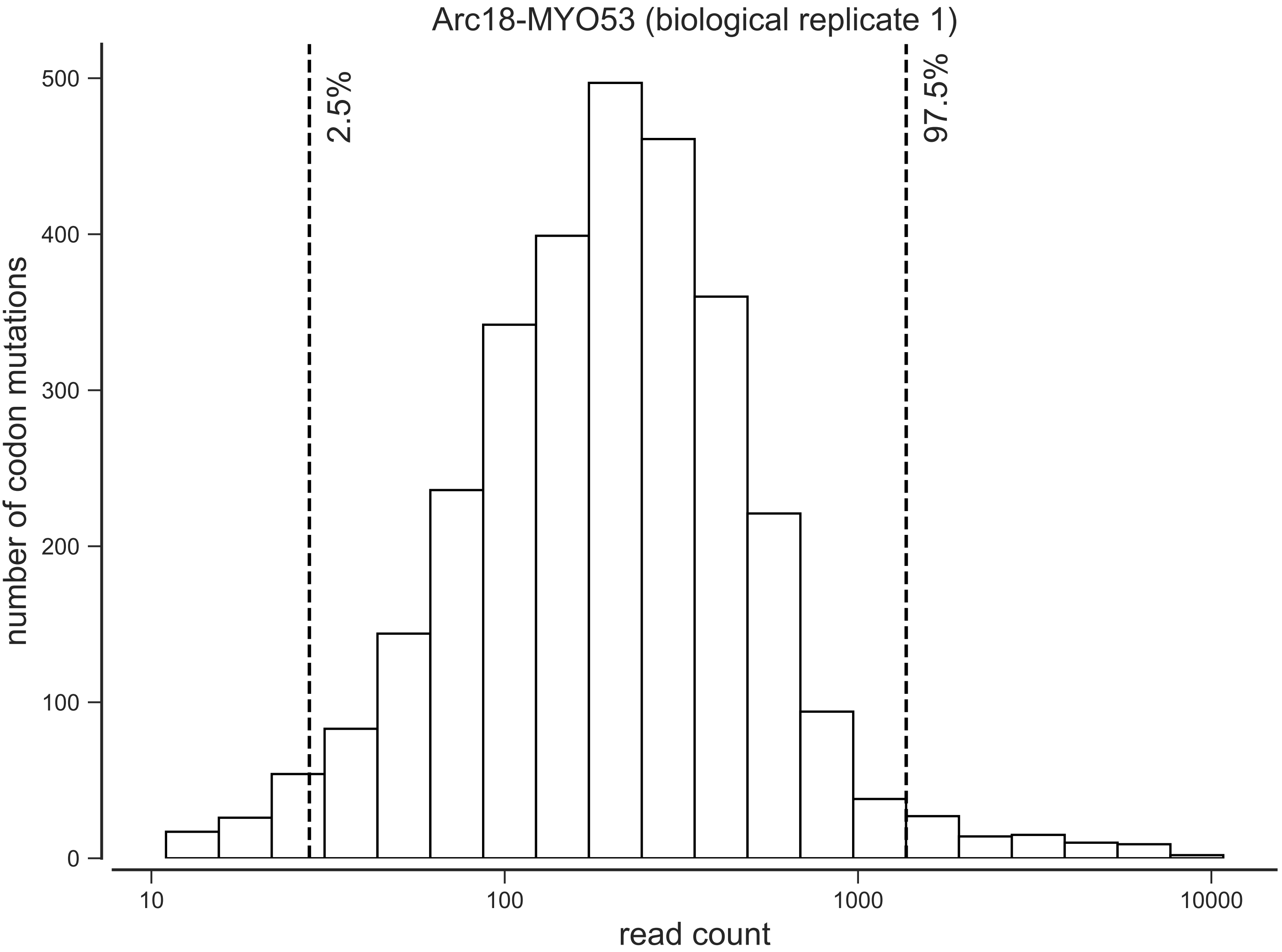
Arc18-MYO53 (biological replicate 1)
2.5%
97.5%
number of codon mutations
0
100
200
300
400
500
10
100
1000
10000
read count

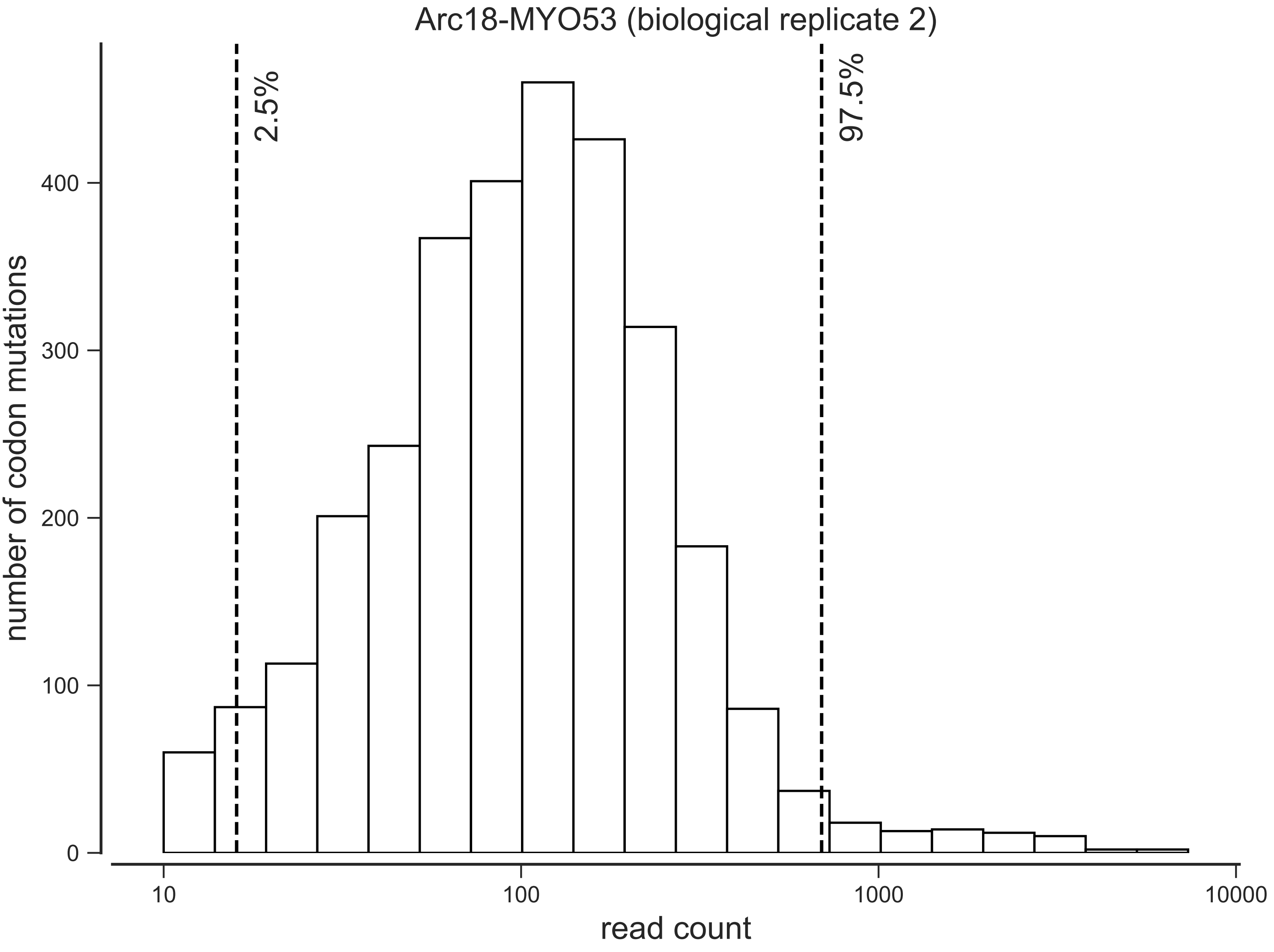

Arc18-MYO53 (biological replicate 2)
2.5%
97.5%
number of codon mutations
0
100
200
300
400
10
100
1000
10000
read count

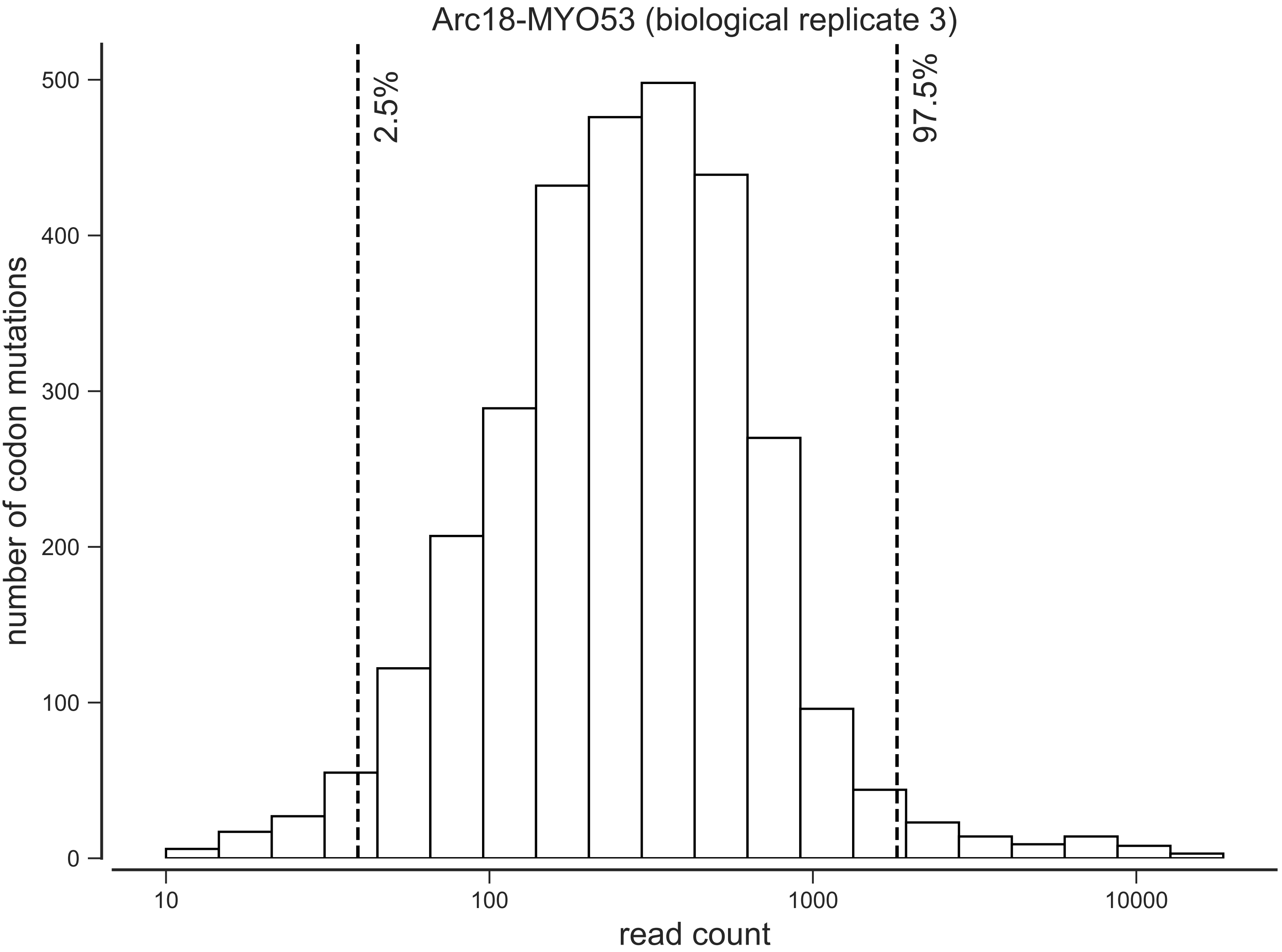

Arc18-MYO53 (biological replicate 3)
number of codon mutations
read count
2.5%
97.5%
0
100
200
300
400
500
10
100
1000
10000

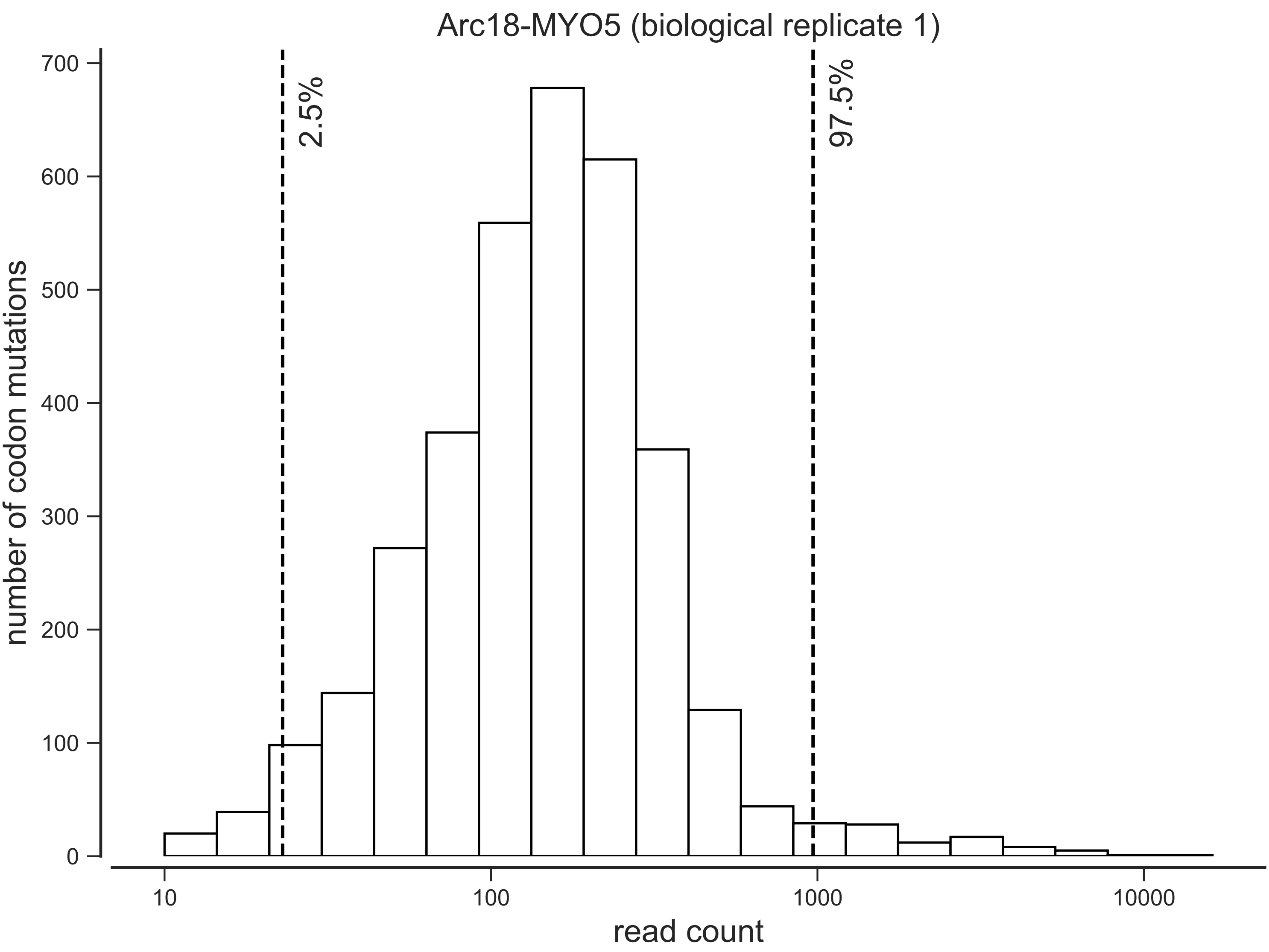

Arc18-MYO5 (biological replicate 1)
2.5%
97.5%
number of codon mutations
read count
0
100
200
300
400
500
600
700
10
100
1000
10000

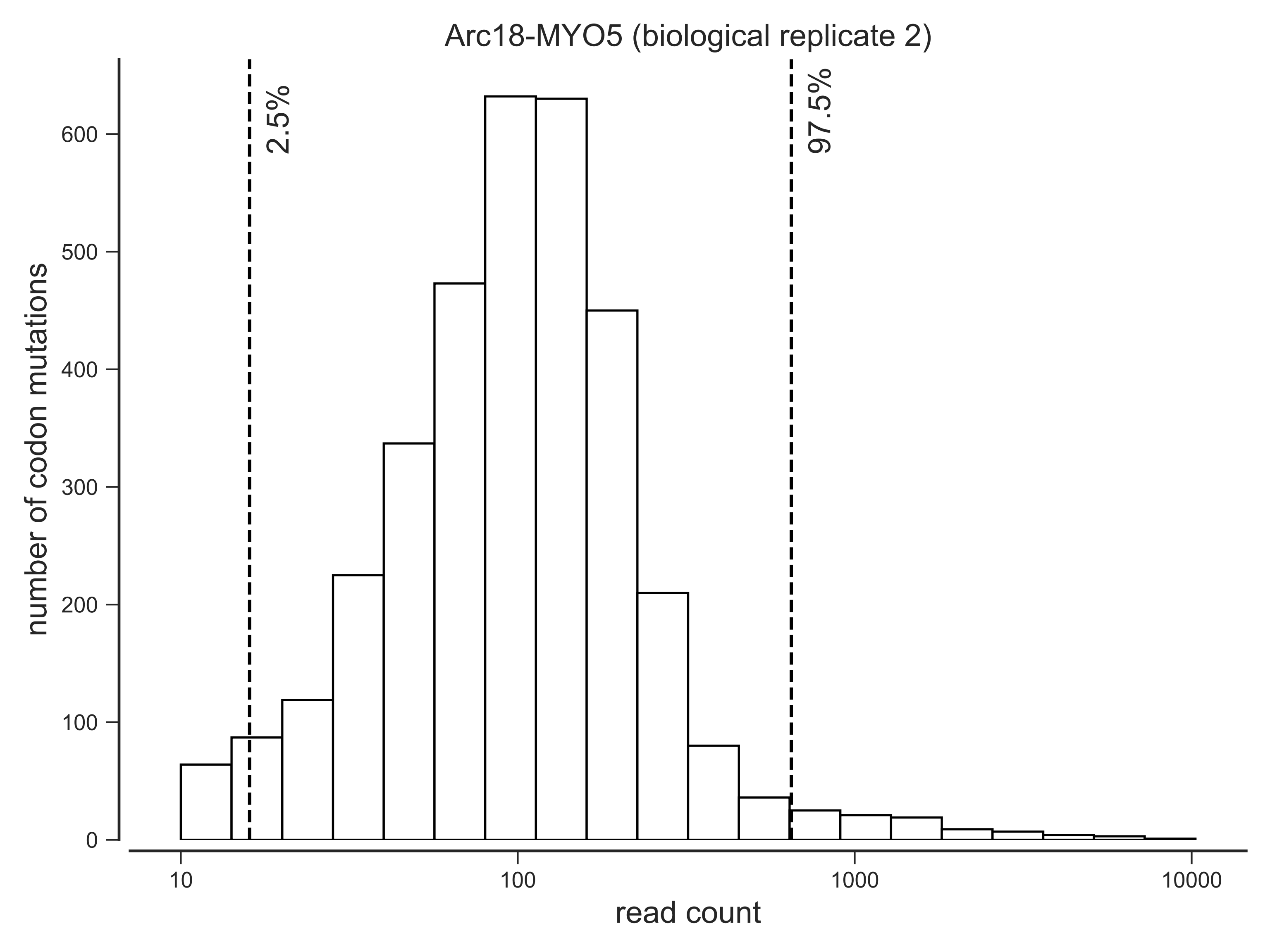

Arc18-MYO5 (biological replicate 2)
2.5%
97.5%
number of codon mutations
0
100
200
300
400
500
600
10
100
1000
10000
read count

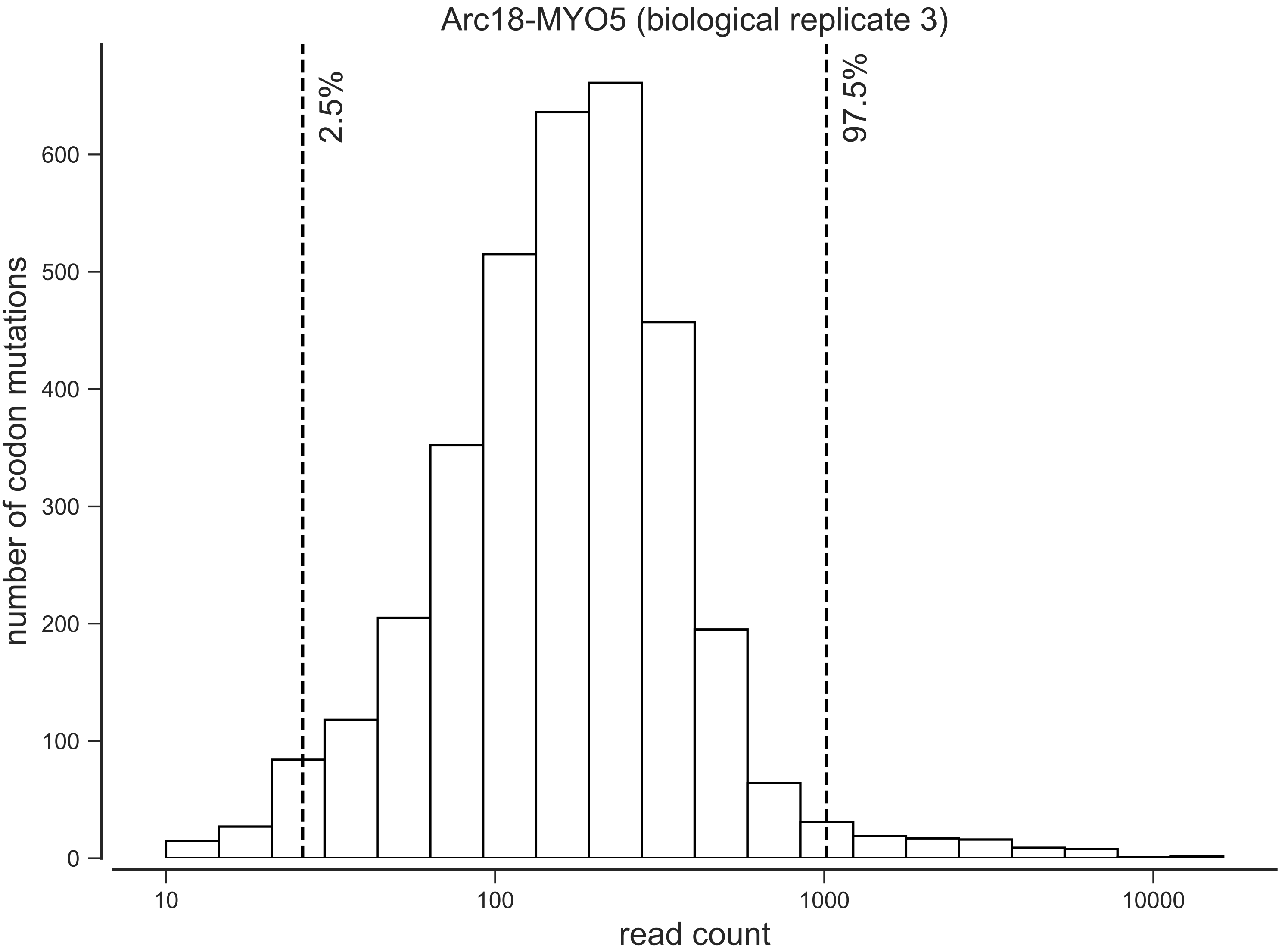

Arc18-MYO5 (biological replicate 3)
2.5%
97.5%
number of codon mutations
0
100
200
300
400
500
600
10
100
1000
10000
read count

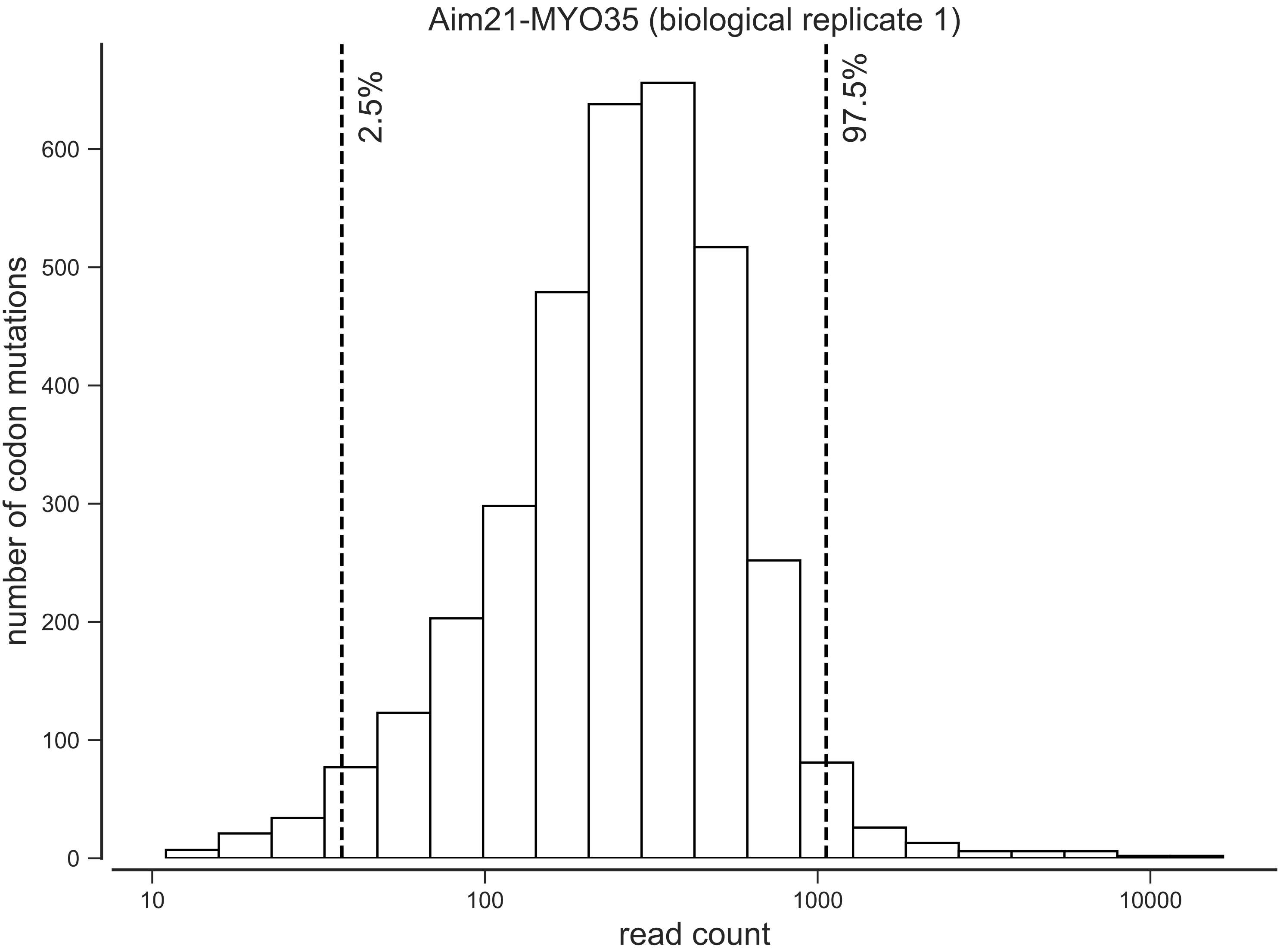

Aim21-MYO35 (biological replicate 1)
2.5%
97.5%
number of codon mutations
0
100
200
300
400
500
600
10
100
1000
10000
read count

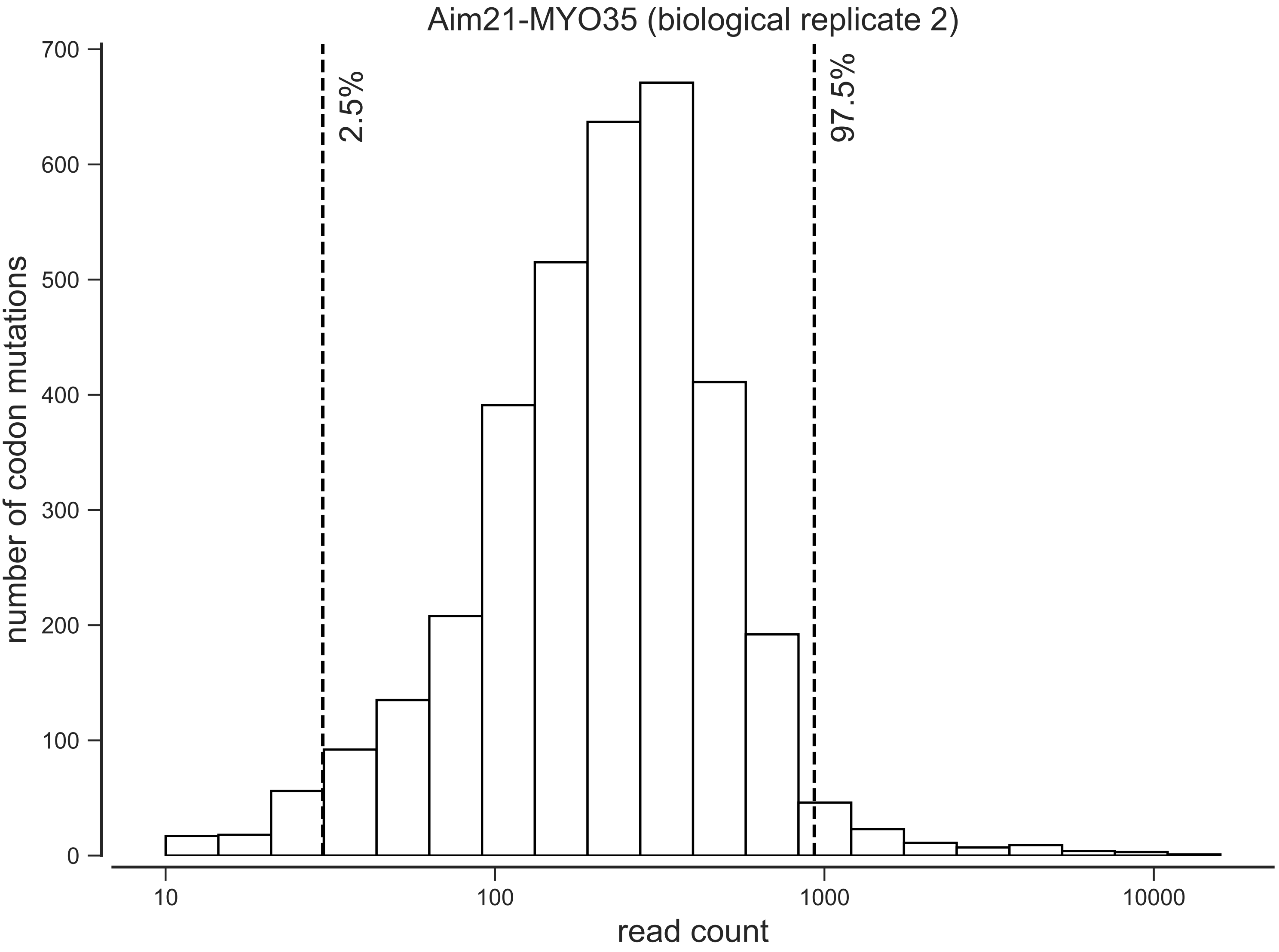

Aim21-MYO35 (biological replicate 2)
number of codon mutations
read count
2.5%
97.5%
0
100
200
300
400
500
600
700
10
100
1000
10000

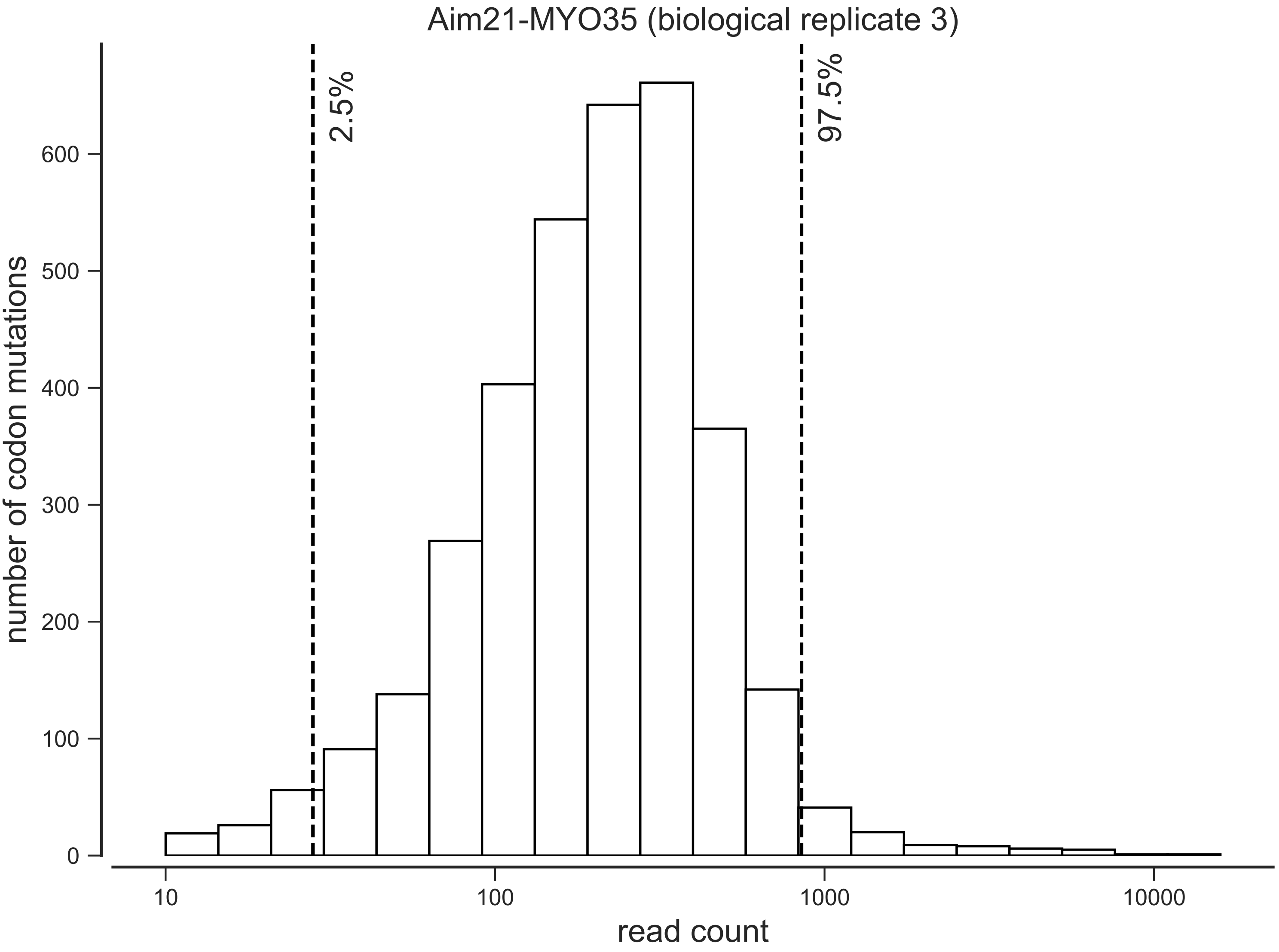

Aim21-MYO35 (biological replicate 3)
2.5%
97.5%
number of codon mutations
0
100
200
300
400
500
600
10
100
1000
10000
read count

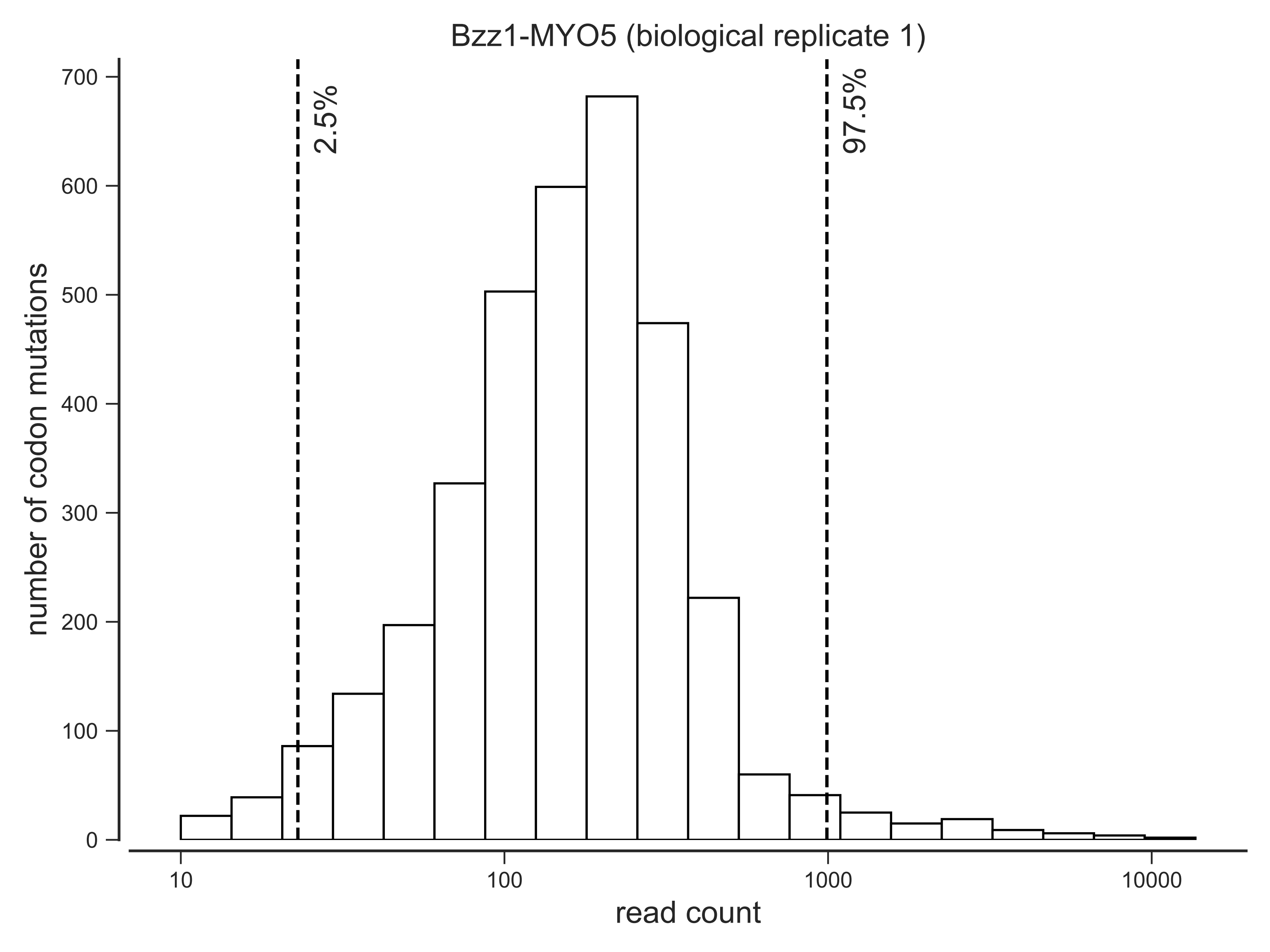

Bzz1-MYO5 (biological replicate 1)
2.5%
97.5%
number of codon mutations
700
600
500
400
300
200
100
0
10
100
1000
10000
read count

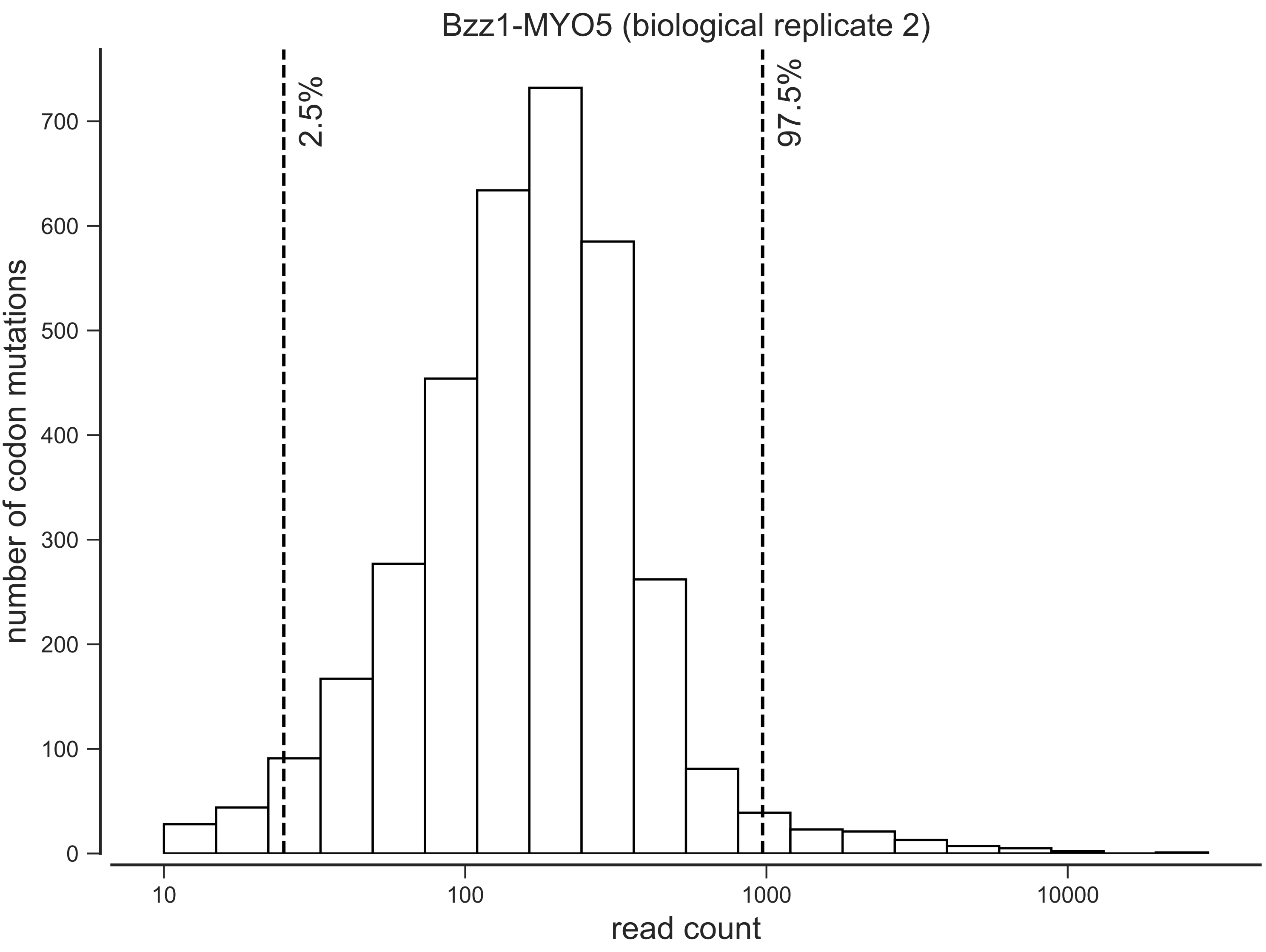

Bzz1-MYO5 (biological replicate 2)
2.5%
97.5%
number of codon mutations
0
100
200
300
400
500
600
700
10
100
1000
10000
read count

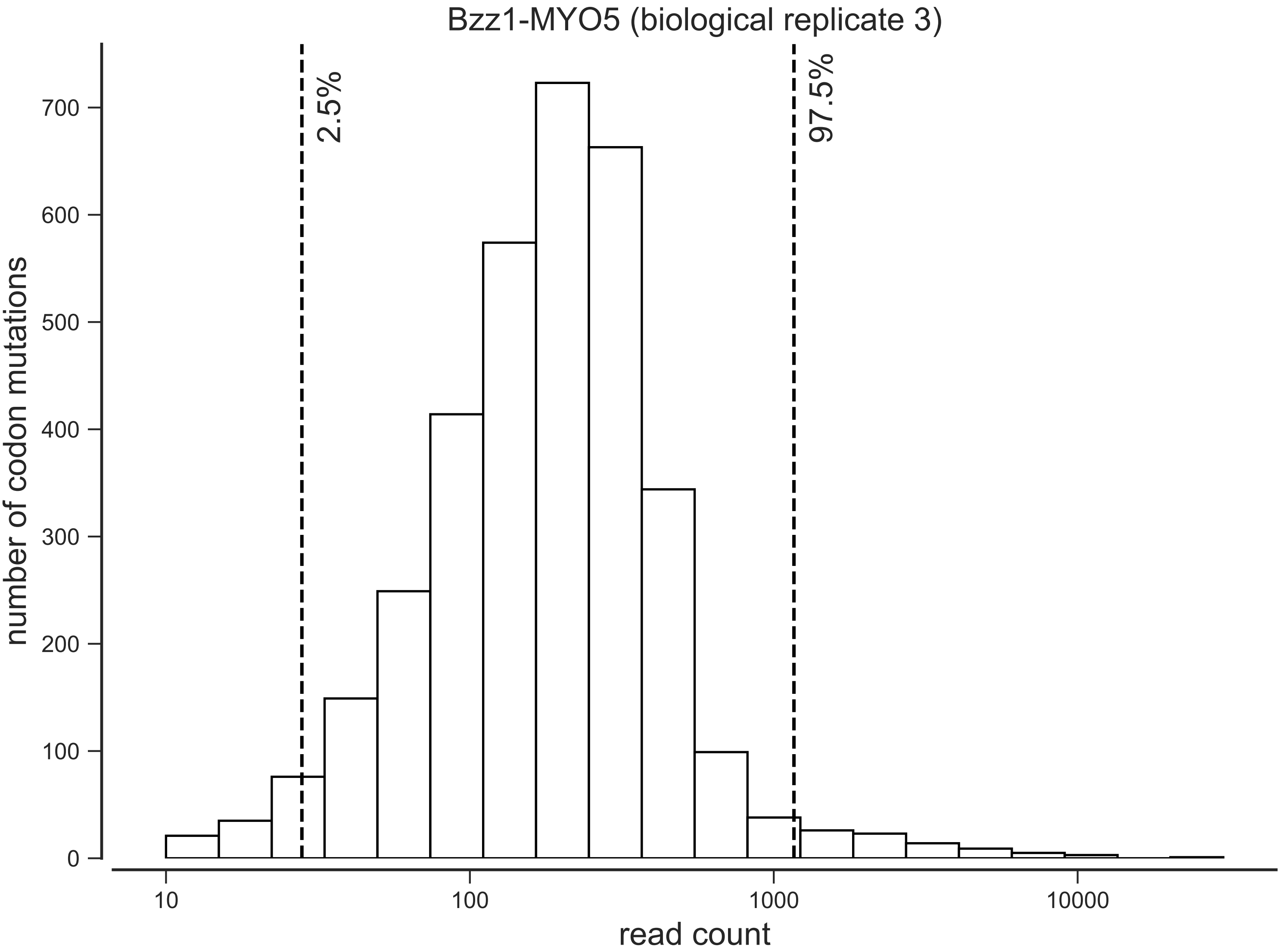

Bzz1-MYO5 (biological replicate 3)
2.5%
97.5%
number of codon mutations
0
100
200
300
400
500
600
700
10
100
1000
10000
read count

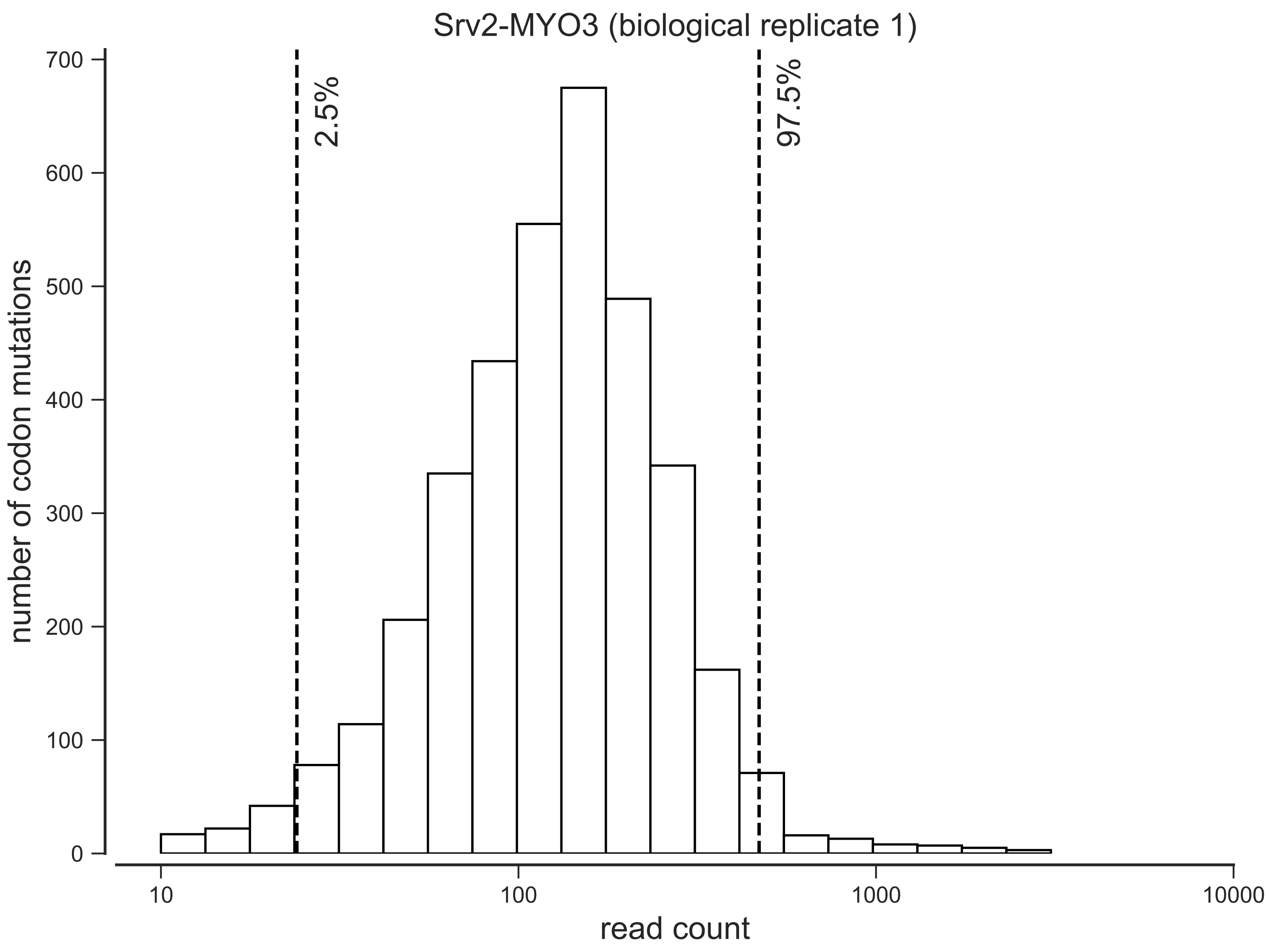
Srv2-MYO3 (biological replicate 1)
2.5%
97.5%
number of codon mutations
0
100
200
300
400
500
600
700
10
100
1000
10000
read count

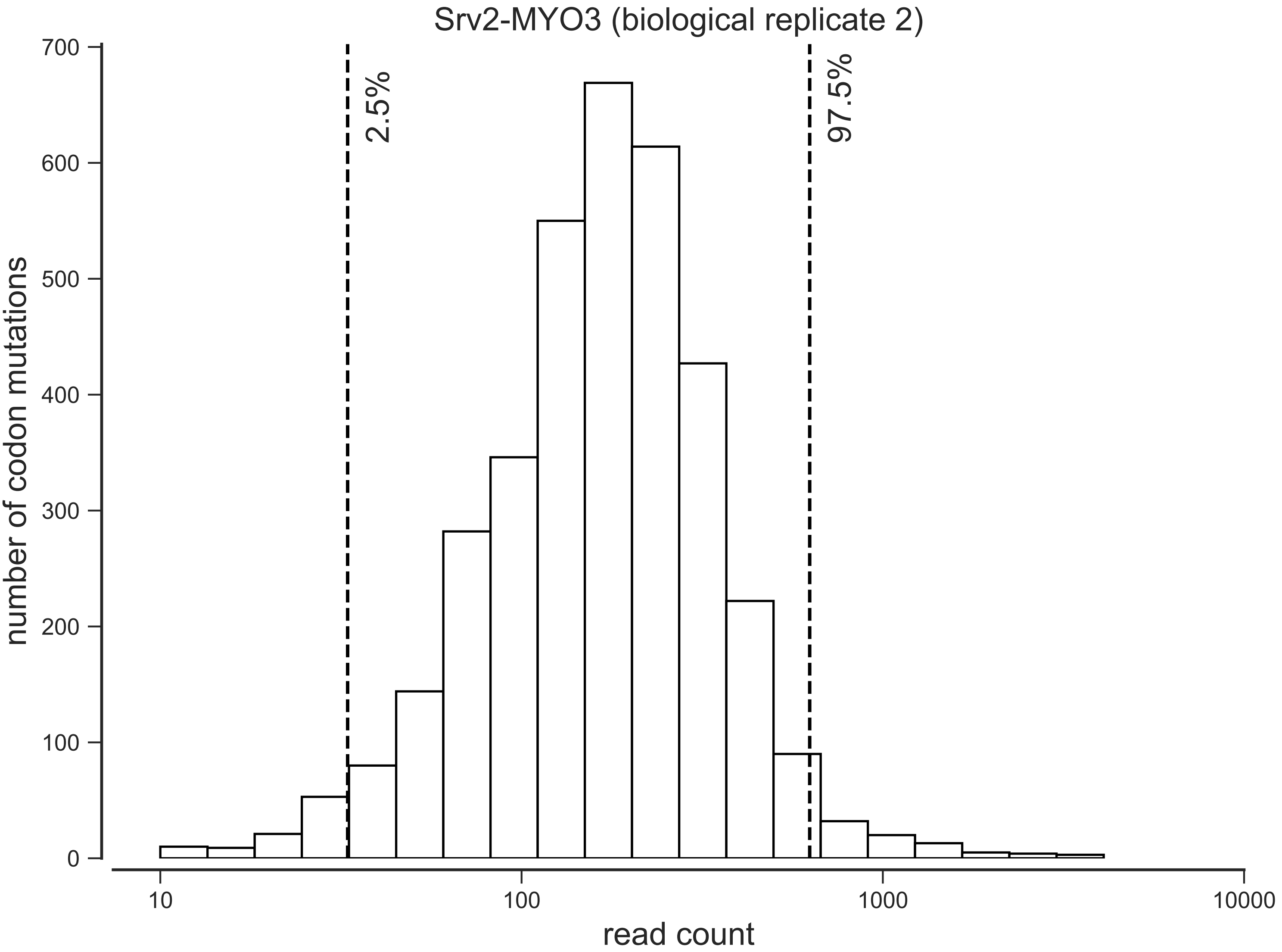
Srv2-MYO3 (biological replicate 2)
number of codon mutations
read count
2.5%
97.5%
0
100
200
300
400
500
600
700
10
100
1000
10000

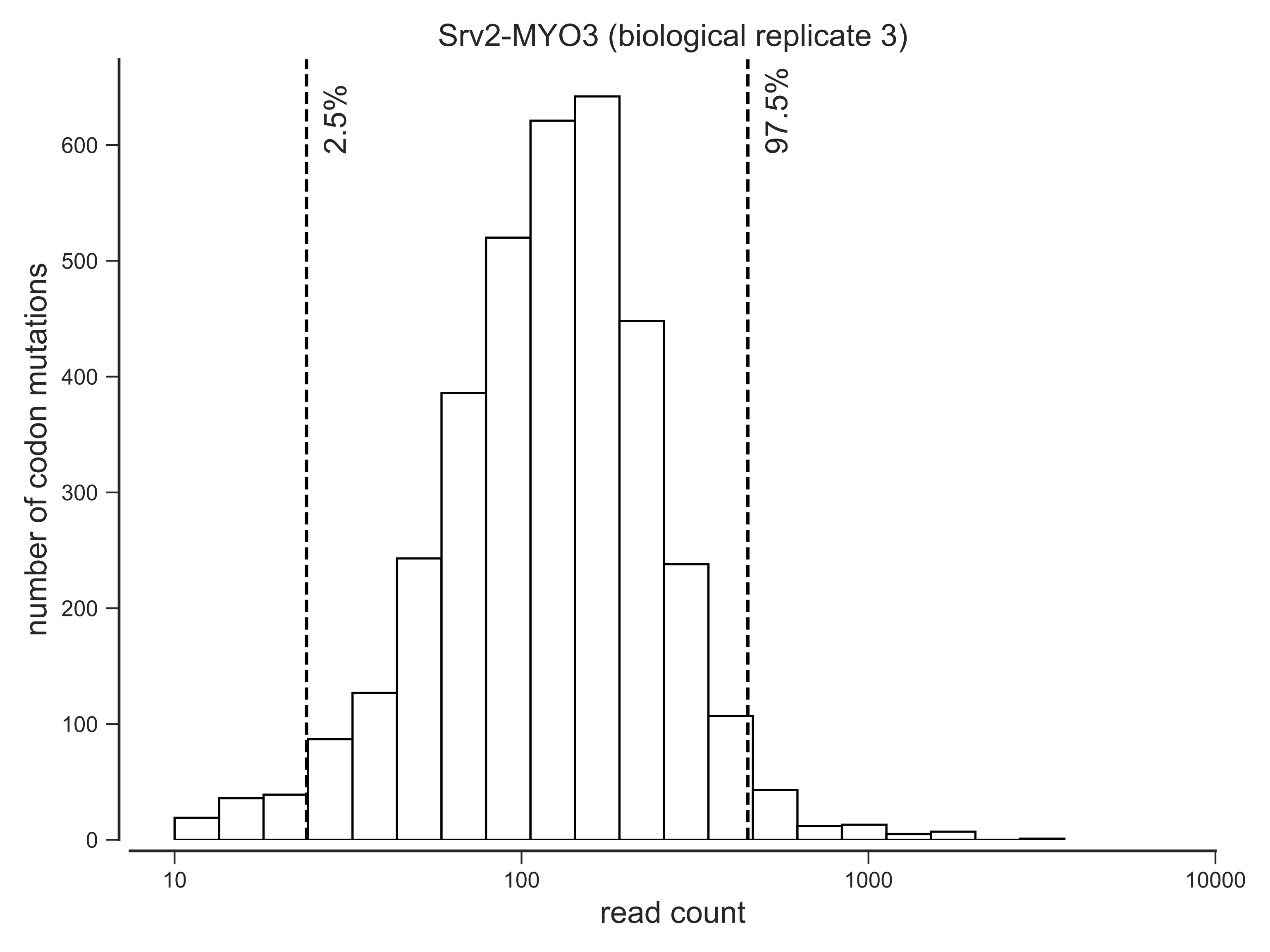

Srv2-MYO3 (biological replicate 3)
2.5%
97.5%
number of codon mutations
0
100
200
300
400
500
600
10
100
1000
10000
read count

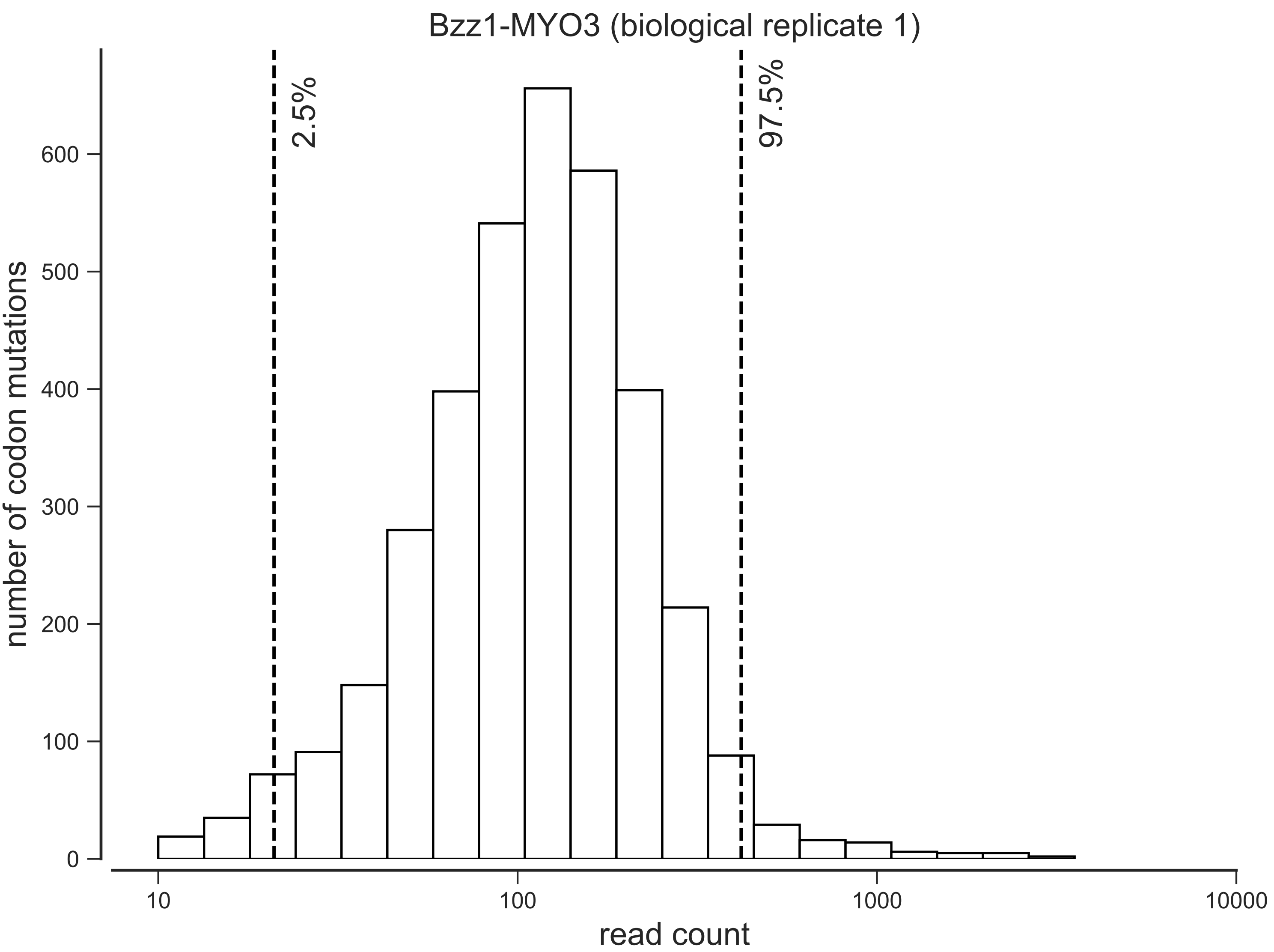

Bzz1-MYO3 (biological replicate 1)
2.5%
97.5%
number of codon mutations
0
100
200
300
400
500
600
10
100
1000
10000
read count

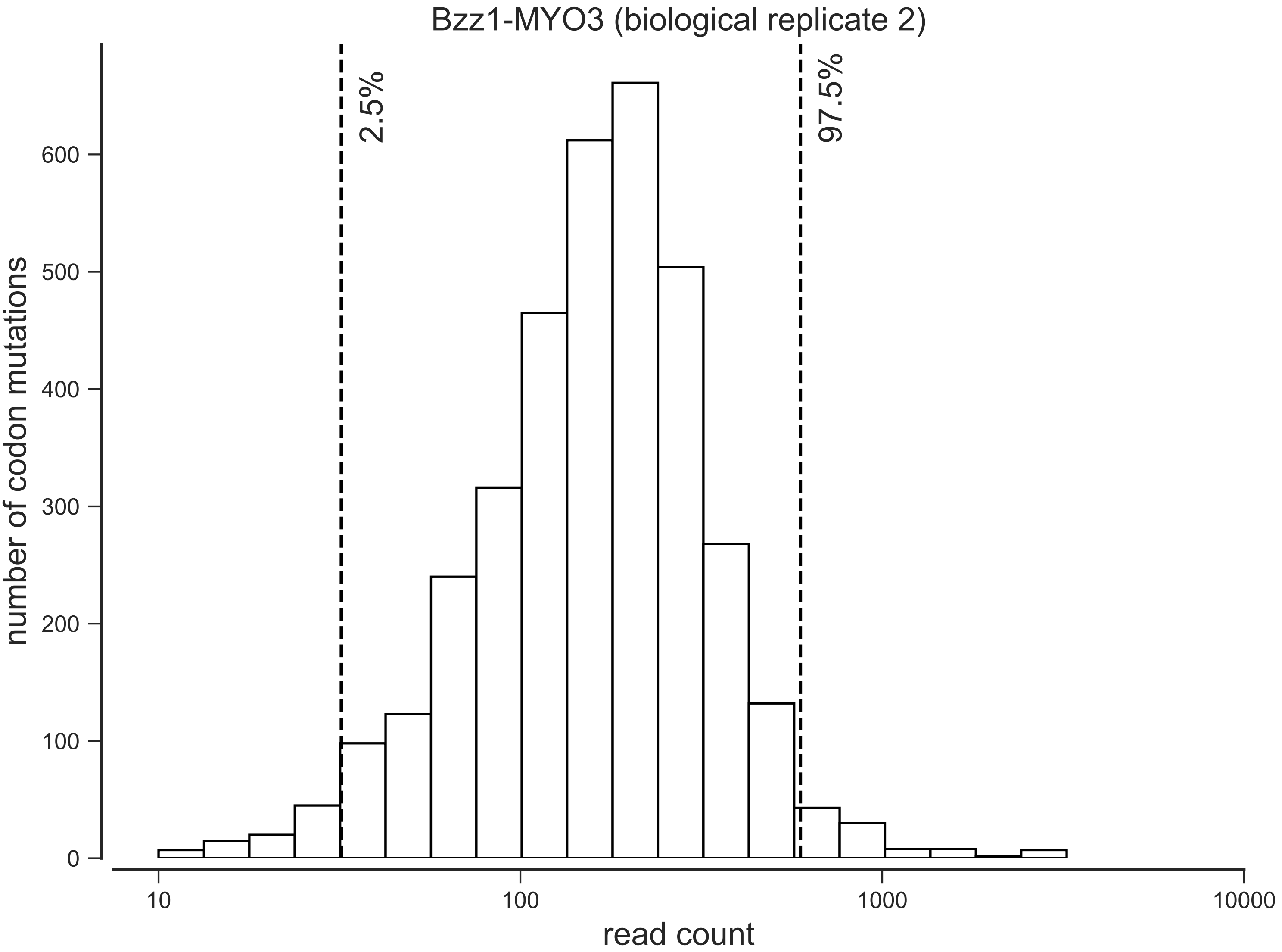

Bzz1-MYO3 (biological replicate 2)
2.5%
97.5%
number of codon mutations
0
100
200
300
400
500
600
10
100
1000
10000
read count

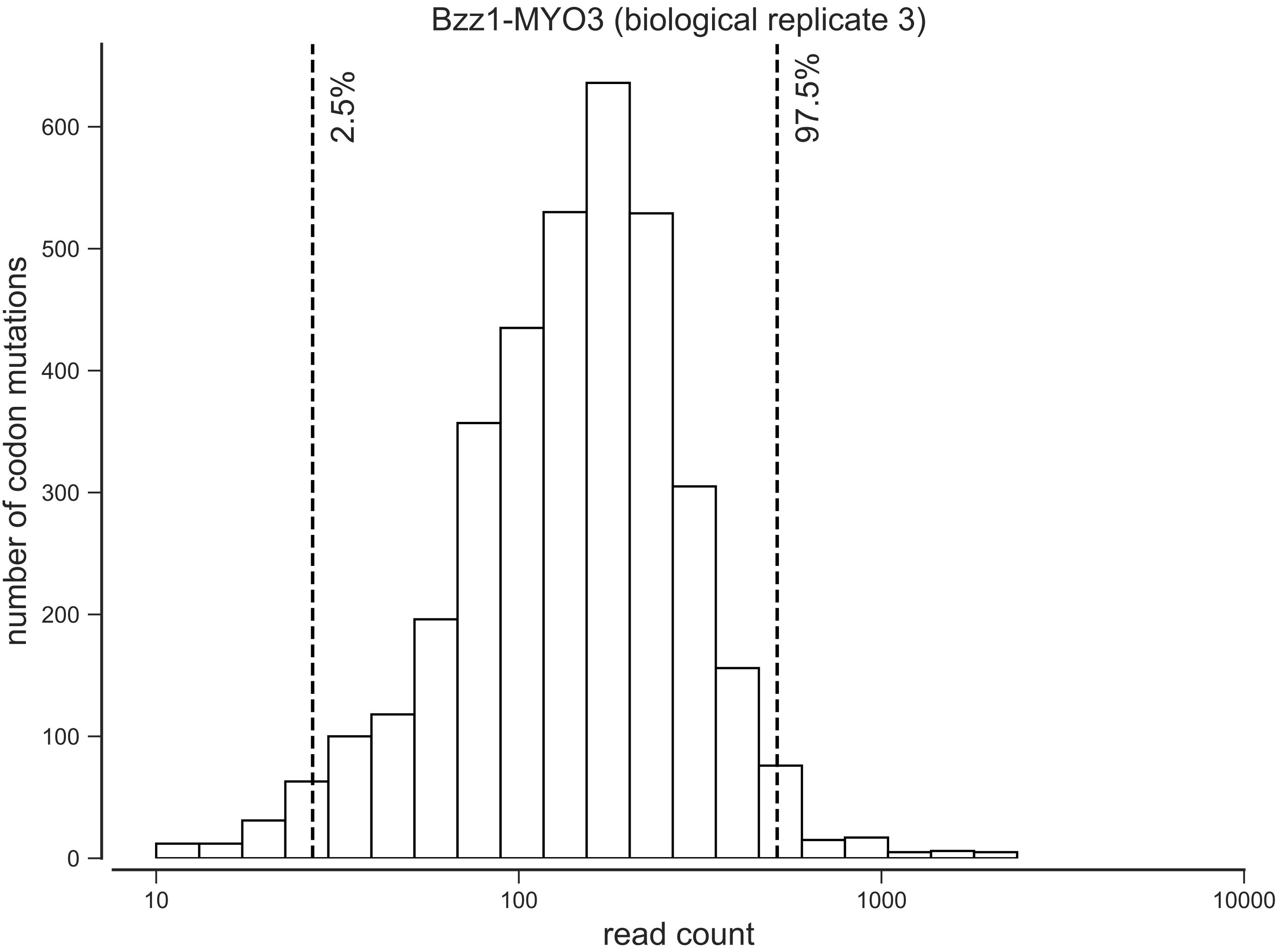

Bzz1-MYO3 (biological replicate 3)
2.5%
97.5%
number of codon mutations
0
100
200
300
400
500
600
10
100
1000
10000
read count

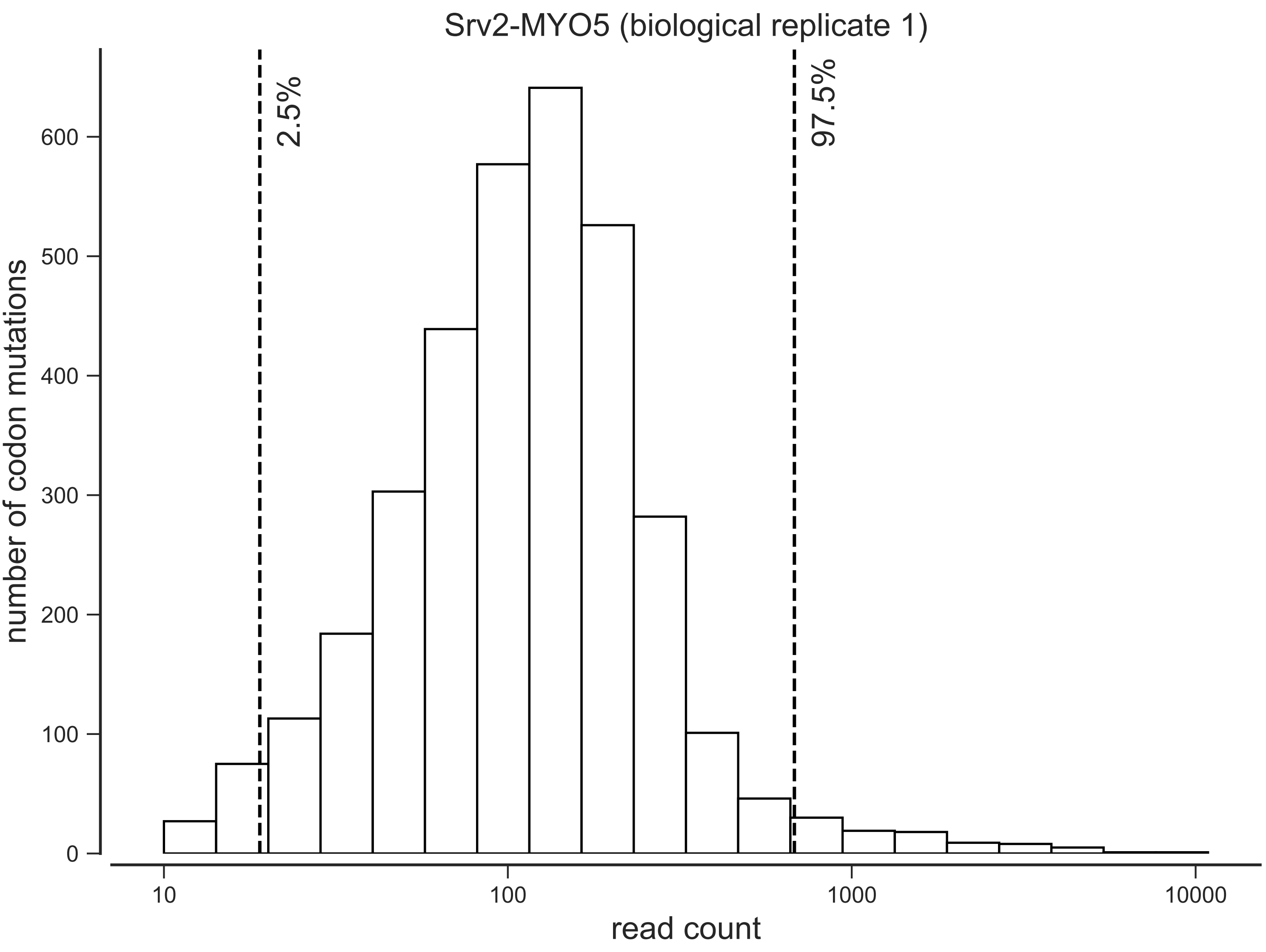
Srv2-MYO5 (biological replicate 1)
2.5%
97.5%
number of codon mutations
0
100
200
300
400
500
600
10
100
1000
10000
read count

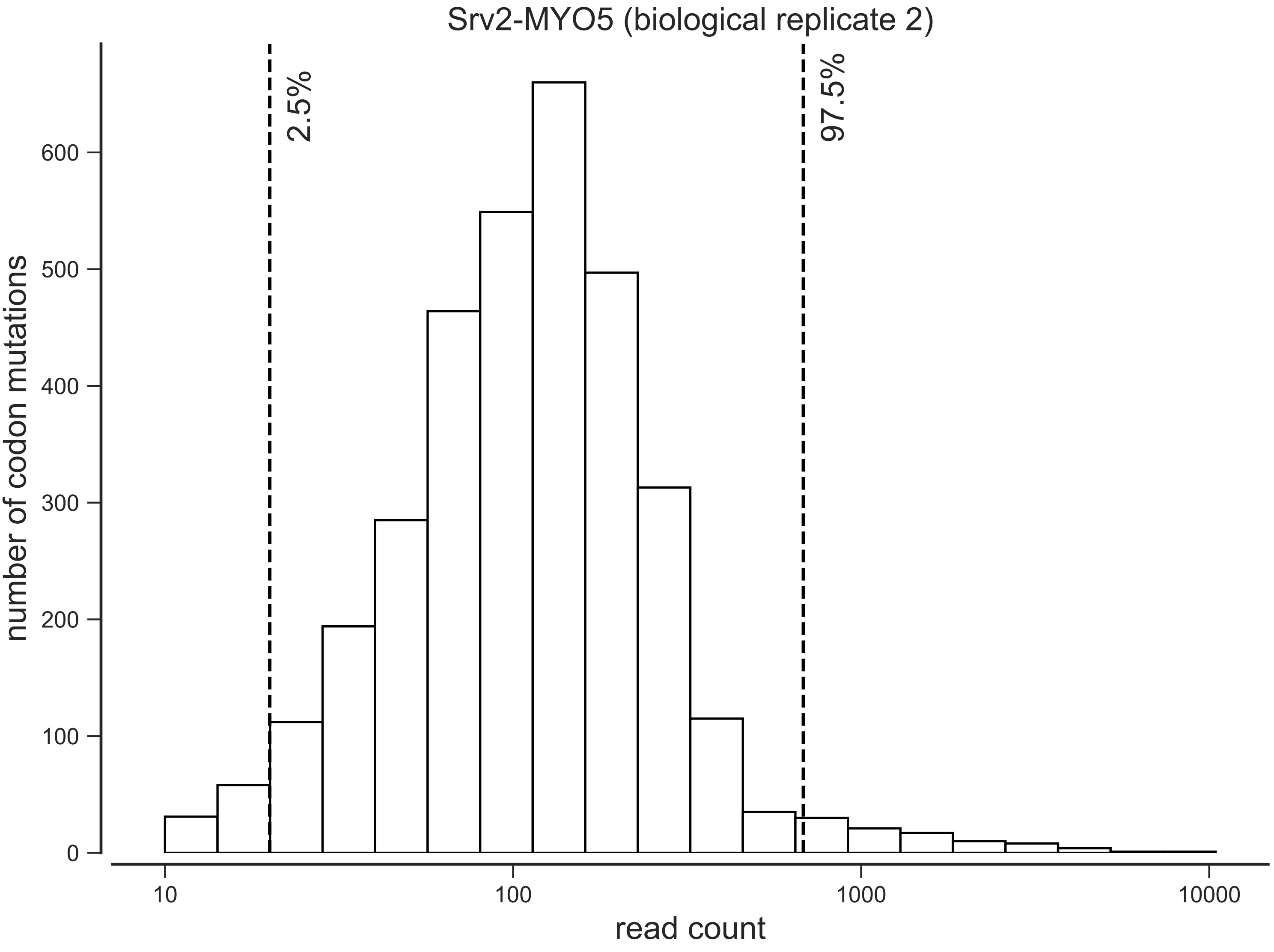

Srv2-MYO5 (biological replicate 2)
2.5%
97.5%
number of codon mutations
0
100
200
300
400
500
600
10
100
1000
10000
read count

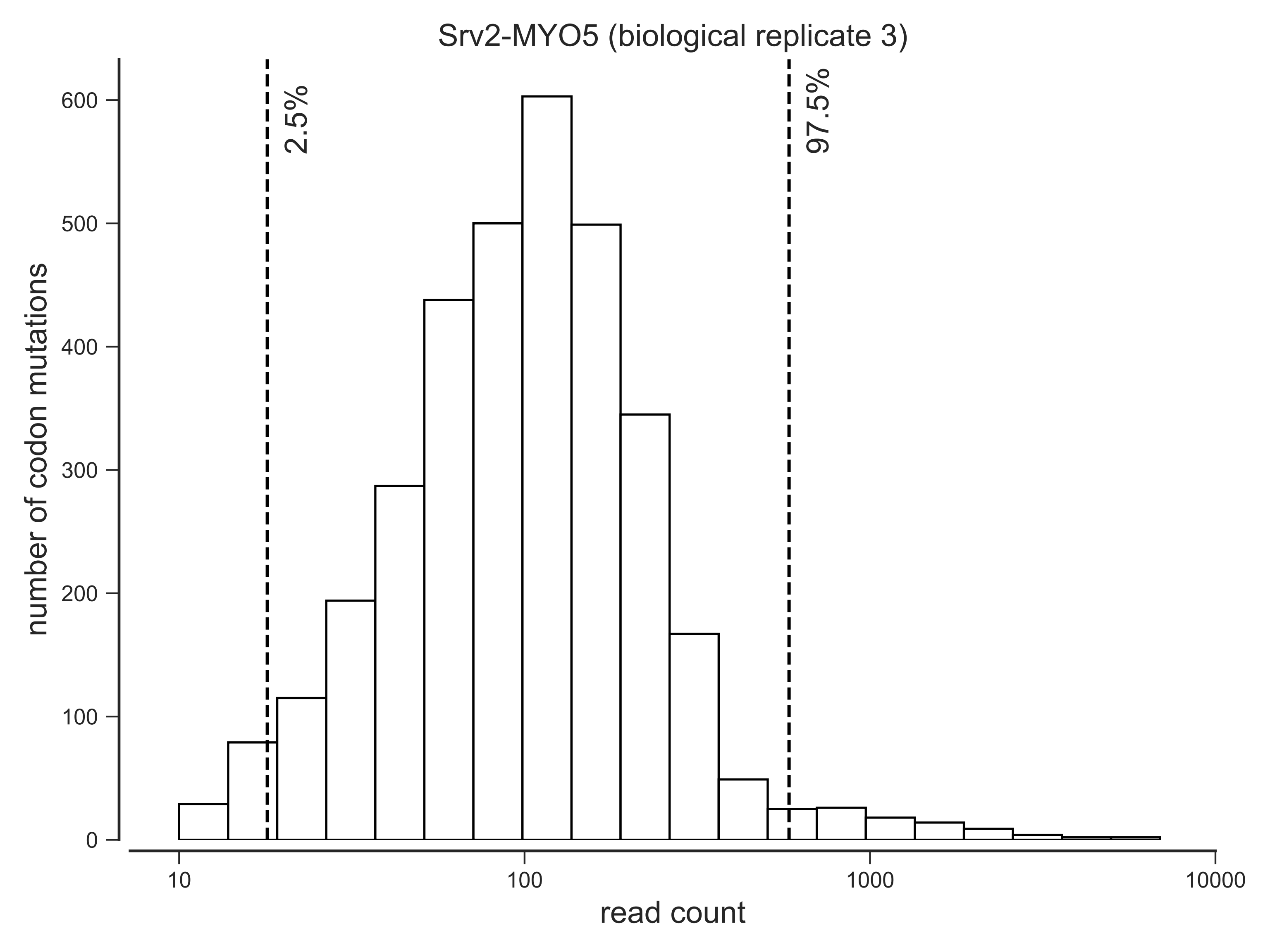
Srv2-MYO5 (biological replicate 3)
2.5%
97.5%
number of codon mutations
0
100
200
300
400
500
600
10
100
1000
10000
read count

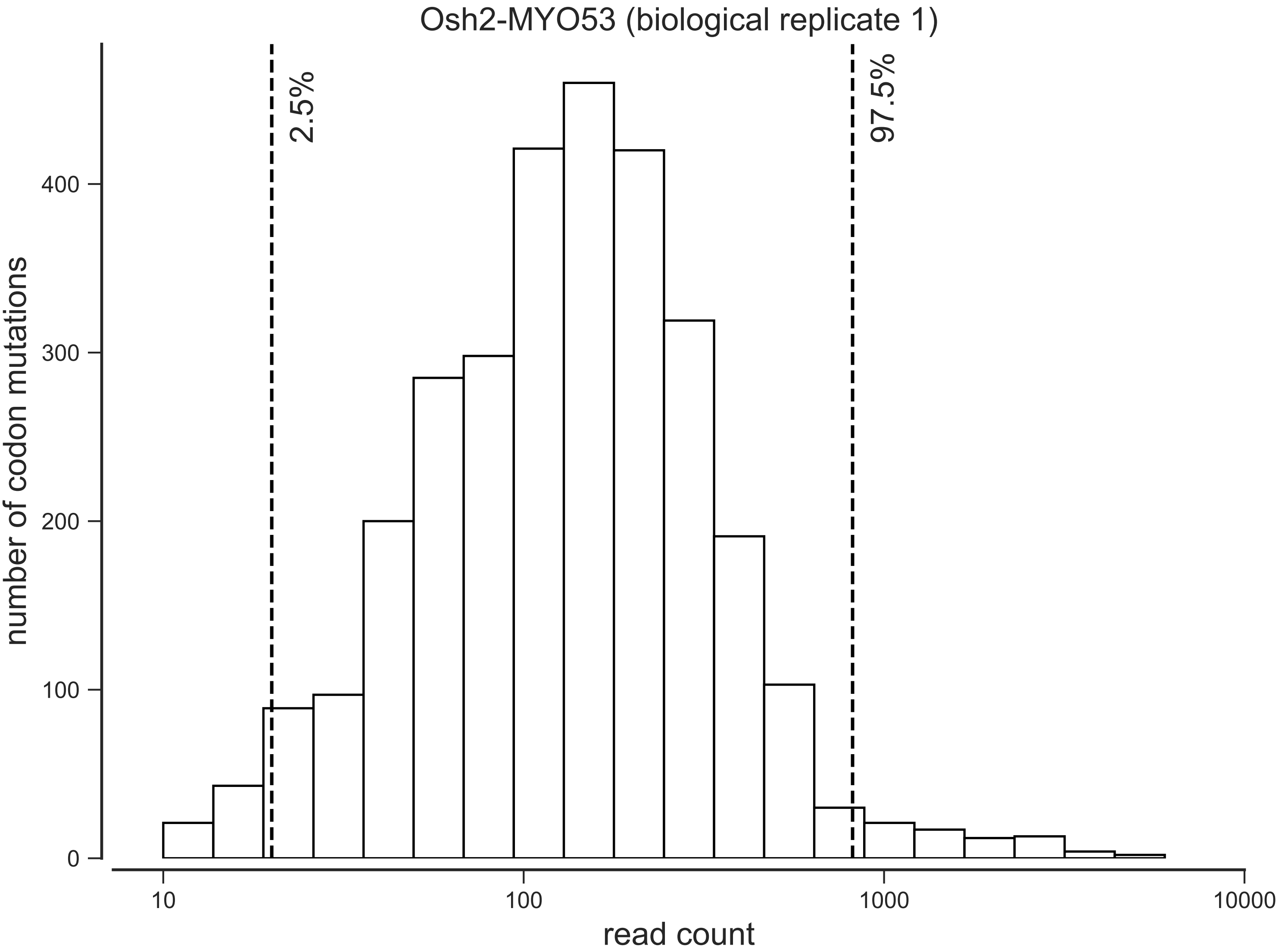
Osh2-MYO53 (biological replicate 1)
2.5%
97.5%
number of codon mutations
0
100
200
300
400
10
100
1000
10000
read count

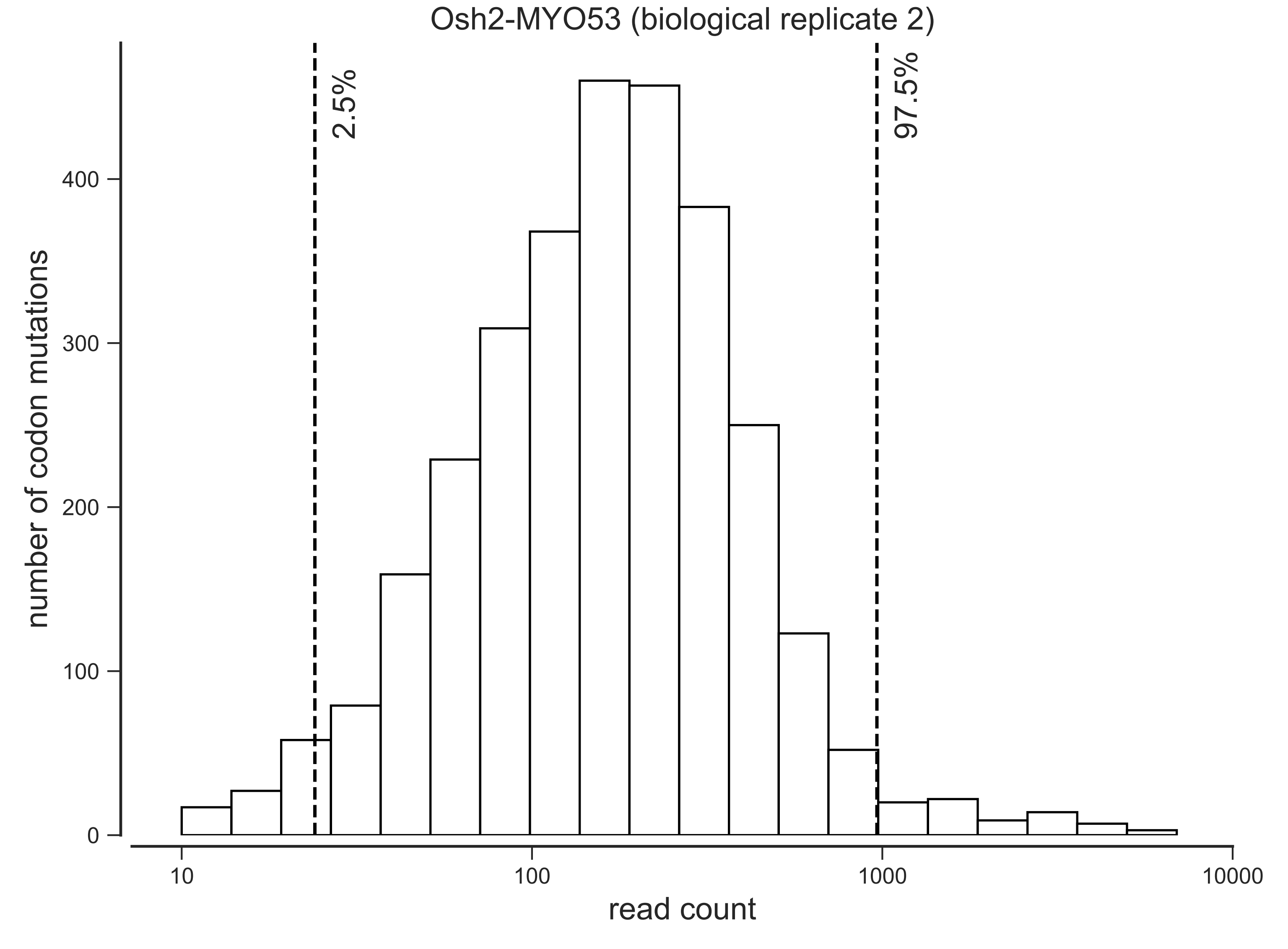
Osh2-MYO53 (biological replicate 2)
2.5%
97.5%
number of codon mutations
0
100
200
300
400
10
100
1000
10000
read count

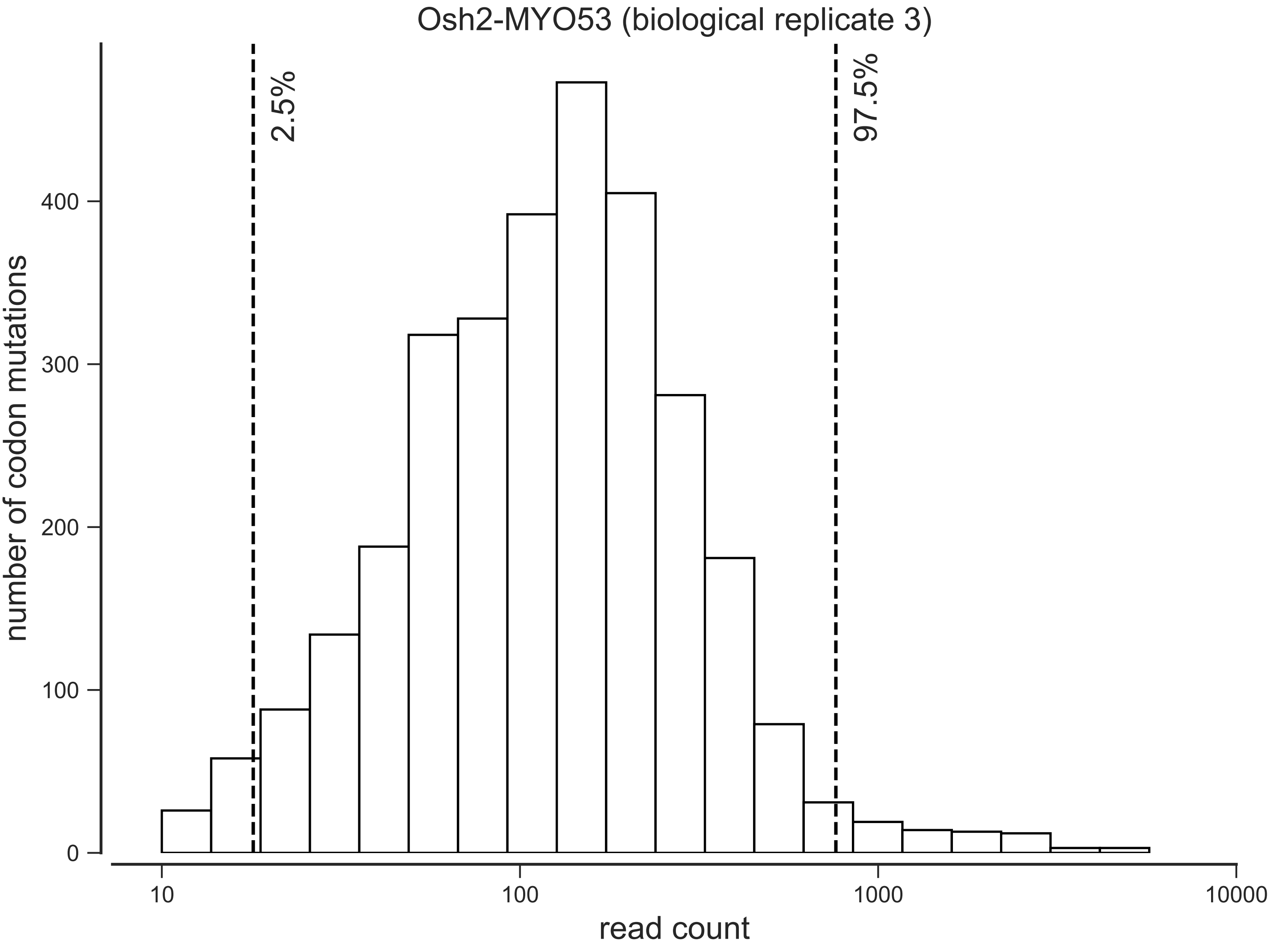
Osh2-MYO53 (biological replicate 3)
2.5%
97.5%
number of codon mutations
0
100
200
300
400
10
100
1000
10000
read count

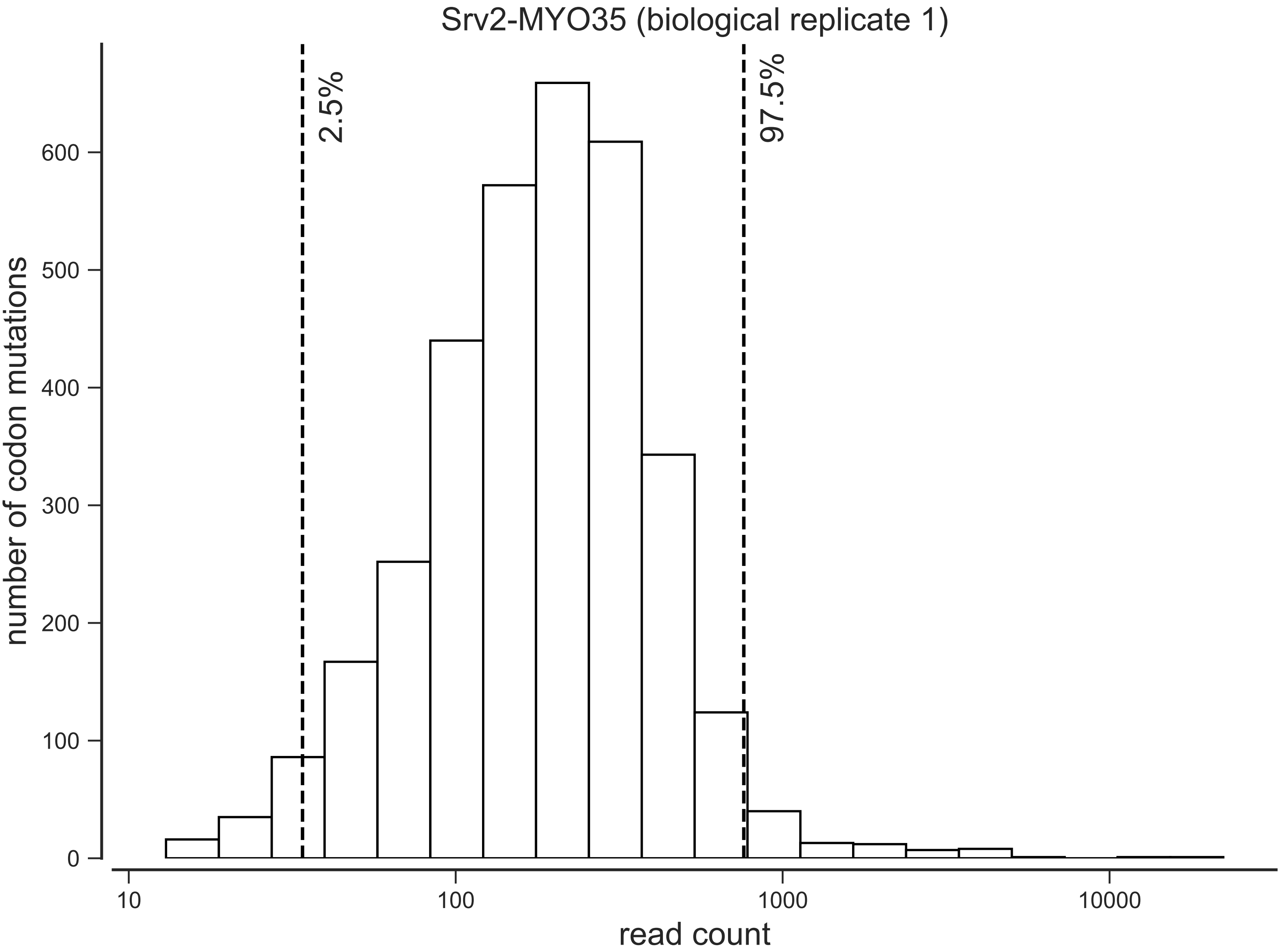

Srv2-MYO35 (biological replicate 1)
2.5%
97.5%
number of codon mutations
0
100
200
300
400
500
600
10
100
1000
10000
read count

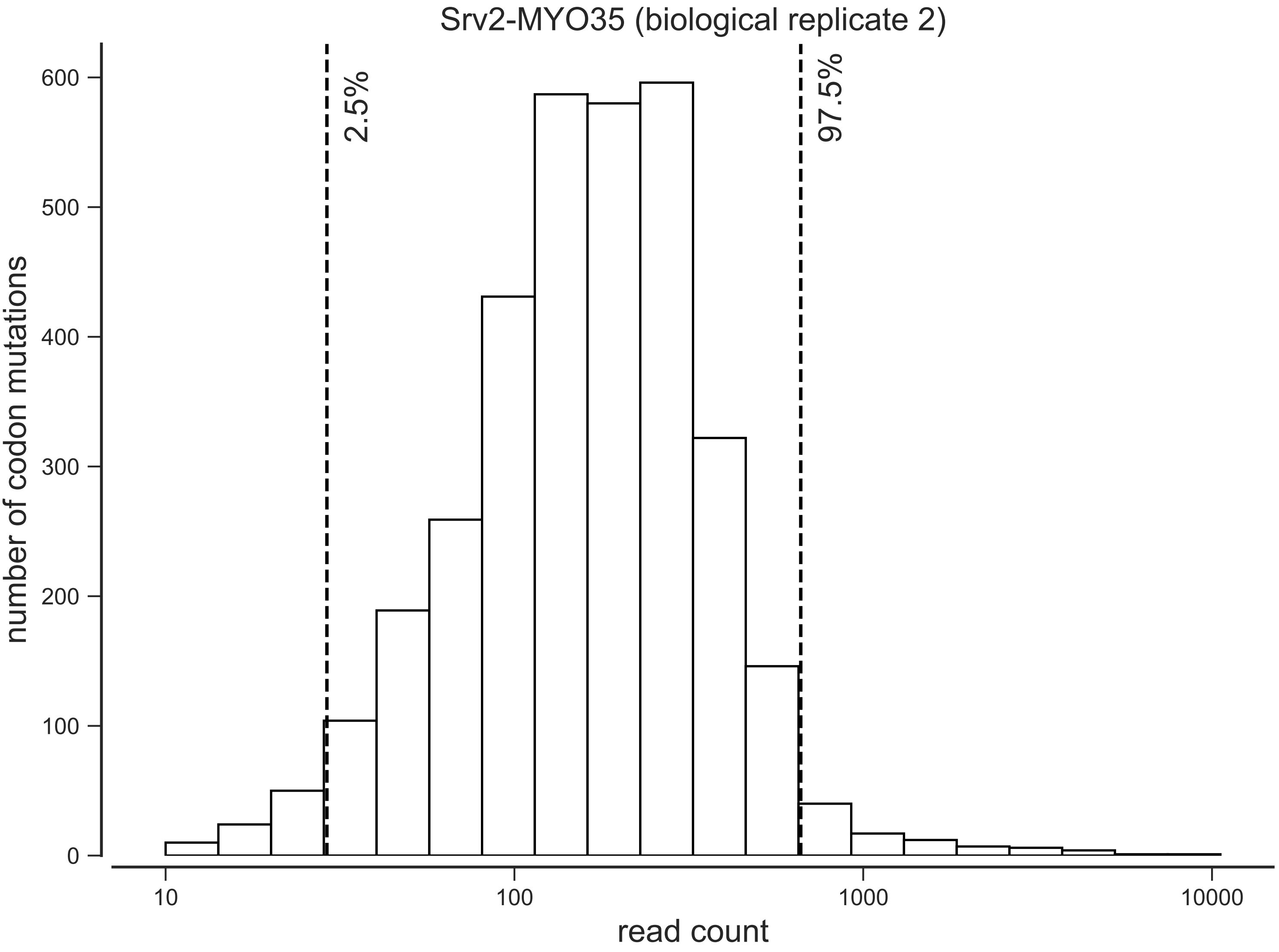
Srv2-MYO35 (biological replicate 2)
2.5%
97.5%
number of codon mutations
0
100
200
300
400
500
600
10
100
1000
10000
read count

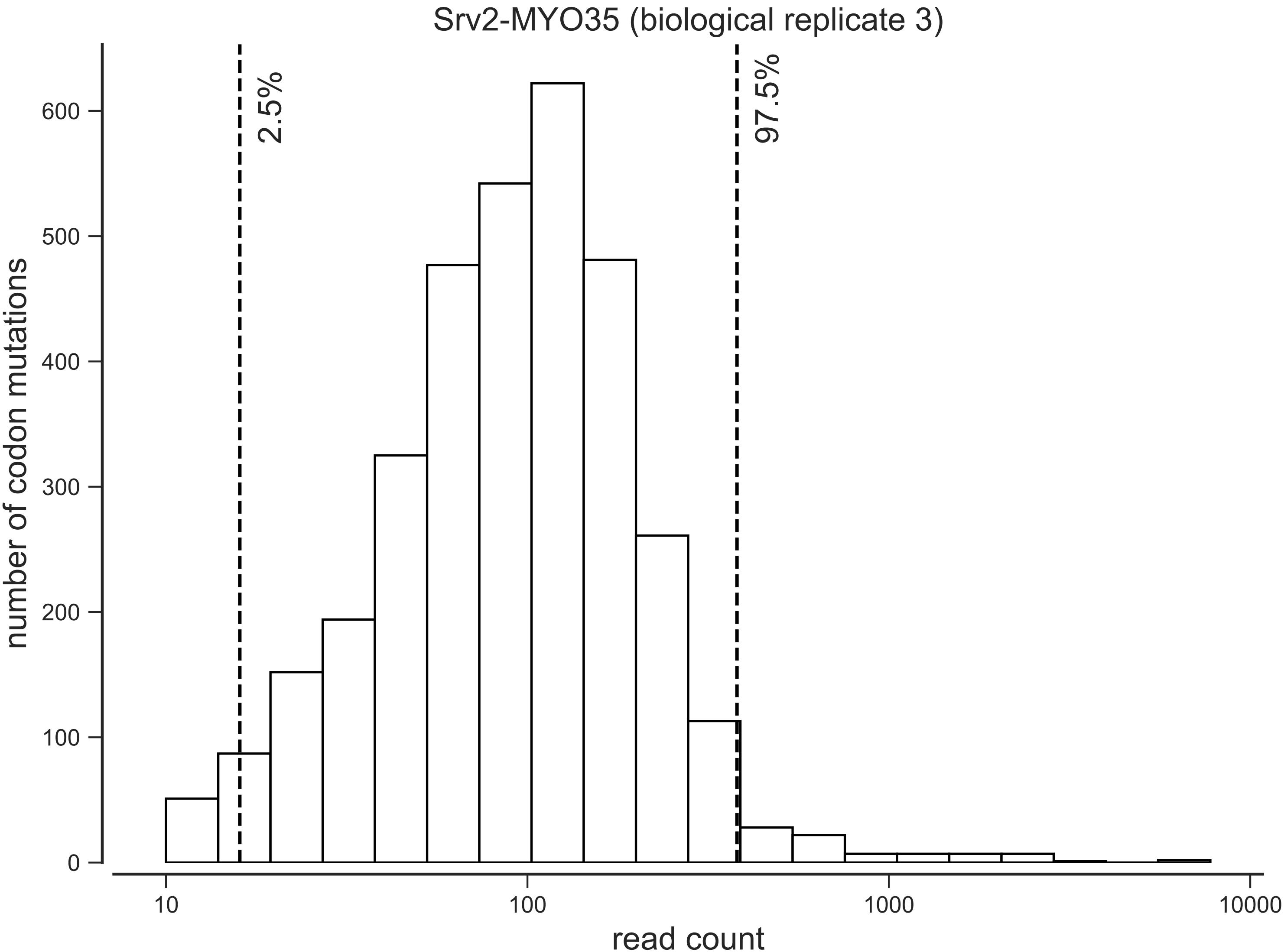

Srv2-MYO35 (biological replicate 3)
2.5%
97.5%
number of codon mutations
0
100
200
300
400
500
600
10
100
1000
10000
read count

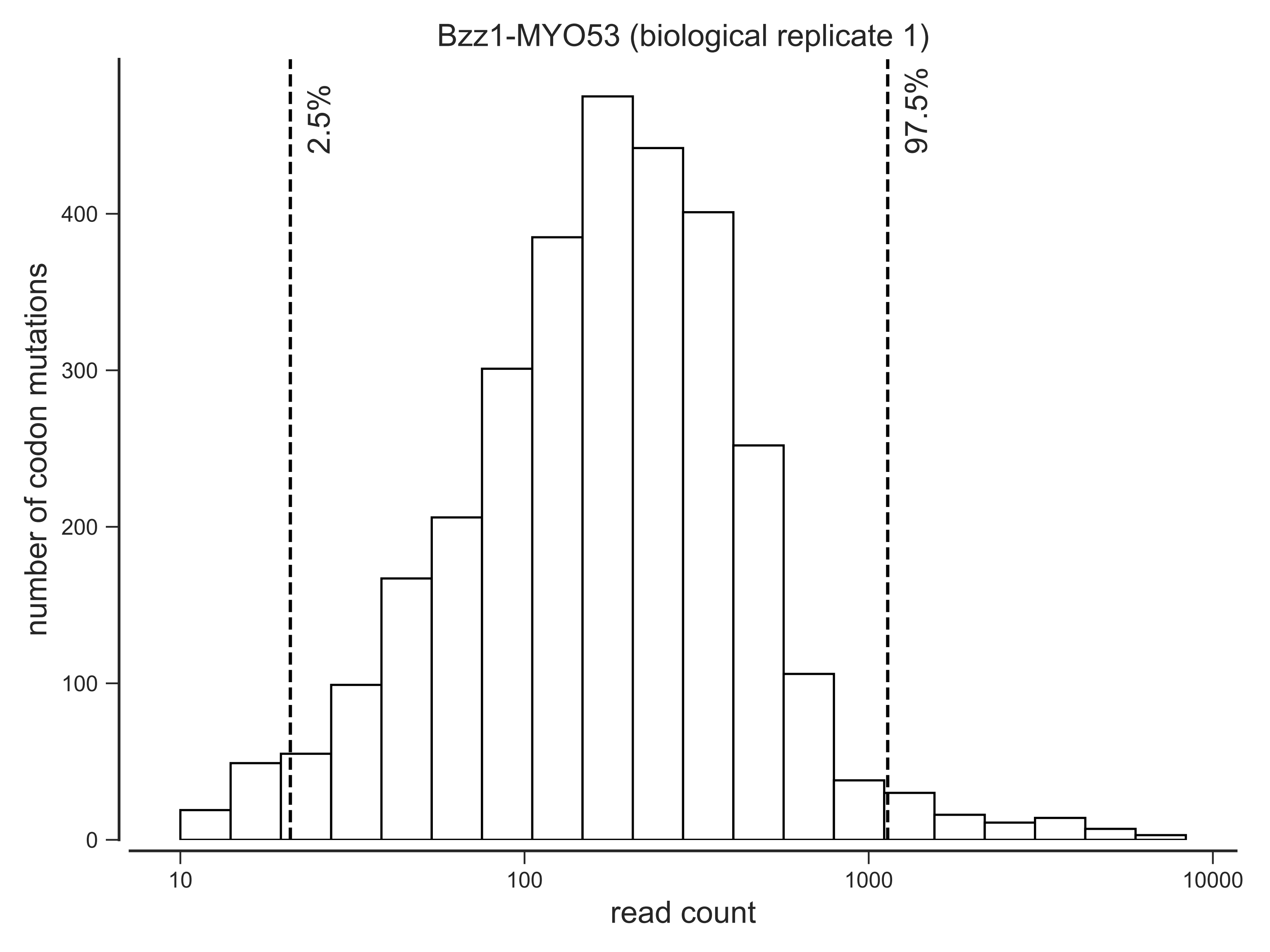
Bzz1-MYO53 (biological replicate 1)
2.5%
97.5%
number of codon mutations
0
100
200
300
400
10
100
1000
10000
read count

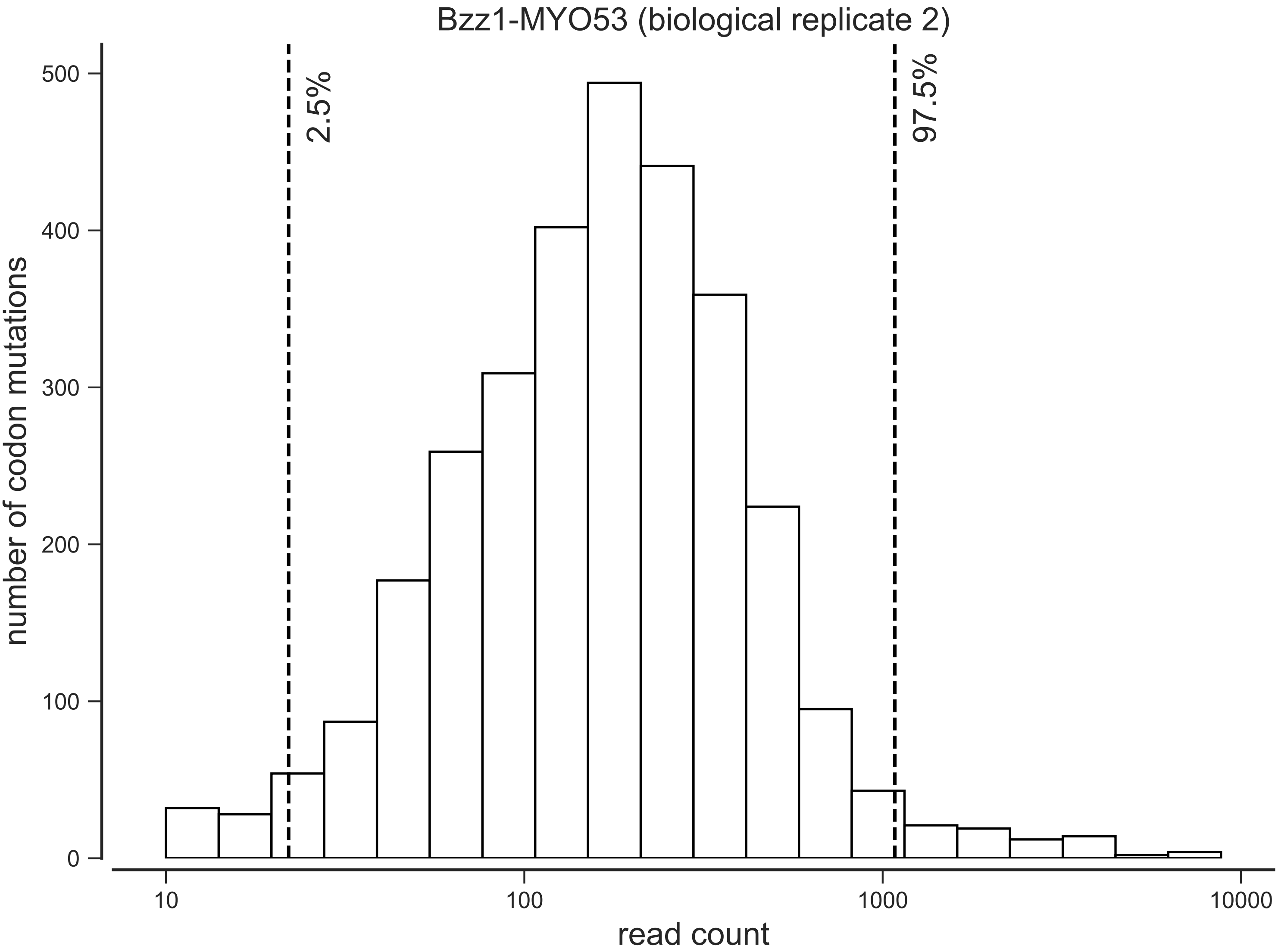

Bzz1-MYO53 (biological replicate 2)
2.5%
97.5%
number of codon mutations
0
100
200
300
400
500
read count
10
100
1000
10000

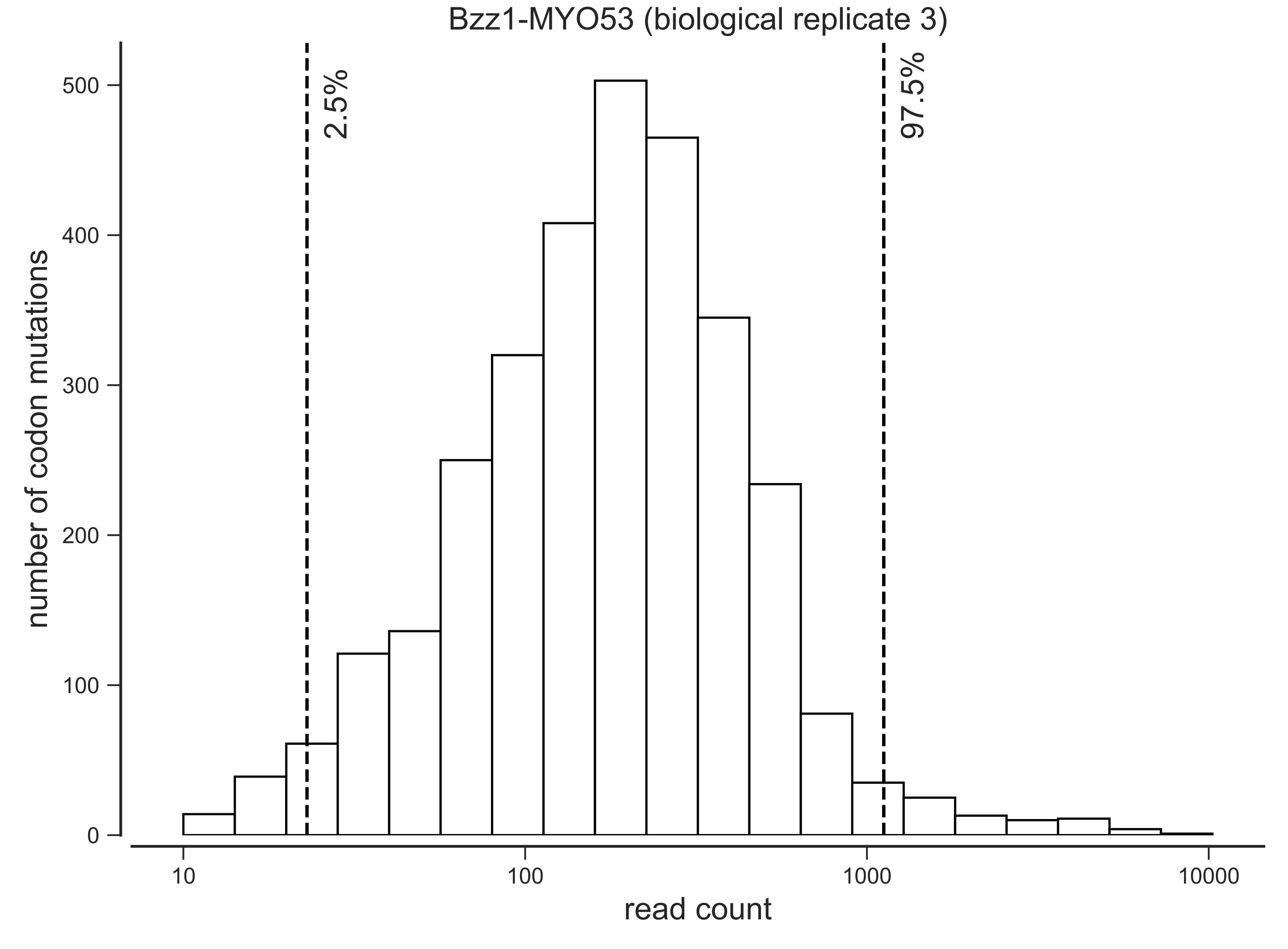
Bzz1-MYO53 (biological replicate 3)
2.5%
97.5%
number of codon mutations
read count
0
100
200
300
400
500
10
100
1000
10000

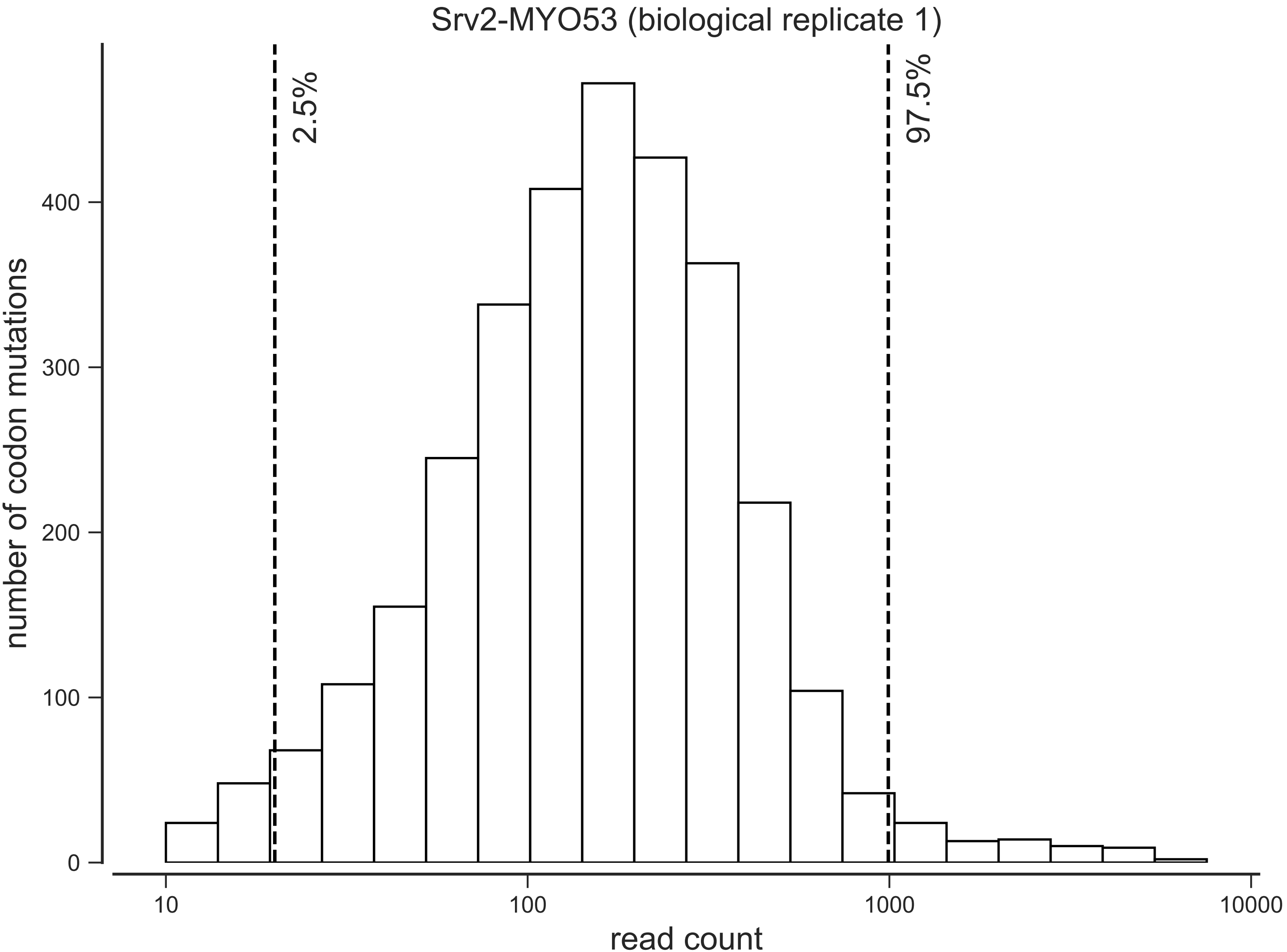

Srv2-MYO53 (biological replicate 1)
2.5%
97.5%
number of codon mutations
0
100
200
300
400
10
100
1000
10000
read count

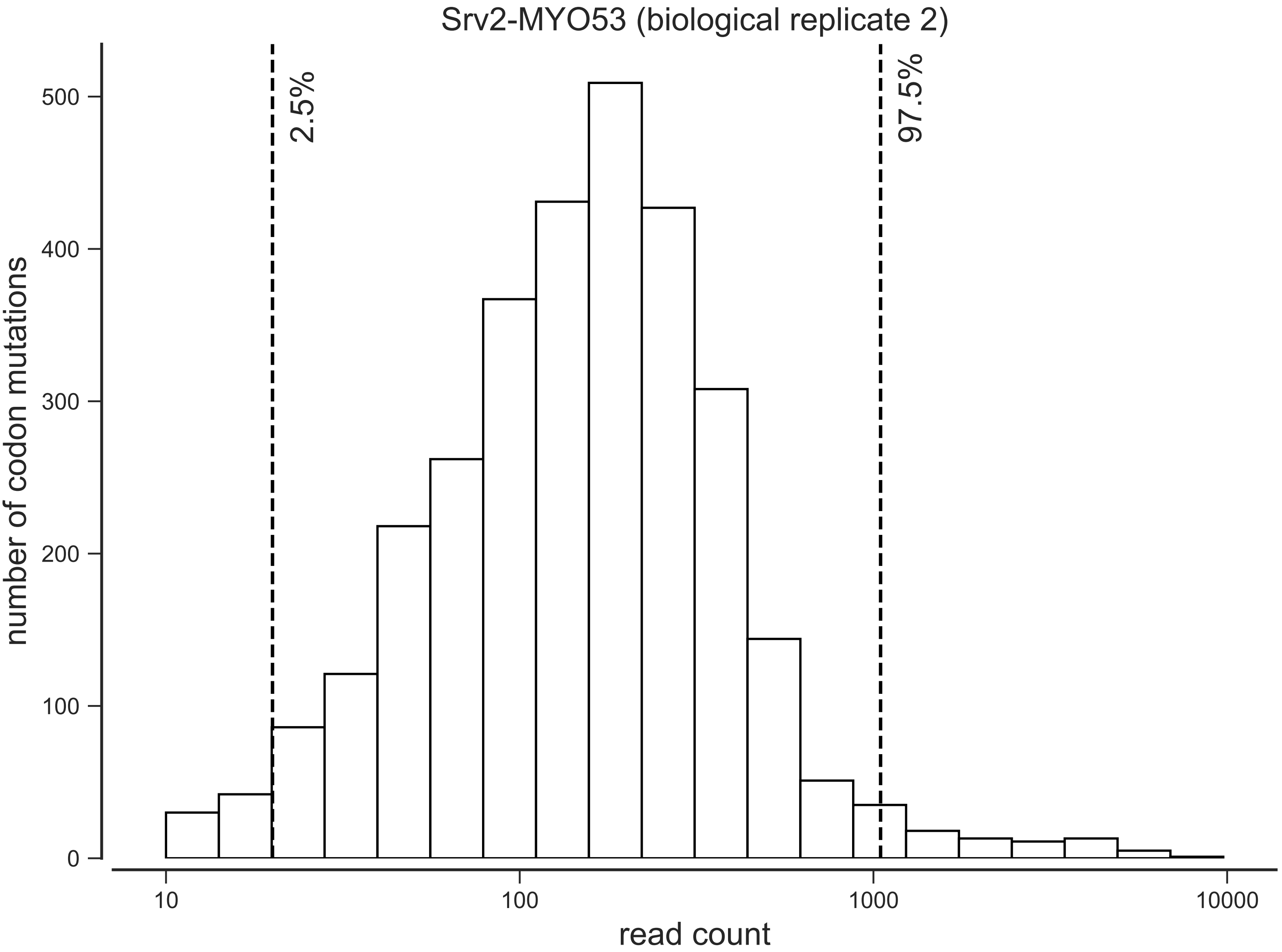

Srv2-MYO53 (biological replicate 2)
2.5%
97.5%
number of codon mutations
0
100
200
300
400
500
10
100
1000
10000
read count

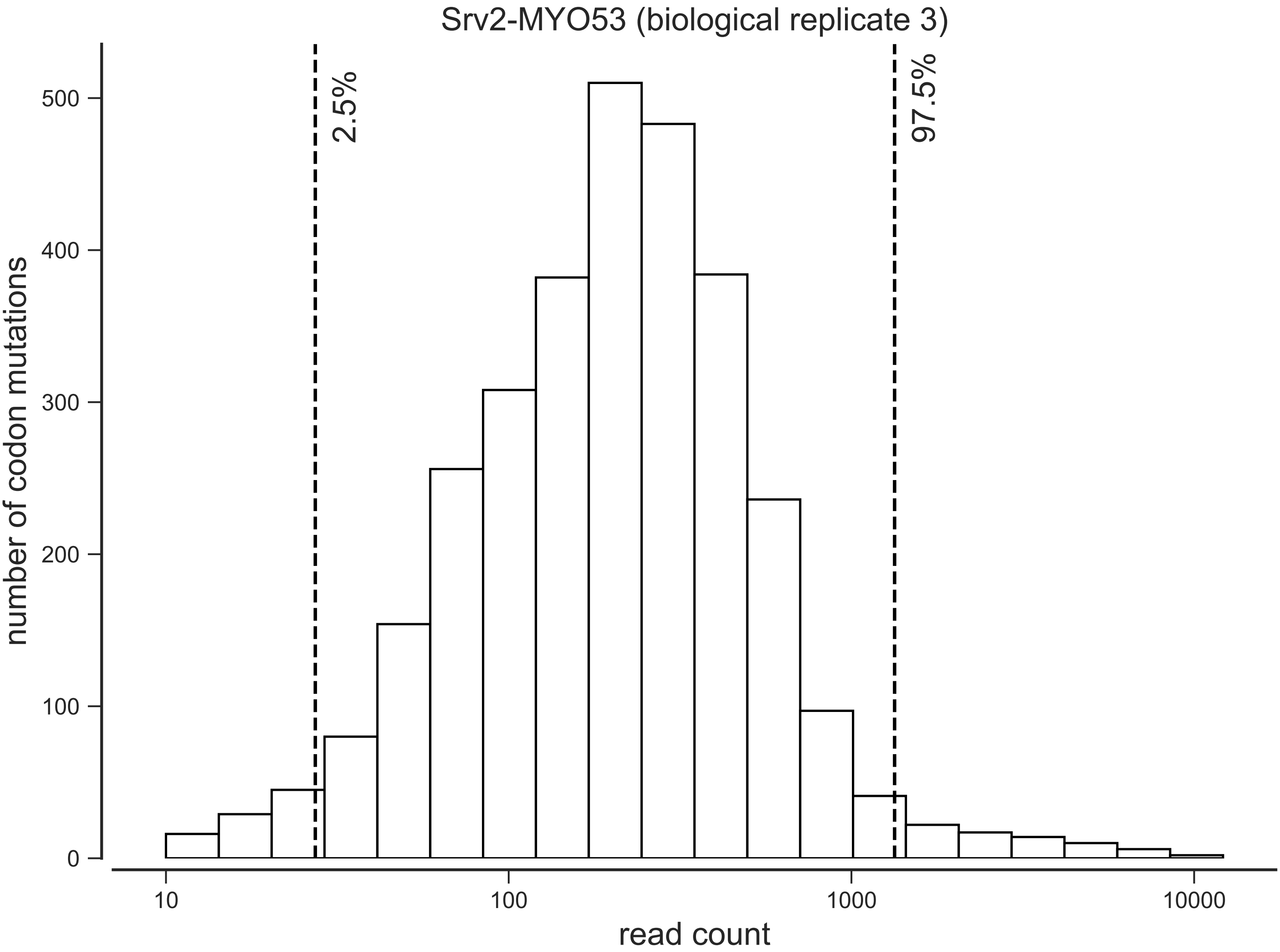

Srv2-MYO53 (biological replicate 3)
number of codon mutations
read count
2.5%
97.5%
0
100
200
300
400
500
10
100
1000
10000

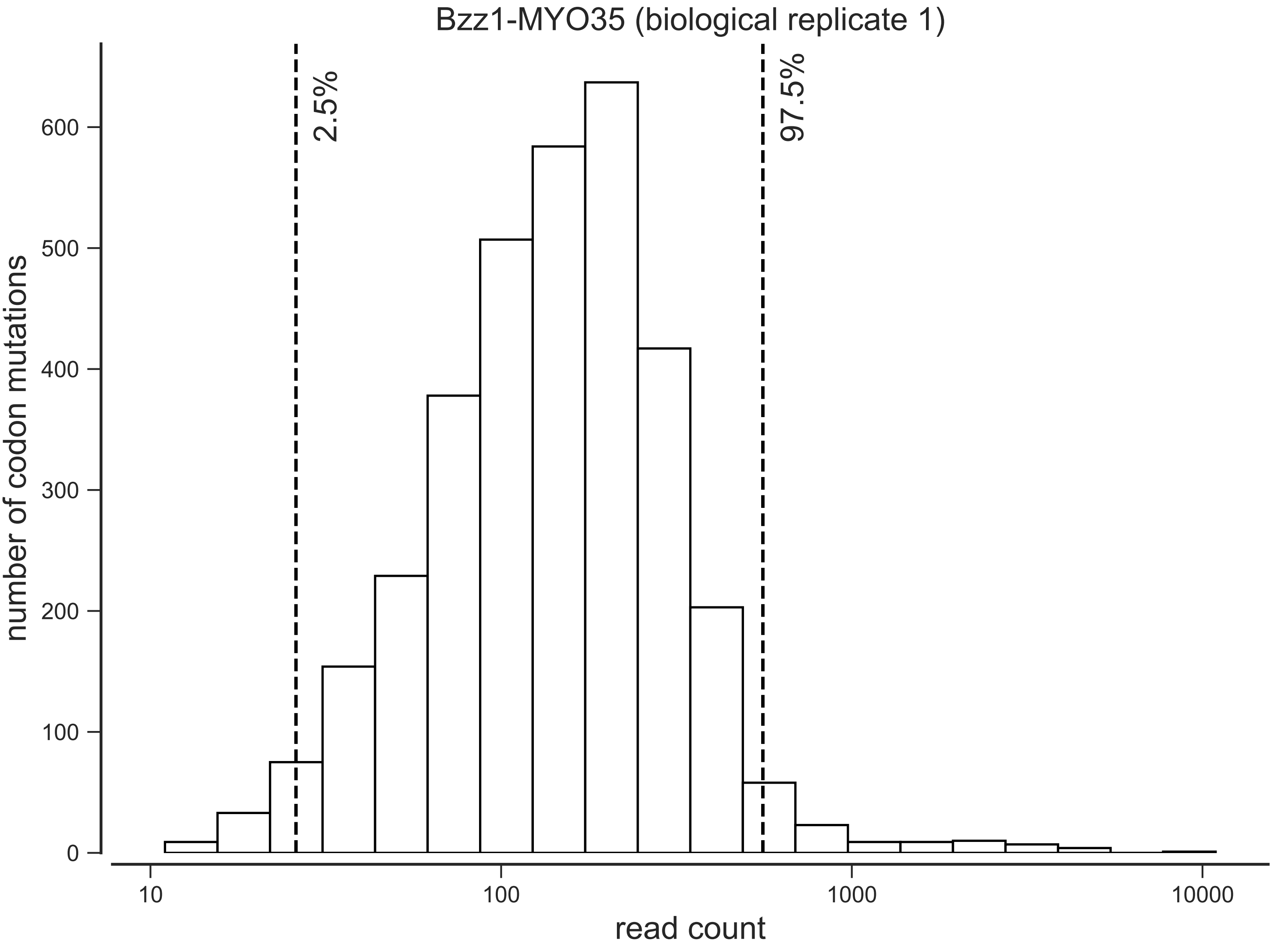
Bzz1-MYO35 (biological replicate 1)
2.5%
97.5%
number of codon mutations
0
100
200
300
400
500
600
10
100
1000
10000
read count

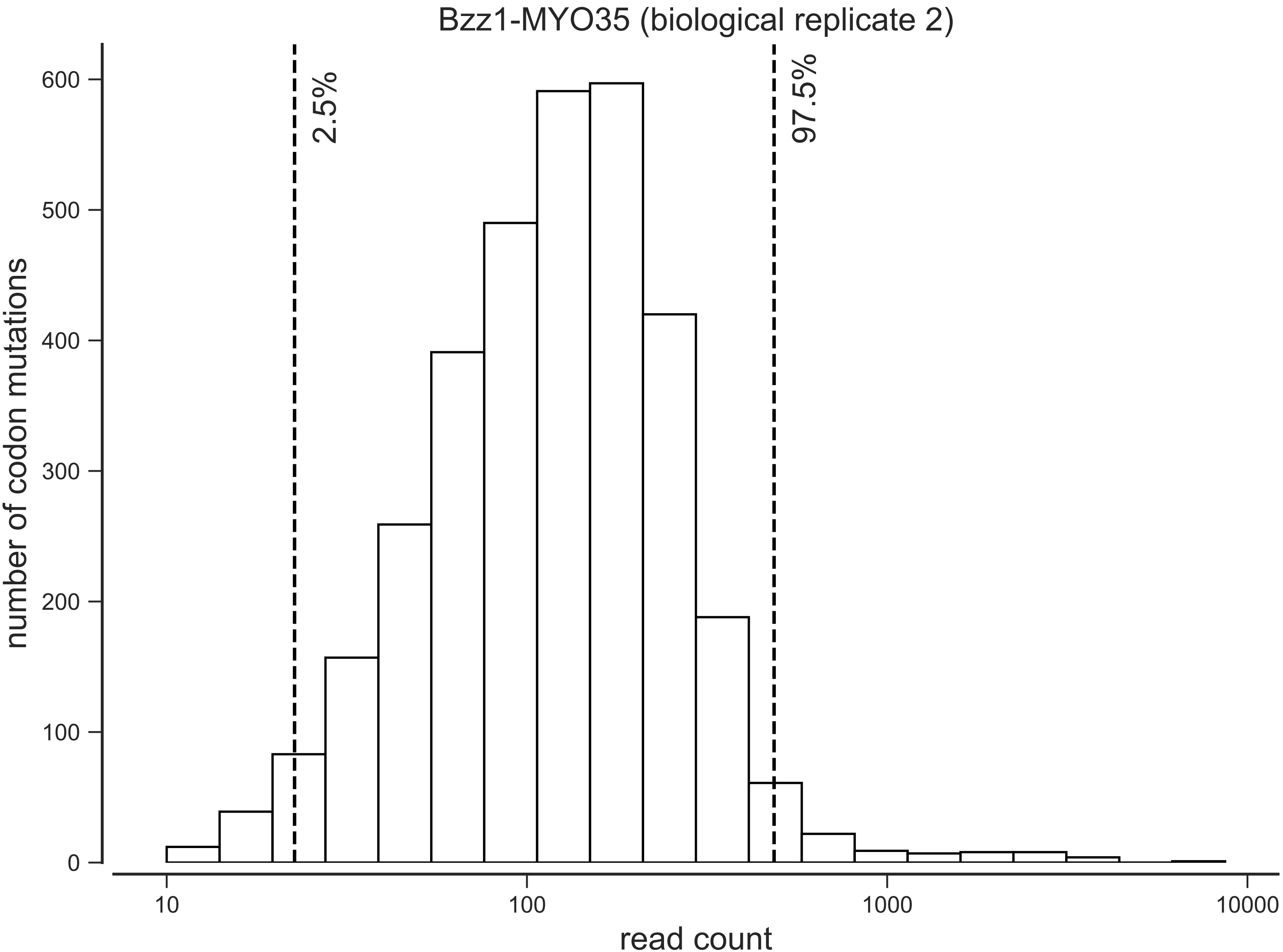
Bzz1-MYO35 (biological replicate 2)
number of codon mutations
read count
2.5%
97.5%
0
100
200
300
400
500
600
10
100
1000
10000

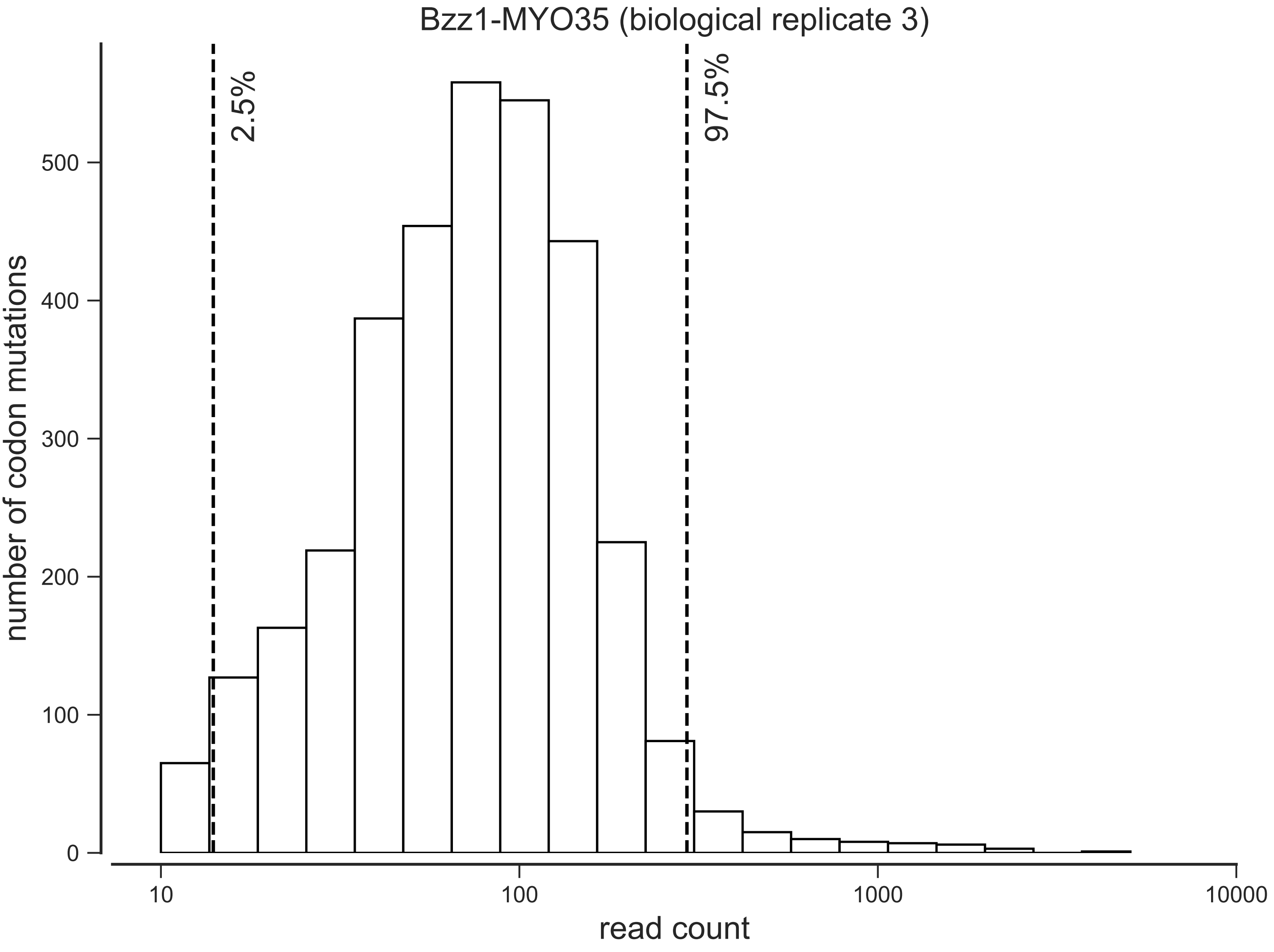

Bzz1-MYO35 (biological replicate 3)
2.5%
97.5%
number of codon mutations
0
100
200
300
400
500
10
100
1000
10000
read count

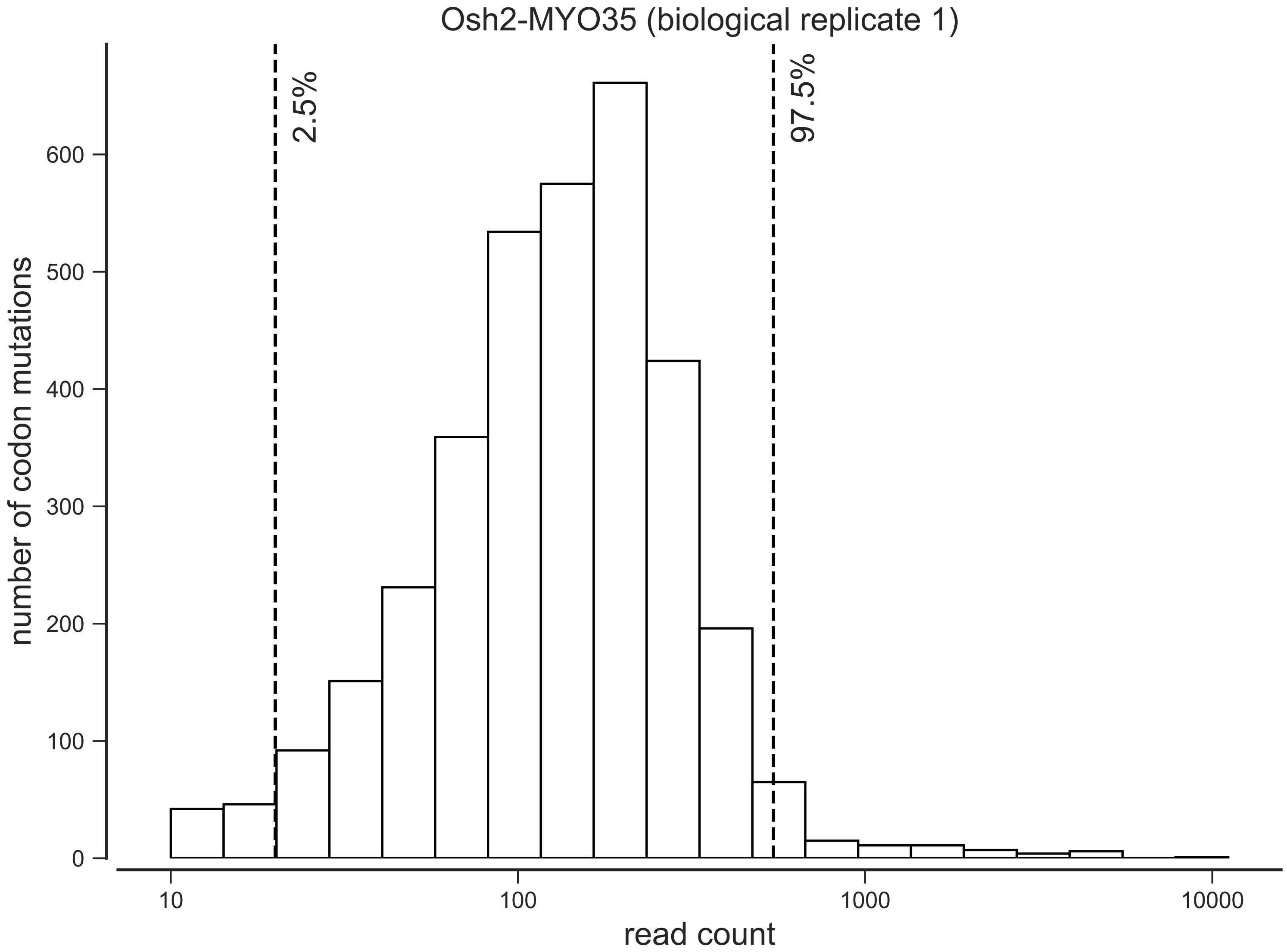

Osh2-MYO35 (biological replicate 1)
2.5%
97.5%
number of codon mutations
0
100
200
300
400
500
600
10
100
1000
10000
read count

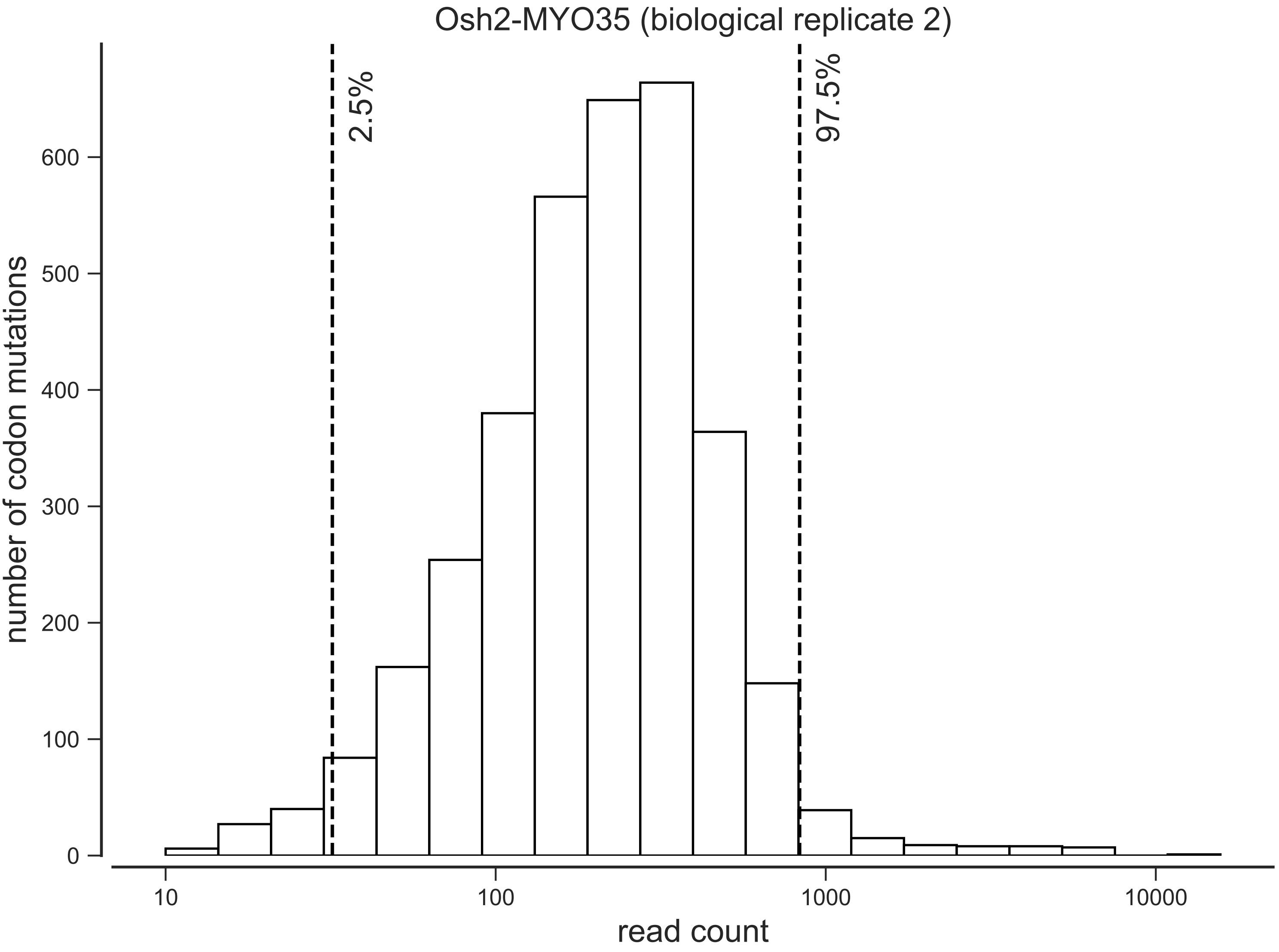
Osh2-MYO35 (biological replicate 2)
2.5%
97.5%
number of codon mutations
0
100
200
300
400
500
600
10
100
1000
10000
read count

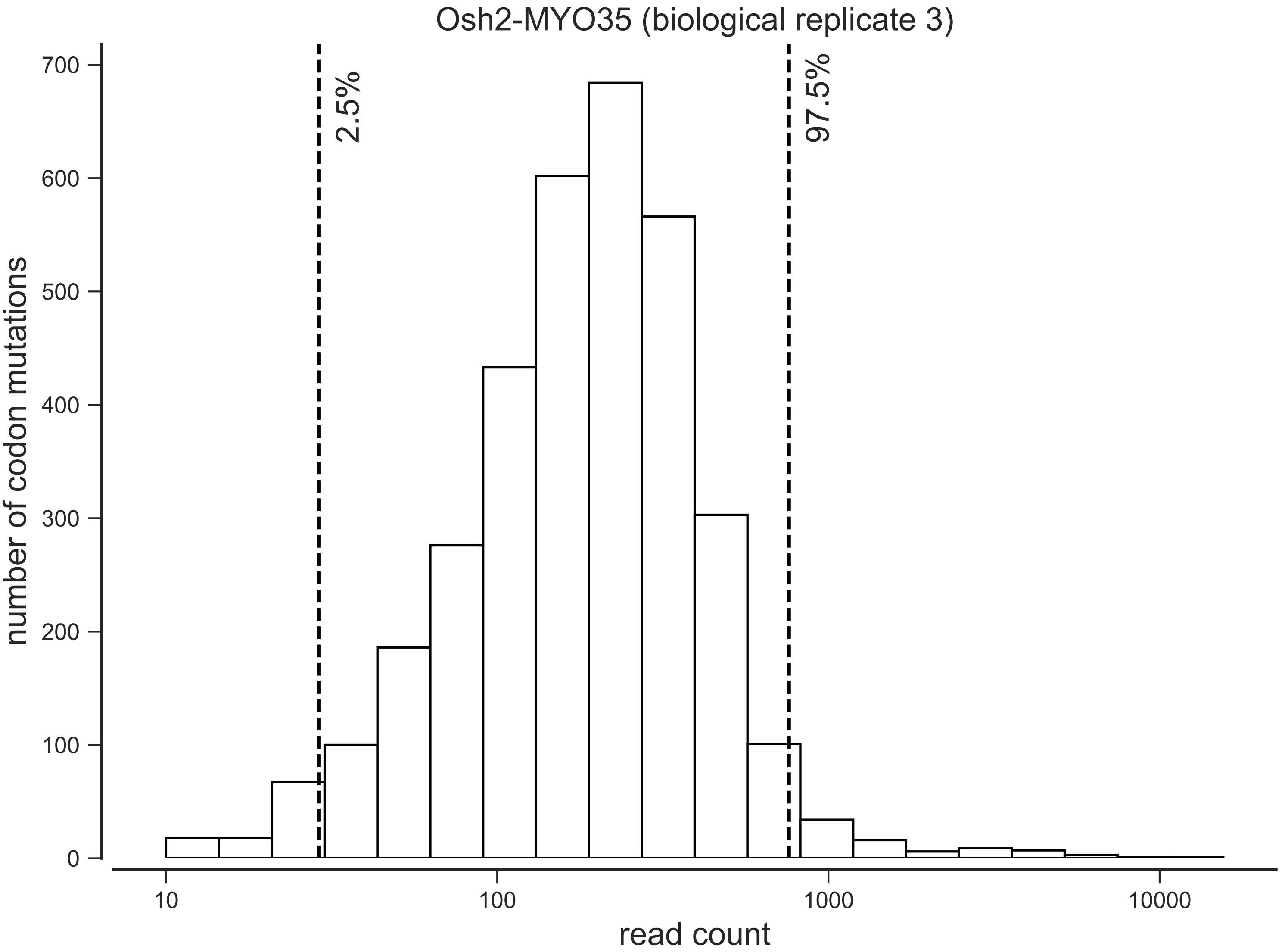

Osh2-MYO35 (biological replicate 3)
2.5%
97.5%
number of codon mutations
0
100
200
300
400
500
600
700
10
100
1000
10000
read count
