## Supplemental File 7 for "Cryptic genetic variation shapes the fate of gene duplicates in a protein interaction network"

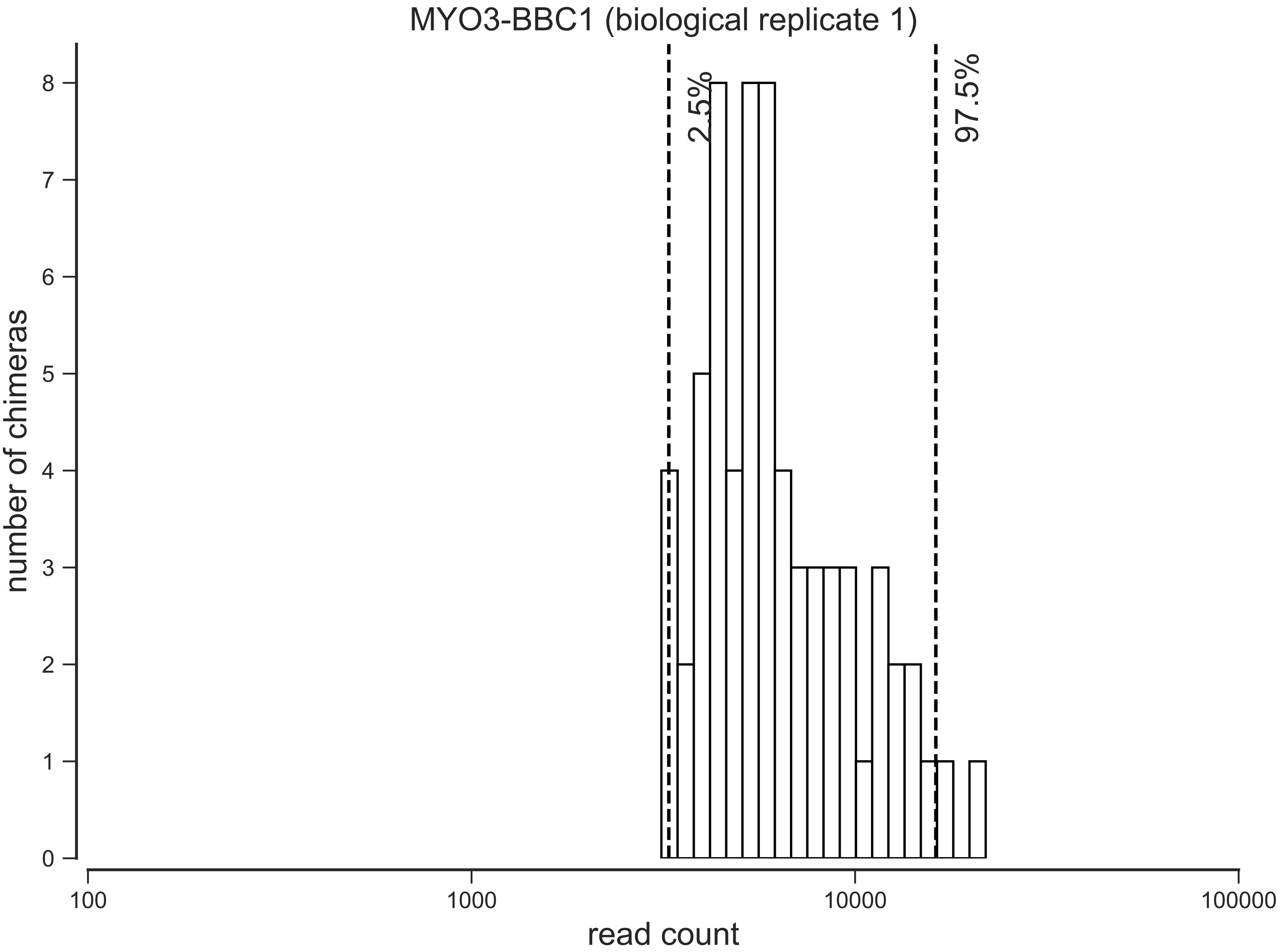

MYO3-BBC1 (biological replicate 1)
2.5%
97.5%
number of chimeras
0
1
2
3
4
5
6
7
8
100
1000
10000
100000
read count

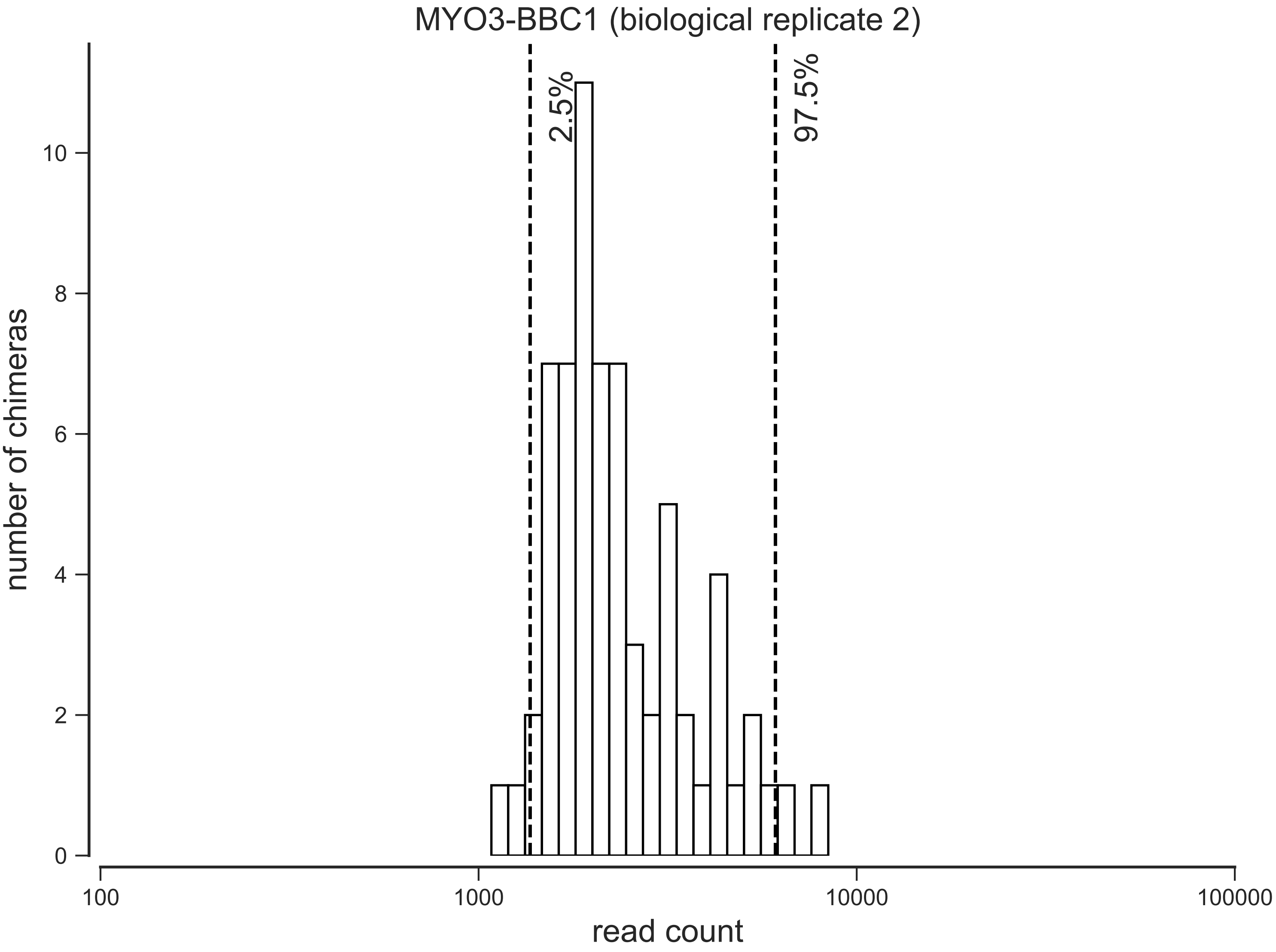

MYO3-BBC1 (biological replicate 2)
2.5%
97.5%
number of chimeras
0
2
4
6
8
10
100
1000
10000
100000
read count

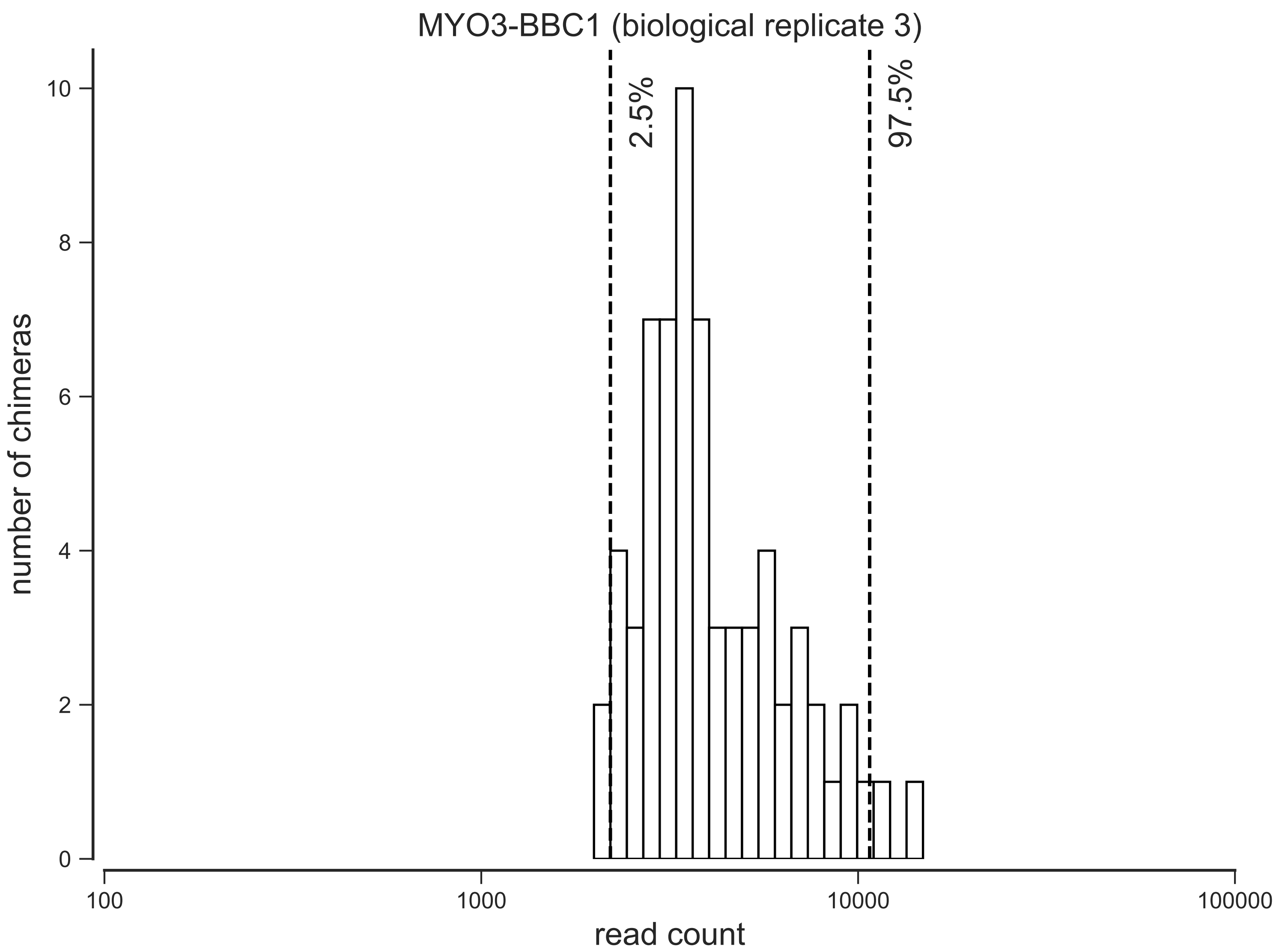
MYO3-BBC1 (biological replicate 3)
2.5%
97.5%
number of chimeras
read count
0
2
4
6
8
10
100
1000
10000
100000

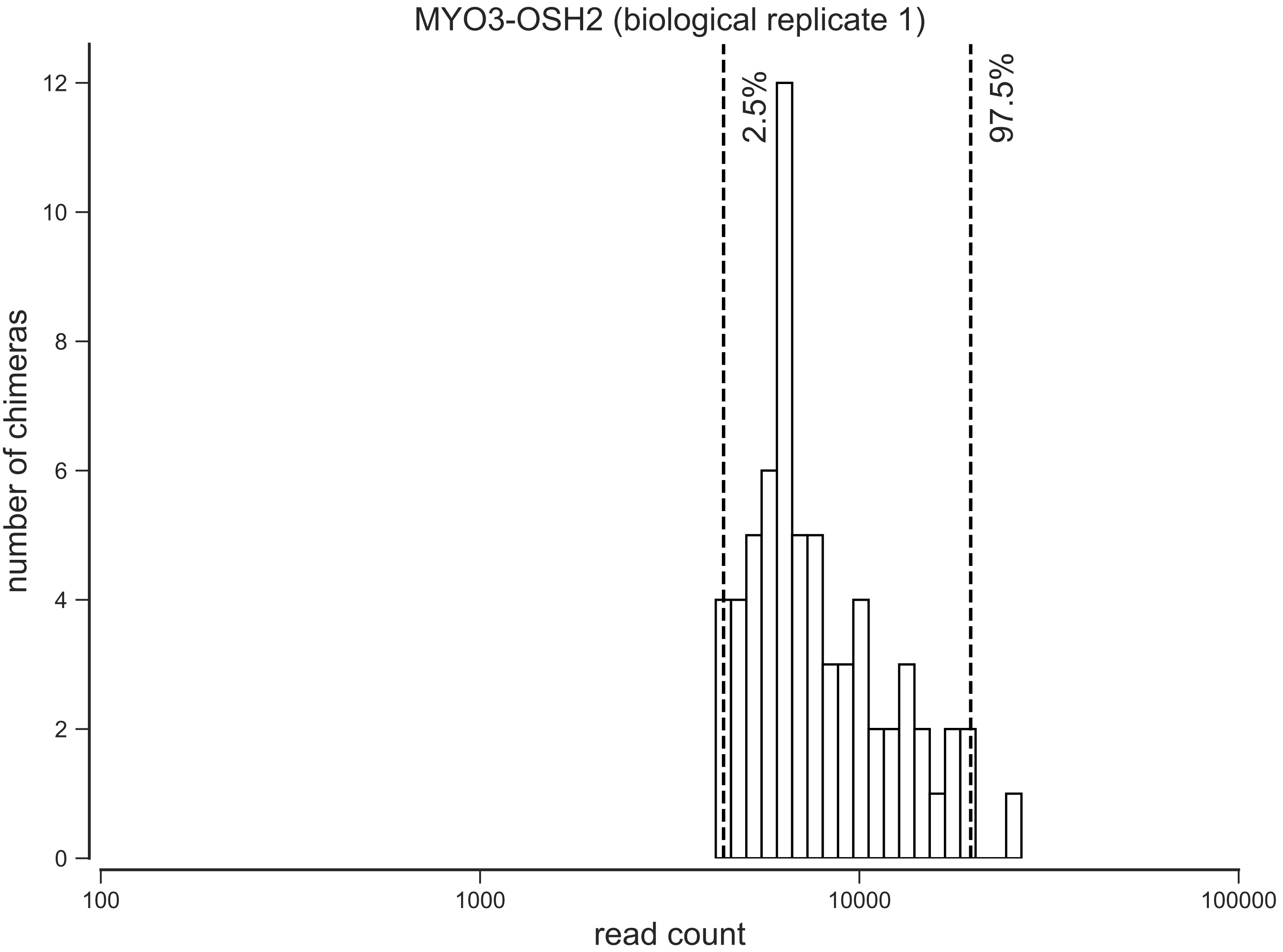

MYO3-OSH2 (biological replicate 1)
number of chimeras
read count
0
2
4
6
8
10
12
100
1000
10000
100000
2.5%
97.5%

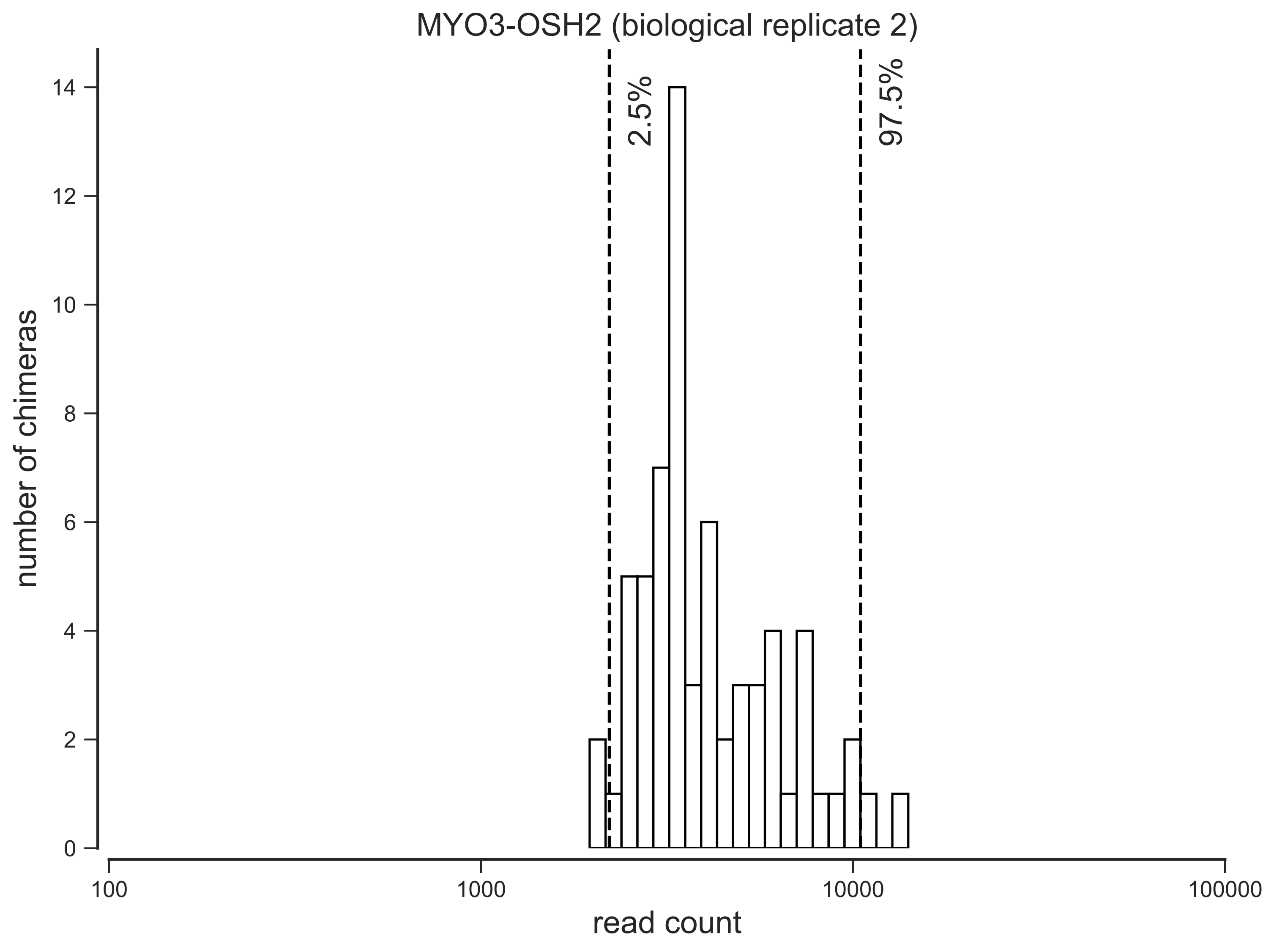

MYO3-OSH2 (biological replicate 2)
2.5%
97.5%
number of chimeras
0
2
4
6
8
10
12
14
100
1000
10000
100000
read count

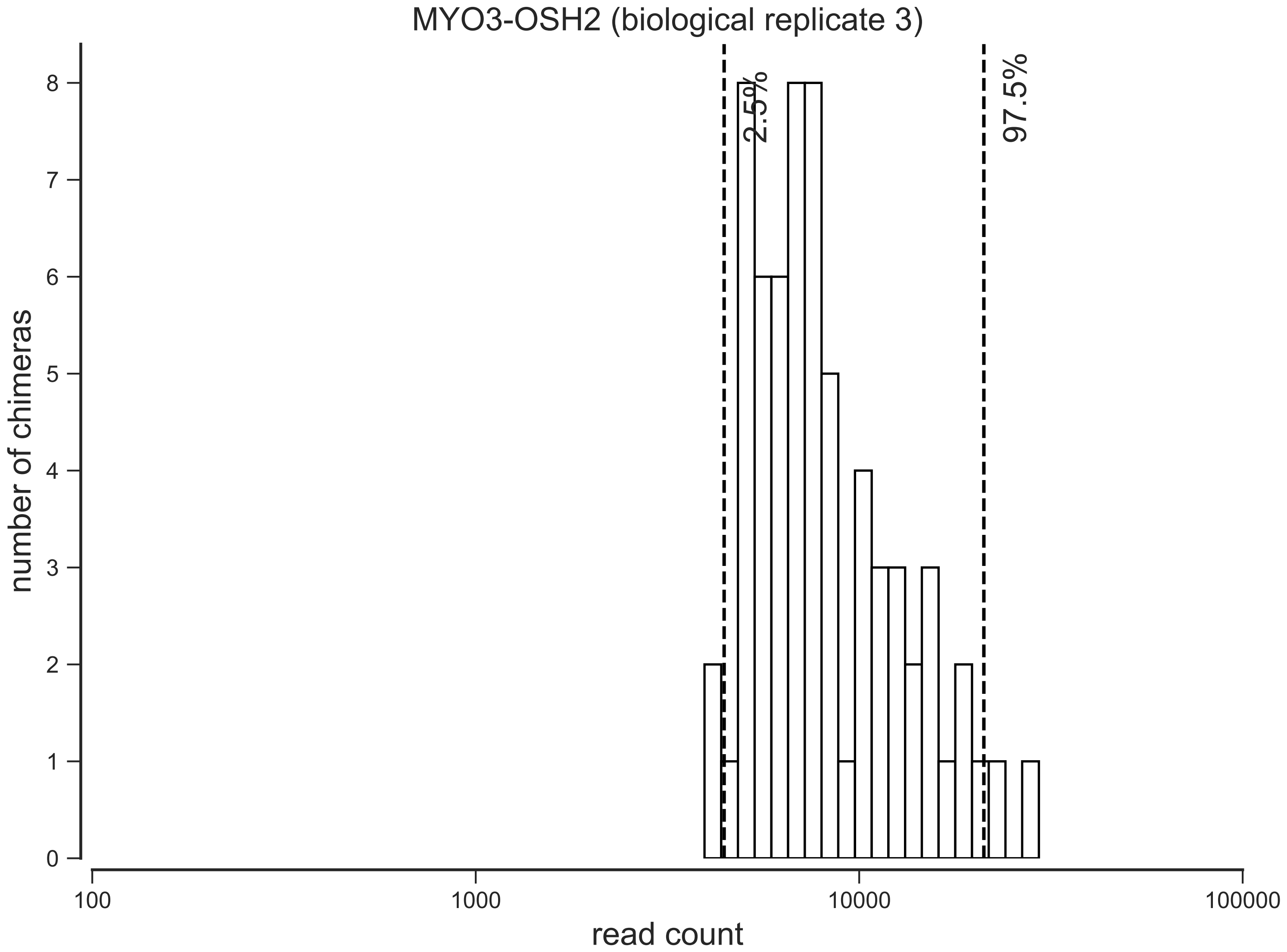

MYO3-OSH2 (biological replicate 3)
2.5%
97.5%
number of chimeras
0
1
2
3
4
5
6
7
8
100
1000
10000
100000
read count

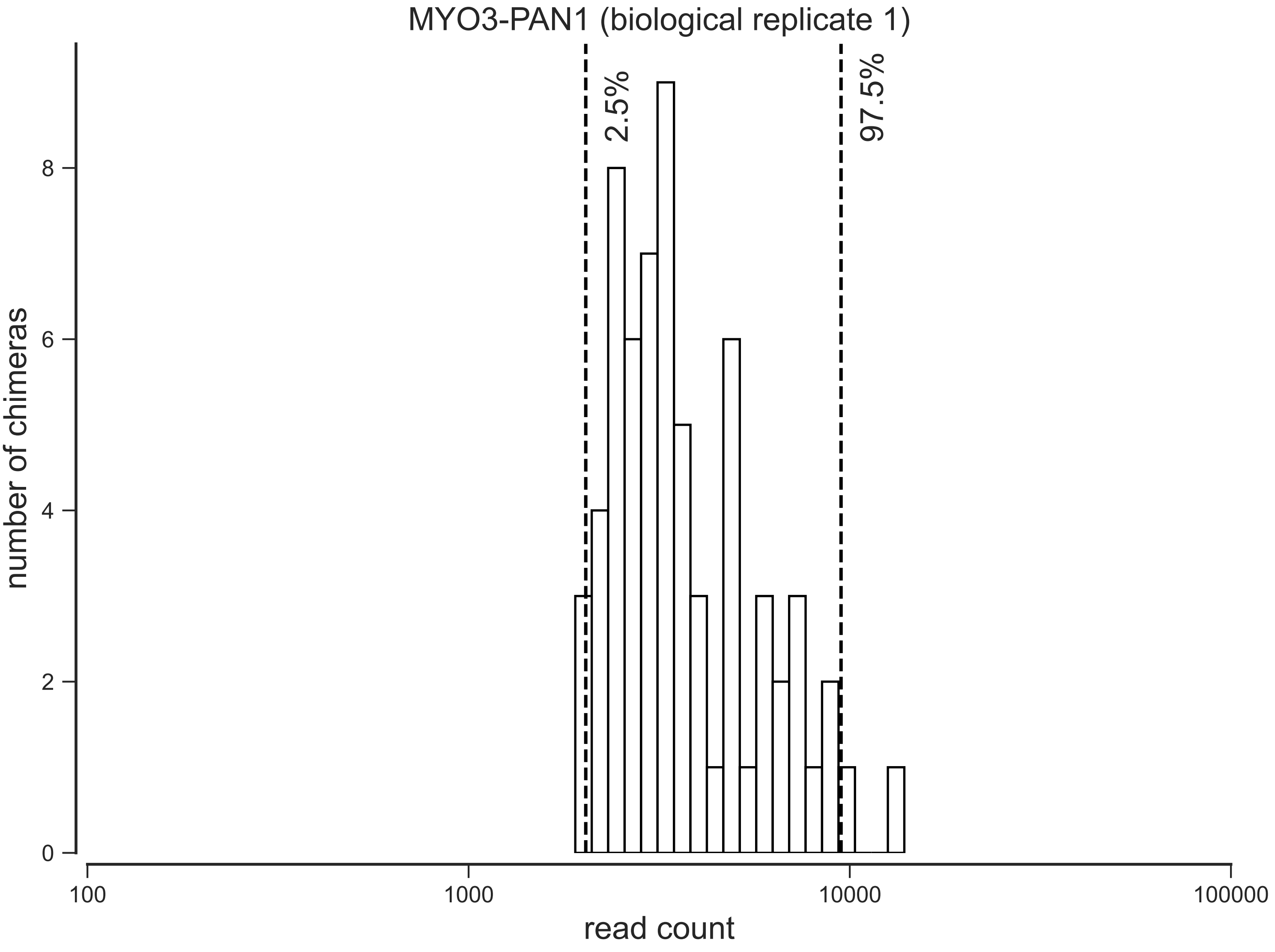
MYO3-PAN1 (biological replicate 1)
2.5%
97.5%
number of chimeras
0
2
4
6
8
100
1000
10000
100000
read count

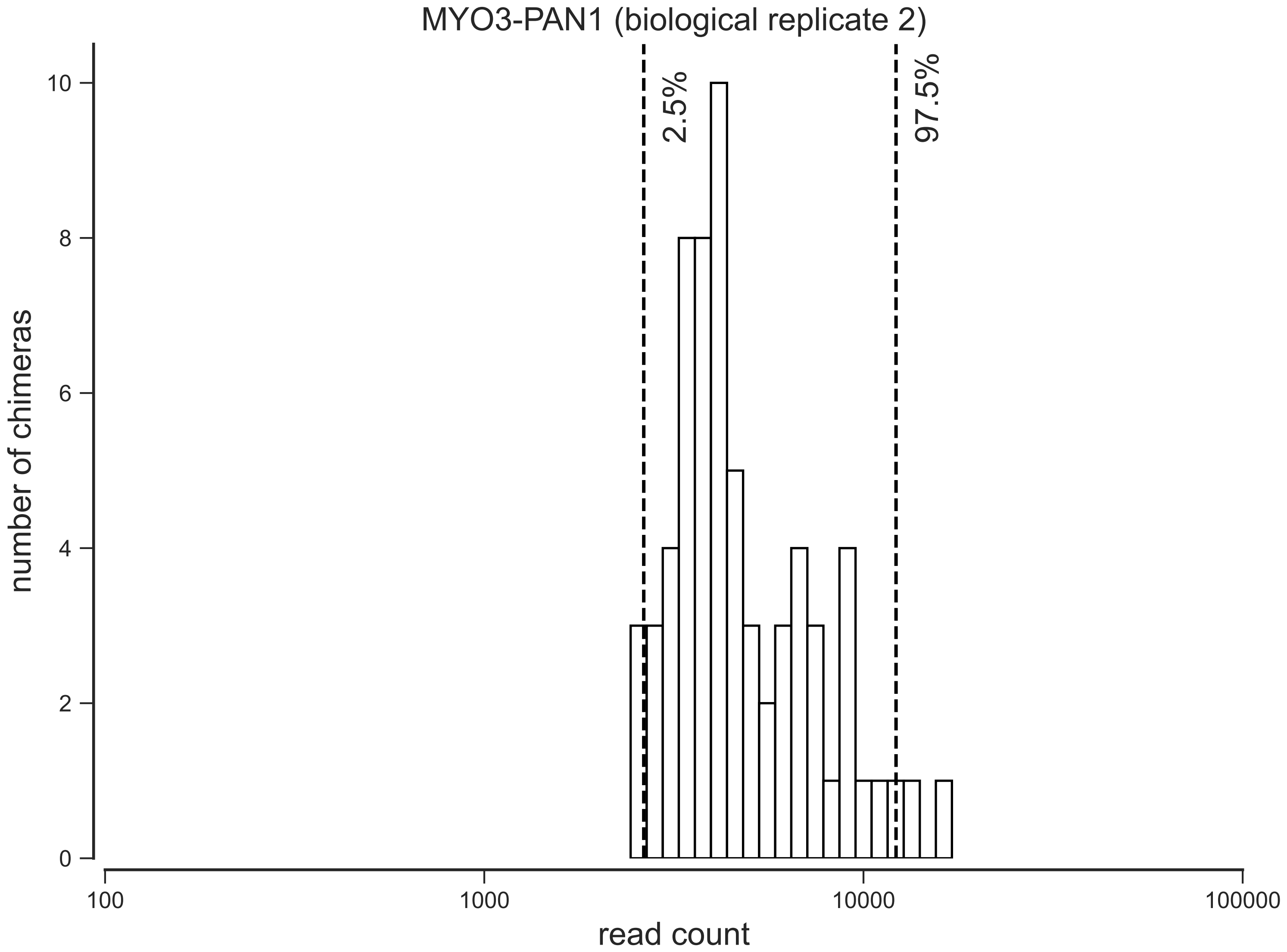
MYO3-PAN1 (biological replicate 2)
2.5%
97.5%
number of chimeras
0
2
4
6
8
10
100
1000
10000
100000
read count

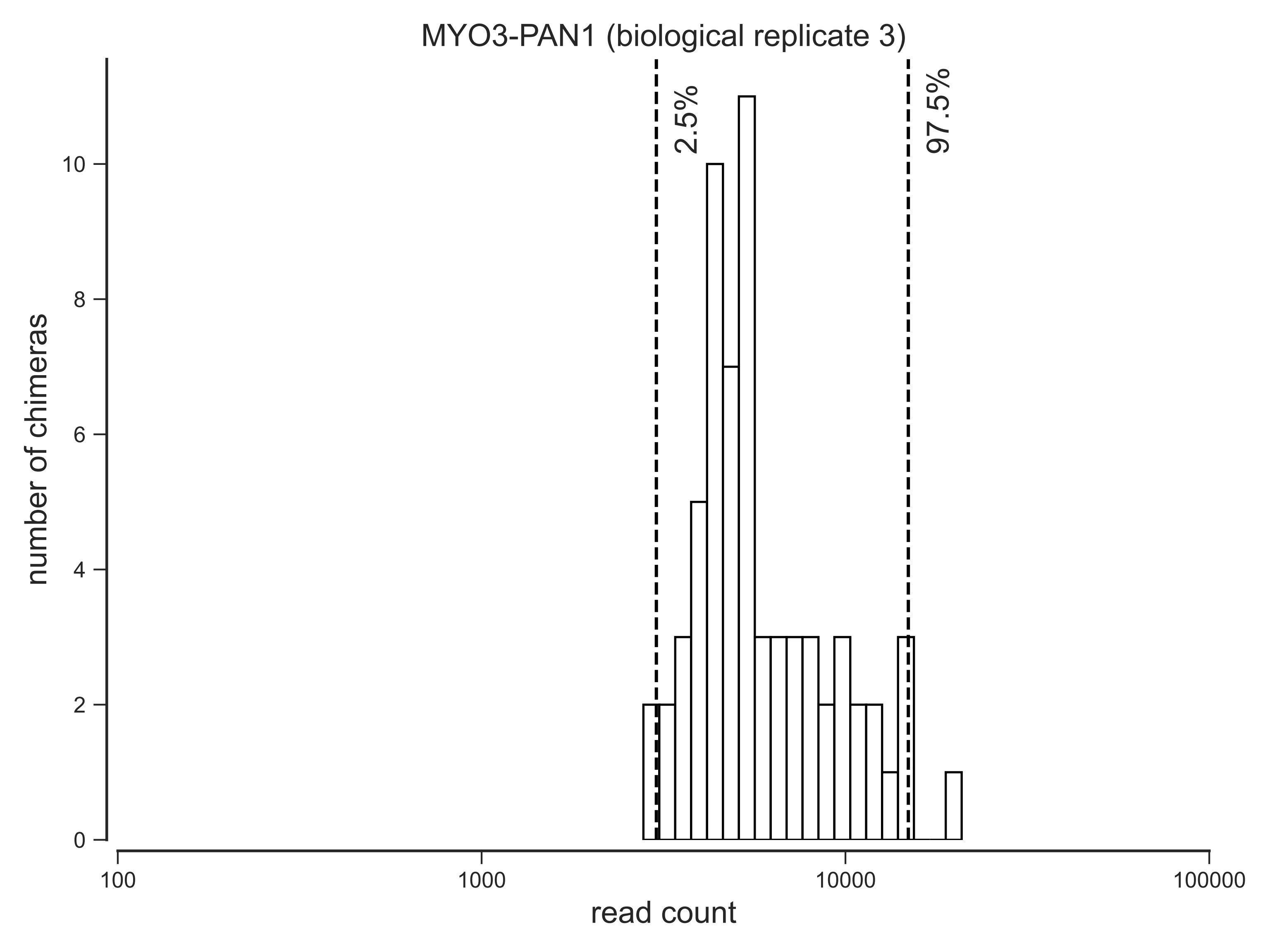
MYO3-PAN1 (biological replicate 3)
2.5%
97.5%
number of chimeras
0
2
4
6
8
10
100
1000
10000
100000
read count

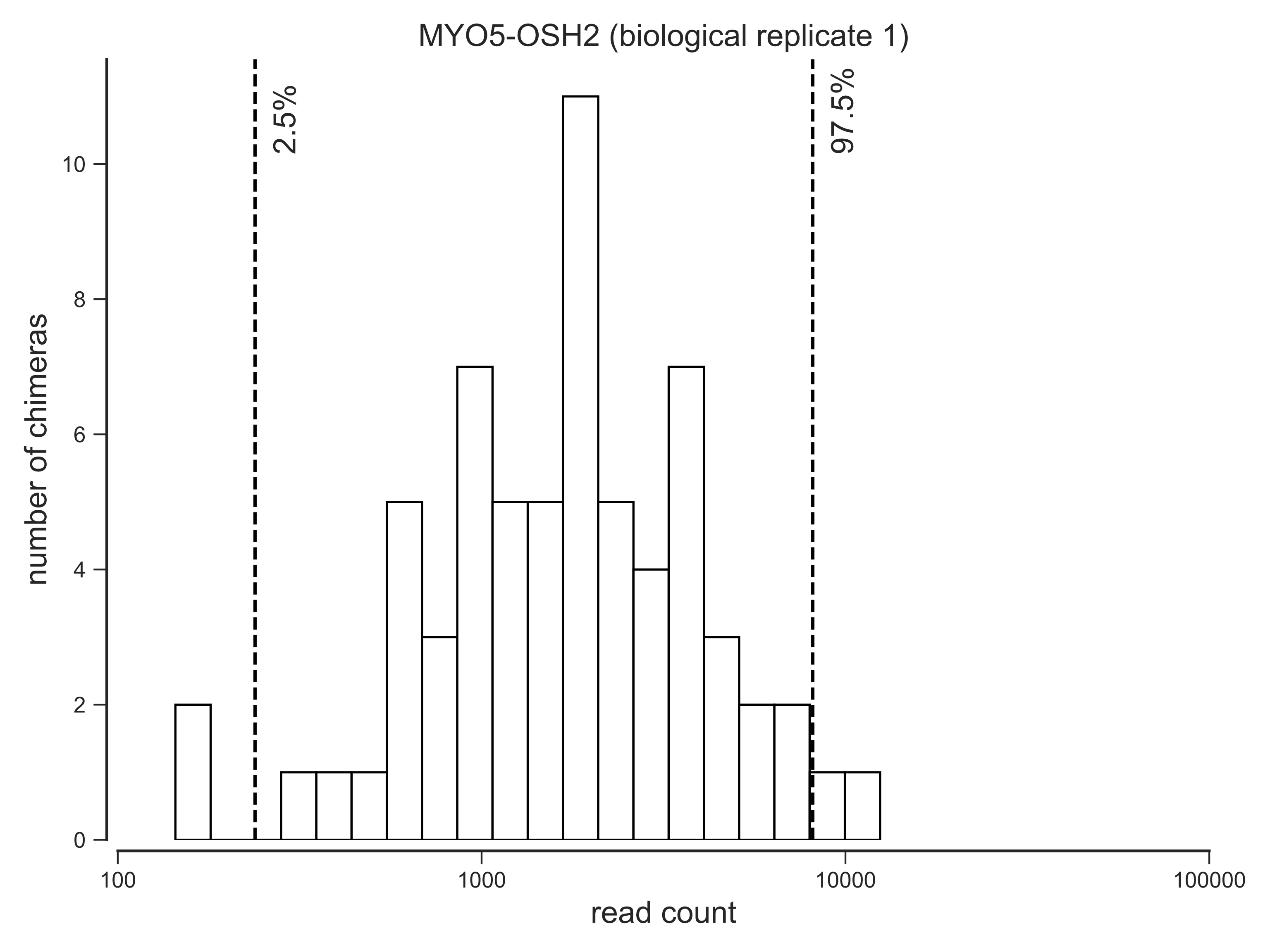

MYO5-OSH2 (biological replicate 1)
2.5%
97.5%
number of chimeras
0
2
4
6
8
10
100
1000
10000
100000
read count

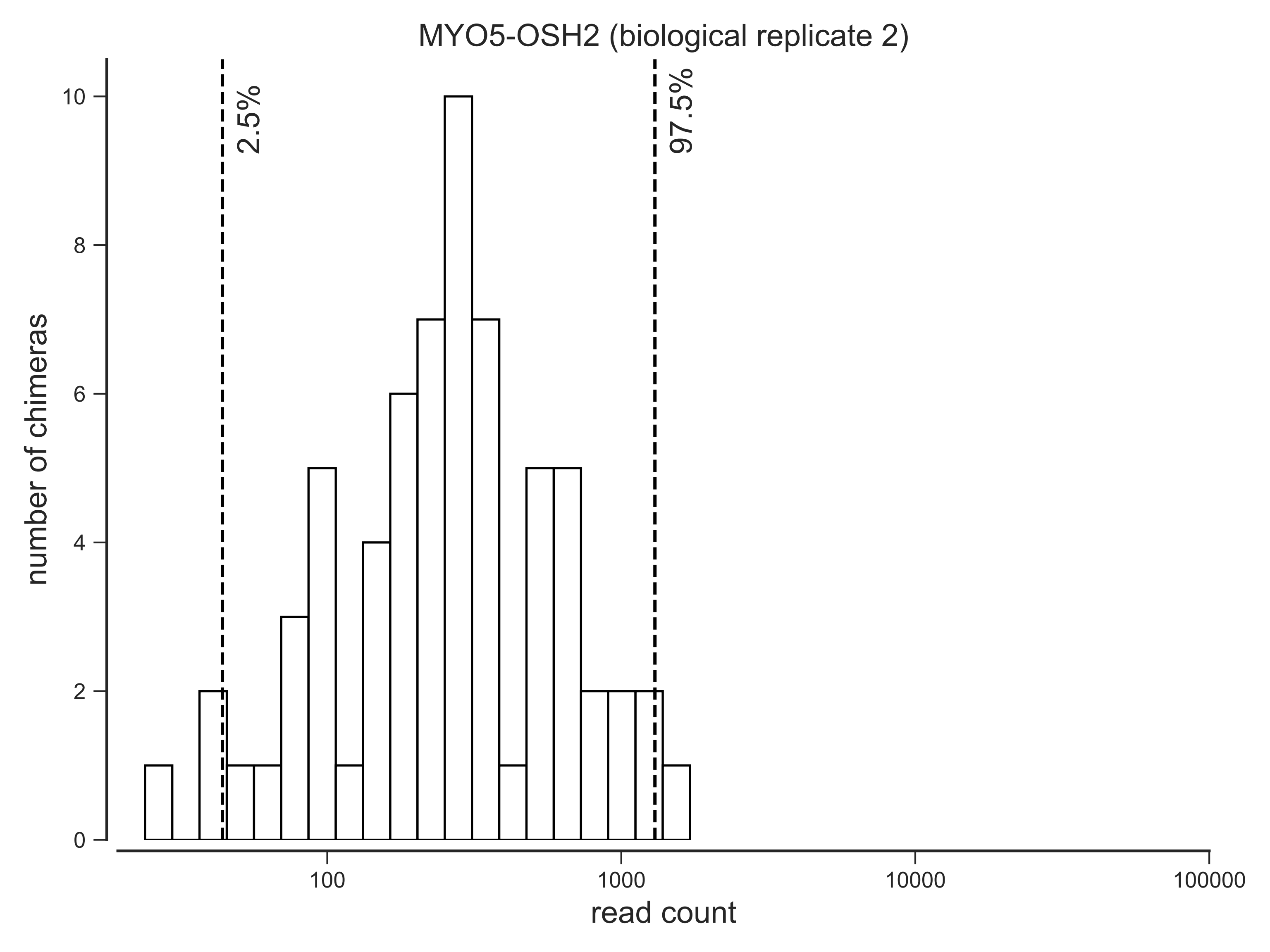
MYO5-OSH2 (biological replicate 2)
2.5%
97.5%
number of chimeras
0
2
4
6
8
10
100
1000
10000
100000
read count

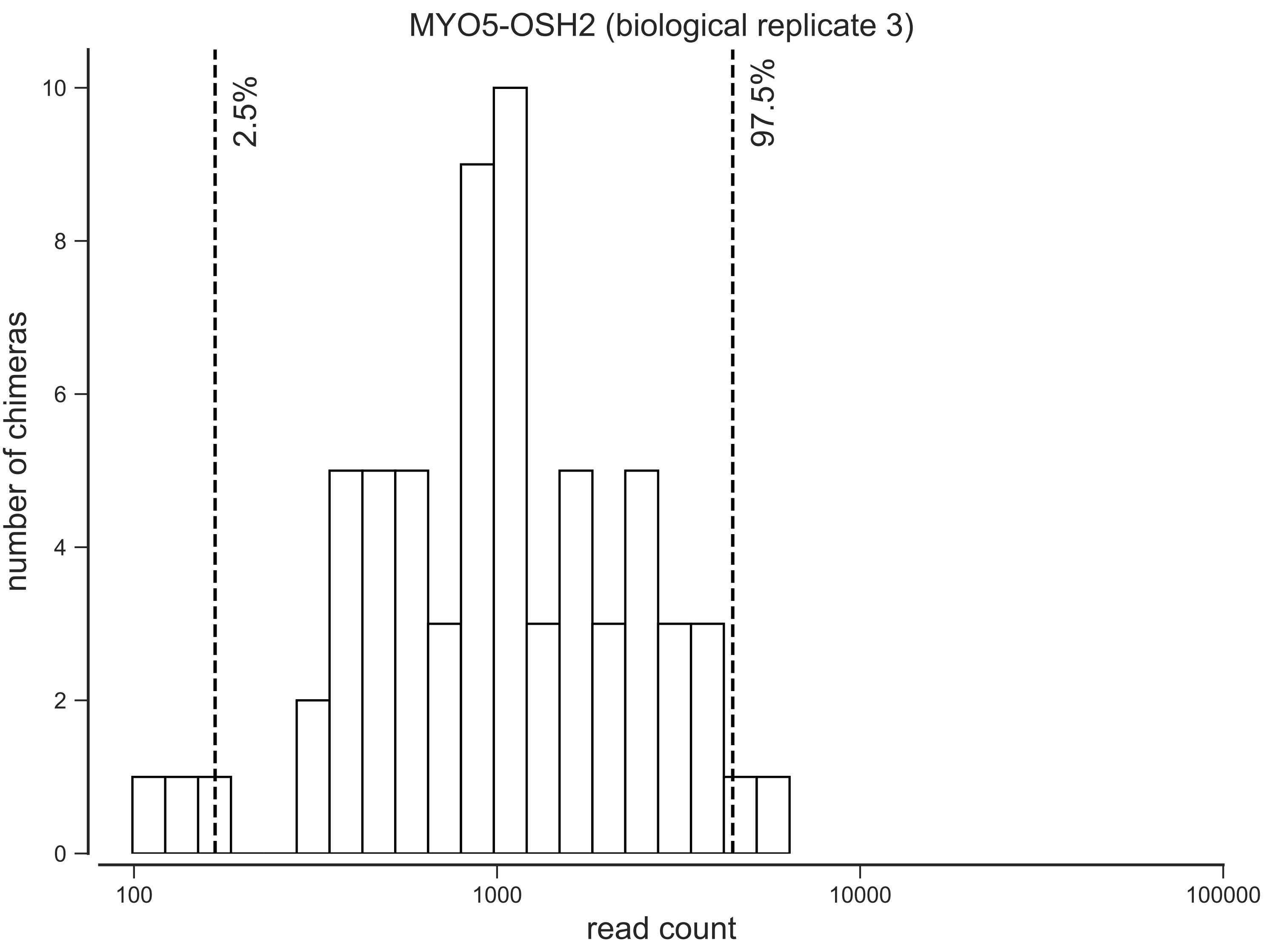
MYO5-OSH2 (biological replicate 3)
2.5%
97.5%
number of chimeras
0
2
4
6
8
10
100
1000
10000
100000
read count

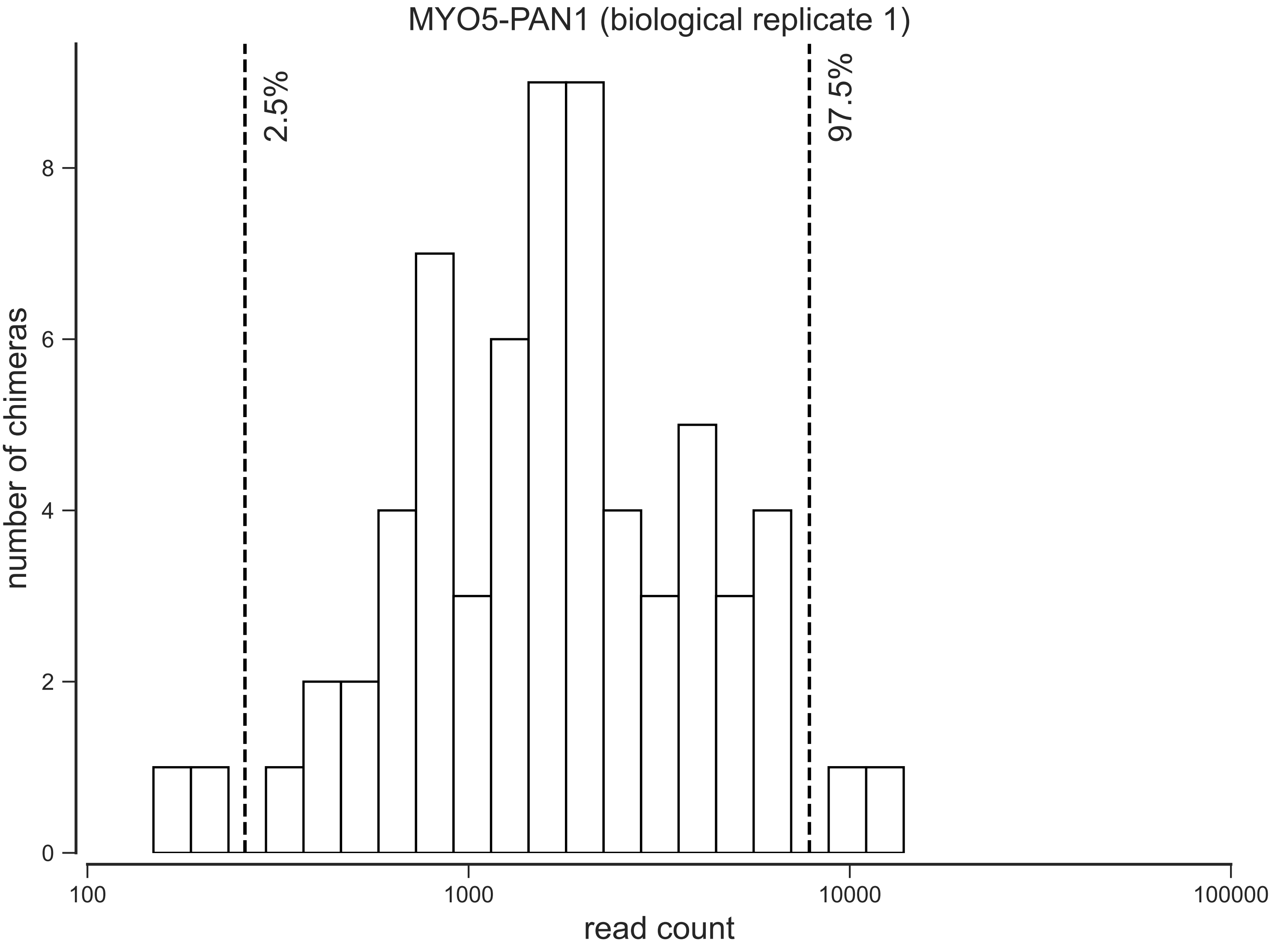
MYO5-PAN1 (biological replicate 1)
2.5%
97.5%
number of chimeras
0
2
4
6
8
100
1000
10000
100000
read count

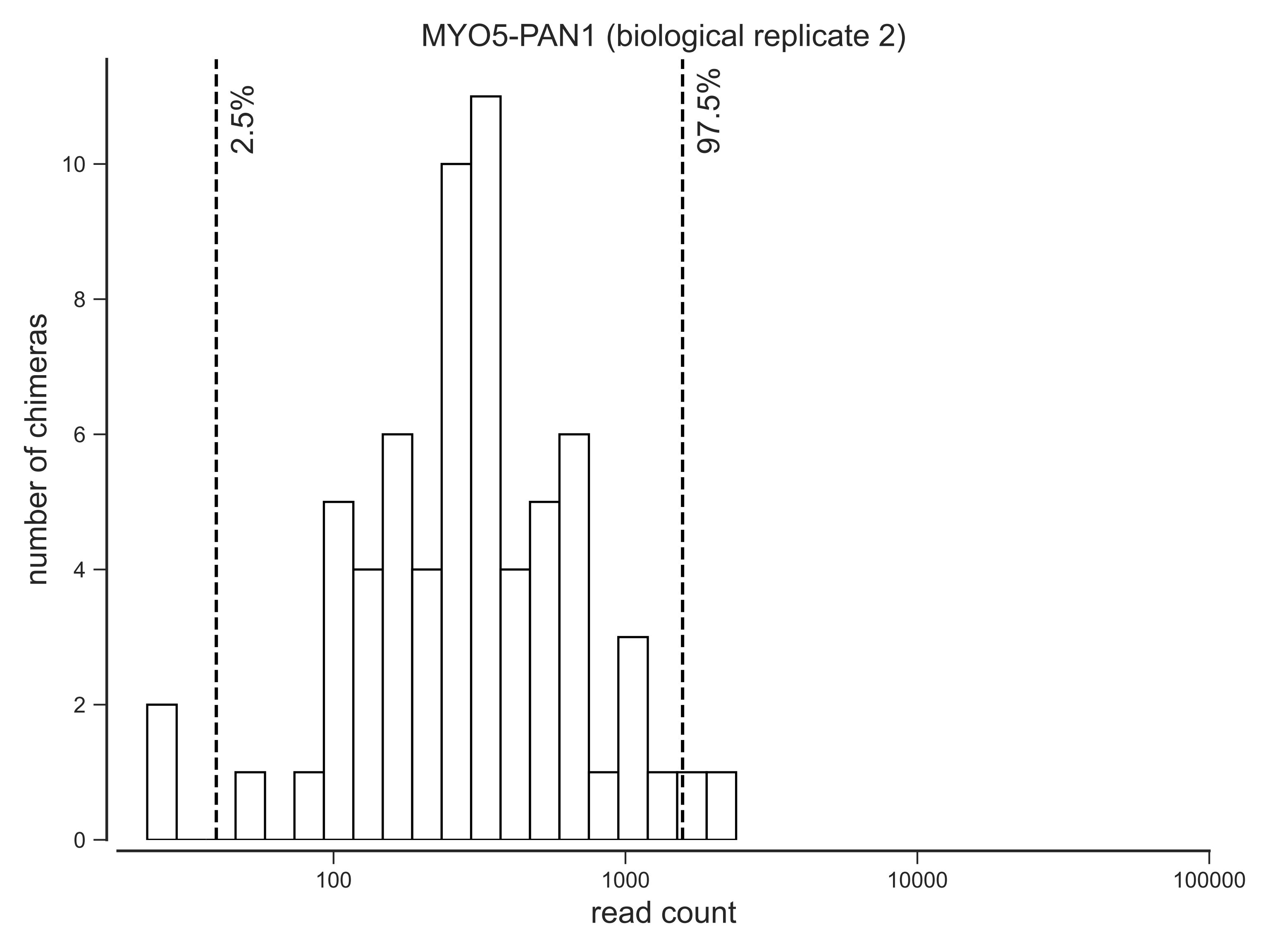

MYO5-PAN1 (biological replicate 2)
2.5%
97.5%
number of chimeras
0
2
4
6
8
10
100
1000
10000
100000
read count

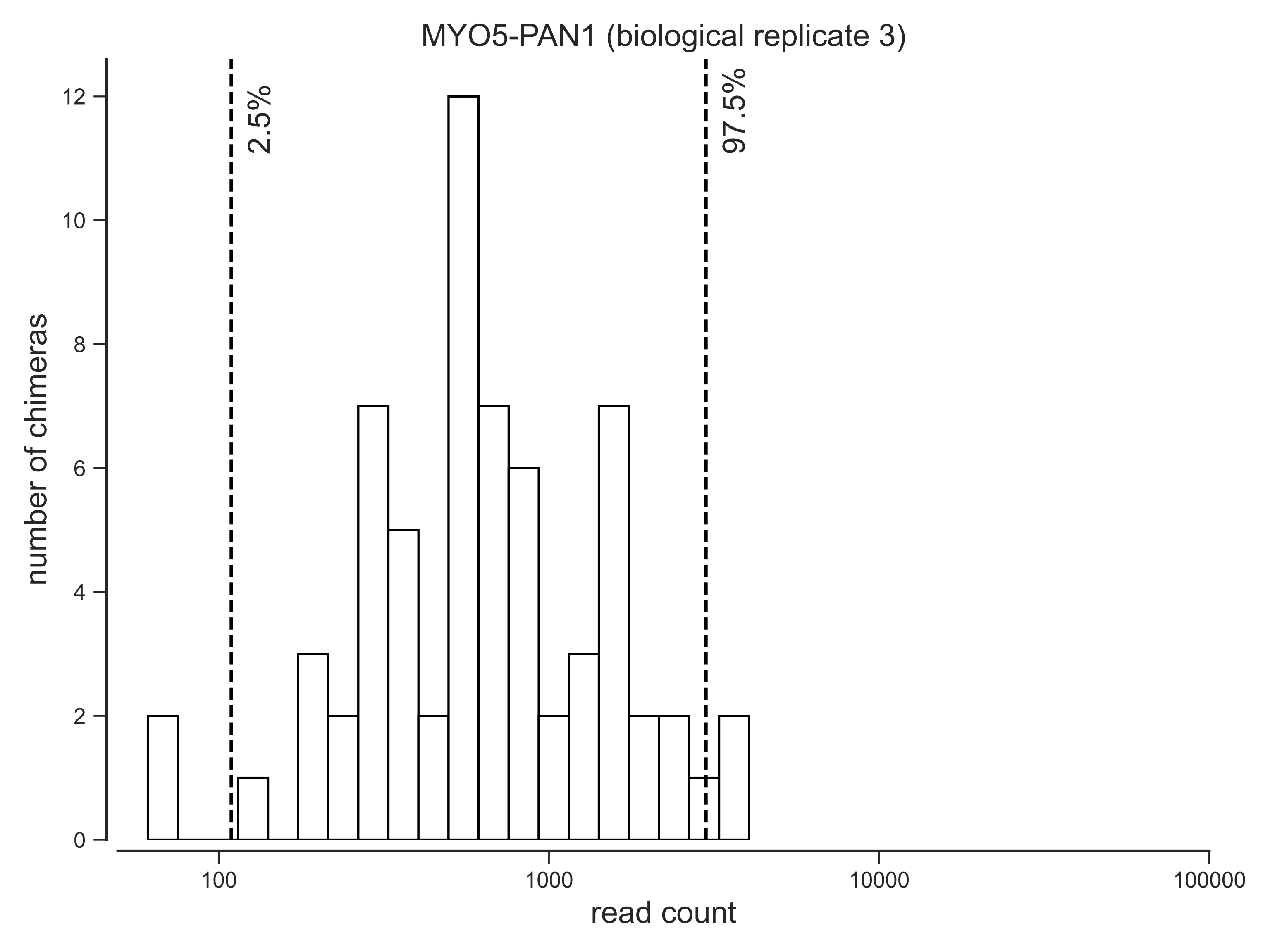
MYO5-PAN1 (biological replicate 3)
2.5%
97.5%
number of chimeras
0
2
4
6
8
10
12
100
1000
10000
100000
read count

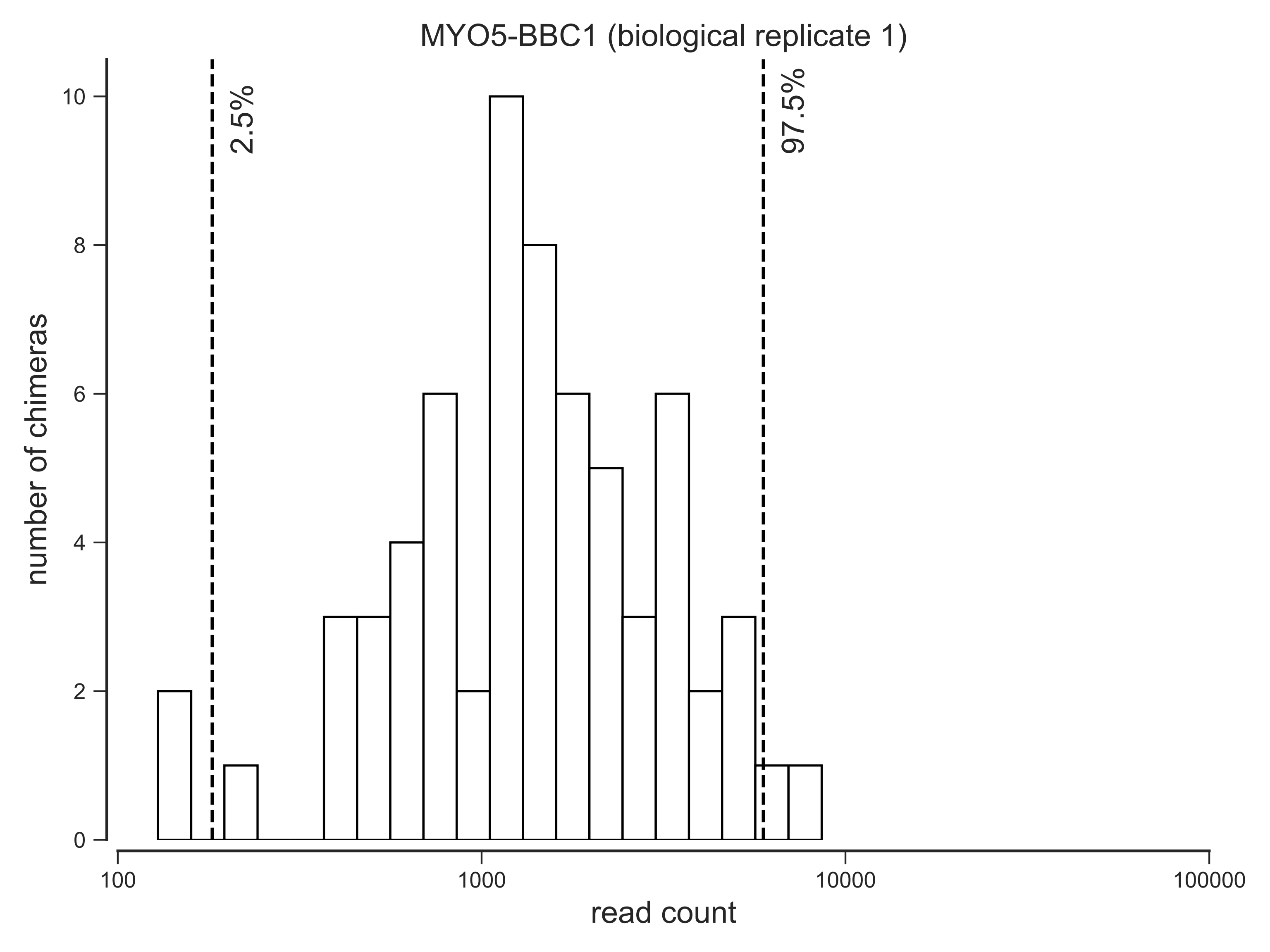
MYO5-BBC1 (biological replicate 1)
2.5%
97.5%
number of chimeras
0
2
4
6
8
10
100
1000
10000
100000
read count

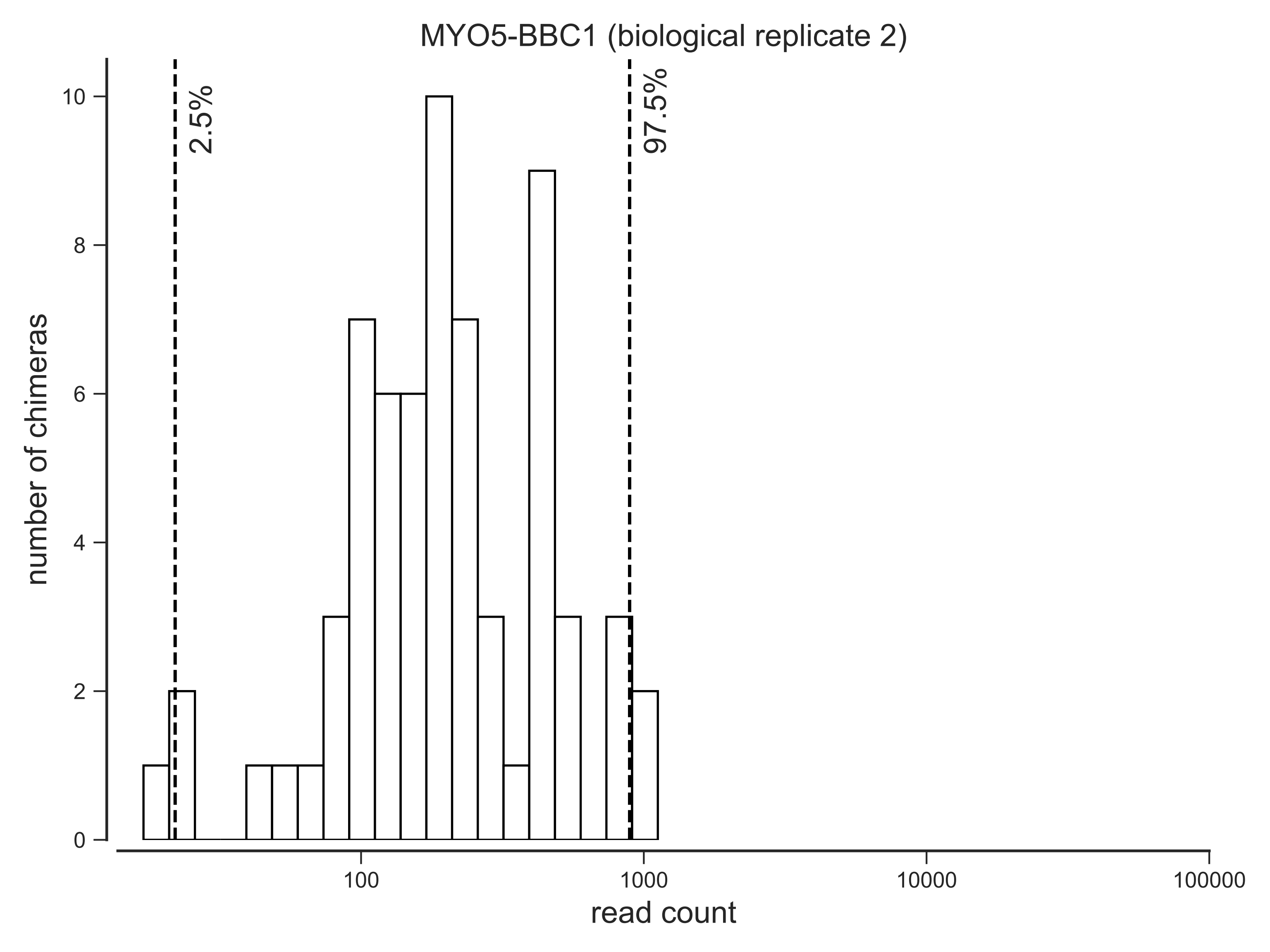
MYO5-BBC1 (biological replicate 2)
2.5%
97.5%
number of chimeras
read count
0
2
4
6
8
10
100
1000
10000
100000

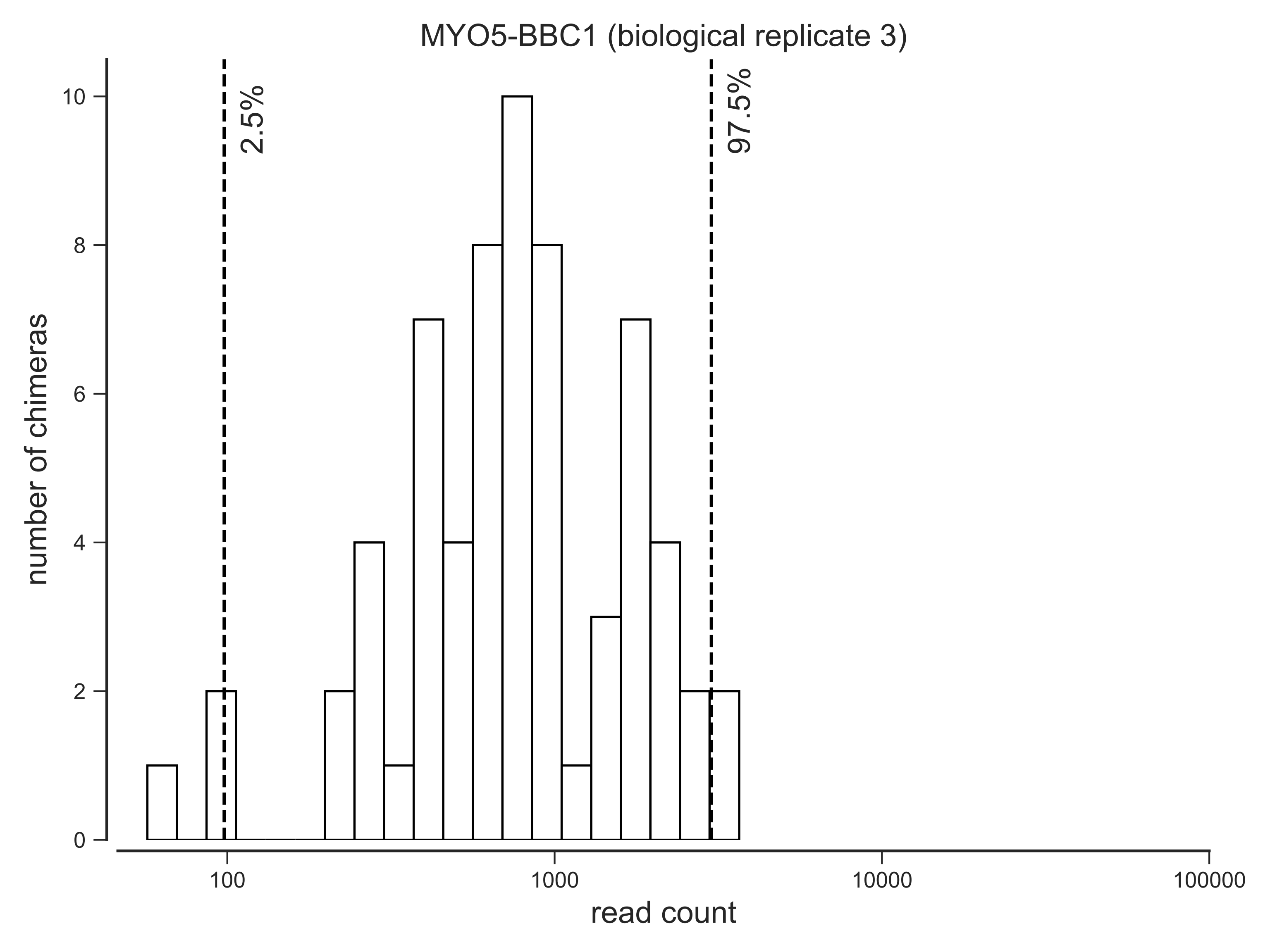
MYO5-BBC1 (biological replicate 3)
2.5%
97.5%
number of chimeras
0
2
4
6
8
10
100
1000
10000
100000
read count
